# Declining intracellular proteostasis capacity drives misfolded protein secretion in senescent human cells

**DOI:** 10.1101/2025.09.07.674107

**Authors:** Todd Watts, Harvey E. Johnston, Yasmeen Al-Mufti, Richard I. Odle, Rachel E. Hodgson, Hanneke Okkenhaug, Hayley Carr, Stephen Cranwell, Estelle Wu, Simon Walker, Simon Andrews, Tatyana Shelkovnikova, Rahul S. Samant

## Abstract

Healthy protein homeostasis (‘proteostasis’) relies on tightly-regulated protein quality-control (PQC) circuits that co-ordinate sequestration and clearance of potentially toxic aggregation-prone proteins, arising from various internal or external stress throughout an organism’s lifespan. At the protein level, proteotoxic stress responses typically involve extensive poly-ubiquitylation and sequestration of aggregation-prone proteins and PQC factors into various protective cytoplasmic and nuclear granules. However, much of our current understanding regarding this aspect of stress responses in humans stems from research in proliferating cells—despite growing evidence that stress responses vary considerably at the transcriptional level across cell proliferation states. Here, we show that the senescent cellular state—considered a major contributor to ageing-associated degeneration due to a chronic inflammatory phenotype—re-wires PQC and expels the misfolded protein load to mitigate proteotoxic stresses. Starting with a multi-dimensional transcriptomics and proteomics approach for measuring levels of total, poly-ubiquitylated, and granule-forming proteins, we have discovered a clear point of divergence between senescent and proliferating or quiescent human cell states in their responses to proteotoxic stress. Although the proteins that were poly-ubiquitylated and degraded during stress were largely conserved across states, the stress-induced sedimentation of a large number of disease-associated RNA-binding proteins (including TDP-43) was impaired only in the senescent state. Strikingly, TDP-43, as well as several other misfolded proteins, were actively secreted through the endo-lysosomal system by a diverse range of senescent cells during acute or chronic stress, through a process that requires the vesicle-associated HSP70 co-chaperone DNAJC5—an established risk factor for several neurodegenerative diseases. Misfolded protein secretion could be rescued by increasing intracellular HSP levels in ‘shallow’ but not ‘deep’ senescence, suggesting that secretion is a proteostatic adaptation that becomes less reversible over time. Our findings reveal an unappreciated aspect of the senescent-cell secretory phenotype, which may have important consequences for the non-cell-autonomous impact of senescence at the level of tissue resilience and frailty.

## INTRODUCTION

The proteostasis network is responsible for maintaining proteome fidelity under steady-state conditions.^1,2^ It must also dynamically respond under conditions of stress to ensure that the cell can resist and recover with minimal damage.^3^ A range of different external and internal cellular stresses cause an increase in protein misfolding, including heat-shock, reactive-oxygen species, UV radiation, and genetic mutation. Such ‘proteotoxic stresses’ typically stimulate multiple components of the proteostasis network to recognise misfolded species— as well as, importantly, proteins that are most prone to misfolding and/or aggregation, such as large multi-domain proteins and beta-sheet-rich proteins.^4,5^ Nascent polypeptide chains emerging from the ribosome are also included in this category.^6^ The proteins are typically post-translationally modified with one or more ubiquitin moieties (monoUb or polyUb), and sequestered into various stress-induced membraneless organelles, such as stress-granules, Q-bodies, p62-bodies, and the JUNQ/aggresome.^7,8^ In contrast with uncontrolled protein aggregation, sequestration into such structures appears to serve a cytoprotective function, not only by keeping misfolding-protein proteins away from the rest of the cellular milieu, but also by enriching them around protein quality-control (PQC) factors that prevent their irreversible aggregation and/or facilitate their processing (i.e., refolding or degradation) upon removal of the stress. Importantly, polyUb conjugation, which occurs not just on the misfolding-prone proteins but also on PQC and granule-trafficking factors themselves, appears to be important for dissolution of the granules during stress-recovery, rather than for formation of the granules.^9^ Therefore, both polyUb conjugation and granule formation are crucial components of the stress response for successful recovery.

Much of what is known about stress responses in human cells is based on experiments performed in transformed cell lines in a rapidly-proliferating state. However, cells within adult human tissues have widely-variable proliferation and turnover rates, in addition to multiple non-proliferating cell niches—e.g., terminally-differentiated, or quiescent (e.g., basal/stem-cell populations).^10,11^ Moreover, with age, cell division rates decline and tissue compositions drifts, including the accumulation of cells transitioned irreversibly to ‘senescent’ states.^12,13^ Cellular senescence (hereafter, senescence) encompasses a spectrum of states with overlapping features, of which irreversible cell-cycle exit, altered cellular morphology, and acquisition of a senescence-associated secretory phenotype (SASP) are core features. While persistence and accumulation of senescent cells is associated with frailty and chronic disease (presumed to be driven in large part by chronic inflammation via the SASP), senescence also serves important physiologic functions in tissue and organismal homeostasis.^14–18^ To take one example, the SASP serves a dual role during wound-healing—a process that declines with age—attracting immune cells for damage clearance and activating stem-cells for tissue regeneration.^19,20^ Therefore, more selective ‘senotherapeutic’ strategies to eradicate aberrant senescent cells, or to mute only the harmful components of the SASP, are highly sought-after, with only limited success, to date.^16^

Given the prominence of non-dividing cells of different types and states in human tissues, the knowledge-gap in understanding of their stress responses is surprising. Although part of this is based on limitations in experimental tractability of primary and/or non-proliferative cells, the high degree of evolutionary conservation of the proteostasis network across the tree of life^21–23^ could suggest general functional conservation of proteostasis strategies under stress. For example, given that the heat-shock response (HSR) is broadly conserved between budding yeast, nematode worms, fruit-flies, mice, and human transformed cell lines, why should we expect any differences in the heat-shock response between primary human cells in different cellular states?

Yet several lines of evidence support the notion that proteostasis networks are wired differently depending on the cell state. For example, fibroblasts made quiescent by serum-starvation or contact-inhibition upregulate lysosomal degradation pathways and rely less than proliferative cells on the ubiquitin–proteasome system for protein degradation.^24–29^ Senescent cells also appear to rely more on lysosomes for protein degradation, and have potential impairment in several transcriptional stress-response programmes, including the HSR and unfolded protein response.^30–36^ However, the impact of any divergence in proteostasis strategies between cellular states on the proteome-wide responses to stress— including at the level of polyUb conjugation and granule formation—has yet to be resolved, despite the crucial roles of such events in successful stress recovery.

Here, we set out to determine whether human fibroblast and epithelial cells in proliferating, reversibly-arrested (quiescent), and irreversibly-arrested (senescent) states respond similarly when exposed to an identical proteotoxic stress. Starting with the surprising finding that both quiescent and senescent states have dampened HSRs at the protein-level yet are more resilient in terms of their survival, we performed multi-dimensional profiling of their transcriptome, proteome, polyUb-enriched proteome, and insoluble proteome, to identify points of divergence at multiple levels of the HSR known to be crucial for successful stress recovery. Broadly, all three states responded similarly at the levels of selective protein degradation and proteome-wide increase in polyUb conjugation, albeit with mildly lowered magnitudes of response in the senescent state. By contrast, stress-induced protein sequestration into insoluble fractions was considerably dampened in the senescent state compared with the quiescent or proliferating states, especially for aggregation-prone nuclear-resident mRNA-binding proteins—despite similar levels of polyUb enrichment. Using the neurodegeneration-associated protein TDP-43 as a case study, we show that senescent cells secrete a sub-population of their aggregation-prone protein load via re-trafficking to the endosome–lysosome system. This process requires the endosome-associated chaperone DNAJC5, and can be abrogated by increasing intracellular proteostasis capacity during stress in shallow but not deep senescence. Our results highlight a novel feature of the senescent-cell secretome, with potential implications for our fundamental understanding of how senescence affects the tissue-wide proteostasis landscape, as well as opportunities for senotherapeutic intervention by curbing this misfolded protein secretion without impacting the canonical SASP.

## RESULTS

### Senescent and quiescent cells display higher proteotoxic stress resilience despite a dampened heat-shock response

Reasoning that the reported impairment in heat-shock response of senescent human cells^30,36–39^ should compromise stress resilience and recovery, we were surprised to find that fibroblasts and epithelia made senescent by various means (Supplementary Fig. S1) were actually more resilient upon heat-shock than their proliferating counterparts, as measured by cell survival after heat-shock removal (Fig. 1a−b; Supplementary Fig. S2a). This resilience was consistent for other proteotoxic stresses, e.g., pharmacologic proteasome inhibition or ER misfolding stress with tunicamycin (Fig. 1c−d, Supplementary Fig. S2b−c). We also noticed that cells made reversibly quiescent by contact inhibition were also more stress resilient in these assays, suggesting that the increased resilience was related to exit from the cell cycle, rather than a feature of senescence.^40^

**Figure 1.**
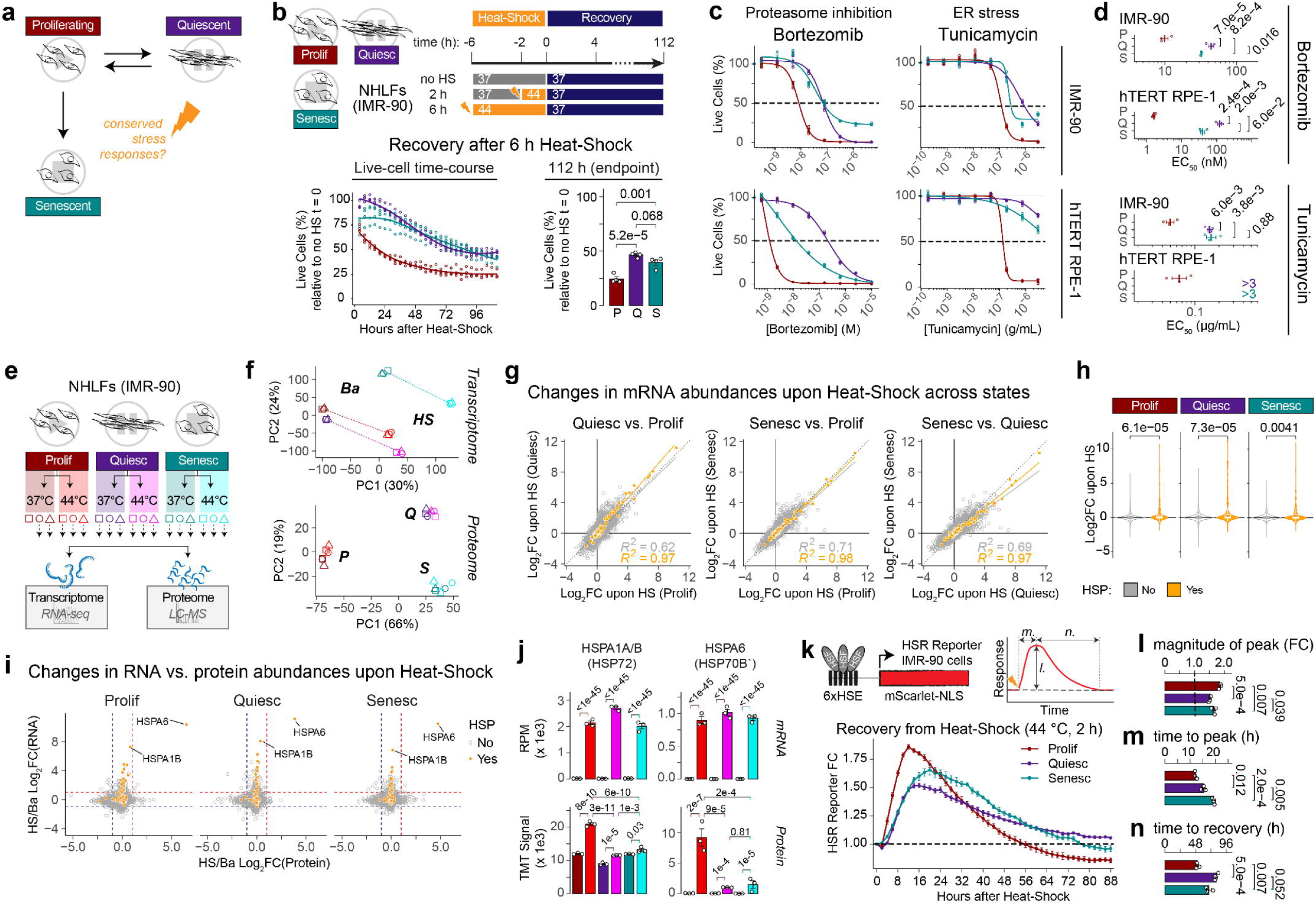
Senescent and quiescent cells display higher proteotoxic stress resilience despite a dampened heat-shock response. **a.** Central question for study: to what extent are responses to acute proteotoxic stress conserved across proliferating, reversibly-quiescent, and irreversibly-senescent cell states? **b.** IMR-90 fibroblasts expressing tdTomato–NLS nuclear marker in proliferating, contact-inhibited quiescent, or DNA-damage-induced senescent states were heat-shocked for 2 h or 6 h at 44 °C, or kept at 37 °C (‘none’), before returning to regular culture conditions (t = 0) with SYTOX Green dead-cell stain, and monitoring every 4 h on an Incucyte SX5 Live-Cell Analysis System. Number of live cells were calculated at each time-point by subtracting SYTOX+ from tdTomato+ nuclei counts, both quantified using the StarDist plugin in ImageJ. Differences in percentage of surviving cells between each state per heat-shock condition at the end of the experiment are shown as a bar-chart. See also Supplementary Fig. S2a. **c–d.** IMR-90 fibroblasts or RPE-1 epithelia in proliferating, contact-inhibited quiescent, or DNA-damage-induced senescent states were treated with a range of concentrations bortezomib or tunicamycin, or DMSO vehicle-treated control, for 72 h, before staining with Hoechst 33342 and SYTOX Orange dead-cell stain, and imaging immediately on an ImageXpress Confocal HT.ai High-Content Imaging System. Number of live cells was calculated as in (b), and normalised as a percentage of the vehicle control for that cell state at the end of the experiment. Dose-response curves and EC_50_ values were calculated using the drc R package. See also Supplementary Fig. S2b–c. **e.** Experimental design for total transcriptome and proteome quantification of IMR-90 fibroblasts. Cells in proliferating, contact-inhibited quiescent, or DNA-damage-induced senescent states were exposed to heat-shock of 44 °C (HS), or kept at 37 °C (Basal, Ba) for 2 h. Samples were harvested immediately, separated in two, and snap-frozen. This process was repeated to obtain three individual biological replicates. The resultant samples (n = 18) were processed together for RNA-sequencing-based transcriptomics or mass-spectrometry-based bottom-up proteomics. **f.** Raw component scores from Principal Component Analysis (PCA) of the top two PCs for each experiment suggest differences in sample clustering between transcriptome and proteome data. Shapes represent individual replicates, and colours represent each state and treatment, as shown in (e). Percentage of variance explained by each PC is indicated in the axis titles. **g.** Heat-shock-induced Log_2_FC values across each cell state for the transcriptome suggests high degree of correlation between states for HSPs (orange) compared with the rest of the transcriptome (grey). For each pairwise comparison, linear regression lines for y ∼ x, and squared Pearson’s correlation coefficients, for HSPs vs. the rest of the transcriptome, are shown. Dotted lines represent y = x. **h.** HSPs have slightly muted induction at the transcript level upon heat-shock in senescent vs. non-senescent states. Global distribution of Log_2_FC values (from DESeq2 differential analysis) upon heat-shock for HSPs (orange) vs. the rest of the transcriptome (grey) in each state are plotted as violin and overlaid boxplots (median + inter-quartile range). Adjusted p-values from non-parametric Kruskal-Wallis test (two-tailed, unpaired) are shown. **i.** HS-induced Log_2_FC values across each cell state for the transcriptome (y-axis) and proteome (x-axis), suggests a low degree of correlation between the two dimensions. Log2FC values were calculated by DESeq2 and DEqMS for transcriptome and proteome dimensions, respectively. HSPs (orange) are highlighted against the rest of the transcriptome (grey). Dashed lines represent Log_2_FC = 1 (red) or Log_2_FC = –1 (blue). **j.** Muted induction of HSP70 chaperones at the protein but not transcript level in both quiescent and senescent states. Global-normalised RPM (transcriptome) or TMT abundances (proteome) for the HSP70s HSPA1A/B (HSP72) or HSPA6 (HSP70B’) are plotted as bar-charts, and adjusted p-values from DESeq2 (transcriptome) or DEqMS (proteome) are shown. **k–n.** Fluorescent reporter suggests muted heat-shock-response (HSR) induction and delayed recovery in quiescent and senescent states. 6xHSEp::mScarlet**–**NLS IMR-90 cells were exposed to the same HS, recovery, and Incucyte imaging conditions as in (b), with the exception of no SYTOX addition, and imaging every 2 rather than 4 hours. Reporter fluorescence intensity fold-change (FC) is based on the integrated signal per well at each time-point vs. t = 0, calculated after surface-fit background subtraction and confluency adjustment. Differences in magnitude of highest FC (l), time taken to reach highest FC (m), and time taken to recover to FC = 1.07 (the value attained by the quiescent state at the end of the experiment) (n), between each state are shown as bar-charts. For b–d, j–n: Open-circle data-points represent individual biological replicates (n = 3/4), and filled-circles or bars +/– error-bars represent mean +/– standard-error. Adjusted p-values from Tukey’s HSD post-hoc test following significant (p < 0.05) one-way ANOVA are shown. For d: ns, P > 0.05; *, p < 0.05; **, p < 0.01; ***, p < 0.001; ****, p < 0.0001.

To determine whether this increased stress resilience in non-proliferative states could be attributed to differences in induction of the cytoprotective HSR at the molecular level, we performed bulk transcriptomics and proteomics of fibroblasts in the three states, with and without heat-shock (Fig. 1e; Supplementary Fig. S3; Tables S1−S3). Despite clear separation between the basal transcriptomes for the three cell states (Fig. 1f, Supplementary Fig. S3a), the mRNA fold-changes upon heat-shock were remarkably similar—especially for HSPs (Fig. 1f–g). The degree of HSP induction, taken as a group, was marginally dampened in the senescent state, although not to the same extent as in a previous transcriptomics study in replicatively-senescent fibroblasts (Fig. 1h, Supplementary Fig. S3b).^30^ Nevertheless, there was a strong correlation in the HSP distributions between the two studies (Supplementary Fig. S3c).

Comparing the transcriptomics with the proteomics data, the typically modest correlation between RNA and protein abundance^41–43^ was notably higher for HSPs (Supplementary Fig. S3d). Despite this, and in contrast with the increase in a substantial proportion of HSPs at the RNA level, only the highly stress-inducible 70 kDa HSP family member HSPA6/HSP70BL showed a clear increase upon heat-shock at the protein level (Fig. 1i, Supplementary Fig. S3e)—and even these seemed slightly delayed in the senescent state in a time-course (Supplementary Fig. S3f).^44^ In addition to the well-established post-transcriptional and post-translational regulation of stress responses,^9,45–48^ we attributed this discrepancy to the signal-delay between mRNA transcription and protein translation captured in the 2-hour heat-shock window, as well as the global arrest in protein translation at such elevated temperatures.

Moreover, there was a clear dampening of the induction of HSPA1/HSP72 and HSPA6/HSP70BL in both quiescent and senescent cells, despite similar induction at the RNA level (Fig. 1j). Such a disconnect between transcriptional and translational/post-translational induction of the heat-shock-response with senescence is strikingly similar to a previous study in replicatively-senescent human mesenchymal stem cells.^36^

Monitoring dynamics of the heat-shock response (HSR) immediately after a sub-lethal 2-hour heat-shock using a fluorescent 6xHSEp::mScarlet reporter IMR-90 line indicated that quiescent and senescent states displayed a reduction in the magnitude of HSR induction, delay in the time taken to achieve the maximum HSR, and slower recovery to baseline HSR levels, relative to the proliferative state (Fig. 1k–n). Our data therefore suggests that non-proliferative cell states display dampened production of cytoprotective HSPs upon heat-shock, yet are surprisingly more tolerant to this and other proteotoxic stressors.

### Stress-induced protein poly-ubiquitylation and degradation are broadly conserved across cell states

Except for impaired protein-level induction of HSPA1/HSP72 and HSPA6/HSP70BL in both quiescent and senescent states upon heat-shock, the global heat-shock–modulated proteome distributions were remarkably similar across the three states (Fig. 2a, Supplementary Fig. S3e, S4a–c). Statistical analysis to identify differential proteins (DPs) across any of the experimental variables, and subsequent clustering based on similarities in DP abundance profiles, mostly identified proteins with different baseline abundances between the three cell states, with only a few clusters representing proteins that changed between basal and heat-shock conditions (Supplementary Fig. S4a–c; Table S4). Over-representation analysis (ORA) identified clear clusters capturing the expected protein-level differences between the three states, including increases in proteins of the extracellular matrix in contact-inhibited quiescence (Clusters 1 and 6),^49,50^ lysosomes in both quiescence and senescence (Cluster 4),^25,51,52^ and sterol biosynthesis pathways in senescence (Supplementary Fig. S4d; Table S4).^53^

**Figure 2.**
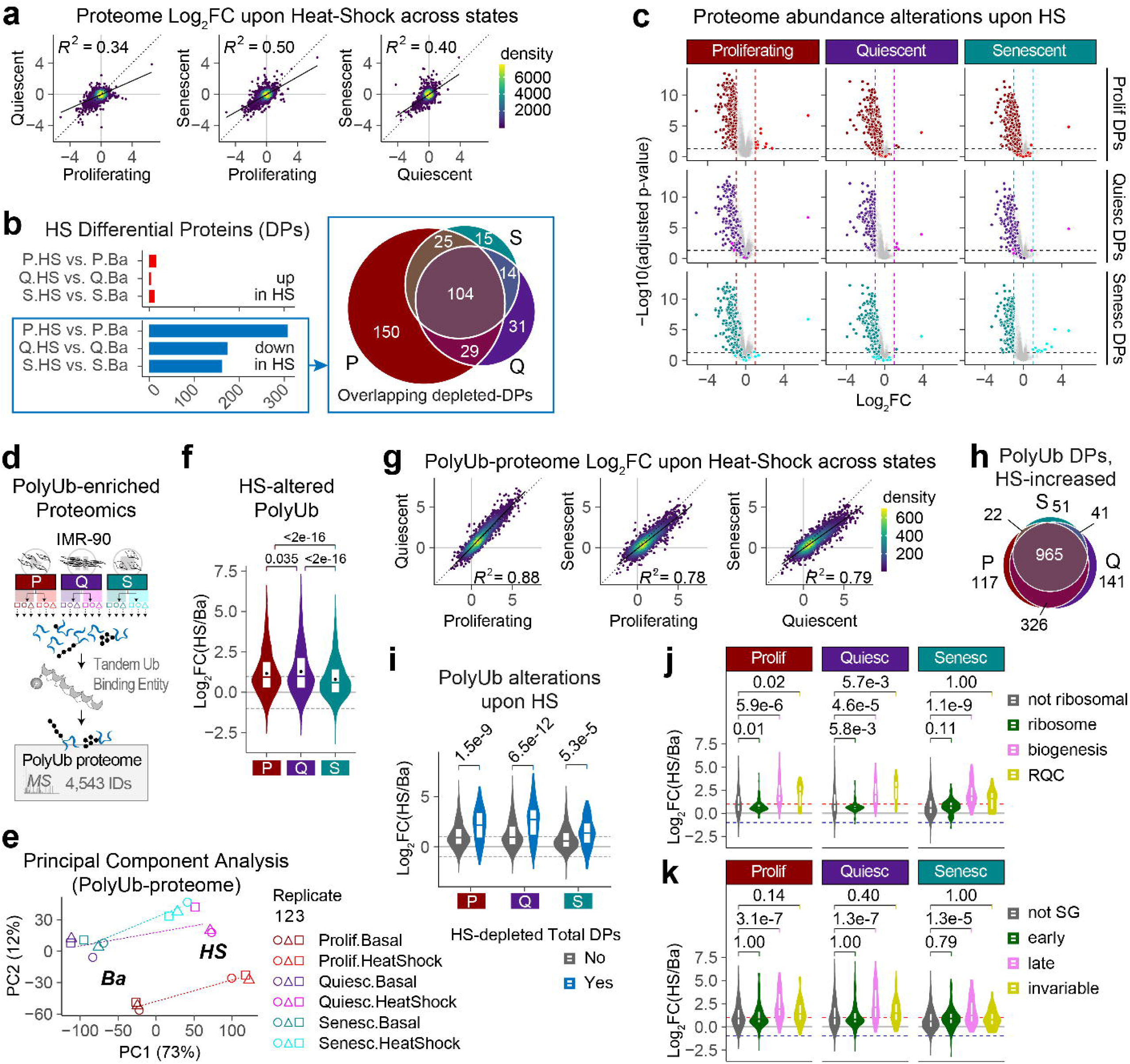
Stress-induced protein poly-ubiquitylation and degradation are broadly conserved across cell states. **a. & g.** Heat-shock-induced Log_2_FC values across each cell state for the total (A) or polyUb-enriched (G) proteomes. For each pairwise comparison, linear regression lines for y ∼ x, and squared Pearson’s correlation coefficients, are shown. Dotted lines represent y = x. **b.** Left, Total numbers of differential proteins (DPs) based on DEqMS-calculated cut-offs of adjusted p < 0.05 and Log2FC >= 1 for increased (up, red) or <= −1 for decreased (down, blue), upon heat-shock (HS) vs. basal (Ba) conditions within proliferating (P), quiescent (Q), or senescent (S) states. Right, Venn diagram showing overlap between heat-shock-depleted proteins across the cellular states. **c.** Volcano plots showing heat-shock-induced total protein-level changes in proliferating, quiescent, and senescent states. The values for Differential Proteins from proliferating (reds), quiescent (purples), or senescent (turquoises) states are plotted onto the other two states, to determine whether direction of changes are conserved across states. Absolute Log2FC = 1, and adjusted p = 0.05, are represented by dashed vertical and horizontal lines, respectively. **d.** Experimental design for tandem-ubiquitin-binding-entity (TUBE)–based poly-ubiquitylated proteome (polyUb-proteome) enrichment and quantification. **e.** Raw component scores from Principal Component Analysis of the top two PCs for the polyUb-proteome suggests clustering of samples first by condition (basal vs. heat-shock), and then by cell state. Percentage of variance explained by each PC is indicated in the axis titles. **f.** Distributions of HS-induced alterations to the polyUb-proteome suggests extensive increase in protein poly-ubiquitylation across all cell states. Black circle represents the mean Log_2_FC value for each distribution. **g.** Venn diagram showing overlap between proteins increased upon heat-shock in the polyUb-proteome across cell states. **h.** Heat-shock-depleted proteins at the total proteome level (blue) have higher polyUb-enrichment upon heat-shock than the rest of the proteome (grey) in all cell states **j–k.** Distribution of heat-shock-induced alterations of ribosome–^154^ (j) or stress-granule–^155^ **a.** (k) annotated proteins in the polyUb–proteome For f, i–k: Global distribution of Log_2_FCs (from DEqMS differential analysis) upon heat-shock in each state are plotted as violin and overlaid boxplots (median + inter-quartile range); adjusted p-values from non-parametric Kruskal-Wallis test (two-tailed, unpaired) are shown; dashed black and grey horizontal lines represent absolute Log_2_FC = 0 and 1, respectively.

Focusing on the 368 proteins that significantly changed upon heat-shock within any of the three states, it was clear that more proteins had significantly decreased rather than increased abundances in the heat-shocked condition for all three states (Fig. 2b)— consistent with the notion that stressed cells prioritise production of a few, key stress-response proteins (e.g., HSPs) at the expense of a broader range of non-essential proteins.^54^ As the abundances of almost all the mRNA transcripts corresponding to heat-shock-depleted proteins were unaltered by heat-shock (Fig. 1i, points to left-hand-side of vertical blue dashed line are not generally below the horizontal blue line), it was likely that this decrease was due to a combination of reduced global protein translation and active stress-induced protein degradation.^9,54,55^ A large subset of the heat-shock-depleted proteins were involved in DNA replication and repair processes, consistent with previous work in other proliferating human cell lines,^54,56^ as well as key stress-related signalling protein complexes (e.g., JAK–STAT, BLOC-2)(Supplementary Fig. S4d, Clusters 7 and 11; Supplementary Fig. S4e). Furthermore, as a group, the heat-shock-depleted proteins had a propensity to be at the lower end of the abundance values (Supplementary Fig. S4f), consistent with the lower relative copy numbers of signalling proteins (vs., e.g., proteostasis factors and general housekeeping proteins).^57,58^

Importantly, we detected no clear differences in either the identity of the proteins, or the magnitude of their depletion, upon heat-shock across the three cell states, with the same proteins that had reduced relative abundance upon heat-shock in one state also changing similarly in the other two states (Fig. 2a–b, Supplementary Fig. S4b–c). Although the overall trends in proteomic down-regulation upon heat-shock appeared to be similar across the three cell states, it was possible that different proteins were being degraded in the different states. Venn diagrams indicated a strong overlap between the heat-shock-depleted DPs between the three states (Figure 2b). There were more depleted proteins in the proliferating state; but the proteins that declined in quiescent and senescent states also declined in the other states, with only a few exceptions. As classification of DPs was based on largely arbitrary cut-offs and could therefore provide an incomplete picture of similarities in trends, we also mapped the down-regulated DPs classified in one state onto the volcano plots of the other two states (Figure 2c). This revealed even clearer trends, e.g., for the proliferating-specific DPs that did not overlap with the other two states in the Venn Diagram: even though they did not pass the fold-change-cut-offs for the quiescent and senescent states, almost all DPs trended towards being depleted even in these states. This commonality between the three states was despite that fact that the heat-shock-depleted DPs were, as a group, less abundant basally in quiescent and senescent states (compared with the proliferating state)(Supplementary Fig. S4g). Therefore, it appears that the set of proteins that is selected for degradation (or de-prioritised for replenishment via translation) during stress appears to be ‘hard-wired’ into the cell’s identity, and does not deviate with changes in the cell’s state— even if acquisition of that state alters the absolute levels of those proteins.

Whereas the changes to total protein levels after 2 hours of heat-shock were expected to be relatively modest, the polyubiquitylation response to heat-shock is expected to show major increases during the same time-scale.^9^ From the same IMR-90 cell preparations used for the transcriptomics and total proteomics, we enriched for the poly-ubiquitylated (polyUb) proteins via an optimised trypsin-resistant tandem-ubiquitin-binding-entity (trTUBE)–based approach (Fig. 2d; Supplementary Fig. S5).^59^ As expected and in contrast with the total proteome, the majority of variation in this trTUBE-enriched proteome (hereafter, polyUb-proteome) separated the basal from the heat-shocked samples (e.g., Fig. 2e, PC1 = 73 %; Supplementary Fig. S5a). We detected a robust and widespread polyUb-proteome increase upon heat-shock across all three cell states (Fig. 2f; Supplementary Fig. S5b–e; Table S5). In the proliferating and quiescent states, over half of the proteins quantified across all 18 samples had at least a two-fold increase in polyUb enrichment upon heat-shock, which is roughly the same order of magnitude as identified in previous work in proliferating cells.^9,55^ This figure dropped to 40 % for the senescent state and correlated with a generally muted HS-induced polyUb distribution (Fig. 2f). As with the total protoeme, there were very few proteins that displayed clear discrepancies in their heat-shock-induced polyUb enrichment across the three states (Fig. 2g–h; Supplementary Fig. S5c–g).

Heat-shock-depleted proteins at the total proteome level had higher polyUb-enrichment upon HS (Fig. 2i), consistent with their targeting for degradation via polyUb conjugation. However, the low general correlation between the total and polyUb-proteome dimensions (Supplementary Fig. S5h) is likely indicative of the diverse role of stress-induced polyUb beyond proteasomal degradation, e.g., K63-linked polyUb for trafficking to stress-induced granules.^9^ Indeed, protective sequestration of protein synthesis–related factors into cytoplasmic and nuclear granules is a major feature of successful stress responses. We found evidence that nucleolar ribosome biogenesis factors (Fig. 2j; Supplementary Figs. S5e–f (Cluster 3), S5i) and canonical cytoplasmic stress-granule (SG) components (Fig. 2k; Supplementary Fig. S5j) were all considerably enriched in the polyUb-proteome—although, in both cases, there were nuances in the sub-categories that were enriched *vs.* the rest of the proteome. Once again, there was no obvious difference between the three states.

This conserved robustness in proteome-wide protein degradation and polyUb responses between cell states also held true for proliferating *vs.* DNA-damage-induced senescent A549 human lung cancer cells exposed to heat-shock, or more prolonged proteotoxic stress using the proteasome inhibitor bortezomib (Supplementary Fig. S6; Tables S6–S9). The heat-shock-treated total proteome for A549 epithelia showed the same trend as in IMR-90 fibroblasts, i.e., a mild depletion in abundance for hundreds of proteins and increase in only a handful of proteins (including the HSP70s HSPA1B and HSPA6) (Supplementary Fig. S6d). By contrast, 24 h proteasome inhibition resulted in more widespread changes, including more proteins with increased total abundance. With the exception of HSPs, there was little correlation and limited overlap between heat-shock– and bortezomib– modulated total or polyUb-proteomes (Supplementary Fig. S6d–f), consistent with the concept that proteome-wide responses are tailored to the nature of the stress.^9,60^ Regardless of the stress, however, the responses of the senescent state mirrored that of the proliferating state within each experiment (Supplementary Fig. S6g–h).

Therefore, across multiple human cell types, and in response to multiple proteotoxic stresses, proteostasis circuits for stress-induced poly-ubiquitylation and degradation appear largely conserved between proliferating and senescent states.

### Stress-induced nuclear RNA-binding-protein sequestration into insoluble granules diverges in the senescent state

Reversible sequestration of RNA and proteins into detergent-insoluble aggregates or granules is another conserved, adaptive component of acute-stress responses,^61,62^ proposed to protect aggregation-prone nascent polypeptides and the protein production machineries themselves from irreversible aggregation during stress. Furthermore, stress-induced granules may serve to organise mRNA transcripts and proteostasis network components in a state poised for successful recovery from stress. To determine whether the same components formed granules upon heat-shock across proliferating, quiescent, and senescent states, we employed an unbiased approach based on low-speed centrifugation to sediment detergent-insoluble granules and aggregates (Fig. 3a–b, Supplementary Fig. S7a).^63^ MS-based bottom-up proteomics identified 3,420 proteins in these sedimented pellets; however, the majority of these were detected at similar levels upon HS (Supplementary Fig. S7b). Focusing on the set of proteins whose sedimentation did increase upon heat-shock, the insoluble proteome displayed clear differences between the senescent and non-senescent states upon heat shock—in direct contrast with the highly-conserved responses across the three states at the total- and polyUb-proteome levels. Only two DPs increased exclusively in the senescent state (Fig. 3c). Pairwise comparisons of the insoluble proteome fold-changes upon heat-shock revealed a strong positive correlation between proliferating and quiescent states; these were considerably weaker between the senescent and proliferating or quiescent states (Fig. 3d). From these distributions, it was clear that the discordance was predominantly due to a lack of heat-shock-induced insoluble fraction accumulation, in the senescent state, of proteins that accumulated similarly upon heat-shock in the quiescent and senescent states.

**Figure 3.**
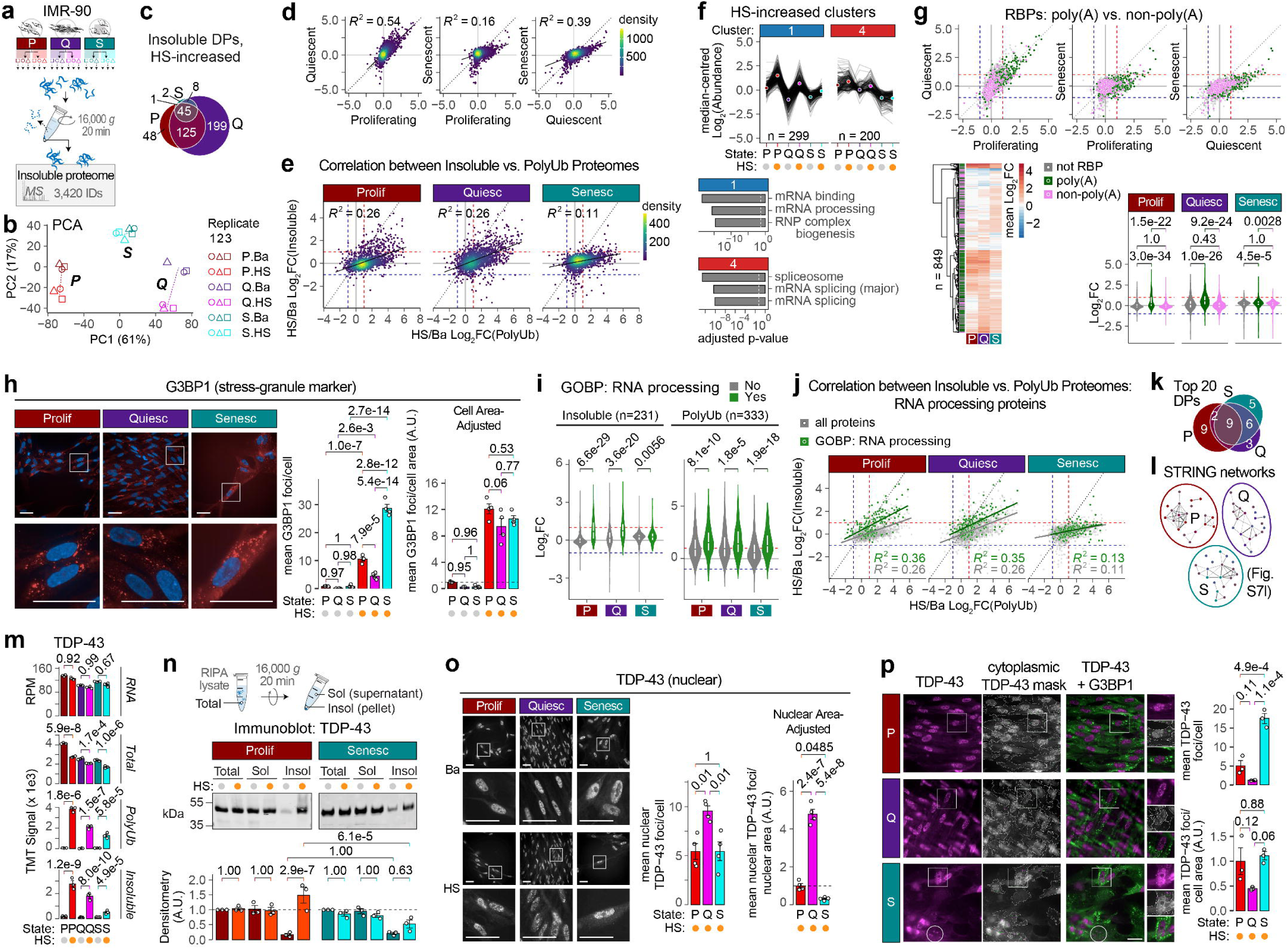
Stress-induced nuclear RNA-binding-protein sequestration into insoluble granules diverges in the senescent state. **a.** Experimental design for low-speed centrifugation-based insoluble proteome enrichment and quantification. **b.** Raw component scores from PCA of the top two PCs for the insoluble proteome suggests clustering of samples first by cell state, and then condition (Ba vs. HS). Separation appears less clear between HS and Ba senescent samples. **c.** Venn diagram showing overlap between proteins increased upon heat-shock in the insoluble proteome across cell states. **d–e.** HS-induced Log_2_FC values across each cell state for the insoluble proteome (d), or for the polyUb (x-axis) vs. insoluble (y-axis) proteomes (e). For each pairwise comparison, linear regression lines for y ∼ x, and squared Pearson’s correlation coefficients, are shown. Dotted lines represent y = x, and density of overlapping points are indicated with the colour scale. For (e), dashed lines represent Log_2_FC = 1 (red) or –1 (blue). **f.** HS-increased insoluble DPs are dampened in the senescent state. Top, traces of median-scaled Log_2_FCs for each DP in the HS-increased clusters 1 and 4 in the insoluble proteome. The mean of the median-scale Log_2_FCs for all proteins in the cluster per condition is overlayed as circles, coloured according to the condition. Bottom, top significant (adjusted p < 0.05, dashed white line) annotation terms by over-representation analysis (ORA) for the proteins in each cluster. ORA was performed using the gprofiler2 R package, with all 8,449 quantified in the IMR-90 total proteome as the custom background. See also Supplementary Fig. S7e–g. **g.** HS-induced insoluble proteome accumulation of mRNA-binding proteins is dampened in the senescent state. HS-induced alterations in the insoluble proteome for each state, with RNA-binding proteins (RBPs) highlighted based on classification by Trendel et al. (2019).^64^ Log_2_FCs for each group are shown as scatter-plots or violin and overlaid boxplots. Heatmap shows row-centred mean Log_2_-transformed normalised TMT abundances in the insoluble proteome for all quantified RBPs. Hierarchical clustering was performed using the complete agglomeration method with Euclidean distance measure for both columns (cell states) and rows (proteins). **h.** SG formation upon HS is conserved across cell states in IMR-90 fibroblasts, when correcting for relative cytoplasm area. IMR-90 fibroblasts in proliferating, contact-inhibited quiescent, or DNA-damage-induced senescent states were heat-shocked (44 °C for 2 h, HS), or kept at 37 °C, followed by immediate fixation. Cells were permeabilised and immuno-stained with anti-G3BP1 (canonical SG marker) and counter-stained with DAPI, before imaging and automated quantification of cytoplasmic SGs (G3BP1 foci) and number of cells (DAPI objects). Bar-charts show mean G3BP1 foci per cell, before or after adjusting for differences in cytoplasmic area (as calculated by the mean cytoplasmic masks for each cell state across all the wells). See also Supplementary Fig. S7j–k. **i–j.** Comparison of proteins belonging to the ‘RNA processing’ GOBP term (green) across insoluble and polyUb–proteomes suggests senescence-specific dampening at the insoluble proteome dimension without marked differences at the polyUb dimensions vs. non-senescent states. Distribution of HS-induced Log_2_FCs in the insoluble or polyUb-enriched proteomes, shown as violin + overlaid boxplots (i) or scatter-plots (j). Dashed lines represent Log_2_FC = 1 (red) or –1 (blue). For (j), regression lines for y ∼ x, and squared Pearson’s correlation coefficients, for ‘RNA processing’ proteins (green) vs. the rest of the proteome (grey), are shown. Dotted lines represent y = x. **k.** Venn diagram showing overlap between the top-20 HS-increased proteins in the insoluble proteome across cell states. **l.** STRING-database protein–protein interaction networks for the top 20 HS-increased insoluble proteome DPs in each cell state. The full STRING network (i.e., both functional and physical interactions) was included at medium confidence (minimum interaction score = 0.4). Fill colour of circles represents the degree of overlap for the protein in the top-20 lists. Line thickness represents the interaction confidence (as evaluated by the STRING database). See also Supplementary Fig. S7l. **m.** Normalised RPM (transcriptome) or TMT abundances (total, polyUb, and insoluble proteomes) for TDP-43 are plotted as bar-charts, and adjusted p-values from DESeq2 (transcriptome) or DEqMS (proteomes) are shown. **n.** HS-induced insoluble fraction accumulation of TDP-43 is dampened in the senescent state, without noticeable differences in the total or soluble fractions. IMR-90 fibroblasts in proliferating or senescent states were heat-shocked (44 °C for 2 h), or kept at basal conditions, and harvested by scraping in PBS. Cell pellets were lysed by freeze-thaw in RIPA buffer, and 1 mg of protein (as estimated by BCA assay) from each sample was processed for either denaturing lysis in 2 % SDS and 8 M urea (‘Total’), or soluble–insoluble fractionation by centrifugation (16,000 *g* for 20 min) into soluble (‘Soluble’) and pellet (‘Insoluble’) fractions. Equal volumes from each fraction were separated by SDS-PAGE, immunoblotted for TDP-43, and bands quantified using a LiCor Odyssey and Image Studio. **o.** TDP-43 nuclear foci formation upon HS diverges across cell states in IMR-90 fibroblasts, when correcting for relative nuclear area. IMR-90 fibroblasts were processed as in (h), but with immuno-staining with anti-TDP-43 instead of anti-G3BP1, and subsequent adjustment for nuclear rather than cytoplasmic mask area. See also Supplementary Fig. S8. **p.** Number of TDP-43 cytoplasmic foci is similar across cell states in IMR-90 fibroblasts, when correcting for relative cytoplasm area, and partially co-localises with G3BP1+ SGs. IMR-90 fibroblasts were processed as in (h), but with immuno-staining with both anti-TDP-43 and anti-G3BP1. For violin and overlaid boxplots in (g) and (i), adjusted p-values from non-parametric Kruskal-Wallis test (two-tailed, unpaired) are shown; dashed black, red, and blue horizontal lines represent Log_2_FC = 0, 1, and –1, respectively. For bar-charts in (h) & (n)–(p), data-points represent individual biological replicates, and bars +/- error-bars represent mean +/- standard-error. Adjusted p-values from Tukey’s HSD post-hoc test following one-way ANOVA are shown, even in cell area-adjusted comparisons in (h) & (p), where ANOVA p-values were not significant.

We confirmed that the heat-shock-enriched insoluble proteome we identified did not simply reflect increases in the total abundance of those proteins during the heat-shock period (Supplementary Fig. S7c). Most of these proteins were also enriched upon heat-shock in the polyUb–proteome (Fig. 3e), consistent with previous observations of stress-induced granule proteins.^9^ In the senescent state, although there was marked reduction in the number of proteins that accumulated in the insoluble proteome dimension upon HS, there was no clear reduction in the polyUb dimension. This observation suggested to us that senescent cells largely maintain stress-induced poly-ubiquitylation of the necessary proteins, but may not sequester these into insoluble granules.

Hierarchical clustering identified two distinct clusters that clearly increased upon heat-shock in the proliferating and quiescent states, and less so in the senescent state (Fig. 3f; Supplementary Fig. S7d–e; Table S10). ORA of proteins from these clusters identified a striking enrichment of RNA-related processes, with Cluster 1 displaying an additional enrichment in inflammation– and infection– related terms (Fig. 3f; Supplementary Fig. S7f–g; Table S10). Our data suggested a general heat-shock-enrichment in insolubility of RNA-binding proteins (RBPs)(Supplementary Fig. S7h), including those associated with both ribosomal and messenger RNA (rRNA and mRNA)–related processes. However, classification according to a high-confidence, integrated human RNA-binding protein (RBP) list^64^—based on UV-cross-linking across three human cell lines—revealed much stronger enrichment for RBPs that interact with polyA tracts (i.e., mature mRNA) vs. other RBPs in our insoluble proteomics data (Fig. 3g). Therefore, mRNA-binding proteins appear to be the major category with increased sedimentation upon heat-shock in proliferating and quiescent states, and impaired in the senescent state.

Many such mRNA-binding proteins are commonly found in stress-induced granules.^65^ We had failed to identify clear differences in the polyUb-enrichment of canonical cytoplasmic SG constituent proteins (Fig. 2k; Supplementary Fig. S5j) upon heat-shock between the three states; the same was true for the insoluble proteome (Supplementary Fig. S7i). Imaging of the canonical SG-marker G3BP1 in IMR-90 fibroblasts and hTERT-immortalised RPE-1 (hereafter, RPE-1) epithelia confirmed that all three states formed SGs under our HS conditions in (Fig. 3h; Supplementary Fig. S7j–k). Although the number of SGs per cell were higher in the senescent state, this scaled with the relative difference in cell size; indeed, the differences in SG numbers upon heat-shock between the three states were no longer significant after adjusting for the area of the cell mask in the experiment (Fig. 3h).

Cytoplasmic SG formation is generally associated with sequestration of mature mRNA, ribosomes, and RBPs related to protein translational control. Immature mRNA is also sequestered in various nuclear granules during stress. Indeed, several mRNA splicing and other RNA processing terms were enriched in both heat-shock-increased ORA clusters for the insoluble proteome (Fig. 3f; Supplementary Fig. S7f–g; Table S10), and proteins annotated in the ‘RNA processing’ GO term were significantly insoluble-enriched upon HS in all three states, although to a lesser extent in the senescent state (Fig. 3i). This group of RNA processing proteins still displayed a strong enrichment in the polyUb-proteome in senescence, providing further evidence that the divergence in RBP insolubility is unlikely to be a consequence of defective polyubiquitylation. Importantly, nine of the top 20 insoluble proteins in each state (based on Log_2_FC upon heat-shock) were shared between all three states (Fig. 3k), and formed similar protein–protein interaction networks (Fig. 3l; Supplementary Fig. S7l). Therefore, the insoluble accumulation defect of RNA processing-related proteins in senescent cells appeared to represent a dampening in the magnitude of the response rather than complete shut-off.

The above top-20 insoluble protein networks revealed a striking enrichment of heterologous nuclear ribonucleoproteins (hnRNPs) in all three states. This structurally and functionally diverse set of proteins are commonly associated with neurodegenerative diseases, especially amyotrophic lateral sclerosis (ALS) and fronto-temporal dementia (FTD).^66^ By far the most well-studied member of this protein class, the nuclear–cytoplasmic shuttling protein TDP-43 (gene name TARDBP), was in the 10 most heat-shock-enriched insoluble proteins for all three states in our data (Fig. 3m). We confirmed that these differences were only at the level of the detergent-insoluble TDP-43 fraction, with no detectable changes in total or detergent-soluble TDP-43 upon heat-shock in proliferating or senescent states (Fig. 3n).

TDP-43 resides in the nucleus in physiologic ‘unstressed’ conditions; upon stress, it forms both nuclear and cytoplasmic granules.^67^ Through immunofluorescence imaging, we detected higher numbers of heat-shock-induced nuclear TDP-43 foci in the quiescent state (Fig. 3o). This agreed with the proteomics data, in which the scale of insoluble proteome enrichment upon heat-shock was most pronounced in the quiescent state (Fig. 3c; Supplementary Fig. S7b). Although the numbers of nuclear TDP-43 foci were similar for the proliferating and senescent states, adjusting for the differences in nuclear size between the states suggested a mild yet significant reduction (p = 0.0485) in nuclear TDP-43 foci per unit area in the senescent state. The same trends were observed with a different TDP-43 antibody raised against the opposite terminal of the protein, and in RPE-1 cells (Supplementary Fig. S8a). We confirmed that this effect was not TDP-43–specific, as a similar state-dependent variation in numbers of nuclear stress-induced foci was observed for MATR3 (Supplementary Fig. S8b), another ALS-associated RBP that co-localises with TDP-43 in neuronal cytoplasmic aggregates.^68^ However, stress-induced granulation was not universal to all neurodegeneration-associated nuclear RBPs, as FUS showed no detectable foci upon heat-shock (Supplementary Fig. S8c).

It is well-established that a fraction of nuclear TDP-43 re-localises to cytoplasmic stress granules (SGs) with acute stress, including heat-shock. A similar trend was observed for cytoplasmic TDP-43 foci upon heat-shock, which largely co-localised with G3BP1 in all three states (Fig. 3p). The numbers of cytoplasmic TDP-43 foci per cell between states correlated inversely with the levels of nuclear TDP-43 foci—although, as with G3BP1 foci quantification (Fig. 3h), adjusting for cell size revealed almost identical numbers of TDP-43 foci per unit area.

### Senescent cells secrete misfolded proteins upon stress via the endosome–lysosome system

Although cytoplasmic TDP-43 foci co-localised with G3BP1 in all three states, we noticed that senescent cells alone consistently contained a distinct G3BP-negative fraction of TDP-43 cytoplasmic foci (e.g., Fig. 3p, white circle).^69^ Rather than representing another distinct cytoplasmic granule or aggregate, however, TDP-43 in the senescent state co-localised with the early endosome marker Rab5 (Fig. 4a). Based on their degree of co-localisation with Rab5, we noted two distinct sub-populations of cells: cells with fewer stress granules had more Rab5-co-localising TDP-43 foci, suggesting that endosomes and SGs represent alternative destinations for extra-nuclear TDP-43 trafficking. Regardless, all senescent cells under heat-shock harboured at least some Rab5-positive TDP-43 foci—in contrast with proliferating and quiescent cells.

**Figure 4.**
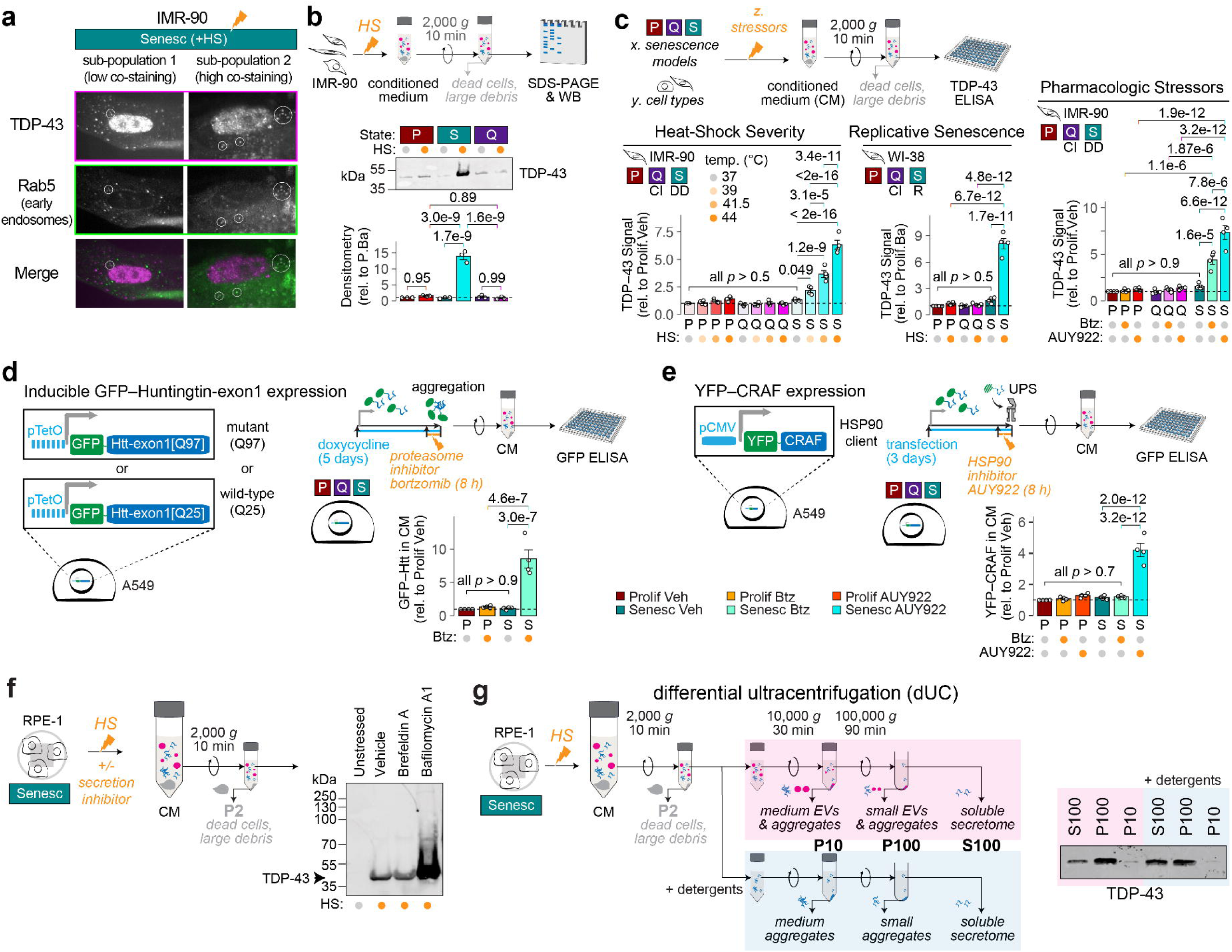
Misfolded proteins in the cytoplasm of senescent cells are secreted via the endosome–lysosome system. **a.** A sub-population of cytoplasmic TDP-43 co-localised with the early-endosome marker Rab5. Senescent IMR-90 cells were heat-shocked and processed as in Fig. 3h, but with immuno-staining with both anti-TDP-43 and anti-Rab5. Two distinct cell sub-populations were identified, based on the degree of co-localisation between TDP-43 and Rab5 foci. **b.** TDP-43 accumulates in the conditioned media (CM) upon HS of senescent cells. IMR-90 fibroblasts in proliferating, contact-inhibited quiescent, or DNA-damage-induced senescent states were heat-shocked (44 °C for 2 h, HS), or kept at 37 °C, in fresh complete growth medium, followed by collection of the CM, centrifugation and passing through a 0.2 µm filter to remove intact cells and debris, and analysis by SDS-PAGE and immunoblotting with anti– TDP-43. CM loading between samples was corrected for cell protein content (based on lysis and BCA assay on cells from which the CM was collected). Immunoblot bands were quantified using a LiCor Odyssey and Image Studio. **c.** Extracellular TDP-43 accumulation is a feature of the senescent cell response to proteotoxic stress, conserved across heat-shock severity (temperature), senescence inducer (replicative; oxidative stress), stress type (heat-shock; proteasome inhibition; HSP90 inhibition), and cell type (fibroblast; epithelial). CM was collected and processed as in (b), but with TDP-43 sandwich ELISA as a readout, according to manufacturer’s instructions. See also Supplementary Fig. S9a. **d.** Misfolded GFP–Htt-exon1[Q97] also accumulates in the CM of senescent A549 epithelia upon proteasome inhibition. A549 human lung adenocarcinoma cells containing a transgenic element for inducible expression of either GFP–Htt-exon1[Q25] or GFP–Htt-exon1[Q97] were established in proliferating or DNA-damage-induced senescent states before induction of Htt-exon1 expression by doxycycline addition for 5 days. Accumulation of proteins was induced by proteasome inhibition for 8 h, and CM was collected and processed as described in (b). Relative levels of GFP–Htt[-exon1[Q25] or GFP–Htt-exon1[Q97] in the CM across samples was quantified by GFP sandwich ELISA. **e.** Misfolded YFP–CRAF also accumulates in the CM of senescent A549 epithelia upon HSP90 inhibition, but not proteasome inhibition. A549 cells transiently-transfected with a plasmid for expression of the HSP90 client YFP–CRAF were established in proliferating or senescent states as in (d), and stressed for 8 h with the proteasome inhibitor bortezomib or HSP90 inhibitor AUY922. CM media collection and quantification was performed as in (d). Note that the GFP ELISA quantifies YFP as well as GFP. **f.** HS-induced extracellular TDP-43 levels are amplified by lysosome deacidification, but not by blocking conventional ER-Golgi-based secretion. DNA-damage-induced senescent RPE-1 epithelia were heat-shocked (44 °C for 2 h, HS), or kept at 37 °C (‘Unstressed’), in fresh complete growth medium containing either the ER-Golgi trafficking inhibitor Brefeldin A, the lysosomal v-ATPase inhibitor Bafilomycin A1, or DMSO-vehicle control. CM was collected, processed, and analysed by SDS-PAGE/immunoblotting as in (b). See also Supplementary Fig. S9c for densitometric quantification. For (c)–(e), data-points represent individual biological replicates, and bars +/- error-bars represent mean +/- standard-error. Adjusted p-values from Tukey’s HSD post-hoc test following significant (p < 0.05) one-way ANOVA are shown. **g.** Extracellular TDP-43 is in a mixture of soluble and small detergent-insoluble aggregates, and may or may not be encapsulated within extracellular vesicles (EVs). CM from heat-shocked senescent RPE-1 cells as in (f) was further fractionated based on a well-established differential ultracentrifugation protocol for crude separation of soluble, small EVs, medium-sized EVs, with parallel processing of the same CM after detergent-based lysis of EVs. Note that the distribution of TDP-43 does not markedly change upon detergent addition, suggesting the presence of a detergent-insensitive aggregated TDP-43 sub-population.

Accumulation of TDP-43 in cytoplasmic neuronal aggregates is associated with several neurodegenerative diseases. It has also been proposed that cytoplasmic TDP-43 can spread between cells via extracellular vesicle (EV) secretion.^70–72^ Therefore, we tested whether the reduction of intracellular insoluble TDP-43 could be due to its secretion out to the conditioned medium. Collected at the end of the 2-hour heat-shock period, the conditioned medium from heat-shocked senescent cells contained substantially more TDP-43 than medium from proliferating or quiescent cells (Fig. 4b). Stress-induced TDP-43 increase in the conditioned media was observed across all of our senescence models—irrespective of cell type or senescence inducer (Fig. 4c, Supplementary Fig. S9a). Furthermore, extracellular TDP-43 accumulation was reproduced with more moderate heat-shock temperatures (39 and 41.5 °C), and the pharmacologic proteotoxic stressors bortezomib (proteasome inhibitor) and AUY922 (inhibitor of the molecular chaperone HSP90, which is involved in TDP-43 turnover).^73,74^ This conserved, senescence-specific increase in TDP-43 levels in the conditioned medium was unlikely to be due to higher levels of death (and therefore release of intracellular proteins into the medium), as senescent cells were more resistant to death following these same proteotoxic stresses compared with proliferating and quiescent cells (Fig. 1b–d). We also detected no increase in the levels of YFP in the medium when YFP-expressing senescent cells were subjected to heat-shock (Supplementary Fig. S9b), confirming that protein release from senescent cells was not an indiscriminate, proteome-wide response to stress.

Based on these findings, we hypothesised that protein release was selective for misfolded proteins, which would include cytoplasmic TDP-43 sub-populations in the absence of stabilising RNAs and/or other RBPs, but would exclude well-folded and highly-stable YFP. In agreement, fusing GFP with the highly aggregation-prone polyglutamine-expanded mutant Huntingtin-exon1 (GFP–Htt-exon1[Q97]), but not its wild-type soluble counterpart (GFP–Htt-exon1[Q25]), increased its extracellular levels upon proteasome inhibition in senescent A549 epithelia (Fig. 4d). Increased release was also observed with YFP fused to the protein kinase CRAF, but only when it was destabilised by inhibition of its chaperone HSP90—and not under the same proteasome inhibition conditions that triggered GFP–Htt-exon1[Q97] release (Fig. 4e). This observation suggests that extracellular release remains selective for the proteins that misfold, which can differ with the nature of the stress.

As all the misfolded proteins we tested—TDP-43, Htt[Q97], and CRAF—are soluble nuclear or cytoplasmic proteins, we thought it unlikely that conventional ER–Golgi-based pathways were involved in their secretion. Indeed, treating senescent cells with the Golgi trafficking inhibitor Brefeldin-A during heat-shock had no impact on extracellular TDP-43 levels (Fig. 4f). By contrast, triggering lysosome deacidification by Bafilomycin-A1 treatment led to a substantial increase. Bafilomycin-A1 treatment has been shown to lead to an increase in small extracellular vesicle (sEV) release, including in the context of pathologic forms of TDP-43.^71,72,75^ To determine whether TDP-43 secretion in senescent cells was in the form of sEVs, we performed differential ultracentrifugation of conditioned medium from senescent RPE-1 cells to enrich fractions containing medium-size EVs, sEVs, and soluble (non-EV-enclosed) proteins (Fig. 4g).^76–78^ Although these fractions suggested that extracellular TDP-43 is present in both soluble and sEV fractions, the propensity of TDP-43 to aggregate made it difficult to distinguish between aggregated and EV-enclosed populations, which could conceivably sediment at the same centrifugation conditions (c.f. 16,000 *g* for 20 min used for enrichment of intracellular detergent-insoluble proteomes). We therefore adjusted the conditioned medium to the equivalent detergent concentrations as the RIPA buffer used for cell lysis and insoluble intracellular proteome enrichment (1 % Triton X-100, 0.5 % SDS, 0.5% sodium deoxycholate), which would also disrupt any membrane-containing vesicles in the conditioned medium. Repeating the differential ultracentrifugation experiment under these conditions did not markedly alter the TDP-43 profile, suggesting the presence of a detergent-resistant extracellular TDP-43 population that may or may not be EV-enclosed.

Our data suggests that insoluble misfolded protein aggregates in the cytoplasm of senescent cells—which, under stress, includes a sub-population of TDP-43—are re-trafficked to the endosome–lysosome system. Secretion of this cargo is presumably driven by lysosomal exocytosis, a physiologic feature of senescence^79,80^ linked to lysosome deacidification,^51,52,81^ which can be stimulated further by pharmacologic agents such as Bafilomycin A1.

### Misfolded protein secretion mediated by the vesicle-associated chaperone DNAJC5 is an adaptation to declining intracellular proteostasis capacity

YFP–CRAF is not an intrinsically aggregation-prone protein, and is intracellularly degraded by the ubiquitin–proteasome system upon HSP90 inhibition in proliferating cells.^82^ Its extracellular accumulation in the senescent state therefore suggests that stress-induced protein secretion is not reserved only for aggregation-prone proteins, but could serve as a general alternative PQC mechanism in senescent cells. In highly-proliferative cells, misfolded proteins are sequestered upon acute proteotoxic stress—including pharmacologic proteasome or HSP90 inhibition—at a juxtanuclear site commonly referred to as the aggresome or JUNQ (juxtanuclear quality-control compartment).^7,83^ One study in adult mouse dermal fibroblasts reported a lack of aggresome formation upon heat-shock or proteasome inhibition in senescent states, although they found the same deficiency in the proliferating state (but not in quiescent and immortalised states).^32^ From our insoluble proteomics data, we identified a striking decrease in components of the centrosome—the major microtubule organising centre (MTOC), and a crucial structure for aggresome formation^84^— during 2 h heat-shock, especially in the senescent state (Supplementary Fig. S7d−e, Cluster 7). Solubilisation is generally a sign of centrosome disassembly and re-localisation of the resident proteins into cytoplasmic pools.^85,86^ Focusing on the centriolar satellite proteins whose depletion has been shown to impair aggresome formation,^84^ their total protein levels were lower in quiescent and senescent states (as expected for non-dividing cells)(Supplementary Fig. S10). Although their total levels did not decrease further with heat-shock, the polyUb-enrichment was increased in the proliferating state for three of the four proteins. The polyUb enrichment was lower in the quiescent state, and almost completely abolished in the senescent state. Together with the fact that the insoluble proteome levels were lowest in the heat-shocked senescent samples for all four proteins, we hypothesised that aggresome formation may be impaired in this state—and that misfolded protein secretion could serve as an adaptive PQC strategy to compensate.

Initially using the juxtanuclear accumulation of proteins tagged with K48-linked polyUb (Ub[K48]) as a marker for the aggresome, we identified aggresome formation in proliferating A549 cells exposed to the proteasome inhibitors bortezomib or MG-132 for 24 h (but not at the earlier time-point of 6 h)(Supplementary Fig. S11a). Through a more stringent and semi-automated image analysis pipeline for aggresome quantification (Supplementary Methods), we confirmed that both A549 and IMR-90 cells in proliferating states formed aggresomes upon proteasome inhibition, although there were far fewer aggresome-positive cells in IMR-90 than in A549 cells (Fig. 5a−d; Supplementary Fig. S11b−c). The percentage of cells with an aggresome was considerably lower in the senescent state of both cell lines, consistent with our hypothesis. Note that, due to the overlapping nature of cells in the contact-inhibited state, it was not possible to quantify aggresomes reliably in this state, although there were clearly strong Ub[K48] foci that resembled aggresomes in a substantial proportion of cells (Supplementary Fig. S11c).

**Figure 5.**
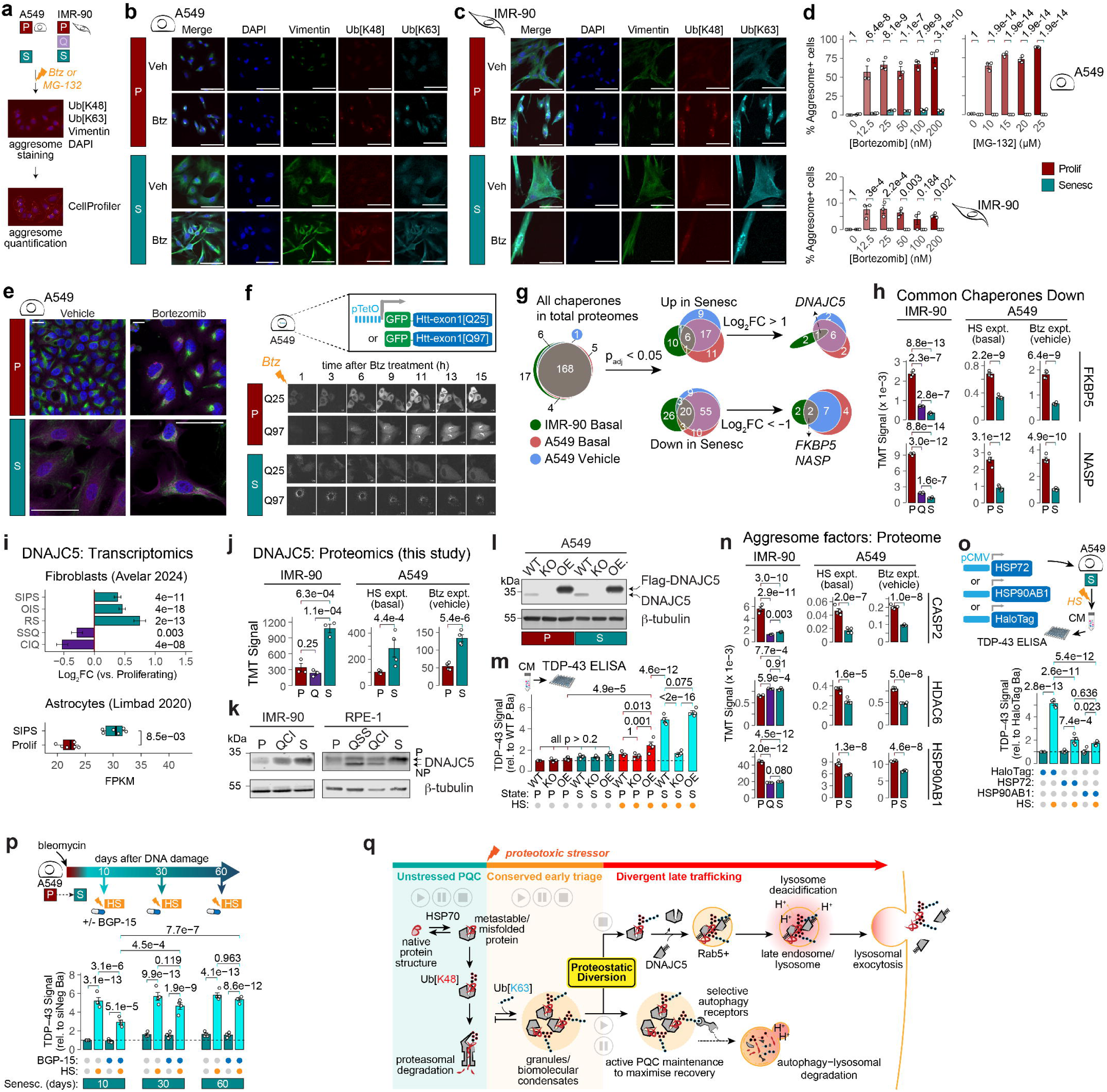
Misfolded protein secretion is a proteostatic adaptation to reduced intracellular PQC capacity in the senescent state. **a.** Imaging-based quantification for aggresomes formed following proteasome inhibition with bortezomib (Btz), or DMSO-vehicle control, for 24 h in A549 human lung adenocarcinoma and IMR-90 human lung fibroblasts in various cell states. At the end of 24 h treatment, cells were fixed, permeabilised, and immuno-stained with anti–Ub[K48], anti–Ub[K63], and anti– Vimentin, and counter-stained with DAPI, before imaging immediately on an ImageXpress Confocal HT.ai High-Content Imaging System. Aggresomes were quantified using a custom pipeline on CellProfiler, based on overlap between Ub[K48], Ub[K63], and vimentin signals at a juxtanuclear region. Note that CellProlifer-based quantification was not performed in the contact-inhibited quiescent state, due to overlapping cell masks between neighbouring cells. **b–c.** Representative images used for aggresome quantification, as described in (a), in A549 (b) or IMR-90 (c) cells, at the 200 nM Btz concentration. Scale-bars = 50 µm. See also Supplementary Fig. S11. **d.** Aggresome formation is severely dampened in the senescent state. Bar-charts show percentage of A549 (top) and IMR-90 (bottom) cells quantified with an aggresome for each cell state and treatment, using the semi-automated CellProfiler pipeline described in (a). **e.** Dispersed stress-induced Ub[K63] foci remain in proximity to vimentin intermediate filaments in the senescent state. A549 cells were treated as in (b) (with 100 nM Btz), with the exception of omitted anti-Ub[K48] immuno-staining, and imaged on a Nikon Ti2 widefield microscope with 20x objective. Scale-bars = 10 µm. See also Supplementary Fig. S12a. **f.** GFP–Htt-exon1[Q97] remains in disperse foci upon proteasome inhibition in senescent A549 cells. A549 cells in proliferating or senescent states expressing GFP–Htt-exon1[Q25] or GFP–Htt-exon1[Q97] for 5 days were treated with 100 nM Btz, and imaged in the GFP channel overnight (16 h) on an Olympus Evident SpinSR spinning-disk confocal microscope using a 40x air-immersion objective. **g.** All molecular chaperones (based on the 259 in the Proteostasis Consortium’s annotation)^154^ quantified in the three total proteomics experiments were filtered according to significant differences in the senescent vs. non-senescent states in unstressed (basal or vehicle-treated) conditions, first by adjusted p-values (as calculated by DEqMS), and then by Log_2_FC cut-offs. Venn Diagrams represent overlap between the three datasets in chaperones that increased (Up) or decreased (Down). Only one and two chaperones were commonly up and down, respectively, in all three datasets after both p-value– and Log_2_FC– based cut-offs. **h.** Bar-charts showing total proteome abundances of the two senescence-increased chaperones shared between all three datasets, using the filtering criteria in (g). **i–k.** Increase in DNAJC5 transcript (i) and protein (j–k) levels is a feature of senescence across senescence inducers and cell types, and is not conserved in quiescent fibroblast states. For (i), fibroblast Log_2_FC, standard-error, and adjusted p-values are as calculated in the meta-analysis in Avelar et al. (2024),^88^ and astrocyte q-value is as calculated by Limbad et al. (2020).^89^ fpkm: ‘fragments per kilobase per million reads’. For (k), 10 µg of protein lysate from IMR-90 or RPE-1 cells in proliferating (P), contact-inhibited quiescent (QCI), serum-starved quiescent (QSS), or DNA-damage-induced senescent (S) states were separated by SDS-PAGE, immunoblotted with anti–DNAJC5 and anti–beta-tubulin (loading control), and bands quantified using a LiCor Odyssey and Image Studio. **l.** Validation of DNAJC5 knock-out and over-expression in A549 cells.^97^ 10 µg of protein lysate from DNAJC5^-/-^ (knock-out, KO), DNAJC5^-/-^::Flag–DNAJC5 (over-expression, OE), or parental wild-type (WT) A549 cells in proliferating or DNA-damage-induced senescent states were separated by SDS-PAGE, immunoblotted with anti–DNAJC5 and anti–beta-tubulin (loading control), and bands quantified using a LiCor Odyssey and Image Studio (shown in Supplementary Fig. S14b). **m.** DNAJC5 is required for HS-induced TDP-43 secretion in senescent A549 cells. DNAJC5 WT, KO, or OE A549 cells in proliferating or DNA-damage-induced senescent states were heat-shocked (HS, 44 °C) or kept at basal conditions, for 2 h, followed by CM collection, centrifugation, and filtration to remove dead cells and large debris. TDP-43 levels in the CM were quantified by TDP-43 ELISA. See also Supplementary Fig. S14c. **n.** The key aggresome-trafficking factors Caspase-2 (CASP2), HDAC6, and HSP90AB1 are at lower protein levels in the senescent state of IMR-90 and/or A549 cells, as quantified in the total proteomes. Note that CASP2 and HSP90AB1 levels are similarly reduced in contact-inhibited quiescent IMR-90 cells. **o.** Heterologous over-expression of HSPA1 (HSP72) or HSP90AB1 (HSP90-beta) significantly reduces extracellular TDP-43 levels in senescent A549 cells upon heat-shock. A549 cells were made senescent by DNA-damage for 7 days, followed by lipid-based transfected with pCMV::mScarlet–HSP72, pCMV::Flag–HA–HSP90AB1, or pCMV::HaloTag for three days. Subsequent HS, CM collection, and TDP-43 quantification was performed as in (m). **p.** Pharmacologic potentiation of the heat-shock-response by BGP-15 significantly reduces extracellular TDP-43 release upon HS in shallow but not deep senescence. A549 cells were made senescent by DNA-damage for 10, 30, or 60 days, followed by acute HS as described in (m), +/- 10 µM BGP-15 (added 30 min before HS). CM collection and TDP-43 quantification was performed as in (m). **q.** Proposed model for the proteostatic diversion in senescent states that leads to misfolded protein secretion into the extracellular space. In unstressed cells, relatively low basal rates of protein misfolding fall within intracellular proteostasis capacities in all three cell states, and is turned over by canonical chaperone-mediated ubiquitin–proteasome system targeting. Acute increases in misfolded protein load, such as during proteotoxic stress, initiate cytoprotective transcriptional and post-transcriptional programmes, most of which are well conserved across cell states. However, whereas proliferative and quiescent states expend considerable resources in maintaining their misfolded proteins and stress-induced protein–RNA granules in reversible forms, senescent states re-traffic the same cargo into the endosome–lysosome system and ultimately outside of the cell, in a process requiring DNAJC5, and exacerbated by lysosomal deacidification, which is a hallmark of senescence. The impact of these extracellular misfolded and aggregation-prone proteins on surrounding cells in the context of tissue function and inflammation is a direction for future work. For all bar-charts, data-points represent individual biological replicates, and bars +/- error-bars represent mean +/- standard-error. For (d) and (o)–(p), adjusted p-values from Tukey’s HSD post-hoc test following significant (p < 0.05) one-way ANOVA are shown. For (h), (j), and (n), adjusted p-values from DEqMS are shown.

In addition to differences in the Ub[K48] localisation, we noticed that the Ub[K63] signal— classically considered a polyUb linkage that acts as a trafficking signal, rather than a proteasomal degradation tag—appeared to be dispersed in smaller foci throughout the cytoplasm of proteasome-inhibited senescent cells, rather than co-localising with Ub[K48] at the juxtanuclear site. Intriguingly, these foci remained in close vicinity to the intermediate filament vimentin, even though vimentin did not collapse to form a characteristic ‘cage’ around the aggresome at the microtubule-organising-centre, as it does in stressed proliferating cells (Fig. 5e; Supplementary Fig. S12a).^7,83^ The disperse localisation was replicated for GFP–Htt[Q97] in senescent state, in contrast with the proliferating state, where the misfolded protein reporter localised at the juxtanuclear site (Fig. 5f; Supplementary Fig. S12b). Therefore, the difference in Ub[K63] foci was most likely due to altered trafficking of the same PQC substrates, rather than different substrates being tagged with Ub[K63] between the two states.

Finally, we attempted to identify the molecular chaperone requirements for this alternative trafficking of misfolded proteins from the intracellular aggresome route to the extracellular secretion route, starting with the basal (i.e., unstressed) distributions of all chaperones quantified in our three total proteomics datasets (Supplementary Fig. S13a). As the secretion phenotype was observed across cell models, and was not observed in quiescence, we filtered for significantly-altered chaperones (adjusted p < 0.05) that changed in the same in the senescent state vs. both other states, and in all three total proteomes. With these criteria, there were 6 and 20 chaperones that consistently increased and decreased, respectively, in the senescent state (Fig. 5g). We noted that many of the major chaperones were filtered out of the list due to being of lower abundance in both the quiescent and senescent states (vs. proliferating), including all 8 subunits of the TRiC/CCT complex, histone chaperones, and the HSP90–R2TP complex (Supplementary Fig. S13b). Applying an additional Log_2_FC filter narrowed down the list to only two common less-abundant chaperones—the histone chaperone NASP, and the steroid-hormone receptor chaperone FKBP5 (Fig. 5h)—and only one common more-abundant chaperone: the J-domain protein DNAJC5.

Mining two meta-analyses of transcriptomics datasets across senescent cell-types and senescence induction methods,^87,88^ we found that DNAJC5 was significantly upregulated in senescent fibroblasts (*vs.* proliferative fibroblasts) at the transcript level, irrespective of senescence inducer (Fig. 5i). DNAJC5 upregulation was also observed in replicatively-senescent astrocytes.^89^ Therefore, transcriptional upregulation of DNAJC5 appears to be a consistent feature of the senescent state. We confirmed this increase at the protein level in IMR-90 and RPE-1 cells (Fig. 5j−k). Importantly, DNAJC5 upregulation was not observed in contact-inhibited or serum-starved quiescent states, either at the transcript or protein levels (Fig. 5i−k).

DNAJC5, also known as Cysteine-String-Protein-alpha (CSPIZl), is tethered to endosomes via palmitoylation of its cysteine-string-domain, and is best known for its role in pre-synaptic vesicle trafficking in neurons. Loss-of-function DNAJC5 mutations are causative for adult-onset neuronal ceroid lipofuscinosis, and the DNAJC5 locus is associated with several other ageing-associated neurodegenerative conditions, including Alzheimer’s, Parkinson’s, and Huntington’s Diseases.^90,91^ Mechanistically, heterologous overexpression of DNAJC5 has been shown to drive unconventional secretion of misfolded proteins such as TDP-43, IZl-synuclein, Htt[polyQ], and Tau, and is therefore proposed as a facilitator of aggregate seeding and spreading.^92–96^ As our data showed that even non-neuronal cells in senescent states displayed increased DNAJAC5 protein levels, we hypothesised that senescence-associated misfolded protein secretion was enabled by re-trafficking to DNAJC5-mediated secretion pathways.

A549 cells with CRISPR-mediated knockout of DNAJC5 (DNAJC5^-/-^)^97^ showed no discernible differences in their ability to become senescent upon DNA-damage vs. the same cells with heterologous ‘rescue’ by overexpression of Flag–DNAJC5 (DNAJC5^-/-^::Flag– DNAJC5) (Fig. 5l; Supplementary Fig. S14a–b), indicating that DNAJC5 was not necessary for acquisition of the senescent state. However, the DNAJC5^-/-^ cells had an almost complete abrogation of stress-induced TDP-43 secretion, which was rescued in the DNAJC5^-/-^::Flag– DNAJC5 cells (Fig. 5m); we reproduced this result with transient siRNA-mediated knockdown in wild-type A549 cells (Supplementary Fig. S14c). Although extracellular TDP-43 levels were restored in senescent DNAJC5^-/-^::Flag–DNAJC5 cells, they were not significantly higher than senescent wild-type A549s, despite having far higher DNAJC5 levels ((Fig. 5l; Supplementary Fig. S14b). Similarly, DNAJC5^-/-^::Flag–DNAJC5 also modestly increased extracellular TDP-43 levels upon stress even in the proliferating state— consistent with previous reports involving DNAJC5 overexpression;^92–96^ yet levels were not at the same magnitude as in the wild-type senescent A549s, despite the much higher DNAJC5 protein levels in these cells. These observations suggest that DNAJC5 is necessary but not sufficient for the misfolded protein phenotype in senescence. Furthermore, DNAJC5 levels are unlikely to be limiting in the senescent state, as increasing its expression levels does not further increase extracellular TDP-43 levels. Therefore, additional context-specific PQC factors drive senescence-associated misfolded protein secretion.

It was possible that declining intracellular PQC capacity was the additional contributor to re-trafficking the misfolded protein load to DNAJC5, which would not be the case in the proliferative state. In addition to decline in insoluble (and presumably assembled) centrosomal proteins discussed above, multiple PQC components involved in cytoplasmic aggresome formation—including the chaperones HSP90AB1/HSP90β, the DUB CASP2/Caspase-2, and the dynein-binding shuttling factor HDAC6^7,98^—were lower at the protein level in the senescent state in IMR-90 and/or A549 cells (Fig. 5n). Furthermore, whereas basal HSP70 levels were similar across the three states, senescent cells were impaired in stress-induced upregulation of HSPA1/HSP72 and HSPA6/HSP70BL (Fig. 1j)— further supporting the hypothesis that the senescent state has a diminished chaperoning capacity at both baseline and stress-inducible levels.^30,36,38,39^ If re-trafficking to DNAJC5 in senescence was a result of this declining in canonical HSP70– and/or HSP90–mediated pathways, we reasoned that restoring levels of these chaperones should dampen the stress-induced secretion phenotype.

First, we artificially overexpressed the chaperones HSPA1/HSP72 or HSP90AB1/HSP90β in senescent wild-type or Htt[Q97]–expressing A549 cells, and monitored stress-induced secretion of TDP-43 or Htt[Q97], respectively (Fig. 5o; Supplementary Fig. S14d). There was a marked reduction in TDP-43 and Htt[Q97] levels in the media of either of the HSP-overexpressing (vs. HaloTag-transfected control) senescent A549 cells. Reduction in extracellular TDP-43 levels could be replicated by pharmacologic potentiation of heat-shock-induced HSF1 target-gene expression with the small-molecule BGP-15 (Supplementary Fig. S14e).^99,100^ However, BGP-15-mediated rescue of TDP-43 secretion appeared to be dependent on the duration of senescence (sometimes referred to as senescence ‘depth’), as indicated by the reduced effect in A549 cells kept senescent for 30 or 60 days after DNA-damage (Fig. 5p).

Taken together, we propose that senescent cells divert their misfolded protein load to DNAJC5-mediated secretory routes as a response to their declining intracellular proteostasis capacity, and that this can be rescued by boosting intracellular PQC—up to a point (Fig. 5q).

## DISCUSSION

Cells within tissues must counter-act stress. Their molecular coping strategies have been assumed to be hard-wired into all eukaryotic cells. We propose here that, although transcriptional programs that respond to proteotoxic stresses might be hard-wired, the protein-level consequences diverge considerably with cellular state. Most prominently, the stress-triggered formation of RNA–protein granules that has been assumed to be a critical component of successful stress responses appears to be dispensable for senescent cells— without any detected compromise in stress resilience. Instead, senescent cells re-traffic these RBPs, as well as other aggregation-prone or misfolded proteins, into the endosome– lysosome system through a triage process requiring the vesicle-associated chaperone DNAJC5. Once in the endo-lysosomal system, these misfolded proteins likely follow the same routes as other cargo, which includes a constitutive level of lysosomal exocytosis that is a feature of the senescent state.^79,80^

Although this secretion route would effectively reduce the misfolded protein burden within senescent cells, the consequence of this increased extracellular load on the surrounding cells, and on the tissue as a whole, needs to be explored. It is possible that this burden could contribute to the chronic stress, inflammation, and tissue dysfunction associated with senescent-cell accumulation, in addition to the conventional SASP factors such as cytokines and chemokines. It is also possible that misfolded protein secretion is an adaptive strategy for multi-cellular proteostasis, by sharing the burden across cells with higher PQC capacity (e.g., surrounding non-senescent cells, or phagocytic immune cells recruited by the SASP). Additionally or alternatively, non-cell-autonomous stress signalling, i.e., communicating stress detected in one cell across a tissue or organism, is a concept that has gained traction in model organisms—especially the nematode worm *C. elegans*—but has limited direct evidence in mammalian systems.^101–105^ It is possible that misfolded protein secretion by senescent cells serves as such a signal to mount protective responses in mammalian tissues as a whole.

The types of proteins that failed to accumulate in the insoluble fraction of senescent cells— i.e., RNA-binding proteins, with intrinsically-disordered regions and high aggregation propensities in the absence of stabilising nucleic acids—overlapped considerably with proteins that have been found to accumulate in aggregates with age across organisms.^5,106,107^ The accumulation of many of these proteins in both intra- and extra-cellular deposits is associated with neurodegeneration, although the extent to which this is driven by loss-of-physiologic-function, vs. gain-of-toxic-function, remains to be determined.^108^ For example, although TDP-43 has long been known to exacerbate pathologic aggregation of several other neurodegeneration-associated proteins, more recent work suggests that loss of TDP-43’s mRNA splicing activity leads to cryptic exon translation to drive formation of non-functional proteins—which would further stress a cell’s proteostatic capacity.^66,109–112^ Intriguingly, DNAJC5 appears to be a splicing target of TDP-43 with a cryptic exon that would render it non-functional.^113–115^ Moreover, TDP-43 in brain pericytes has been shown to regulate transcription of IL-6—a classical SASP factor.^116^ Therefore, it is possible that the senescent-cell-context creates negative-feedback loops where stress-induced loss of TDP-43 due to its aggregation and secretion leads to loss of critical SASP mediators, such as DNAJC5 and IL-6, which in turn results in reduced TDP-43 secretion. DNAJC5 is transcriptionally upregulated in senescent astrocytes,^89^ which are linked not just to TDP-43 seeding in neurodegeneration,^117^ but also to neuroinflammation via potentiation of the cGAS–STING pathway.^118^

Within this neuronal context, we must also decipher the role played by secretion of such proteins in seeding and/or spreading across cells. Impairments to the endosome–lysosome system increase TDP-43’s aberrant localisation and ALS–FTD pathology, and TDP-43 loss-of-function inhibits endosomal trafficking—potentially creating, in this case, a positive-feedback loop to drive pathology.^119–121^ The direct relationship between Rab5 and TDP-43 is unclear: acute knockdown of TDP-43 appears to have no impact on Rab5+ endosomes (but does on Rab11+ endosomes),^122^ and one study in a sporadic ALS model found an upregulation of Rab5+ endosomes in the vicinity of—but not co-localising with—TDP-43 cytoplasmic aggregates.^123^

Except for the divergence in misfolded protein fates, the similarity between other components of the HSR between states was initially surprising. Earlier global comparisons with replicative senescence reported attenuated stress responses at transcriptional and/or translational levels.^30,36^ One study in the WI-38 human fetal lung fibroblast line, in response to the identical heat-shock conditions used here, reported mild dampening of heat-shock-induced HSP induction, reduction in proteasome activity, and marked impairment in alternative mRNA splicing, in the senescent state.^30^ Our findings point towards similar trends, with slightly lower degrees of HSP induction, lower numbers of significantly-depleted proteins and significantly-increase polyUb proteins, and impairment of RBP insolubility (including spliceosome components) upon HS in senescent vs. both non-senescent states. It is also possible that these differences would be more pronounced in replicative senescence—which involves serial passaging and thus more chronic changes to the proteostasis network than the acute, stress-induced premature senescence models used here. Note that another study in replicatively-senescent human mesenchymal stem cells proposed a decline in intracellular misfolded protein degradation capacity (e.g., via decline in levels of the HSP70-associated E3 ubiquitin ligase CHIP/STUB1),^36^ which also fits with our model.

In addition to decline in levels of the canonical intracellular proteostasis network, we also observed loss of cell-division-related structures that are also critical for intracellular PQC in proliferating cells. For example, senescent states had clearly lower levels of centriolar satellite proteins in the insoluble proteome, which in their case is generally associated with assembled (insoluble) vs. unassembled (soluble) sub-populations. Based on the literature, the centrosome/MTOC displays reduced immuno-staining during heat-shock,^124–128^ although this has been attributed to a decrease, rather than increase, in solubility of centrosomal proteins such as pericentrin/PCNT.^126^ The accumulation of aggregation-prone misfolded proteins at this location has not been linked directly to these changes in MTOC components; however, the fact that HS-induced MTOC protein disruption can be rescued by over-expression of HSP70 suggests at least an indirect connection between aggresome assembly and changes in MTOC solubility.^128^

We thus propose that the decline in intracellular proteostasis capacity of senescent states creates an impetus for alternative PQC strategies. The endosome–lysosome system, which is already re-modelled as part of the senescence programme, appears a logical re-trafficking route. Indeed, one study found that certain aspects of the SASP appear to be mediated via lysosomal exocytosis.^79^ Notably, DNAJC5 was identified within lysosomes targeted via chaperone-mediated autophagy. It is possible that unconventional secretion of misfolded, poly-ubiquitylated proteins is an as yet unappreciated part of this same phenotype. There is also emerging evidence for more wide-spread endosomal dysfunction in senescence, with accumulation of endosomal proteins—including Rab5—in atypical hybrid cytoplasmic structures.^129^ As pharmacologic suppression of endosomal trafficking and lysosome exocytosis has recently been shown to have senolytic effects,^80^ stimulating proteostasis collapse by intracellular aggregate accumulation could add to this therapeutic window.

Our work adds an additional dimension to current debate on the development and deployment of senotherapeutic strategies for healthspan. Senescent cells appear to encompass a spectrum of states, with degrees of overlap between features varying with cell and tissue type, senescent inducer, and senescent ‘depth’.^130^ It is also likely that several senescence-associated features are shared to some extent by non-senescent cells within the human body. Therefore, finding common vulnerabilities for all senescent cells, without limiting toxicity to any non-senescent cell populations, presents a considerable challenge.

Even if we did discover common vulnerabilities, it remains to be seen whether eradicating all senescent cells is therapeutically desirable. As well as the important physiologic roles of senescent cells in processes that are already attenuated in older people (e.g., wound-healing, tumours suppression), senescent cells can comprise a non-negligible proportion of cell mass in certain tissues with age.^13^ It is therefore unclear whether all tissues can successfully tolerate removal of the entire senescent cell burden, even from a purely biomechanical perspective.

An alternative strategy that could have less drastic side-effects is through ‘senomorphics’, i.e., therapeutics that blunt 9 specific pathologic features of chronic senescent cell accumulation. The most obvious senomorphic approach is shutting off the SASP, which would serve to reduce chronic inflammation. However, the nature of the SASP also varies considerable across senescence models,^131^ and presents similar challenges to senolytics, both from the perspective of common and selective pathways for modulation, and from balancing effects of beneficial vs. harmful effects. For example, it is possible to imagine a scenario where blunting the SASP cloaks senescent cells even more effectively from the immune system, and ultimately results in the increase in total senescent-cell burden, making certain tissues even more dysfunctional. If, as our data suggests, misfolded protein secretion is a more universal feature of senescent cells under stress, and provided that it serves no healthspan-promoting function in tissue homeostasis, senomorphics that target this feature specifically could be a well-tolerated method of reducing the systemic proteostasis burden. Our BGP-15 data suggest that the timing of such approaches need consideration, as long-term senescent cells may be less malleable to proteostasis-boosting strategies—adding yet another layer of complexity to senotherapeutic development.

## METHODS

### Molecular cloning

All plasmid construction is described in the Supplementary Methods.

### Cell lines

IMR-90 human fetal lung fibroblasts at population doubling level (PDL) of 7.78 were acquired from Coriell Institute for Medical Research. A549 human lung adenocarcinoma cells (epithelial, alveolar type II) were purchased from ATCC. hTERT RPE-1 cells were a kind gift from René Medema and sent by Rene Medema and Rob Klompmaker (The Netherlands Cancer Institute, Netherlands). Generation of stable tdTomato−NLS, 6xHSE::mScarlet−NLS and Htt-exon1[polyQ] cell lines is described in the Supplementary Methods. DNAJC5^-/-^ and DNAJC5^-/-^::Flag–DNAJC5 A549 cell lines were gifted by Eric Faudry and Ina Attree (Institut de Biologie Structurale, Grenoble, France).^97^

All cell lines were regularly tested for Mycoplasma and authenticated by short-tandem repeat profiling (IDEXX Bioanalytics, Germany).

IMR-90, WI-38, and A549 cells were cultured in DMEM supplemented with 10 % fetal bovine serum, 2 mM L-Glutamine, 0.1 mM non-essential amino acids, and 100 Units of penicillin and streptomycin. RPE-1 cells were cultured in DMEM/F12 with the same supplements as above. All cells were cultured at 37 °C in a humidified incubator with 5 % CO_2_ and sub-cultured at 70–80 % confluency.

### Chemicals, compounds, and key reagents

Details of all chemicals and compounds used are included in the Key Resources Table.

### Senescence and quiescence induction

All senescence induction protocols were based on well-established methods from the literature. For DNA-damage (DD)-induced senescence with doxorubicin, proliferating cells were treated with 250 nM (IMR-90), 750 nM (A549), or 1 µM (RPE-1) doxorubicin for one day, followed by media changes every two to three days for a total of ten days.^132^ For DD-induced senescence with bleomycin, A549 cells flasks treated with 50 µg/mL bleomycin for five days, followed by five days with their regular complete media (media change at day 7 or 8).^133,134^ For oxidative stress–induced senescence, cells were treated with a 2 h pulse of 200 µM hydrogen peroxide (dissolved from a 9 M stock in the cell type’s regular complete growth media) every two days for a total of three treatments, followed by six days without hydrogen peroxide and media changes every three days. For replicative senescence, cells were serially passaged in their regular complete growth media until they reached their Hayflick Limit, initially determined *via* brightfield microscopy by an increase in mean cell area and a static total cell number over two weeks of culture, and confirmed by the battery of senescence assays described below. Unless otherwise stated, all states labelled ‘Senescent’ in the figures were induced by DNA damage, using the relevant protocol for the cell line.

Senescence was confirmed by a combination of higher SA-beta-galactosidase (SA-bG) activity, lack of EdU incorporation, larger nuclear area, and increase in number of 53-BP1 foci, as per recommendations from the senescence community.^135–137^

To establish contact-inhibited quiescence, proliferating cells were seeded at 30,000 cells/cm², and kept in the same culture vessel for ten days. For serum-starvation-induced quiescence, proliferating cells were seeded and allowed to adhere overnight in their complete growth media (with 10 % FBS), before replacement with media containing 0.2 % FBS for 10 (IMR-90) or 4 (RPE-1) days. For both quiescence models, media was changed on the same days as for the senescent models (i.e., every 2−3 days). Quiescence was confirmed by lack of EdU incorporation, and discriminated from senescence by the ability to re-enter the cell cycle upon return to sub-confluent culture conditions, as well as lack of nuclear size increase. Note that contact-inhibited quiescent states also stained positive for the SA-bG assay, as reported previously.^138^

Proliferating control cells were seeded at the same density as the senescent cells 24 h prior to subsequent assays.

### Senescence-Associated beta-Galactosidase activity assay

Cells were fixed in 4% paraformaldehyde, and senescence-associated beta-Galactosidase (SA-bG) activity staining was performed immediately (see Supplementary Methods for detailed protocol).^139^

### EdU cell proliferation assay

Cell cycle arrest was assessed by 10 µM EdU (Invitrogen; Cat# C10634) incorporation for 24 h, followed by detection using picolyl-azide-sulfo-Cy5 (Jena Biosciences, Germany; Cat# CLK-1177) (see Supplementary Methods for detailed protocol). For quiescence recovery experiments, cells were trypsinised, re-seeded at 25% confluency into a fresh culture vessel, and allowed to recover for two days before EdU addition.

### Nuclear area increase

Nuclear area was measured by Hoechst 33342 or DAPI staining and quantified using the ImageXpress Confocal HT.ai High-Content Imaging System (Molecular Devices, CA USA) and StarDist plugin (v0.3.0)^140^ with default settings in ImageJ (v1.54p).

### Immunofluorescence of DNA-damage foci

DNA-damage foci were visualised on fixed and permeabilised cells by immunofluorescence staining with rabbit anti–53-BP1 (1:500 dilution), and counter-staining with DAPI (1:1,000) and mouse anti–vimentin (1:500).

### Live-cell confluency monitoring

Cell confluency was quantified on an IncuCyte SX5 (Sartorius, Germany) set up inside a humidified incubator under standard cell-culture conditions, using the Incucyte AI Confluence analysis module with default settings.

### Heat-shock treatments

For heat-shock experiments, complete growth medium pre-warmed to the heat-shock temperature was added to cells immediately before movement into a humidified CO_2_ incubator set at the heat-shock temperature (most commonly 44 °C) for the allotted time, before proceeding to downstream assays. For longitudinal assays, IMR-90 cells expressing the tdTomato–NLS nuclear marker were seeding into clear, TC-treated flat-bottom 96-well microplates (Thermo; Cat# 167008), allowed to adhere overnight, and incubated at 44 °C to induce heat-shock. After the allotted heat-shock time, the cell-culture media was replaced with media containing 0.5 µM SYTOX Green dead-cell stain (Invitrogen; Cat# S7020) pre-warmed at 37 °C, and each well was imaged every 4 h on an Incucyte SX5 Live-Cell Analysis System (G/O/NIR filter set). Cell counts from the obtained images were calculated using the StarDist plugin (v0.3.0)^140^ with default settings in ImageJ (v1.54p). SYTOX-positive cells were excluded from live counts using manual intensity thresholding. Percentage viability was calculated by normalizing the cell count to the untreated (non-heat-shocked) control count at the start of imaging.

### Small-molecule proteotoxic stressor treatment

Cells were seeded in 96-well PhenoPlates (Revvity, UK; Cat# 6055300), allowed to adhere overnight, and incubated in pre-warmed media (37 °C) with bortezomib or tunicamycin for the indicated time. Mock vehicle-control treatments were performed in parallel for each small**-**molecule.

### High-content imaging−based live-cell quantification and analysis

Cells seeded in 96-well PhenoPlates, induced into the required cell states, and exposed to stresses (e.g., heat-shock; small-molecule treatment), were stained for 15 min with 10 µM Hoechst 33342 and 0.5 µM SYTOX Green or SYTOX Orange (Invitrogen, Cat# S34861) dead-cell stains, pre-warmed at 37 °C in their complete growth media. Staining media was replaced with fresh pre-warmed complete growth media before imaging in an ImageXpress Confocal HT.ai High-Content Imaging System. The plate schematics were mapped to the imaging system manually prior to image acquisition. Images were acquired using the x10 air objective, with 9 fields of view per well. Acquired images were analysed by CellProfiler/ImageJ. Calculated live-cell numbers were normalised as a percentage of either the mock-vehicle control at the same time-point as the treatments (e.g., t = 72 h), or the live-cell numbers at the starts of the experiment (t = 0 h). Final live-cell percentages were plotted on R using ggplot2, and dose-response curves and EC_50_ values were calculated using the drc package (v3.0-1).

### Transcriptomics and proteomics analyses

#### Experimental Design and Statistical Rationale

##### Choice of cell lines & model

DNA-damage-induced senescence was used in order to generate sufficient cellular material for all three dimensions of the proteomics sample preparation to be performed from the same batch. The DNA-damage agents doxorubicin and bleomycin were used according to well-established protocols in IMR-90 human fetal lung fibroblasts and A549 human lung adenocarcinoma cells, respectively.^132,133,141–144^

##### Heat-shock conditions

Heat-shock conditions were the same as used in a transcriptomics analysis of a similar human fetal lung fibroblast cell line (WI-38) in proliferating and replicatively-senescent states.^30^

##### Number of replicates

For IMR-90 cells, three biological replicates were prepared in order to enable all six conditions to be incorporated into a single TMT18-plex. Based on the variance detected in our previous experiments, we were confident three replicates would provide sufficient statistical power given the expected effect sizes. For A549 cells, where the quiescent state was not deemed relevant, four biological replicates were prepared. For each biological replicate, the treatments were always performed in parallel with the controls for each cellular state (i.e., Prolif.Basal_1, Prolif.HS_1, Quiesc.Basal_1, Quiesc.HS_1, Senesc.Basal_1, and Senesc.HS_1 were all performed in parallel).

##### Mass spectrometry–based proteomics strategy

Cell lysates from each sample were prepared for bottom-up proteomics via the glass bead–based SP4 method.^145^ Isobaric labelling by TMT18-plex was performed to minimise missing values and maximise quantitative comparisons between each sample. Spectra were analysed by MS^3^ to mitigate ratio compression.

### Heat-shock or bortezomib treatments

For the heat-shock experiments in IMR-90 and A549 cells, two 15 cm plates each of proliferating (∼ 80 % confluency) and contact-inhibited quiescent cells (IMR-90 cells only), and eight 15 cm plates of senescent cells (∼ 80 % confluency), were heat-shocked at 44 °C in a humidified incubator for 2 hours, per replicate. At the same time, the same number of plates representing the ‘Basal’ conditions were moved to a different humidified incubator set at 37 °C. After 2 h, cells were rinsed briefly three times with 20 mL PBS to remove excess serum proteins, and processed using denaturing or non-denaturing lysis conditions, depending on the downstream application. For the bortezomib experiment, A549 cells prepared in proliferating or senescent states as described above were treated with 100 nM bortezomib and incubated at 37 °C for 24 h, before proceeding to denaturing lysis.

### Total proteome sample preparation

Cells were scraped on ice in ∼4 pellet volumes of ice-cold denaturing lysis buffer (50 mM HEPES pH 8.0, 8 M urea, 1 % SDS, 1 % Triton X-100, 1 % IGEPAL CA-630, 1 % Tween 20, 1 % sodium deoxycholate, 150 mM NaCl) supplemented with 1 x Roche cOmplete EDTA-free Protease Inhibitor Cocktail, 1 mM PMSF, 40 mM 2-chloroacetamide, 10 µM PR-619, and 20 U/µL Pierce Universal Nuclease before snap-freezing in liquid nitrogen. Lysates were thawed on ice, sonicated on ice with a microtip probe (20 W, 12 cycles: 5 s on, 5 s off), and cleared by centrifugation (16,000 *g*, 4 °C, 5 min). Total protein concentration in each lysate was estimated by BCA assay, and adjusted to ∼6 mg/mL in more denaturing lysis buffer before snap-freezing again and storing at −70 °C.

Samples were prepared for bottom-up proteomics by SP4, as described previously.^145^ Briefly, 100 µg (IMR-90) or 50 µg (A549) of each sample frozen in denaturing buffer was thawed on ice, before being reduced with 10 mM TCEP (5 min, 80 °C) and 40 mM 2-chloroacetamide (15 min, room temperature). 1 mL of glass beads (Sigma Cat# 440345) suspended in acetonitrile (final bead concentration: 2.5 mg/mL) were initially washed with acetonitrile, 100 mM ammonium bicarbonate (ABC), and 2x with milliQ water, before resuspension in 100% acetonitrile. 4 volumes of these washed and resuspended beads were added to the reduced and alkylated cell lysates and mixed in a Thermomixer Comfort (5 s at 400 rpm) to precipitate proteins onto the beads. Precipitate-coated beads were pelleted (5 min, 16,000 *g*) and carefully washed (i.e., without resuspension) 3x with 80 % ethanol (with centrifugation for 2 min at 16,000 *g* in between each wash). Bead-bound precipitates were then digested in a Thermomixer Comfort (18 h, 1,000 rpm, 37 °C) with sequencing-grade modified trypsin (Promega Cat# V5111) at 1:100 enzyme:protein ratio in 50 mM HEPES pH 8.0. Glass beads were separated from the digested peptides by centrifugation for 5 min at 16,000 *g*, before proceeding to TMT labelling.

### Transcriptomics and non-denaturing cell lysate preparations

For transcriptome, polyUb–proteome, and insoluble proteome samples, adherent cells were collected after 3x PBS rinses by scraping in ice-cold PBS, split into two 15 mL falcon tubes, pelleted (200 *g*, 4°C, 5 min), and snap-frozen in liquid nitrogen.

RNA from one pellet was extracted in Trizol using standard protocols, and RNA quality was assessed using TapeStation (Agilent Technologies, CA USA), with all samples showing RNA Integrity Numbers > 9.5. Library preparation was performed at Cancer Research UK– Cambridge Institute Genomics Facility using the Illumina Stranded mRNA Kit, and 50 bp paired-end sequencing was conducted on a NovaSeq X, generating > 20 million reads per sample.

For polyUb–proteome and insoluble proteome analyses, the other snap-frozen cell pellets were initially homogenised, with all handling and buffers on ice, in approximately 5 cell pellet volumes of IP buffer (50 mM HEPES pH 8, 150 mM NaCl and 1 % IGEPAL) supplemented with 1x cOmplete EDTA-free Protease Inhibitor Cocktails, 1 mM PMSF, 40 mM 2-chloroacetamide, 10 μM PR-619, and 2.5 U/µL Pierce Universal Nuclease, and sonicated on ice with a microtip probe (20 W, 12 cycles: 5 s on, 5 s off). For polyUb–proteome enrichment, 250 µL of the sonicated lysates were adjusted to denaturing lysis buffer (50 mM HEPES pH 8, 150 mM NaCl, 8 M urea, 1 % SDS, 1 % Triton X-100, 1 % IGEPAL CA-630, 1% Tween 20, 1 % Na deoxycholate). For the pellet and supernatant isolation, homogenates were adjusted to a modified RIPA buffer (50 mM HEPES pH 8, 150 mM NaCl, 0.5 % SDS, 0.5 % Na deoxycholate, and 1 % IGEPAL CA-630). The protein concentration of the initial lysate was estimated by BCA from a SP4 protein clean-up and digestion of the samples in denaturing lysis buffer. Briefly, a 50 μL aliquot was diluted with 50 μL water, prior to SP4 processing and digestion with 1 µg of sequencing-grade modified trypsin for 18 h at 37 °C.

### PolyUb-enriched proteome sample preparation

20 µL (IMR-90) or 10 µL (A549) of HaloTag—[tev]—trTUBE_6_-conjugated magnetic beads (prepared as described in Supplementary Methods) were incubated with 500 μg (IMR-90) or 100 μg (A549) of cell lysate diluted in immunoprecipitation (IP) buffer (18 h, 4 °C, end-over-end rotation at 20 rpm). Beads were gathered by a magnet, washed 3x with 1 mL of TUBE wash buffer and another 3x with 1 mL 50 mM HEPES pH 8 (each wash with 30x end-over-end inversions at ∼30 rpm). For the final wash, beads were transferred to a new 1.5 mL low-bind microfuge tube, and fully aspirated with the aid of brief centrifugation at 2000 *g* for 15 s. Captured proteins were eluted from the beads by TEV protease-mediated cleavage of the trTUBE_6_ (0.1 µL (1 U) of TEV protease (NEB) in 20 µL 50 mM HEPES pH 8) and incubated in a Thermomixer Comfort (2 h, 30 °C, 1,200 rpm). Cleaved beads were gathered by a magnet, with the supernatant containing eluted proteins collected into a 0.5 mL low-bind microfuge tube and digested overnight in a Thermomixer Comfort (2 h, 1,000 rpm, 47 °C) with 50 ng of sequencing-grade modified trypsin in 50 mM HEPES pH 8.

### Insoluble proteome sample preparation

RIPA-adjusted cell lysate (0.6 µg/µL in 600 µL total volume) was first cleared of large debris by centrifugation (2000 *g*, 5 min, 4°C). 500 µL of the pre-cleared protein lysate were transferred to a new tube, and insoluble proteins enriched by centrifugation (16,000 *g*, 30 mins, 4°C). The pellet was washed 3x with 1mL RIPA buffer without resuspension, with centrifugation at 16,000 *g* for 10 mins at 4 °C between each wash. After the final wash was carefully aspirated, pellets were resuspended in 50 µL denaturing lysis buffer with 10 mM TCEP and 20 mM 2-chloroacetamide, sonicated in a water-bath for 15 min to aid resolubilisation, incubated on a Thermomixer Comfort (15 min, 1,200 rpm, 80 °C) for reduction, followed by 15 min at room temperature for alkylation. 10 µL of each sample was cleaned by SP4 and digested in a Thermomixer Comfort (18 h, 1,200 rpm, 37 °C) with 100 ng sequence-grade modified trypsin in 10 µL 50 mM HEPES pH 8.

### TMT labelling and peptide fractionation

TMTpro 18-plex labelling of trypsin-digested peptides was performed for 1 h in 50 mM HEPES pH 8.0, followed by quenching by adjusting to a final concentration of 0.4 % (v/v) hydroxylamine (Thermo Scientific, Cat# 90115) and incubating for 15 min at room temperature. At this stage, equal volumes of all samples were pooled together, and the multiplexed peptide mixture was dried in a Savant SpeedVac vacuum concentrator (Thermo Scientific). Dried peptides were resuspended in a solution of 2 % acetonitrile and 0.1 % triethylamine in Optima LC/MS-grade water (Fisher Chemical, Cat# W6-500), and fractionated with a Pierce High pH Reversed-Phase Peptide Fractionation Kit (Thermo; Cat# 84868), according to the manufacturer’s instructions. Peptides were resolubilised a solution of 5 % acetonitrile and 0.1 % formic acid in Optima LC/MS-grade water, for LC-MS injection.

For IMR-90 or A549 total proteome digests, 30 µg or 10 µg of peptides were labelled with 100 µg or 50 µg of TMTpro reagent, respectively, separated into 12 peptide fractions, and 4 µL of the 30 µL from each fraction analysed by LC-MS/MS.

For IMR-90 or A549 polyUb-enriched proteome digests, all peptide material was labelled with 100 µg or 25 µg of TMTpro reagent, separated into 8 or 4 peptide fractions, and 5 µL or 6 µL, respectively, of the 15 µL from each fraction analysed by LC-MS.

For IMR-90 insoluble proteome digests, all peptide material was labelled with 100 µg of TMTpro reagent, separated into 12 peptide fractions, and 3 µL of the 20 µL from each fraction analysed by LC-MS.

### LC-MS/MS and analysis of spectra

TMT-labelled high-pH peptide fractions were analysed by Orbitrap Eclipse MS (Thermo Scientific) with on-line separation on a reversed-phase nanoLC column (450 mm × 0.075 mm ID) packed with ReprosilPur C18AQ (Dr Maisch, 3 μm particles) at 40 °C. A 180 min (or 120 min for IMR-90 total proteome) gradient of 4–35% ACN, 0.1% FA at 300 nL/min was delivered via a Dionex UltiMate 3000 nanoHPLC system. Mass spectra were acquired in SPS MS^3^ mode and FAIMS (−45, −55, −65, and −75 V) with the following settings: MS^1^ —120k resolution, max IT 50 ms, AGC target 400,000; MS^2^ — IW 0.7 CID fragmentation, CE 35%, max IT 35 ms, ion trap rapid scan rate, AGC target 10,000; MS^3^ reporter quantitation —10 2 m/z notches, HCD fragmentation, CE 55%, 50k resolution, max IT 86 ms, AGC target 100,000.

The raw proteomics data have been deposited to the ProteomeXchange Consortium (IMR-90 heat-shock: PXD067225; A549 heat-shock: PXD067153; A549 bortezomib: PXD067220).

### Processing of raw spectrum files

Thermo raw spectrum files were processed with Proteome Discoverer (Thermo Scientific, v. 2.5.0.400) using Sequest HT (2.0.0.24, x64), Percolator (3.05.0), and IMP-ptmRS (1.0.0.0), searching against UniProt Human Swissprot database (UniProtKB 2023-06), supplemented with sequence for common contaminants,^146^ TEV protease, and HaloTag–[tev]–trTUBE_6_. In order to simplify protein-grouping of peptides belonging to ubiquitin, the polyubiquitin-B gene (UBB; P0CG47) was replaced with a single di-Ubiquitin sequence, and all other ubiquitin sequences with the relevant fusion genes (entire UBB gene, and N-terminal of UBA52 and RPS27A) within the search file were removed. The settings used for analysis were as follows: enzymes set as Trypsin/P, with a maximum of two missed cleavages; fixed modification was carbamidomethyl (Cys) and TMTpro (Lys, peptide N-term); variable modifications were acetyl, Met-loss, or Met loss+acetyl (protein N-term), and Oxidation (Met). For polyUb-enriched proteomes, TMTpro (Lys) and GlyGly+TMTpro (Lys) were set as additional variable modifications. Mass tolerances were set to 10 ppm (MS^1^, FTMS), 0.5 Da (MS^2^, ITMS), and 20 ppm (MS^3^, FTMS); false discovery rate (FDR) for both protein and peptide identifications was 0.01. For quantitation filtering, a co-isolation threshold of 50 % average reporter ion signal/noise >= 5 and SPS matches >= 50 % were applied.

### Processing of RNA-seq transcriptomics raw files

Total RNA was extracted using Trizol, following standard protocols. RNA quality was assessed using TapeStation, with all sample showing RIN > 9.5. Library preparation was performed using the Illumina Stranded mRNA Kit, and 50 bp paired-end sequencing was conducted on a NovaSeq X, generating > 20 million reads per sample. Reads were trimmed with TrimGalore v0.6.10 using default parameters. Trimmed reads were aligned to the Ensembl GRCh38 genome using hisat2 (v2.2.1), guided by introns extracted from Ensembl release 87. Hisat used default options plus --no-softclip, --nomixed, --no_discordant. Reads were quantitated using SeqMonk (v1.48.1) using human genome GRCh38 (v113) using the RNASeq quantitation to generate raw read counts for an opposing strand specific library.

### Initial data filtering and exploration

All initial data filtering, exploration, and statistical analyses of the protein-level data were performed using R (https://cran.r-project.org/, v4.3.1), unless otherwise stated. Detailed analysis notes, specific R packages, and R code used for initial analysis are included in Supplementary File 1.

For the IMR-90 multi-dimensional proteomics experiment, the exported csv file from the ProteomeDiscoverer analysis (Supplemnetary Table S1) was used as the input dataset for all downstream statistical analysis reported here. Of the 10,388 protein-group entries, we filtered out entries identified as potential contaminants, entries belonging to Y-chromosome genes (IMR-90 is a female-derived cell line), and entries that were only identified by a single peptide-spectrum match (PSM). The resulting data-frame of 8,669 proteins was used for downstream analysis.

The same filtering strategy was employed for the A549 multi-dimensional proteomics experiments, with the 9,200 or 9,541 initial protein-group entries from ProteomeDiscoverer (Supplementary Tables S6–S7) filtered down to 7,273 or 7,494 proteins for downstream analysis of the heat-shock or bortezomib experiments, respectively.

### Normalisation

For the total proteome data, the globally-normalised protein abundance values calculated from ProteomeDiscoverer were used from downstream analysis.^147^ For the polyUb and insoluble proteome data, global normalisation was deemed inappropriate, as global normalisation assumed relatively similar total protein amounts between samples. This would not be the case for the polyUb-enriched proteomes, as heat-shock is known to lead to global increase in polyUb chains,^9,55^ thus leading to more poly-ubiquitylated protein material to be captured for the heat-shocked samples than the non-heat-shocked samples, given equal total lysate protein input for polyUb enrichment. A similar rationale could apply to the insoluble proteome.^60,61^ Therefore, for these two proteome dimensions, a more conservative normalisation was performed through size-factor normalisation on the raw (unnormalised) protein abundance values, by defining the 10% least-changed proteins in each dimension (i.e., the proteins with the lowest 10% coefficients of variation (CVs)), calculating size-factors for each sample based on how much the median of the mean raw abundances for the 10% low-CV proteins in that sample deviated from the mean raw abundances of those proteins across all samples, and correcting the raw abundances for all proteins in that sample by the calculated size-factor.

### Missing values

Imputation was only performed in the globally-normalised total proteome data, and only for the proteins with a maximum of 6 missing values, with the R package DEP,^148^ using a manual left-censored Missing Not-At-Random method against a normal distribution with a left-shift of 1.8 and scale of 0.3.

### Differential analysis

The R package DEqMS was used for statistical analysis of differential abundances between cell states and treatment conditions.^149^ Differential proteins (DPs) were defined as those with adjusted p < 0.05 & absolute log_2_-transformed fold-changes (Log_2_FC) >= 1 for any selected pairwise comparison. Results from these analyses are available in Tables S2 (IMR-90), S8 (A549 heat-shock), and S9 (A549 bortezomib).

### Downstream analysis of transcriptomics and proteomics data

All downstream analysis of the proteomics data was performed in R (v4.5.0) with tidyverse (v2.0.0), using the packages and versions indicated below. Correlation heatmaps using ggcorrplot (v0.1.4.1); Heatmaps using pheatmap (v1.0.12); PCA using the built-in ‘prcomp’ function in base R; Over-representation analysis (ORA) using gprolifer2 (v0.2.3) with annotation pulled from org.Hs.eg.db (v3.20.0); Venn diagrams using a combination of VennDiagram (v.1.7.3) and eulerr (v7.0.2); all other plots with ggplot2 (v3.5.1), and statistical tests using rstatix (v0.7.2). Protein–protein interaction networks were generated with the STRING online database (https://string-db.org/, version 12),^150^ with the custom parameters and backgrounds as indicated in the relevant Figure Legends. Complete R packages and versions are listed in the Key Resources Table.

### Heat-Shock-Response reporter induction

IMR-90 cells expressing a fluorescent heat-shock-response reporter (6xHSEp::mScarlet–NLS) were subjected to heat-shock at 44 °C for 2 h, followed by return to 37 °C and imaging every 2 h on an Incucyte SX5 system. Confluency was quantified using the Incucyte AI Confluence analysis module with default settings. Fluorescence intensity was calculated after surface-fit background subtraction and normalized to confluency, expressed as the integrated signal per well.

### Reverse-Transcriptase quantitative PCR (RT-qPCR)

RNA was extracted from cells using the Phasemaker TRIzol Reagent and Phasemaker Tubes Complete System (Thermo, Cat# A33251), with GlycoBlue Coprecipitant (Thermo, Cat# AM9516) added to aid precipitant visualisation. To eliminate residual DNA, samples were treated with the Invitrogen DNA-free DNA Removal Kit (Thermo, Cat# AM1906). RNA was then reverse-transcribed into cDNA using the High-Capacity cDNA Reverse Transcription Kit (Thermo, Cat# 4368814) following the manufacturer’s protocol, with no-reverse-transcriptase controls included to confirm the absence of DNA contaminants. RT-qPCR was performed using iTaq Universal SYBR Green Supermix (Bio-Rad, Cat# 1725121) with 0.4 µM of primers for PUM1 (housekeeping gene^151^ that we determined to vary negligibly across all three cell states, and +/- heat-shock for 2 h at 44 °C) and HSPA1A on a CFX Opus 96 Real-Time PCR System (Bio-Rad, Cat# 12011319), with the following cycling conditions: (1) initial denaturation at 95 °C for 30 s, (2) denaturation at 95 °C for 5 s, (3) annealing at 58.4 °C (optimized for the primers used) for 30 s, and (4) repeat steps 2–3 for 39 cycles. Gene expression was quantified using the delta-delta CT method for relative quantification, and normalised to the PUM1 values for each sample. Primer sequences are listed in the Key Resources Table, were verified to target the desired gene products (on BLAST), produced a single melting curve, with efficiencies between 90–110 % and R^2^ > 0.9.

### High-content imaging and quantification of RBP foci

IMR-90 fibroblasts or RPE-1 epithelia in proliferating, contact-inhibited quiescent, or doxorubicin-induced senescent states were prepared on 96-well PhenoPlates as described earlier. After fixing with 4% paraformaldehyde in PBS for 10 min at room temperature, cells were washed 3x in cold PBS, permeabilised in 100 % methanol for 10 min at room temperature, and washed once more in cold PBS. Immunostaining for RBPs was performed with antibodies (see Key Resources table), all at 1:1,000 dilution. Imaging was performed on a PerkinElmer Opera Phenix High-Content Screening System (Revvity, UK) using a 40x water objective. Automated quantification of RBP foci quantification, and fluorescence intensity in the nucleus of cells, was performed using custom automated pipelines^67^ on Harmony 4.9 High-Content Imaging and Analysis Software (Revvity, UK). To calculate cytoplasmic area, images were pre-processed in ImageJ by subtracting background and enhancing local contrast, and cytoplasmic regions were segmented using the Trainable Weka Segmentation plugin (v4.0.0),^152^ trained to distinguish cytoplasm from background. Resulting binary masks were denoised using the ‘Remove Outliers’ function and used to quantify cytoplasmic area.

### Protein detection by SDS-PAGE and immunoblotting

For comparison of relative levels of proteins in cell lysates, cells were harvested as described for the total proteome sample preparation, except in denaturing lysis buffer without urea. 10 µg of protein (as estimated by BCA assay) were loaded per sample. For comparison of soluble vs. insoluble fractions, equal mass of BCA-estimated protein from each sample were enriched in parallel, followed by equivalent downstream processing for SDS-PAGE, regardless of differences in total mass of enriched protein between samples. For CM comparisons, CM loading was normalised to total protein mass of the cells from which the CM was collected, i.e., by harvesting the cells after CM removal in denaturing lysis buffer followed by BCA assay to calculate a scaling factor for each sample, and adjusting (with fresh complete medium) the volume of the CM from each sample by the scaling factor. After normalisation, CM was concentrated in a Pierce Protein Concentrator with 3-kDa Molecular-Weight-Cut-Off (Thermo Scientific; Cat# 88526) to ∼100 µL.

All samples were prepared for SDS-PAGE by dilution in 4x Protein Loading Buffer (125 mM Tris-HCl, pH 6.8, 50 % glycerol, 4 % SDS, 0.2 % (w/v) Orange G (Sigma; Cat# 861286)) with 100 mM DTT (final concentration), heated at 95 °C for 5 min, separated by SDS-PAGE in 10% Tris-Glycine gels, transferred onto 0.45 µm Immobilon-FL low-fluorescence PVDF (Sigma; Cat# IPFL00010), and blocked for 1 h at room temperature with TBS-T + 5 % Bovine Serum Albumin Fraction V (Melford, UK; Cat# A30075) (TBS-T/BSA). Primary antibody incubation (anti–DNAJC5: 1:1,000 dilution; anti–TDP-43: 1:3,000 dilution; anti–beta-tubulin: 1:4,000 dilution) was performed in TBS-T/BSA overnight at 4 °C, followed by 3x 5 min washes with TBS-T at room temperature, secondary antibody incubation (all at 1:10,000 dilution) in TBS-T/BSA for 1 h at room temperature. After 3 more washes with TBS-T at room temperature, immunoblots were imaged on a LiCor Odyssey CLx Imager (LI-COR Biotech, NE USA), and quantified using Image Studio (v6.0; LI-COR Biotech, NE USA).

### TDP-43 and GFP sandwich ELISAs

2 mL of CM from a single well of a 6-well plate was centrifuged (2,000 *g* for 10 min at 4 °C) and filtered through a 0.2 µm filter to remove dead cells and large debris. Vesicles within this filtered CM were disrupted by adjusting the CM to a final concentration of 1 % (v/v) Triton X-100, 0.5 % (w/v) sodium deoxycholate, and 0.5 % (w/v) SDS. The resultant CM was concentrated in a Pierce Protein Concentrator with 3-kDa Molecular-Weight-Cut-Off (Thermo Scientific; Cat# 88515) to 100 µL, and added directly to a single well of a 96-well assay plate from the Human TDP-43 ELISA Kit (Proteintech; Cat# KE00005), or the ChromoTek GFP-Trap Multiwell plate (Proteintech; Cat# gtp), and relative TDP-43 or GFP levels in the samples were quantified following the manufacturer-recommended sandwich-ELISA protocols.

### Imaging-based aggresome quantification

Cells seeded into 96-well PhenoPlates in proliferating, contact-inhibited quiescent, or DNA-damage-induced senescent states were treated for 24 h with proteasome inhibitors bortezomib or MG-132 (or DMSO vehicle) to induce misfolded protein accumulation. At the end of drug treatment, cells were fixed in 4 % formaldehyde at room temperature for 15 min, and permeabilised in 0.1 % Triton X-100 in ice-cold PBS for 5 min. Following 3x washes in ice-cold PBS, cells were incubated in antibody blocking solution (1% BSA in ice-cold PBS) for 1 h, and incubated with anti-Ub[K48]-JF549, anti-Ub[K63], and anti-Vimentin (all at 1:500 dilution) in antibody buffer (0.1% BSA in ice-cold PBS) overnight at 4 °C in the dark. Following 3x washes in ice-cold PBS, cells were incubated with anti-rabbit Alexa Fluor 647 and anti-mouse Alexa Fluor 488 secondary antibodies (both at 1:1,000 dilution) in antibody buffer at room temperature in the dark. Following 1x wash in ice-cold PBS, cells were stained with 0.5 µg/mL DAPI in antibody buffer for 10 min at room temperature in the dark, followed by 3x washes in ice-cold PBS. Cells were imaged immediately on the ImageXpress HT.ai using the 40x water-immersion lens, with 9 fields-of-view per well. A 23-step custom CellProfiler (v3.1.9)^153^ pipeline was used for automated aggresome quantification using a combination of all four channels, i.e., overlap between Ub[K48], Ub[K63], and vimentin signals at a juxtanuclear region (based on vicinity to the DAPI-based nuclear mask)(see Supplementary Methods).

### Live-cell imaging of GFP–Htt-exon1 A549 cells

For live-cell imaging, pTetO::GFP–Htt-exon1[Q25] or pTetO::GFP–Htt-exon1[Q97] A549 cells were established in proliferating or DNA-damage-induced senescent states in PhenoPlates before induction of Htt-exon1 expression by addition of 30 ng/mL doxycycline for 5 days (media change with fresh doxycycline on day 3). On day 5, media was replaced with fresh doxycycline-supplemented media with 100 nM Btz, and cells were imaged overnight (16 h) on an Olympus Evident SpinSR spinning-disk confocal microscope using a 40x air-immersion objective, with environmental controls set to 37 °C with 5 % CO_2_.

### siRNA-based DNAJC5 knockdown

25 nM of ON-TARGETplus Human siRNA SMARTPool targeting DNAJC5 (Horizon Discovery, UK; Cat# L-024098-01), or a Non-Target Control Pool (Horizon Discovery, UK; Cat# D-001810-10), were transfected into A549 cells seeded overnight into 15 cm dishes, using DharmaFECT 1 Transfection Reagent (Horizon Discovery, UK; Cat# T-2001) according to manufacturer’s recommended protocols, in Gibco Opti-MEM I Reduced Serum Medium (Thermo, Cat# 31985070), for 4 hours, followed by replacement with regular complete growth medium, and cultured for 3 days to allow DNAJC5 knock-down, prior to downstream experiments.

### Transient HSP72, HSP90, or HaloTag over-expression

20 µg of pCMV::mScarlet– HSPA1A, pCMV::Flag–HA–HSP90AB1, or pCMV::HaloTag, were transfected into A549 cells seeded overnight into 15 cm dishes, using Lipofectamine 2000 reagent (Thermo, Cat# 11668027) according to manufacturer’s recommended protocols, in Gibco Opti-MEM I Reduced Serum Medium, for 4 hours, followed by replacement with regular complete growth medium, and cultured for 3 days to allow over-expression, prior to downstream experiments.

## Supporting information

Key Resources Table

Stable S7

## ABBREVIATIONS

ALS: Amyotrophic Lateral Sclerosis
CID: Collision-Induced Dissociation
CORUM: Comprehensive Resource of Mammalian protein complexes
CDK: Cyclin-dependent kinase
CM: Conditioned Medium
DAPI: 4’, 6-diamidino-2-phenyindole
DDA: ata-Dependent Acquisition
DNAJC5/CSP: DnaJ homolog subfamily C member 5/Cysteine String Protein alpha
DP: Differential Protein
DMSO: Dimethyl Sulfoxide
EdU: 5-Ethynyl-2’-deoxyUridine
ELISA: Enzyme-Linked Immunosorbent Assay
EV: Extracellular Vesicle
FC: Fold Change
FDR: False Discovery Rate
FPKM: Fragments per Kilobase per Million
FTD: Fronto-Temporal Dementia
FUS: Fused in Sarcoma
G3BP1: Ras GTPase-Activating Protein–Binding Protein 1
GFP: Green Fluorescent Protein
GO: Gene Ontology
hnRNP: heterologous nuclear Ribo-Nucleo-Protein
HS: Heat-Shock
HSE: Heat-Shock Element
HSF1: Heat-Shock Factor protein 1
HSP: Heat-Shock Protein
HSR: Heat-Shock Response
hTERT: human Telomerase Reverse Transcriptase
Htt-ex1[Q25]: Huntingtin-exon1, with polyglutamine tract length of 25
Htt-ex1[Q97]: Huntingtin-exon1, with polyglutamine tract length of 97
IL-6: Interleukin-6
JUNQ: Juxtanuclear Quality Control Compartment
mRNA: messenger RNA
MATR3: Matrin 3
MTOC: Microtubule Organising Centre
NLS: Nuclear Localisation Sequence
ORA: Over-Representation Analysis
PCA: Principal Component Analysis
polyUb: poly-Ubiquitin
PQC: Protein Quality Control
RBP: RNA-Binding Protein
RIPA: Radio-Immuno-Precipitation Assay
RPE: Retinal Pigment Epithelia
RPM: Reads Per Million
rRNA: ribosomal RNA
RT-qPCR: Reverse-Transcriptase quantitative Polymerase Chain Reaction
SA-bG: Senescence-Associated beta-Galactosidase
SASP: Senescence-Associated Secretory Phenotype
SDS: Sodium Dodecyl Sulfate
sEV: small Extracellular Vesicle
SG: Stress Granule
STRING: Search Tool for the Retrieval of Interacting Genes
TARDBP/TDP-43: Trans-Active Response DNA-Binding Protein, 43 kDa
TMT: Tandem Mass Tag
TRiC/CCT: T-complex Ring Complex/Chaperonin Containing TCP-1
trTUBE: trypsin-resistant Tandem-Ubiquitin-Binding-Entity
Ub[K48]: Lys48-linked Ubiquitin
Ub[K63]: Lys63-linked Ubiquitin
YFP: Yellow Fluorescent Protein

## ACKNOWLEDGEMENTS

We thank Ritwick Sawarkar and Mengjia Li for processing of samples for transcriptomics (MRC-Toxicology Unit, Cambridge, UK); BI Proteomics Facility for all mass-spectrometry runs; Eric Faudry and Ina Attree (Institut de Biologie Structurale, Grenoble, France) for DNAJC5 knock-out and over-expressing A549 cells; Vincent Masto and Judith Frydman (Stanford University, CA USA) for plasmids containing Htt-exon1[Q25] and Htt-exon1[Q97] sequences for sub-cloning.

RSS and HEJ were supported by BI’s BBSRC Institute Strategic Programmes Grant for Signalling (BB/P013384/1 and BB/Y006925/1); HO, HC, SW, and SA by BI’s BBSRC Core Capability Grant (BB/CCG2310/1); TW by a BBSRC Training Grant (BB/X511559/1); YAM by a Training Grant from BI and Cambridge University School of Biological Sciences via The Vice-Chancellor’s Fund; EW by a BI Science Policy Committee Project Grant; RIO by the Cambridge University Hospitals NHS Foundation Trust. This work was also supported by a BBSRC Institute Development Grant (BB/IDG2310/1).

