## Supplementary File S1 for "Declining intracellular proteostasis capacity drives misfolded protein secretion in senescent human cells"

#### Processing of multi-dimensional proteomics data

**References** **85**

### 1 Introduction

We have performed several multi-omics experiments to determine whether there is a difference in the proteotoxic stress responses of cells in different states: proliferating, quiescent, or senescent.

For IMR-90 normal human lung fibroblasts (NHLFs), we have all three cell states—proliferating, contact-inhibited quiescent, and DNA-damage-induced senescent—with heat-shock as the proteotoxic stress (2 hours at 44 ◦C). For A549 lung epithelial adenocarcinoma cells, we only have two states—proliferating and DNA-damage-induced senescent—but one additional proteotoxic stress: treatment with the clinically-approved proteasome inhibitor bortezomib (100 nM for 24 h).

```r
knitr::include_graphics("../experimentalDesign.png")
```

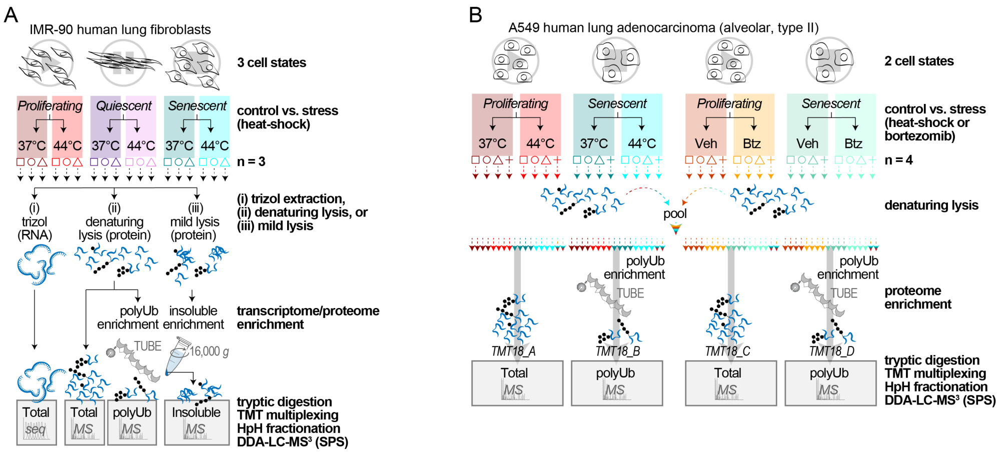

Figure 1: Experimental design for multi-omics characterisation of proteoxic stress responses in IMR-90 human fetal lung fibroblasts (A) or A549 lung epithelial adenocarcinoma (B), across different cellular states.

This document describes the processing steps applied to all of the proteomics data across both cell lines. The RNA-seq data were processed by Simon Andrews (Bioinformatics Group, Babraham Institute), as is standard practice for all sequencing data at Babraham Institute.

The starting point for this analysis is the ProteomeDiscoverer (PD) protein-level output from Harvey Johnston (Post-doctoral Research Associate, Signalling Programme, Babraham Institute). Details of prior sample preparation and processing are described in the Methods.

### 2 IMR-90 Heat-Shock Experiment

#### 2.1 Initial Data Clean-Up and Filtering

```r
read_csv("../../IMR90_PSQ_HS_TotUbiIns/data.csv") -> data
dim(data)
```

```
## [1] 10388   257
```

The initial data-frame has 10,388 entries (i.e., proteins identified in the experiment), and 257 columns, containing various data and metadata.

The protein-level "abundance" quantitations are based on relative TMT signal:noise ratios for each channel. Both raw and normalised (using the default global normalisation function in PD) abundances are included.

```
# find abundance columns
grep("^Abundance:", names(data), value = TRUE) %>%
  head()
```

```
## [1] "Abundance: F1: 126, Sample, 01_P37"  "Abundance: F1: 127C, Sample, 01_P37"
## [3] "Abundance: F1: 128C, Sample, 01_P37" "Abundance: F1: 129C, Sample, 02_S37"
## [5] "Abundance: F1: 130C, Sample, 02_S37" "Abundance: F1: 131C, Sample, 02_S37"
```

```
# find normalised abundance columns
grep("(Normalized)", names(data), value = TRUE) %>%
  head()
```

```
## [1] "Abundances (Normalized): F1: 126, Sample, 01_P37"
## [2] "Abundances (Normalized): F1: 127C, Sample, 01_P37"
## [3] "Abundances (Normalized): F1: 128C, Sample, 01_P37"
## [4] "Abundances (Normalized): F1: 129C, Sample, 02_S37"
## [5] "Abundances (Normalized): F1: 130C, Sample, 02_S37"
## [6] "Abundances (Normalized): F1: 131C, Sample, 02_S37"
```

There are 54 abundance columns: one for each TMT18-plex sample, across total (F1), polyUb (F2), and insoluble (F3) proteomes.

We can use these values to create our experimental_design, which will be important for categorising samples.

```
# create experimental design
data %>%
  dplyr::select(starts_with("Abundance:")) %>%
  colnames() %>%
  enframe(name = NULL, value = "sample") %>%
  mutate(sampleID=str_replace(sample,"[tpu]$","")) %>%

  separate(sampleID,
           into = c("temp1", "temp2", "TMT_temp3_Con"), sep = ":", remove = F) %>%

  mutate(TMT_temp3_Con = str_trim(TMT_temp3_Con, side = "both")) %>%

  separate(`TMT_temp3_Con`, into = c("TMT", "temp3", "condition"), sep = ",") %>%

  dplyr::select(-temp1, -temp2, -temp3) %>%

  dplyr::mutate(condition = substr(condition, 5, nchar(condition))) %>%

  mutate(`condition` = str_sub(`condition`, 1, 3)) %>%

  dplyr::mutate(condition = str_replace(condition, "P", "Prolif.")) %>%
  dplyr::mutate(condition = str_replace(condition, "Q", "Quiesc.")) %>%
  dplyr::mutate(condition = str_replace(condition, "S", "Senesc.")) %>%

  dplyr::mutate(condition = str_replace(condition, "37", "Basal")) %>%
  dplyr::mutate(condition = str_replace(condition, "44", "HeatShock")) %>%
```

```
  separate(condition, into = c("state", "treatment"), remove = F) -> experimental_design

nreps <- 3
experimental_design$replicate <- rep(c(1:nreps), (nrow(experimental_design)/nreps))

experimental_design %>%
  unite("label", c(condition, replicate), sep = "_", remove = F) -> experimental_design

experimental_design %>%
  write_tsv("IMR90/experimental_design.txt")

head(experimental_design)
```

```
## # A tibble: 6 x 8
##   sample                sampleID TMT   label condition state treatment replicate
##   <chr>                 <chr>    <chr> <chr> <chr>     <chr> <chr>         <int>
## 1 Abundance: F1: 126, ~ Abundan~ 126   Prol~ Prolif.B~ Prol~ Basal             1
## 2 Abundance: F1: 127C,~ Abundan~ 127C  Prol~ Prolif.B~ Prol~ Basal             2
## 3 Abundance: F1: 128C,~ Abundan~ 128C  Prol~ Prolif.B~ Prol~ Basal             3
## 4 Abundance: F1: 129C,~ Abundan~ 129C  Sene~ Senesc.B~ Sene~ Basal             1
## 5 Abundance: F1: 130C,~ Abundan~ 130C  Sene~ Senesc.B~ Sene~ Basal             2
## 6 Abundance: F1: 131C,~ Abundan~ 131C  Sene~ Senesc.B~ Sene~ Basal             3
```

##### 2.1.1 Clean-up of Gene and Accession identifiers

**2.1.1.1 Filling in missing Gene Symbols** There are almost always missing Gene Symbols from a PD protein-level output (i.e., UniProt Accessions that were not matched to a Gene name).

```
data %>%
  rename("Gene" = `Gene Symbol`) -> data

data %>%
  group_by(is.na(Gene)) %>%
  dplyr::count()
```

```
## # A tibble: 2 x 2
## # Groups:   is.na(Gene) [2]
##   `is.na(Gene)`     n
##   <lgl>         <int>
## 1 FALSE         10321
## 2 TRUE             67
```

We can write a function to attempt to assign these 67 missing Genes automatically from UniProt.

```
# write a function to get the Gene name from uniprot
 get_gene_from_uniprot <- function(accession) {
   uniprot_text <- NA
   tryCatch({scan(paste0("http://rest.uniprot.org/uniprotkb/", accession, ".txt"),
          what = character(), sep = "\n", quiet = TRUE) -> uniprot_text},
     warning = function(x){}, error = function(x){})
   if (any(is.na(uniprot_text))) {return(NA)}
```

```r
  grep("^GN", uniprot_text, value = TRUE)[1] -> gene_name
  str_replace(gene_name, ";.*", "") -> gene_name
  str_replace(gene_name, "^.*=", "") -> gene_name
  str_replace(gene_name, " .*", "") -> gene_name
  return(gene_name)
}

# use get_gene_from_uniprot
data %>%
  mutate(Gene = replace(Gene, is.na(Gene), sapply(
    Accession[is.na(Gene)], get_gene_from_uniprot))) -> data

# find number of missing Genes remaining
data %>%
  group_by(is.na(Gene)) %>%
  dplyr::count()
```

```
## # A tibble: 2 x 2
## # Groups:   is.na(Gene) [2]
##   `is.na(Gene)`     n
##   <lgl>         <int>
## 1 FALSE         10374
## 2 TRUE             14
```

These 14 remaining missing Gene names need to be annotated manually.

```r
data %>%
  dplyr::filter(is.na(Gene)) %>%
  dplyr::select(Accession, Description)
```

```
## # A tibble: 14 x 2
##    Accession       Description
##    <chr>           <chr>
##  1 Cont_X00000     Halo-TR-TUBE protein OS=Escherichia coli OX=0000 GN=HaloTEVT~
##  2 Cont_P00761     Trypsin OS=Sus scrofa OX=9823 PE=1 SV=1
##  3 Q6ZSR9          Uncharacterized protein FLJ45252 OS=Homo sapiens OX=9606 PE=~
##  4 Cont_G5E513     Uncharacterized protein OS=Bos taurus OX=9913 PE=1 SV=2
##  5 Cont_A0A3Q1M3L6 Uncharacterized protein OS=Bos taurus OX=9913 PE=1 SV=1
##  6 Cont_A0A3Q1M032 Uncharacterized protein OS=Bos taurus OX=9913 PE=1 SV=1
##  7 Cont_P50448     Factor XIIa inhibitor OS=Bos taurus OX=9913 PE=1 SV=1
##  8 Cont_Q29437     Primary amine oxidase, liver isozyme OS=Bos taurus OX=9913 P~
##  9 Cont_P00767     Chymotrypsinogen B OS=Bos taurus OX=9913 PE=1 SV=1
## 10 Cont_Q1A7A4     Complement component C5a (Fragment) OS=Bos taurus OX=9913 PE~
## 11 A0A8V8TPW1      Uncharacterized protein OS=Homo sapiens OX=9606 PE=1 SV=2
## 12 Q8NFD4          Uncharacterized protein FLJ76381 OS=Homo sapiens OX=9606 PE=~
## 13 A0A8V8TPP0      Uncharacterized protein OS=Homo sapiens OX=9606 PE=4 SV=1
## 14 Q6ZQT0          Putative uncharacterized protein FLJ45035 OS=Homo sapiens OX~
```

Most of the missing Gene names are contaminants. For these, we will copy the Accession into the Gene column.

```r
data %>%
  mutate(Gene = case_when(grepl("^Cont_", Accession) ~ Accession,
                          TRUE ~ Gene)) -> data

data %>%
  dplyr::filter(is.na(Gene)) %>%
  dplyr::select(Accession, Description)
```

```
## # A tibble: 5 x 2
##   Accession  Description
##   <chr>      <chr>
## 1 Q6ZSR9     Uncharacterized protein FLJ45252 OS=Homo sapiens OX=9606 PE=2 SV=2
## 2 A0A8V8TPW1 Uncharacterized protein OS=Homo sapiens OX=9606 PE=1 SV=2
## 3 Q8NFD4     Uncharacterized protein FLJ76381 OS=Homo sapiens OX=9606 PE=2 SV=1
## 4 A0A8V8TPP0 Uncharacterized protein OS=Homo sapiens OX=9606 PE=4 SV=1
## 5 Q6ZQT0     Putative uncharacterized protein FLJ45035 OS=Homo sapiens OX=9606 ~
```

For the remaining missing Genes, we will manually fill these in by searching for them on UniProt, or by adding the symbols we want.

```r
data %>%
  mutate(Gene = replace(Gene, Accession == "A0A8V8TPP0", "A0A8V8TPP0")) %>%
  mutate(Gene = replace(Gene, Accession == "A0A8V8TPW1", "A0A8V8TPW1")) %>%
  mutate(Gene = replace(Gene, Accession == "Q6ZQT0", "FLJ45035")) %>%
  mutate(Gene = replace(Gene, Accession == "Q6ZSR9", "FLJ45252")) %>%
  mutate(Gene = replace(Gene, Accession == "Q8NFD4", "FLJ76381")) -> data

data %>%
  dplyr::filter(is.na(Gene)) %>%
  dplyr::count()
```

```
## # A tibble: 1 x 1
##       n
##   <int>
## 1     0
```

After annotating all the missing Genes, we will also change the name of the polyUbiquitin gene (currently "Ubiqp; Ubiqp_0; Ubiqp_1") to "UBB". Note that the customised FASTA file used for the PD search had removed the ubiquitin sequence from all other genes (UBA52, RPS27A, and UBC), so all tryptic peptides matching the ubiquitin amino-acid sequence will have been grouped to this single Gene.

```r
data %>%
  mutate(Gene = replace(Gene, Accession == "P0CG47", "UBB")) -> data
```

**2.1.1.2 Remove Y-chromosome genes** Due to the nature of protein grouping, non-unique peptides can be assigned to Y-chromosome genes—even in cases where the cells are all female (as in IMR-90 cells). To simplify the analysis (and remove these false-assignments), we will remove all Y-chromosome Gene entries from the data.

We can annotate the chromosome locations of all the Genes in our data. Note that connecting to the ensembl mirror using bioMart is not always possible, so we will instead use a csv file we downloaded containing a list of all the Y-chromosome genes from bioMart.

```r
# Uncomment if using bioMart (i.e., if we don't have a saved list of genes)
# library(biomaRt)
#
# # Define biomart object
# mart <- useEnsembl(biomart="ensembl", dataset="hsapiens_gene_ensembl", mirror = "www")
#
# normal.chroms <- c(1:22, "X", "Y", "M")
#
# # Filter on HGNC symbol and chromosome
# my.symbols <- data$Gene
#
# my.regions <- getBM(c("hgnc_symbol", "chromosome_name"),
#                      filters = c("hgnc_symbol", "chromosome_name"),
#                      values = list(hgnc_symbol=my.symbols, chromosome_name=normal.chroms),
#                      mart = mart)
#
# # Filter only the Y chromosome genes
# my.regions %>%
#  dplyr::filter(chromosome_name=="Y") %>%
#  dplyr::select(geneName) %>%
#  distinct() %>%
#  pull(geneName) -> Ygenes

# If loading straight from a local csv file
# Load list of genes
Ygenes <- read_csv("../../ygenes.csv") %>%
  dplyr::select(geneName) %>%
  distinct() %>%
  pull(geneName)

# Filter the Ygenes that were actually in our data
data %>%
  dplyr::filter(Gene %in% Ygenes) %>%
  pull(Gene)
```

```
##  [1] "DDX3Y"                "USP9Y"                "SLC25A6"
##  [4] "ASMTL"                "DHRSX"                "RPS4Y1"
##  [7] "EIF1AX"               "KDM5D"                "EIF1AY"
## [10] "VAMP7"                "AKAP17A"              "ZBED1"
## [13] "CD99"                 "NLGN4Y"               "GTPBP6"
## [16] "PPP2R3B"              "PLCXD1"               "BPY2; BPY2B; BPY2C"
```

Some of these genes can actually be X-chromosome variants. We will only remove the proteins representing genes that can only be present on the Y chromosome.

```r
data %>%
  dplyr::filter(!Gene %in% c("DDX3Y", "RPS4Y1", "USP9Y", "EIF1AY", "KDM5D", "NLGN4Y")
                ) -> data
```

**2.1.1.3 Making duplicated Genes unique** Many of the downstream processing and plotting steps can't handle duplicated Genes; so these need to be edited.

```r
# Check for duplicated Accessions or Genes
data %>%
  group_by(Accession) %>%
  filter(n()>1) %>%
  dplyr::count()
```

```
## # A tibble: 0 x 2
## # Groups:   Accession [0]
## # i 2 variables: Accession <chr>, n <int>
```

```r
data %>%
  group_by(Gene) %>%
  filter(n()>1) %>%
  dplyr::count()
```

```
## # A tibble: 2 x 2
## # Groups:   Gene [2]
##   Gene            n
##   <chr>       <int>
## 1 CDKN2A          2
## 2 PALM2AKAP2      2
```

There are no duplicated Accessions, but there are 2 duplicated Genes. We need to make these unique.

```r
data %>%
  dplyr::group_by(Gene) %>%
  dplyr::count() %>%
  ungroup() %>%
  filter(n>1) %>%
  arrange(desc(n)) %>%
  left_join(data %>% dplyr::select(Accession, Gene, `# Peptides`))
```

```
## # A tibble: 4 x 4
##   Gene            n Accession `# Peptides`
##   <chr>       <int> <chr>            <dbl>
## 1 CDKN2A          2 P42771               7
## 2 CDKN2A          2 Q8N726               2
## 3 PALM2AKAP2      2 Q9Y2D5              37
## 4 PALM2AKAP2      2 Q8IXS6              17
```

We will distinguish between these manually, changing the name of the Gene with the fewer peptide identifications ("# Peptides").

```r
data %>%
  mutate(Gene = replace(Gene, Accession == "Q8N726", "ARF")) %>%
  mutate(Gene = replace(Gene, Accession == "Q8IXS6", "PALM2")) -> data

data %>%
  group_by(Gene) %>%
  filter(n()>1) %>%
  dplyr::count()
```

```
## # A tibble: 0 x 2
## # Groups:   Gene [0]
## # i 2 variables: Gene <chr>, n <int>
```

Excellent, now both Accession and Gene entries are unique.

##### 2.1.2 Adding proteome annotations

First, we must convert the data into long format, and add another column to specify whether the abundances are raw or globally-normalised values.

```
data %>%
  dplyr::select(Accession, Gene, Description, Contaminant,
         starts_with("Abundance:"), starts_with("Abundances (Normalized)")) %>%
  distinct(Accession, .keep_all = TRUE) %>%
  filter(!is.na(Accession)) %>%
  pivot_longer(
    cols = -c(Accession, Gene, Description, Contaminant),
    names_to = "sample",
    values_to = "abundance") -> dataLong

dataLong %>%
  dplyr::mutate(normalisation =
                  case_when(
                    grepl("Normalized", sample) ~ "normGlobal",
                    .default = "raw")) %>%

  mutate(across(
    'sample', str_replace, 'Abundances \\(Normalized\\)', 'Abundance')) -> dataLong
```

We will also name the different proteomes based on the “F” annotations in the “sample” column, and then add the sample annotations from experimental_design.

```
dataLong %>%
    mutate(proteome = case_when(
      str_detect(sample, "F1") ~ "total",
      str_detect(sample, "F2") ~ "polyUb",
      str_detect(sample, "F3") ~ "insoluble"
    )) %>%
    left_join(experimental_design) %>%
  dplyr::select(-sampleID) -> dataLong
```

##### 2.1.3 Removing contaminants

Next, we will remove contaminants.

```
# count contaminants
dataLong %>%
  group_by(Contaminant) %>%
  distinct(Accession) %>%
  dplyr::count()
```

```
## # A tibble: 2 x 2
## # Groups:   Contaminant [2]
##   Contaminant     n
##   <lgl>       <int>
## 1 FALSE       10204
## 2 TRUE          178
```

Let's plot these 178 contaminants, to see how abundant they are across our samples.

```r
# plot contaminants
dataLong %>%
  ggplot(aes(x = `label`, y = log2(`abundance`))) +
  geom_jitter(alpha = 0.05, size = 0.5) +
  geom_jitter(data = ~filter(.x, Contaminant == "TRUE"),
              colour = "red", size = 0.5, alpha = 0.5) +
  ggtitle("Abundance of Contaminants") +
  coord_flip() +
  theme(axis.title.y = element_blank()) +
    facet_grid(normalisation ~ proteome)
```

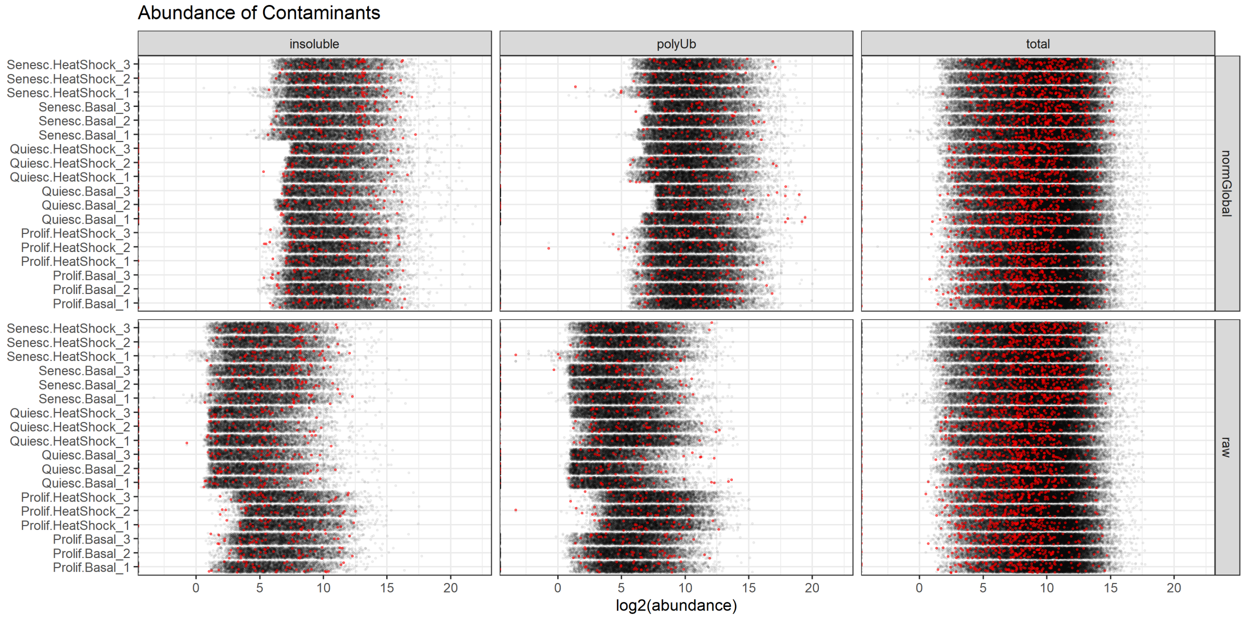

Figure 2: The raw abundance distribution of known contaminants in the IMR-90 dataset.

Based on these plots, it doesn't seem like there is any clear bias in the distribution of contaminants between the samples. We can safely remove these contaminants at this stage.

```r
# remove contaminants
dataLong %>%
  filter(!Contaminant) %>%
  dplyr::select(-Contaminant) -> dataLong

dataLong %>%
  unite(norm_proteome_label, c(normalisation, proteome, label), sep = "_") %>%
```

```
  dplyr::select(Gene, Accession, norm_proteome_label, abundance) %>%
  pivot_wider(names_from = norm_proteome_label, values_from = abundance) -> dataNoConts
```

##### 2.1.4 Dealing with single PSM/peptide identifications

The quantification of proteins identified by a single peptide is likely to be noisy and unreliable. But removing all such proteins could result in biasing data (e.g., against very short proteins that are more likely to have single quantifiable peptides).

The DEqMS differential analysis pipeline (Zhu et al. 2020) takes into account the number of peptides (or peptide-spectrum matches, PSMs) identified as an extra variable, thus allowing us to keep data on large and consistent differences across samples, even when only a single peptide was used for the quantitation.

Nevertheless, we should have some level of filtering. In this case, as we have three proteomes, we can make the decision that proteins identified by multiple PSMs across the proteomes should be kept, i.e., only those proteins that had a single PSM even when the quantitations from all three proteomes are combined should be removed as very low confidence identifications.

```
data %>%
  dplyr::select(Gene, Accession, `# Peptides`, `# PSMs`) %>%
  right_join(dataNoConts) -> dataNoConts

# Plot number of proteins identified by a single PSM
dataNoConts %>%
  dplyr::rename(numPSMs = `# PSMs`) %>%
  group_by(numPSMs > 1) %>%
  dplyr::count() %>%
  ungroup() %>%
  dplyr::filter(`numPSMs > 1` == FALSE) %>%
  dplyr::select(`n`) -> singletonNumber

dataNoConts %>%
    ggplot(aes(x = `# PSMs`)) +
    geom_bar(fill = "grey", colour = "black", size = 0.2) +
    geom_bar(data = ~filter(.x, `# PSMs` == 1), fill = "red", colour = "black") +
    coord_cartesian(xlim = c(0, 200)) +
  annotate("text", label = paste0(
    singletonNumber, " proteins identified by a single PSM"),
    x = 80, y = 380, color = "red", size = 4) +
    xlab("number of unique PSMs") +
  ylab("count(Proteins)") -> plot_numPSMs

# Count number of proteins identified by a single peptide
dataNoConts %>%
  dplyr::rename(numPeptides = `# Peptides`) %>%
  group_by(numPeptides > 1) %>%
  dplyr::count() %>%
  ungroup() %>%
  dplyr::filter(`numPeptides > 1` == FALSE) %>%
  dplyr::select(`n`) -> singletonNumber

dataNoConts %>%
    ggplot(aes(x = `# Peptides`)) +
```

```
    geom_bar(fill = "grey", colour = "black", size = 0.2) +
    geom_bar(data = ~filter(.x, `# Peptides` == 1), fill = "red", colour = "black") +
    coord_cartesian(xlim = c(0, 200)) +
  annotate("text", label = paste0(
    singletonNumber, " proteins identified by a single peptide"),
    x = 80, y = 600, color = "red", size = 4) +
    xlab("number of unique Peptides") +
  ylab("count(Proteins)") -> plot_numPeptides

plot_numPSMs + plot_numPeptides +
  plot_layout(nrow = 2) + plot_annotation(tag_levels = 'A')
```

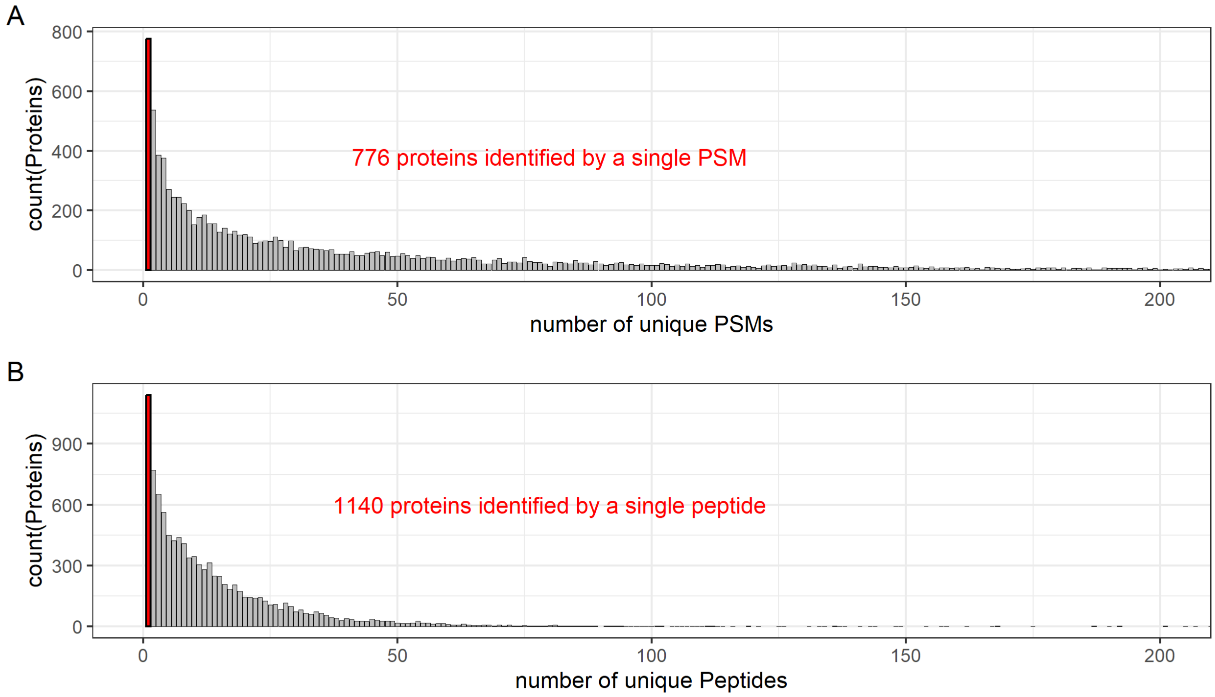

Figure 3: Histogram showing distribution of proteins identified by the number of unique PSMs (A) or peptides (B) in the IMR-90 dataset.

After removing these proteins identified by a single PSM, we can count how many proteins identified with single peptides remain.

```
dataNoConts %>%
  filter(`# PSMs` > 1) -> dataNoContsNoSinglePSMs

# Replot peptides histogram
# Count number of proteins identified by a single peptide
dataNoContsNoSinglePSMs %>%
  dplyr::rename(numPeptides = `# Peptides`) %>%
  group_by(numPeptides > 1) %>%
  dplyr::count() %>%
  ungroup() %>%
  dplyr::filter(`numPeptides > 1` == FALSE) %>%
```

```
  dplyr::select(`n`) -> singletonNumber

dataNoContsNoSinglePSMs %>%
  ggplot(aes(x = `# Peptides`)) +
  geom_bar(fill = "grey", colour = "black", size = 0.2) +
  geom_bar(data = ~filter(.x, `# Peptides` == 1), fill = "red", colour = "black") +
  annotate("text", label = paste0(
    singletonNumber, " proteins remaining identified by a single peptide"),
           x = 100, y = 380, color = "red", size = 4) +
  coord_cartesian(xlim = c(0, 200)) +
  xlab("number of unique Peptides") +
ylab("count(Proteins)") +
  ggtitle("Number of observed peptides (after removal of singleton-PSM proteins)")
```

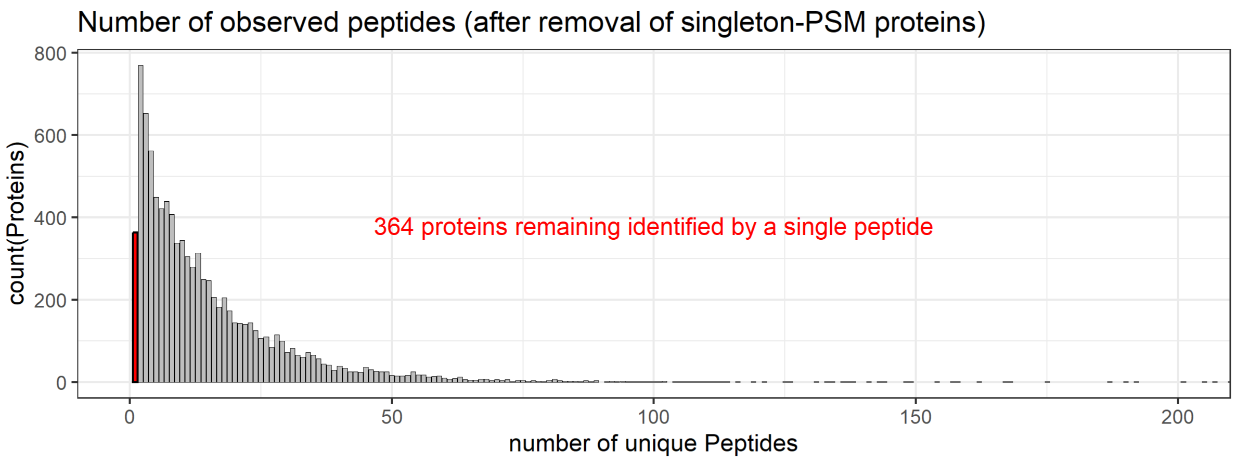

Figure 4: Histogram showing distribution of proteins identified in the IMR-90 experiment by the number of unique peptides, after the 776 proteins identified by single PSMs had been removed.

This data-frame will be saved as **dataFiltered.csv**.

```
dataNoContsNoSinglePSMs %>%
  dplyr::select(-`# PSMs`, -`# Peptides`) %>%

  write_csv("IMR90/dataFiltered.csv",
            na = "NA", append = FALSE, col_names = TRUE, escape = "double")

dim(dataNoContsNoSinglePSMs)
```

```
## [1] 9428  112
```

#### 2.2 Data Normalisation

##### 2.2.1 Assessing global normalisation

As discussed elsewhere (Palomba et al. 2021), the PD global normalisation is generally sufficient for most total proteome experiments, where roughly equal loading is expected between all samples. However, in this experiment, there are at least two proteomes where equal loading would not be expected.

For example, after heat-shock, there is a global increase in poly-ubiquitylation; therefore, from an equal amount of initial starting cell material (in this case, 0.5 mg protein, as estimated by BCA and then adjusted using the total proteome signal), we would expect far more poly-ubiquitylated material captured for the heat-shocked samples. Any global normalisation at the level of the quantified proteome would mask such differences—and perhaps create artefacts.

Let's visualise the total protein abundances across the 18 samples for each of the three proteomics experiments.

We will first create a custom colour scale, which will be used throughout this analysis.

```
six_sample_colours <- c('red4', 'red', 'purple4', 'violet', 'turquoise4', 'cyan')

# a function 'col2hex' to find the HEX codes for these colours
# (e.g., for replicating them in Illustrator)
col2hex <- function(x, alpha = FALSE) {
  args <- as.data.frame(t(col2rgb(x, alpha = alpha)))
  args <- c(args, list(names = x, maxColorValue = 255))
  do.call(rgb, args)
}

col2hex(six_sample_colours)
```

```
##       red4        red    purple4     violet turquoise4       cyan
##  "#8B0000"  "#FF0000"  "#551A8B"  "#EE82EE"  "#00868B"  "#00FFFF"
```

```
dataNoContsNoSinglePSMs %>%
  dplyr::select(-`# Peptides`, -`# PSMs`) %>%
  pivot_longer(cols = -c(Gene, Accession),
               names_to = "norm_proteome_label",
               values_to = "abundance") %>%
  separate(norm_proteome_label,
           into = c("normalisation", "proteome", "state.treatment", "rep"),
           sep = "_") %>%
  unite(state.treatment_rep,
        c(`state.treatment`, `rep`),
        sep = "_", remove = FALSE) %>%
  separate(state.treatment, into = c("state", "treatment"),
           sep = "\\.", remove = FALSE) -> dataLong

dataLong %>%
  ggplot(aes(x = state.treatment_rep, y = log2(abundance), fill = state.treatment)) +
  geom_boxplot(outlier.size = 0.4, linewidth = 0.2) +
  coord_flip() +
  scale_fill_manual(values = six_sample_colours) +
  theme(axis.title.y = element_blank(),
        axis.text.y = element_text(size = 6),
        axis.ticks.y = element_blank()) +
  facet_grid(proteome ~ normalisation)
```

Based on the total plots, the raw distributions generally look fine, but clearly there are some replicates that probably have more loading than others. This is what we need to correct with normalisation.

As expected, the global normalisation from ProteomeDiscoverer has performed well for the total proteome—especially as the aim of the total proteome is compare based on equal loading. However, for the polyUb

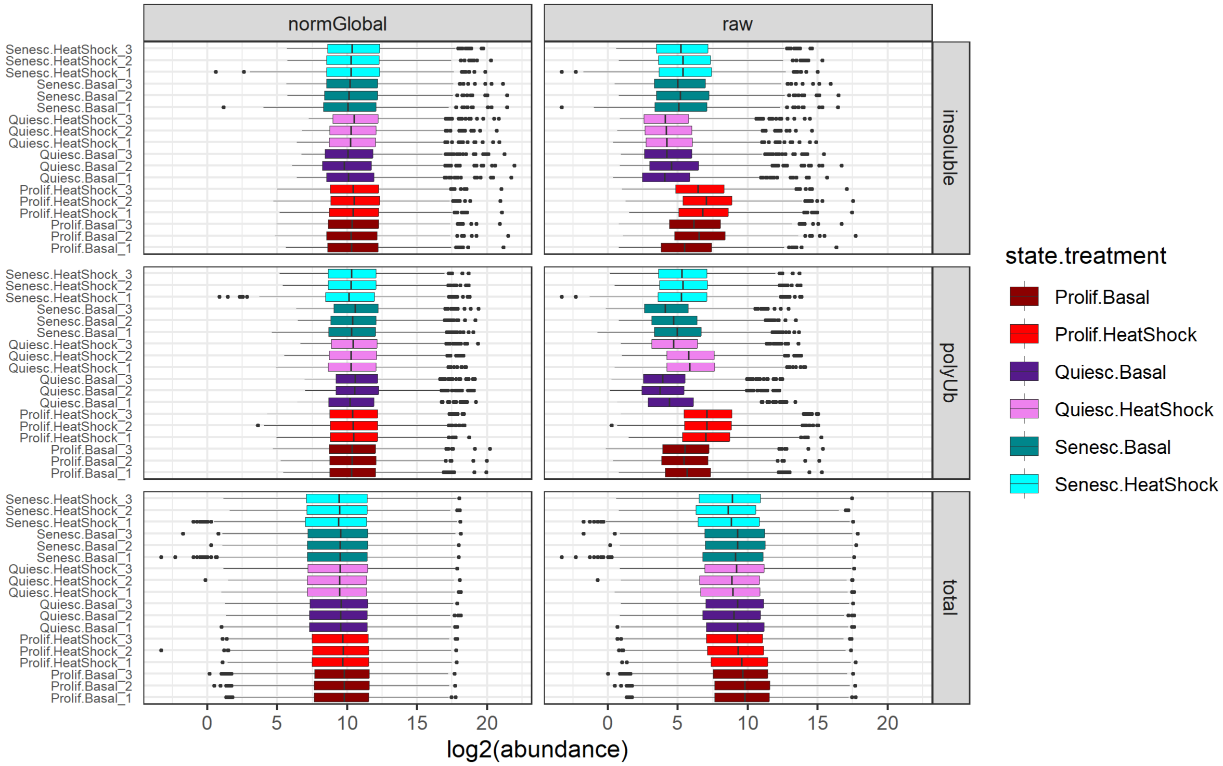

Figure 5: Raw and normalised log2-tranformed abundance distributions of all proteins identified in each sample from the IMR-90 experiment.

proteome (and also, most likely, for the insoluble proteome), it is too harsh; it only works with the assumption that most of the proteins identified are unchanged across our samples. This is clearly not a biologically-sound assumption here, as heat-shock is expected to have a much high level of polyUb-enrichment (which is borne out by the raw data).

We need to formulate a more gentle normalisation for these two proteomes. One possible strategy is to reverse the assumption of global normalisation: rather than assuming that the majority of the proteome is unchanged between samples, we can assume that there is at least a minority of the proteome that is unchanged—even between basal and heat-shocked conditions. Following through with this assumption, we can define the 10 % of the proteins in each proteome that have the lowest coefficients of variation (CVs) across the 18 samples.

We can plot how heat-shock changes the individual protein abundances.

```
# pivot the raw and normGlobal parameters into different columns
dataLong %>%
  pivot_wider(names_from = normalisation, values_from = abundance) -> dataLong

# plot just the raw values
dataLong %>%
  dplyr::mutate(log2raw = log2(raw)) %>%
  group_by(Gene, treatment, state, proteome) %>%
  summarise(log2raw = mean(log2raw)) %>%
  ungroup() %>%
  pivot_wider(names_from = treatment, values_from = log2raw) %>%

  ggplot(aes(x = `Basal`, y = `HeatShock`)) +
  geom_point(size = 0.5, alpha = 0.2) +
  geom_abline(slope = 1, intercept = 0,
              colour = "red", linewidth = 1, alpha = 0.5) +
  facet_grid(cols = vars(state), rows = vars(proteome)) +
  xlab("Log2Abundance (Basal)") +
  ylab("Log2Abundance (Heat-Shock)")
```

Before we try the low-CV-based normalisation, we will separate the three proteomes. In this step, we are also dealing with missing values (zeros) temporarily by replacing them with the lowest non-zero value in each proteome.

```
dataLong %>%
  dplyr::filter(proteome == "total") %>%
  group_by(Gene) %>%
  dplyr::filter(!any(is.na(raw))) %>%
  group_by(state.treatment_rep) %>%
  mutate(raw = replace(raw,raw == 0, min(raw[raw > 0]))) %>%
  ungroup() %>%
  mutate(log2raw = log2(raw)) -> totalome

dataLong %>%
  dplyr::filter(proteome == "polyUb") %>%
  group_by(Gene) %>%
  dplyr::filter(!any(is.na(raw))) %>%
  group_by(state.treatment_rep) %>%
  mutate(raw = replace(raw,raw == 0, min(raw[raw > 0]))) %>%
  ungroup() %>%
  mutate(log2raw = log2(raw)) -> ubome
```

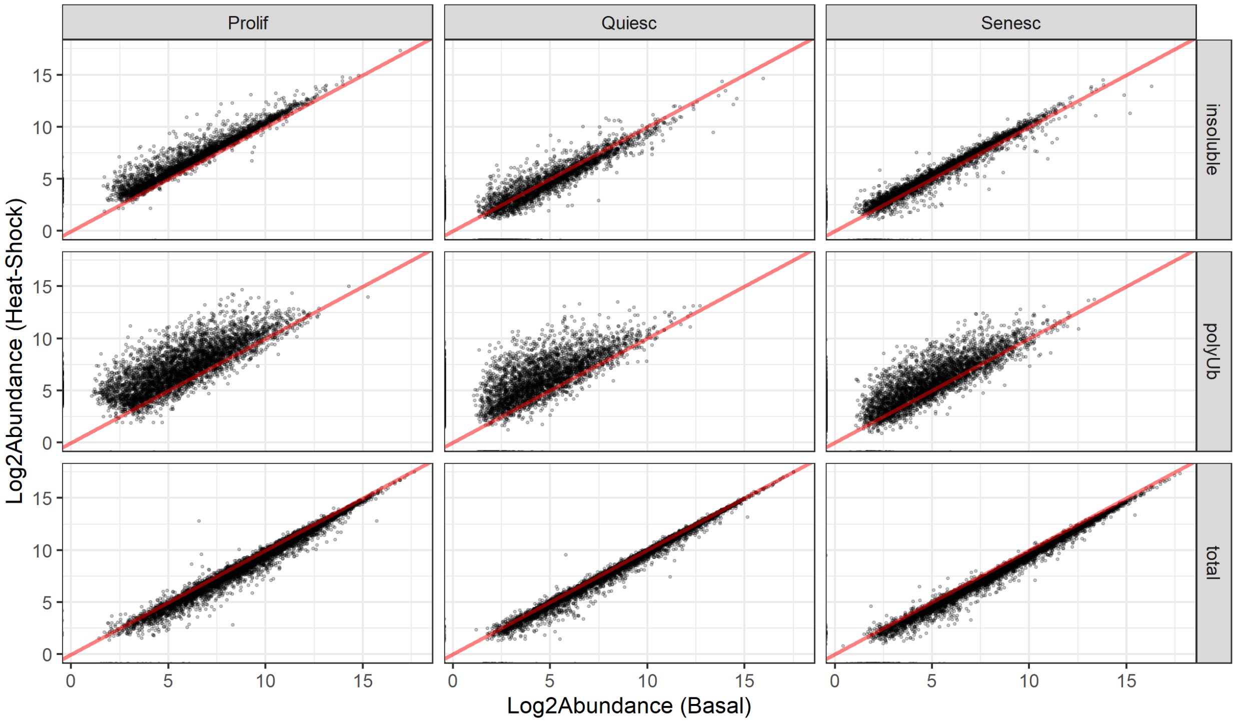

Figure 6: Scatter-plots of mean raw protein abundances in the IMR-90 dataset. Each point represents the mean raw abundance of an individual protein in the basal vs. heat-shocked samples.

```
dataLong %>%
  dplyr::filter(proteome == "insoluble") %>%
  group_by(Gene) %>%
  dplyr::filter(!any(is.na(raw))) %>%
  group_by(state.treatment_rep) %>%
  mutate(raw = replace(raw,raw == 0, min(raw[raw > 0]))) %>%
  ungroup() %>%
  mutate(log2raw = log2(raw)) -> insolome
```

We also need to summarise the mean abundances per condition.

```
totalome %>%
  group_by(Gene, treatment, state) %>%
  summarise(
    log2raw = mean(log2raw)
  ) %>%
  ungroup() %>%
  pivot_wider(
    names_from = treatment,
    values_from = log2raw
  ) -> totalome_per_condition

ubome %>%
  group_by(Gene, treatment, state) %>%
  summarise(
```

```
    log2raw = mean(log2raw)
  ) %>%
  ungroup() %>%
  pivot_wider(
    names_from = treatment,
    values_from = log2raw
  ) -> ubome_per_condition

insolome %>%
  group_by(Gene, treatment, state) %>%
  summarise(
    log2raw = mean(log2raw)
  ) %>%
  ungroup() %>%
  pivot_wider(
    names_from = treatment,
    values_from = log2raw
  ) -> insolome_per_condition
```

##### 2.2.2 Calculate low-CV genes

We will write a function to calculate the CVs, and use this function to generate a list of the genes with the 10% lowest coefficients of variance, for each of the three proteomes separately.

```
cv <- function(x, na.rm = FALSE)  sd(x, na.rm = na.rm)/mean(x, na.rm = na.rm)

dataLong %>%
  group_by(proteome, Gene) %>%
  dplyr::filter(!any(is.na(raw))) %>%
  ungroup() %>%
  group_by(proteome, state.treatment_rep) %>%
  mutate(raw = replace(raw,raw == 0, min(raw[raw > 0]))) %>%
  ungroup() %>%
  group_by(proteome, Gene) %>%
  summarise(cv = cv(raw)) %>%
  arrange(cv) %>%
  filter(cv < quantile(cv, 0.1)) %>% # change this number to alter the % CVs to group
  dplyr::select(-cv) -> lowCVgenes

lowCVgenes %>%
  group_by(proteome) %>%
  summarise(nProteins = n())
```

```
## # A tibble: 3 x 2
##   proteome  nProteins
##   <chr>         <int>
## 1 insoluble       342
## 2 polyUb          455
## 3 total           847
```

Plotting these on the raw scatterplots allows us to visualise what the proteins with the lowest 10% CVs are, just to make sure they seem sensible.

```r
# total
lowCVgenes %>%
  dplyr::filter(proteome == "total") %>%
  pull(Gene) -> lowCVgenes_total

totalome_per_condition %>%
  mutate(stable = Gene %in% lowCVgenes_total) %>%
  arrange(stable) %>%
  ggplot(aes(x = `Basal`, y = `HeatShock`, colour = stable, alpha = stable)) +
  geom_point(size = 0.5, show.legend = FALSE) +
  geom_abline(slope = 1, intercept = 0, colour = "black", linewidth = 1, alpha = 0.5) +
  facet_grid(rows = vars(state)) +
  scale_colour_manual(values = c("darkgrey", "red")) +
  scale_alpha_manual(values = c(0.6, 0.6)) +
  ggtitle("Total") -> scatter_totalome

# polyUb
lowCVgenes %>%
  dplyr::filter(proteome == "polyUb") %>%
  pull(Gene) -> lowCVgenes_polyUb

ubome_per_condition %>%
  mutate(stable = Gene %in% lowCVgenes_polyUb) %>%
  arrange(stable) %>%
  ggplot(aes(x = `Basal`, y = `HeatShock`, colour = stable, alpha = stable)) +
  geom_point(size = 0.5, show.legend = FALSE) +
  geom_abline(slope = 1, intercept = 0, colour = "black", linewidth = 1, alpha = 0.5) +
  facet_grid(rows = vars(state)) +
  scale_colour_manual(values = c("darkgrey", "red")) +
  scale_alpha_manual(values = c(0.6, 0.6)) +
  ggtitle("polyUb") -> scatter_ubome

# insoluble
lowCVgenes %>%
  dplyr::filter(proteome == "insoluble") %>%
  pull(Gene) -> lowCVgenes_insoluble

insolome_per_condition %>%
  mutate(stable = Gene %in% lowCVgenes_insoluble) %>%
  arrange(stable) %>%
  ggplot(aes(x = `Basal`, y = `HeatShock`, colour = stable, alpha = stable)) +
  geom_point(size = 0.5, show.legend = FALSE) +
  geom_abline(slope = 1, intercept = 0, colour = "black", linewidth = 1, alpha = 0.5) +
  facet_grid(rows = vars(state)) +
  scale_colour_manual(values = c("darkgrey", "red")) +
  scale_alpha_manual(values = c(0.6, 0.6)) +
  ggtitle("Insoluble") -> scatter_insolome

scatter_totalome + scatter_ubome + scatter_insolome + plot_layout(ncol = 3)
```

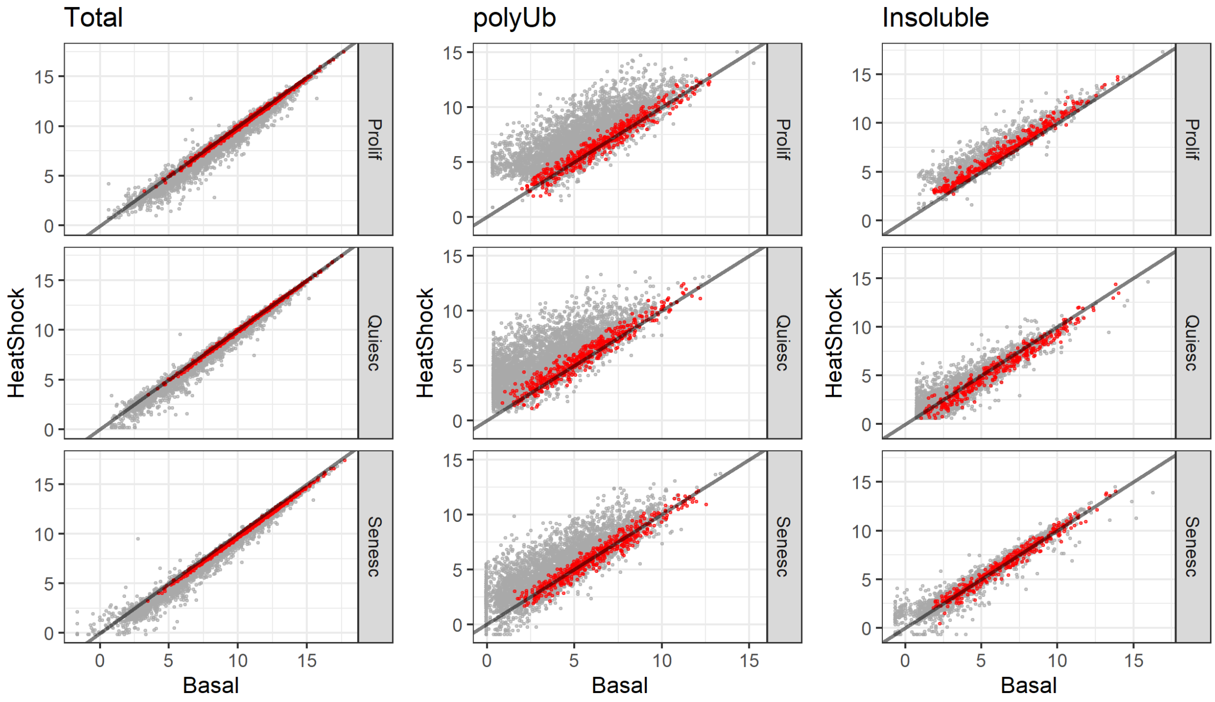

Figure 7: Lowest 10% CV proteins mapped onto scatter-plots of mean raw protein abundances for the IMR-90 dataset. Each point represents the mean raw abundance of an individual protein in the basal vs. heat-shocked samples. The proteins with the 10% lowest CVs for each proteome are plotted in red.

##### 2.2.3 Calculate size-factors based on low-CV genes

Based on the low-CV gene list above, we can calculate the size-factors needed for normalising each proteome.

```
totalome %>%
  group_by(Gene) %>%
  summarise(mean_raw = mean(log2raw)) %>%
  right_join(totalome) %>%
  mutate(diff = log2raw - mean_raw) %>%
  filter(Gene %in% lowCVgenes_total) %>%
  group_by(state, state.treatment, state.treatment_rep) %>%
  summarise(size_factor = median(diff)) %>%
  dplyr::mutate(proteome = "total") -> cv_sizeFactors_total

ubome %>%
  group_by(Gene) %>%
  summarise(mean_raw = mean(log2raw)) %>%
  right_join(ubome) %>%
  mutate(diff = log2raw - mean_raw) %>%
  filter(Gene %in% lowCVgenes_polyUb) %>%
  group_by(state, state.treatment, state.treatment_rep) %>%
  summarise(size_factor = median(diff)) %>%
  dplyr::mutate(proteome = "polyUb")-> cv_sizeFactors_polyUb

insolome %>%
```

```
  group_by(Gene) %>%
  summarise(mean_raw = mean(log2raw)) %>%
  right_join(insolome) %>%
  mutate(diff = log2raw - mean_raw) %>%
  filter(Gene %in% lowCVgenes_insoluble) %>%
  group_by(state, state.treatment, state.treatment_rep) %>%
  summarise(size_factor = median(diff)) %>%
  dplyr::mutate(proteome = "insoluble") -> cv_sizeFactors_insoluble

# plot the size factors
bind_rows(cv_sizeFactors_total,
          cv_sizeFactors_polyUb,
          cv_sizeFactors_insoluble) -> cv_sizeFactors_ALL

cv_sizeFactors_ALL %>%
  ggplot(aes(x = state.treatment_rep, y = size_factor,
             fill = state.treatment, colour = state.treatment)) +
  geom_col(width = 0.1, show.legend = FALSE) +
  geom_hline(yintercept = 0) +
  geom_point(show.legend = FALSE) +
  scale_fill_manual(values = six_sample_colours) +
  scale_colour_manual(values = six_sample_colours) +
  coord_flip() +
  theme(axis.title.y = element_blank(),
        axis.text.y = element_text(size = 6)) +
  facet_wrap(~factor(proteome, levels = c("total", "polyUb", "insoluble"))) +
  ylab("Size factor for normalisation")
```

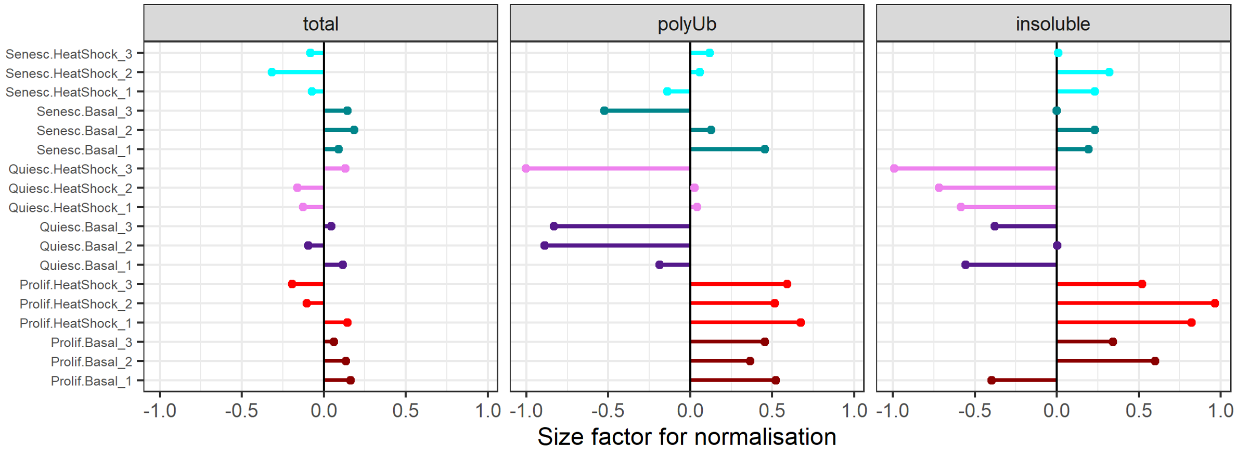

Figure 8: Size factors required for normalisation of each sample in the IMR-90 dataset, based on the mean 10% low-CV protein values.

##### 2.2.4 Normalising data based on low-CV-calculated size-factors

Finally, we can use these calculated size-factors to normalise the data.

```r
# normalise based on size factors
dataLong %>%
  mutate(across(is.numeric, na_if, 0)) %>%
  dplyr::mutate(log2raw = log2(raw)) -> dataLong

dataLong %>%
  mutate(across(is.numeric, na_if, 0)) %>%
  dplyr::mutate(log2normGlobal = log2(normGlobal)) -> dataLong

dataLong %>%
  left_join(cv_sizeFactors_ALL) %>%
  mutate(log2normCV = log2raw-size_factor) %>%
  mutate(normCV = 2^log2normCV) %>%
  dplyr::select(-size_factor) %>%
  ungroup() -> dataNormalised
```

Plot how this normalisation has affected the proteome distributions.

```r
dataNormalised %>%
  dplyr::select(
    proteome, Gene, raw, normGlobal, normCV, state.treatment, state.treatment_rep) %>%
  pivot_longer(cols = c(raw, normGlobal, normCV),
               names_to = "abundance_type", values_to = "abundance") %>%
  dplyr::mutate(abundance_type = factor(
    abundance_type, levels = c("raw", "normGlobal", "normCV"))) %>%
  dplyr::mutate(proteome = factor(
    proteome, levels = c("total", "polyUb", "insoluble"))) %>%

  ggplot(aes(x = state.treatment_rep, y = log2(abundance), fill = state.treatment)) +
  geom_boxplot(outlier.size = 0.4, linewidth = 0.2) +
  coord_flip() +
  scale_fill_manual(values = six_sample_colours) +
  theme(axis.title.y = element_blank(),
        axis.text.y = element_text(size = 6),
        axis.ticks.y = element_blank()) +
  facet_grid(proteome~abundance_type, scales = "free")
```

```r
dataNormalised %>%
  group_by(Gene, treatment, state, proteome) %>%
  summarise(
    log2normCV = mean(log2normCV)
  ) %>%
  ungroup() %>%
  pivot_wider(
    names_from = treatment,
    values_from = log2normCV
  ) %>%

  ggplot(aes(x = `Basal`, y = `HeatShock`)) +
  geom_point(size = 0.5, alpha = 0.2) +
  geom_abline(slope = 1, intercept = 0, colour = "red", linewidth = 1, alpha = 0.5) +
  ggtitle("Mean CV-Normalised Abundances") +
```

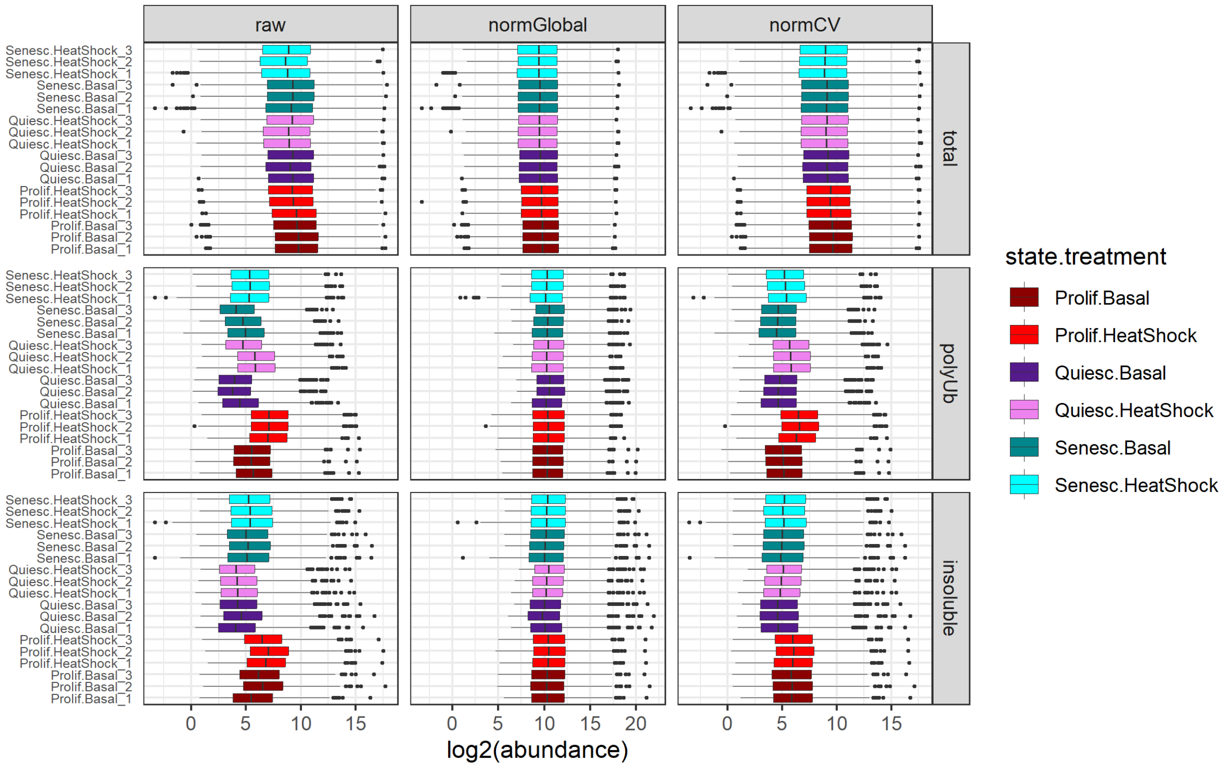

Figure 9: Raw and normalised protein abundance distributions in each sample of the IMR-90 dataset.

```
  facet_grid(cols = vars(state),
             rows = vars(proteome)) +
  xlab("Log2Abundance (Basal)") +
  ylab("Log2Abundance (Heat-Shock)")
```

Comparing these CV-normalised distributions with the global-normalised distributions earlier, it seems clear that the low-CV normalisation approach preserves most of the heat-shock-induced changes that are lost with global normalisation.

For the remainder of the analysis, we will proceed with the CV-normalised abundances for the polyUb and insoluble proteomes, and keep the global-normalised total proteomes.

We can save all three of the these values (raw, global-normalised, and low-CV-normalised) as **dataNormalised__All.csv** file, and just the normalised values that will be used for further analysis (global-normalised for total; CV-normalised for polyUb and insoluble) as **dataNormalised__Selected.csv**.

```
dataNormalised %>%
  dplyr::filter(!is.na(raw)) %>%
  unite(proteome_state.treatment_rep, c("proteome", "state.treatment_rep"),
        sep = "_") -> dataNormalised

dataNormalised %>%
  dplyr::select(c(Gene, proteome_state.treatment_rep, raw)) %>%
  dplyr::mutate(
    proteome_state.treatment_rep = paste0("raw_", proteome_state.treatment_rep)) %>%
  pivot_wider(names_from = proteome_state.treatment_rep, values_from = raw) %>%
```

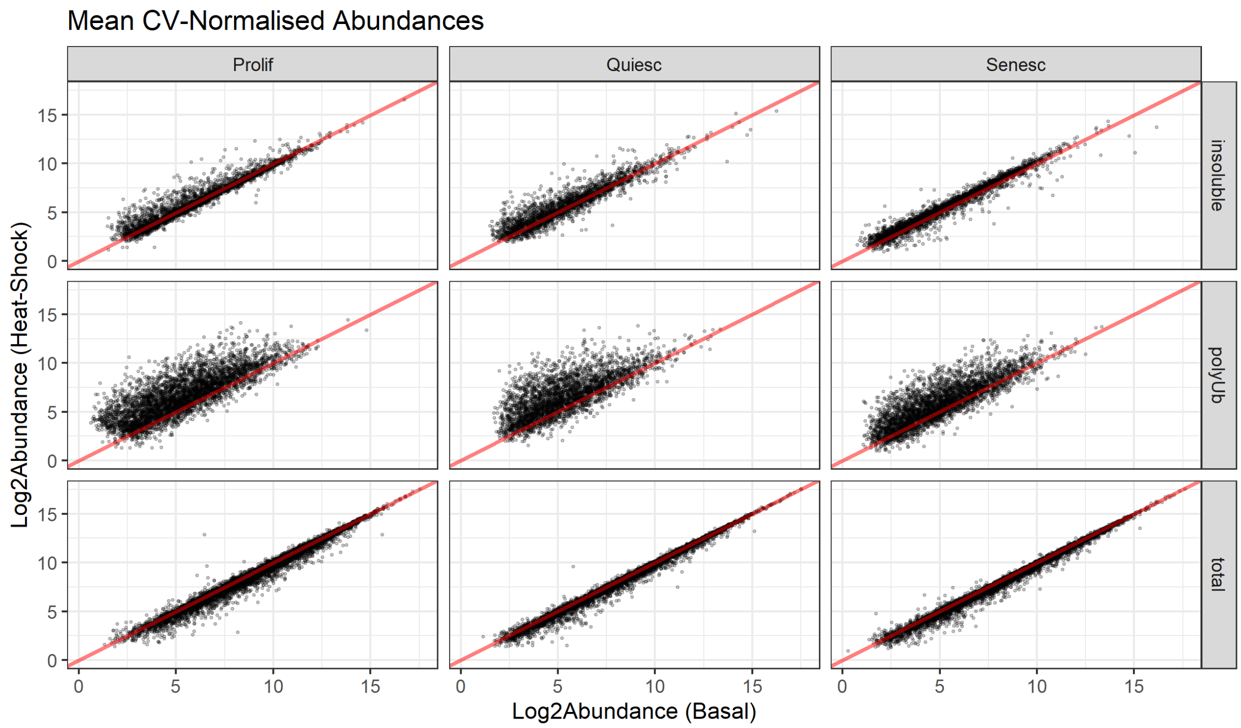

Figure 10: Scatter-plots of mean CV-normalised protein abundances for each state in the IMR-90 experiment. Each point represents the mean CV-normalised abundance of an individual protein in the basal vs. heat-shocked samples.

```
  arrange(Gene) -> dataNormalised_raw

dataNormalised %>%
  dplyr::select(c(Gene, proteome_state.treatment_rep, normGlobal)) %>%
  dplyr::mutate(proteome_state.treatment_rep = paste0(
    "normGlobal_", proteome_state.treatment_rep)) %>%
  pivot_wider(names_from = proteome_state.treatment_rep, values_from = normGlobal) %>%
  arrange(Gene) -> dataNormalised_normGlobal

dataNormalised %>%
  dplyr::select(c(Gene, proteome_state.treatment_rep, normCV)) %>%
  dplyr::mutate(proteome_state.treatment_rep = paste0(
    "normCV_", proteome_state.treatment_rep)) %>%
  pivot_wider(names_from = proteome_state.treatment_rep, values_from = normCV) %>%
  arrange(Gene) -> dataNormalised_normCV

reduce(list(dataNormalised_raw, dataNormalised_normGlobal, dataNormalised_normCV),
       full_join) -> dataNormalised_All

# save all abundance values as "dataNormalised_All.csv"
dataNormalised_All %>%
  write_csv("IMR90/dataNormalised_All.csv",
            na = "NA", append = FALSE, col_names = TRUE, escape = "double")

# save only selected normalised abundances as "dataNormalised_Selected.csv"
```

```r
dataNormalised_All %>%
  dplyr::select(-starts_with("raw")) %>%
  dplyr::select(-starts_with(c("normCV_total"))) %>%
  dplyr::select(-starts_with(c("normGlobal_polyUb", "normGlobal_insoluble"))) %>%
  rename_with(~ gsub("normCV_", "", .x, fixed = TRUE)) %>%
  rename_with(~ gsub("normGlobal_", "", .x, fixed = TRUE)) -> dataNormalised_selected

dataNormalised_selected %>%
  write_csv("IMR90/dataNormalised_Selected.csv",
            na = "NA", append = FALSE, col_names = TRUE, escape = "double")
```

#### 2.3 Missing Values

We can now decide how to process missing values.

##### 2.3.1 Overlapping proteins

Let's first compare how many overlapping proteins were identified across our proteomes (Fig. 11). An upset plot is what is required, using the UpSetR package (Conway, Lex, and Gehlenborg 2017).

```r
library(UpSetR)

# make a data frame with 1s and 0s for protein presence in a proteome.
dataNormalised_selected %>%
  dplyr::select(Gene, starts_with(c("total","polyUb","insoluble"))) %>%
  pivot_longer(cols = starts_with(c("total","polyUb","insoluble")),
               names_to = "Sample",
               values_to = "Abundance") %>%
  separate(Sample, into = c("Proteome", "Sample.Treatment", "Replicate"), sep = "_") %>%
  unite(Sample.Treatment_Rep, c("Sample.Treatment", "Replicate"), sep = "_") %>%

  dplyr::select(-Sample.Treatment_Rep) %>%
  group_by(Gene, Proteome) %>%
  summarise(
    present = case_when(
      all(is.na(Abundance)) ~ as.integer(0),
      .default = as.integer(1))
  ) %>%
  ungroup() %>%
  pivot_wider(names_from = Proteome, values_from = present) %>%
  as.data.frame() -> upset_input_proteomes

upset(upset_input_proteomes,
      sets = c("insoluble", "polyUb", "total"), keep.order = TRUE,
      order.by = "degree", empty.intersections = "on",
      point.size = 3, line.size = 1,
      sets.x.label = "Proteins Quantified",
      mainbar.y.label = "Unique Proteins")
```

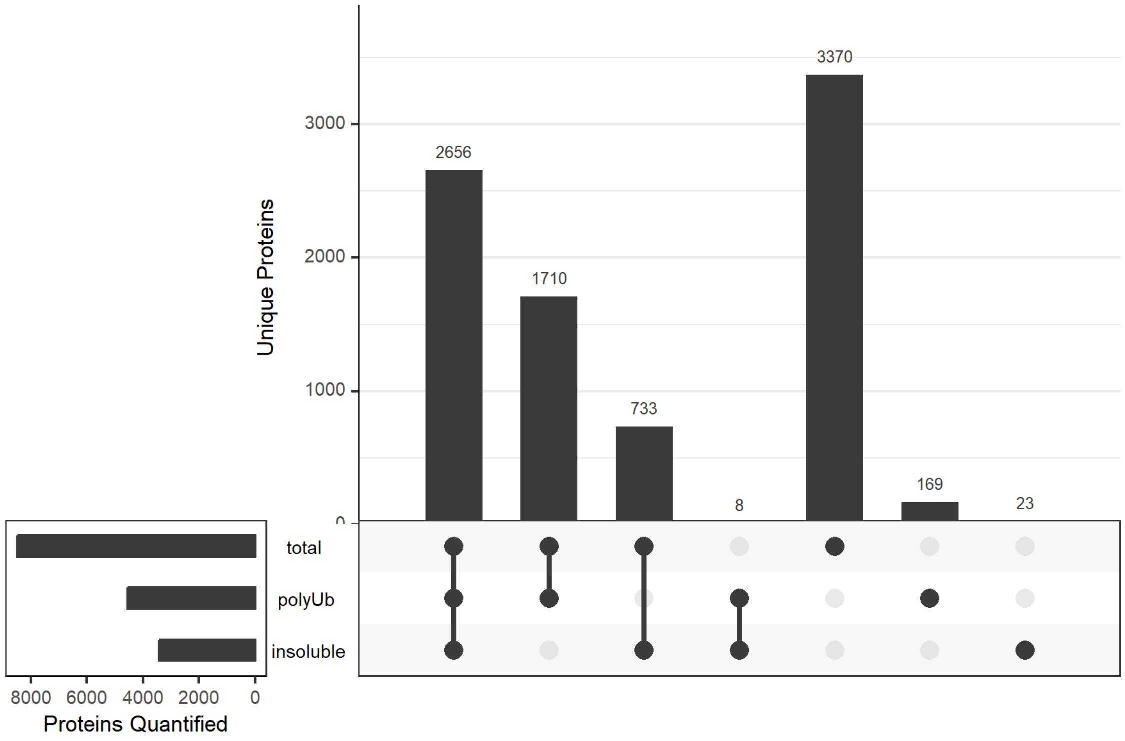

Figure 11: UpSet plot of overlap between proteins quantified in any sample between the three proteomes for the IMR-90 experiment.

##### 2.3.2 Channel occupancy

We also need to find the number of proteins with incomplete channel occupancy (i.e., NAs in one or more of the 18 samples).

The initial PD analysis already filtered out proteins that were quantified (i.e., non-NA values) in fewer than 6 of the 18 channels. We might want to be more stringent with our filtering.

```
dataNormalised_selected %>%
  pivot_longer(cols = -Gene, names_to = "sample", values_to = "abundance") %>%
  separate(sample, into = c("proteome", "state.treatment", "replicate"), sep="_") %>%
  separate(state.treatment, into = c("state", "treatment"), sep = "\\.", remove=F) %>%
  group_by(proteome, Gene) %>%
  ungroup() %>%
  unite("label", c(`state.treatment`, `replicate`), sep = "_", remove = F) -> dataLong

dataLong %>%
  mutate(abundance = na_if(abundance, 0)) %>%
  drop_na() %>%
  group_by(proteome, Gene) %>%
  dplyr::count() %>%
  group_by(proteome, n) %>%
  dplyr::count() %>%

  ggplot(aes(x = n, y = nn)) +
  geom_col(show.legend = F, width = 0.5) +
```

```
  scale_x_continuous(breaks = seq(6, 18, by = 1)) +
  ylab("count(Proteins)") +
  xlab("Identified in number of samples") +
  theme(axis.text.x = element_text(angle = 90, vjust = 0.5, hjust = 1)) +
  facet_wrap(~factor(proteome, levels = c("total", "polyUb", "insoluble")),
             scales = "free_y")
```

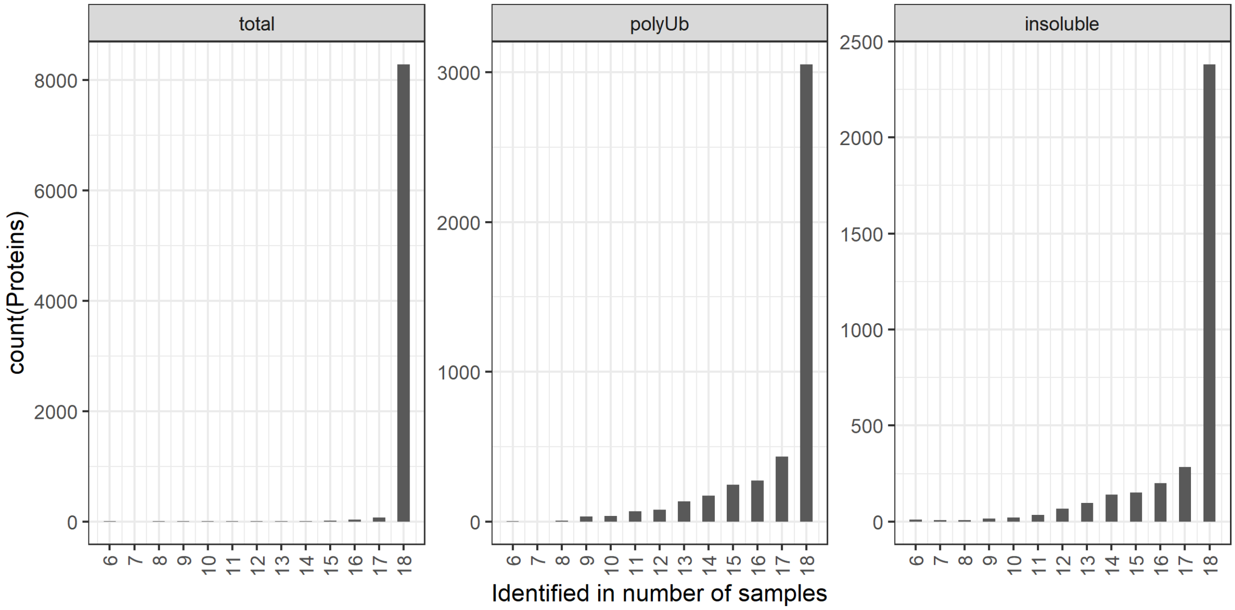

Figure 12: Channel occupancy for each proteome in the IMR-90 experiment.

It is also useful to visualise the number of proteins that were quantified in each sample, and the number of missing values per state.treatment.

```
dataLong %>%
  mutate(abundance = na_if(abundance, 0)) %>%
  drop_na() %>%
  group_by(proteome, label) %>%
  dplyr::count() %>%
  left_join(experimental_design %>%
              dplyr::select(label, condition) %>%
              distinct(label, .keep_all = TRUE)) %>%

  ggplot(aes(x = label, y = n, fill = condition)) +
  geom_col(show.legend = F, width = 0.5) +
  scale_fill_manual(values = six_sample_colours) +
  ylab("count(Proteins)") +
  theme(axis.text.x = element_text(angle = 90, vjust = 0.5, hjust=1, size = 6),
        axis.title.x = element_blank()) +
  facet_wrap(~factor(proteome, levels=c("total", "polyUb", "insoluble")),
             scales = "free_y")
```

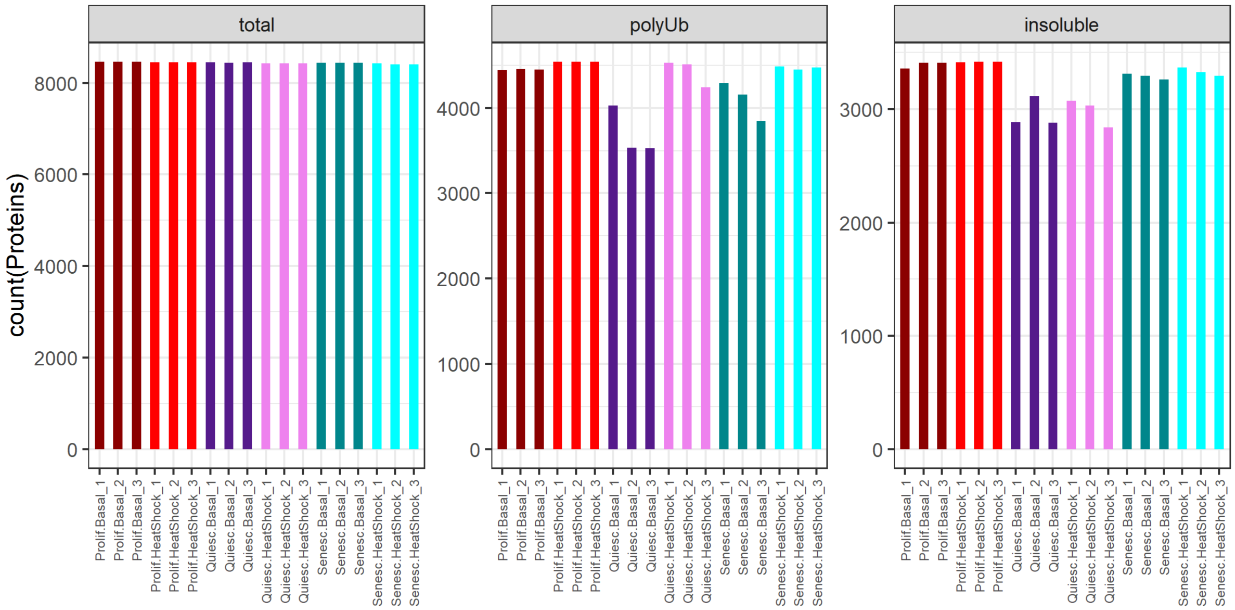

Figure 13: Number of proteins quantified in each sample of the IMR-90 dataset.

```
dataLong %>%
  group_by(Gene, proteome) %>%
  dplyr::filter(!all(is.na(abundance))) %>%

  dplyr::group_by(Gene, proteome, state.treatment) %>%
  dplyr::summarise(num_NA = sum(is.na(abundance))) %>%
  ungroup() %>%
  dplyr::filter(num_NA > 0) %>%
  dplyr::mutate(num_NA = as.factor(num_NA)) %>%

  ggplot(aes(x = state.treatment, fill = num_NA)) +
  geom_bar() +
  ylab("Number of proteins with missing values") +
  theme(axis.text.x = element_text(angle = 45, hjust = 1),
        axis.title.x = element_blank()) +
  facet_grid(~factor(proteome, levels = c("total", "polyUb", "insoluble")))
```

These plots make a few points. The total proteome has very few missing values. The polyUb proteome has missing values mostly in the basal condition and not in the heat-shock condition, as expected (with the slight exception of Quiesc.HeatShock_3). The insoluble proteome has missing values mostly in the quiescent state (both basal and heat-shock).

Therefore, for both polyUb and insoluble proteomes, these are Missing Not At Random (MNAR), i.e., proteins that were not quantified in specific conditions (e.g., in quiescent or basal samples only). MNAR can be indicative of proteins below the detection limit in specific samples—which is what we hypothesise is going on here.

We need to decide whether (and how) to impute these missing values.

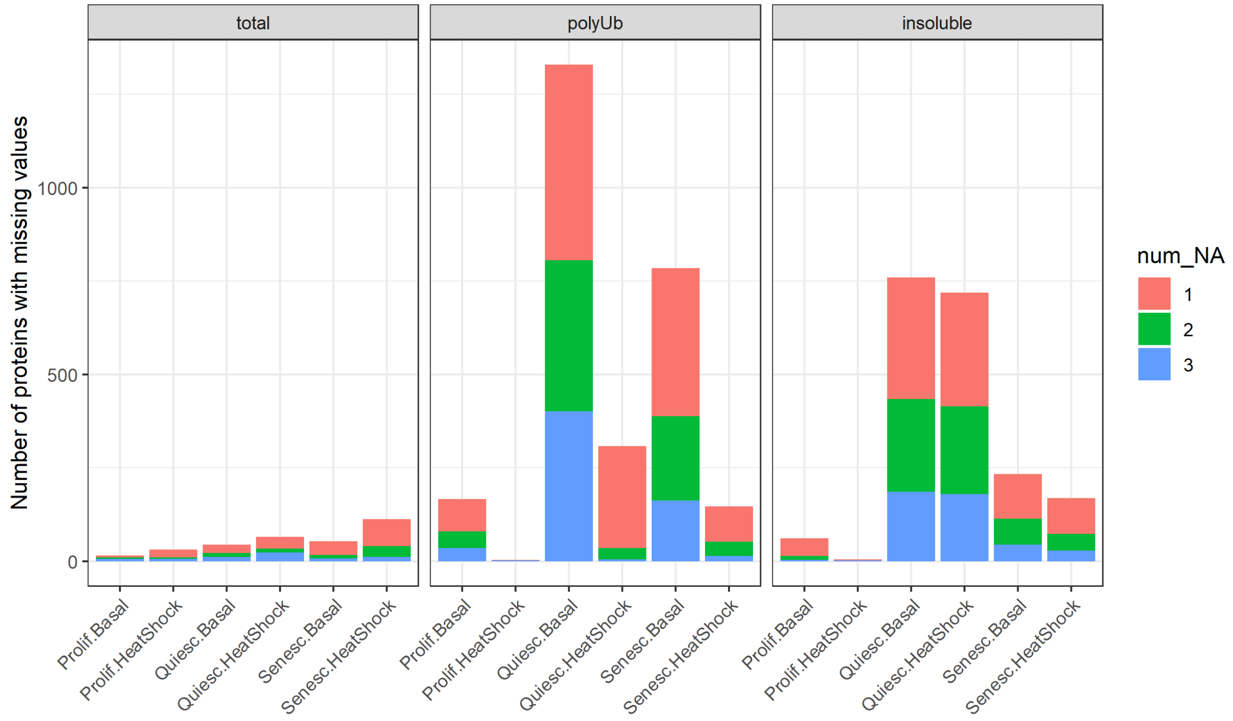

Figure 14: Number of missing values in each state.treatment per proteome of the IMR-90 dataset.

##### 2.3.3 Imputing missing values

We can use the built-in imputation functions in the R package DEP (Zhang et al. 2018).

```
library(DEP)
```

From the DEP vignette: "Many Bioconductor packages use SummarizedExperiment objects as input and/or output. This class of objects contains and coordinates the actual (assay) data, information on the samples as well as feature annotation."

"The experimental design must contain 'label', 'condition' and 'replicate' columns. The 'label' column contains the identifiers of the different samples and they should correspond to the column names containing the assay data. The 'condition' and 'replicate' columns contain the annotation of these samples as defined by the user."

First, we separate out the proteomes.

For this, we need the Accessions, Gene Symbols, and abundance values. We can re-enter this information from **dataFiltered.csv** before separating the proteomes.

```
read_csv("IMR90/dataFiltered.csv") %>%
  dplyr::select(Gene, Accession) %>%
  right_join(dataNormalised_selected) -> data

data %>%
  dplyr::select(Gene, Accession, starts_with(c("total", "polyUb", "insoluble"))) %>%
  dplyr::rename(name = Gene) %>%
  dplyr::rename(ID = Accession) -> data
```

```
data %>%
  dplyr::select(name, ID, starts_with("total")) %>%
  dplyr::filter_at(vars(starts_with("total")), any_vars(!is.na(.))) %>%
  select_all(~gsub("total_", "", .)) -> data_total

data %>%
  dplyr::select(name, ID, starts_with("polyUb")) %>%
  dplyr::filter_at(vars(starts_with("polyUb")), any_vars(!is.na(.))) %>%
  select_all(~gsub("polyUb_", "", .)) -> data_polyUb

data %>%
  dplyr::select(name, ID, starts_with("insoluble")) %>%
  dplyr::filter_at(vars(starts_with("insoluble")), any_vars(!is.na(.))) %>%
  select_all(~gsub("insoluble_", "", .)) -> data_insoluble
```

To generate the SummarizedExperiment object *SE_for_DEP* from *data_total*, we use the experimental design table we made for the study design. We can do this once for all four proteomes, as the annotations are the same.

```
# specify the abundance column names for the SummarizedExperiment
abundance_columns <- grep("Prolif|Quiesc|Senesc", colnames(data_total))

experimental_design %>%
  dplyr::select(label, condition, replicate) %>%
  distinct() -> experimental_design

SEforDEP_total <- make_se(data_total, abundance_columns, experimental_design)
SEforDEP_polyUb <- make_se(data_polyUb, abundance_columns, experimental_design)
SEforDEP_insoluble <- make_se(data_insoluble, abundance_columns, experimental_design)
```

**2.3.3.1 Deciding on imputation model** A different way of exploring the pattern of missing values is through a heatmap with values missing (0) or not (1). Only proteins with at least one missing value are visualised.

```
plot_missval(SEforDEP_total)
```

```
plot_missval(SEforDEP_polyUb)
```

```
plot_missval(SEforDEP_insoluble)
```

We will also check whether missing values are biased towards less intense proteins, as is typically the case with data-dependent-acquisition (DDA)- based proteomics datasets. We can plot intensity distributions and cumulative fraction of proteins with and without missing values.

```
plot_detect(SEforDEP_total)
```

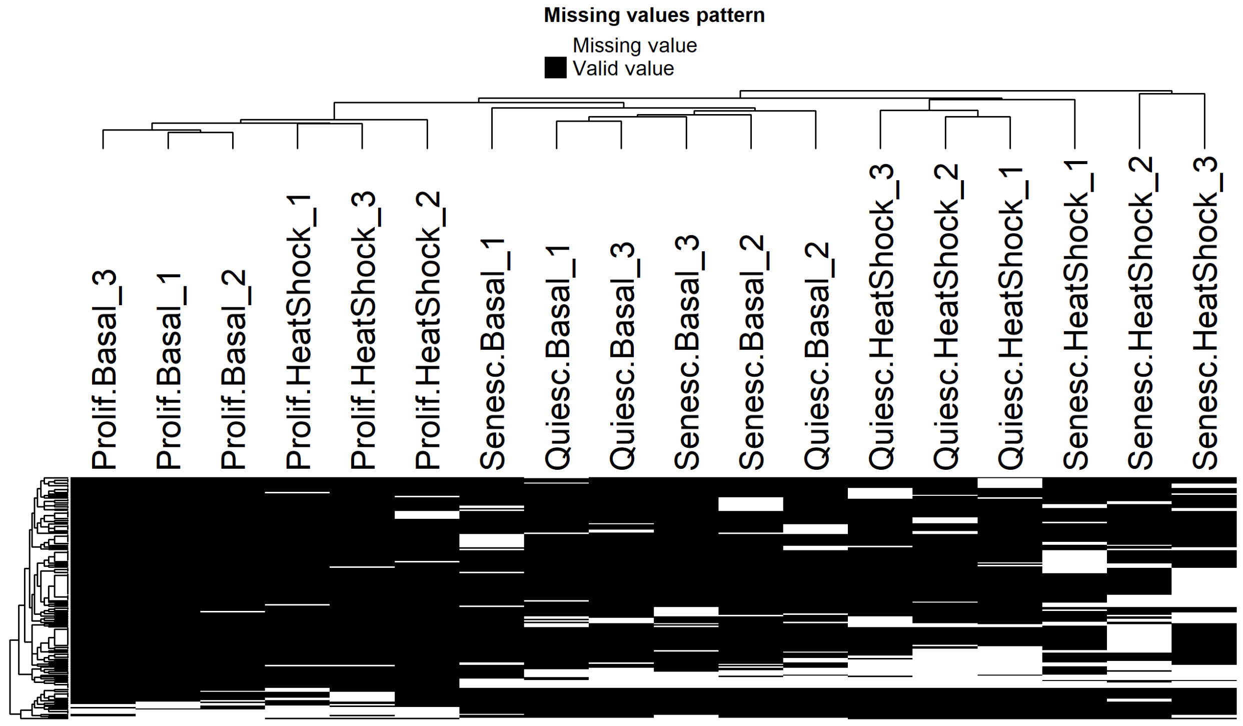

Figure 15: Heatmaps of proteins in the total proteome of the IMR-90 dataset that have at least one missing value. Samples where abundance values are missing (white) or present (black) are shown.

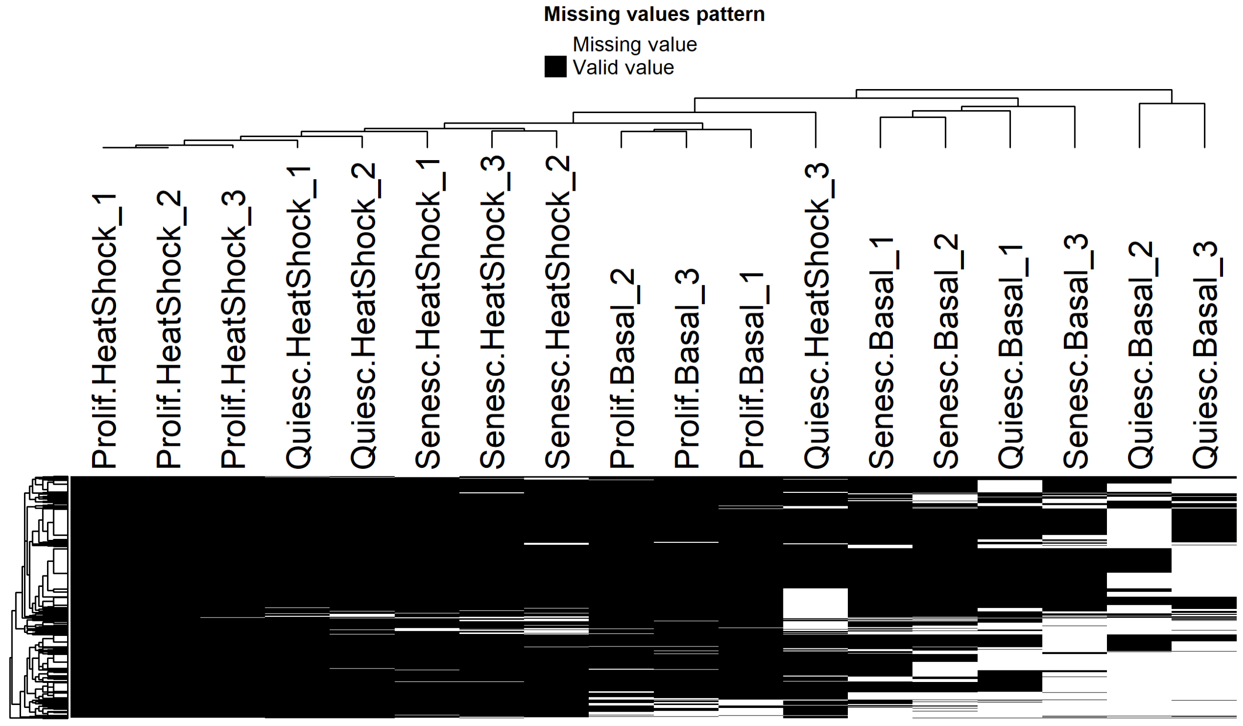

Figure 16: Heatmaps of proteins in the polyUb proteome of the IMR-90 dataset that have at least one missing value. Samples where abundance values are missing (white) or present (black) are shown.

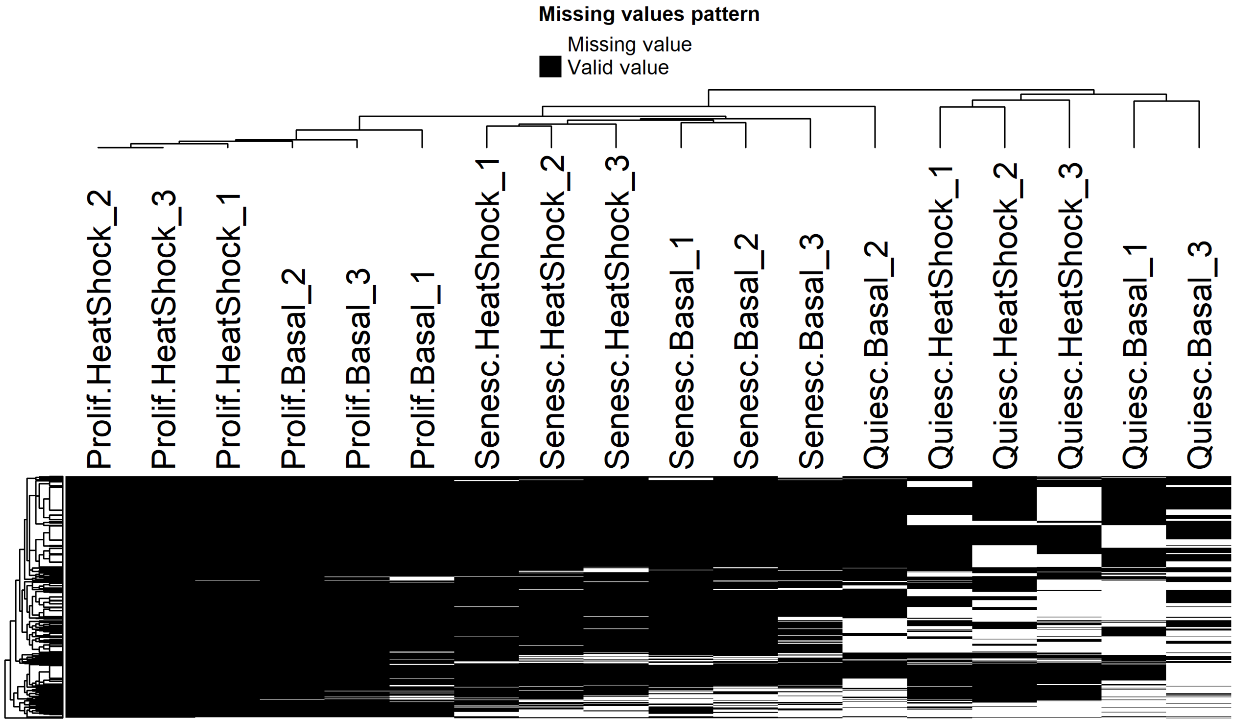

Figure 17: Heatmaps of proteins in the insoluble proteome of the IMR-90 dataset that have at least one missing value. Samples where abundance values are missing (white) or present (black) are shown.

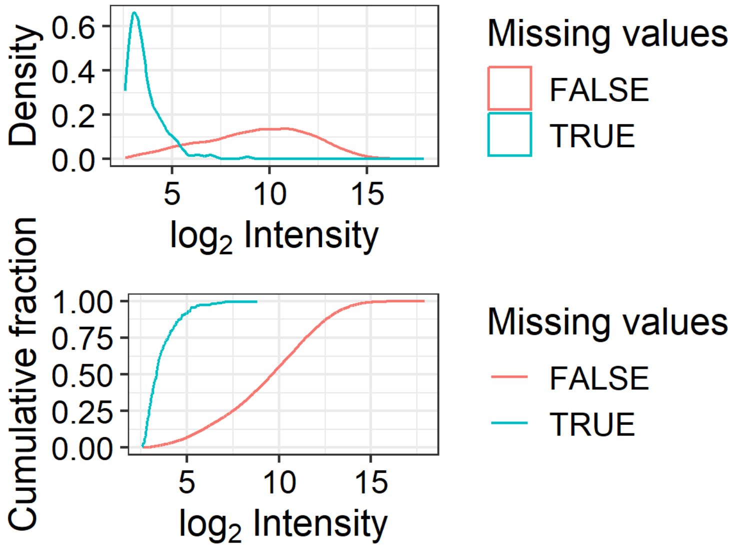

Figure 18: Density (top) and cumulative density (bottom) distributions of log2-transformed intensities, for proteins with (turquoise) or without (red) one or more missing values, in th total proteome of the IMR-90 dataset.

```r
plot_detect(SEforDEP_polyUb)
```

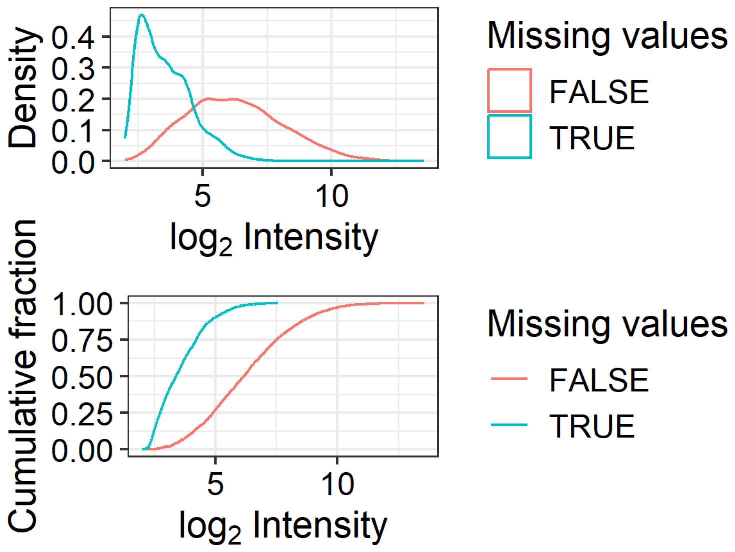

Figure 19: Density (top) and cumulative density (bottom) distributions of log2-transformed intensities, for proteins with (turquoise) or without (red) one or more missing values, in th polyUb proteome of the IMR-90 dataset.

```r
plot_detect(SEforDEP_insoluble)
```

Although it varies a little bit across proteomes, proteins with missing values do on average have low intensities.

This type of data (MNAR, and close-to-detection-limit) should be imputed by a left-censored imputation method available within the DEP package, e.g., quantile regression-based left-censored function ("QRILC"), or random draws from a left-shifted distribution ("MinProb" and "man").

Before we proceed to imputation, we will set some quality filters so that only proteins quantified in at least 50 % of the samples, and in all three replicates of at least one condition, are kept in the data.

```r
SEforDEP_total <- filter_proteins(SEforDEP_total, "fraction", min = 0.5)
SEforDEP_total <- filter_proteins(SEforDEP_total, "condition", thr = 0)
SEforDEP_polyUb <- filter_proteins(SEforDEP_polyUb, "fraction", min = 0.5)
SEforDEP_polyUb <- filter_proteins(SEforDEP_polyUb, "condition", thr = 0)
SEforDEP_insoluble <- filter_proteins(SEforDEP_insoluble, "fraction", min = 0.5)
SEforDEP_insoluble <- filter_proteins(SEforDEP_insoluble, "condition", thr = 0)
```

```r
# MNAR Imputation 1: MinProb
## Impute missing data using random draws from a Gaussian distribution
## centred around a minimal value
MinProb_total <- impute(SEforDEP_total, fun = "MinProb", q = 0.01)
```

###### 2.3.3.2 Assessing imputation models

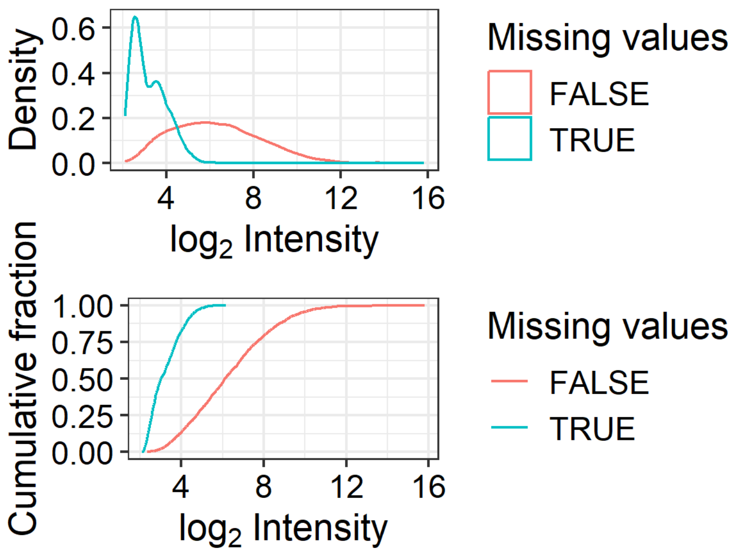

Figure 20: Density (top) and cumulative density (bottom) distributions of log2-transformed intensities, for proteins with (turquoise) or without (red) one or more missing values, in th insoluble proteome of the IMR-90 dataset.

```
## [1] 0.4193667
```

```
MinProb_polyUb <- impute(SEforDEP_polyUb, fun = "MinProb", q = 0.01)
```

```
## [1] 0.956232
```

```
MinProb_insoluble <- impute(SEforDEP_insoluble, fun = "MinProb", q = 0.01)
```

```
## [1] 0.870139
```

```
# MNAR Imputation 2: Manual
## Impute missing data using random draws from a manually-defined
## left-shifted Gaussian distribution
Manual_total <- impute(SEforDEP_total, fun = "man", shift = 1.8, scale = 0.3)
Manual_polyUb <- impute(SEforDEP_polyUb, fun = "man", shift = 1.8, scale = 0.3)
Manual_insoluble <- impute(SEforDEP_insoluble, fun = "man", shift = 1.8, scale = 0.3)

# MNAR Imputation 3: QRILC
## Impute missing data using quantile regression-based
## left-censored function
QRILC_total <- impute(SEforDEP_total, fun = "QRILC")
QRILC_polyUb <- impute(SEforDEP_polyUb, fun = "QRILC")
QRILC_insoluble <- impute(SEforDEP_insoluble, fun = "QRILC")
```

```
plot_imputation(SEforDEP_total, MinProb_total, Manual_total, QRILC_total) +
  ggtitle("Total") -> imputations_total
plot_imputation(SEforDEP_polyUb, MinProb_polyUb, Manual_polyUb, QRILC_polyUb) +
  ggtitle("polyUb") -> imputations_polyUb
```

```r
plot_imputation(
  SEforDEP_insoluble, MinProb_insoluble, Manual_insoluble, QRILC_insoluble) +
  ggtitle("Insoluble") -> imputations_insoluble

imputations_total + imputations_polyUb + imputations_insoluble +
  plot_layout(ncol = 2) + plot_annotation(tag_levels = 'A')
```

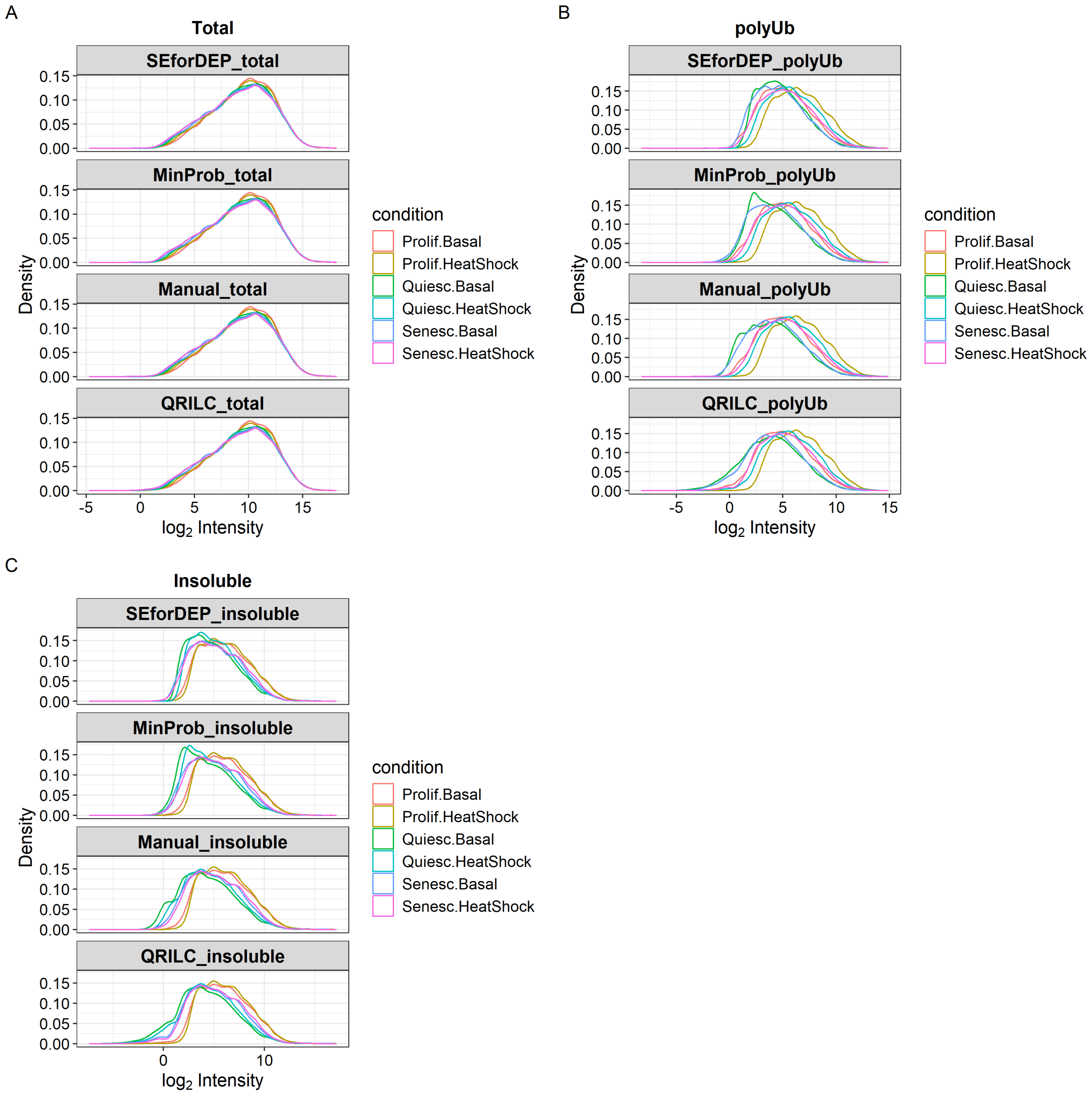

Figure 21: Density distributions of log2-transformed intensities with each imputation model across total (A), polyUb (B), and insoluble (C) proteomes, for the IMR-90 dataset.

What we seek is the imputation method that changes the unimputed data (the "SEforDEP" facet) the least. The MinProb models add shoulders to the polyUb and insoluble proteomes, and therefore should not be considered. The Manual and QRILC models seem acceptable, but with slightly different trade-offs or artefacts.

We will proceed with the QRILC imputation, and save it as **dataImputed.csv**. Note that this is the only step in the workflow where there is a degree of stochasticity introduced, i.e., re-running the code will yield slightly different numbers every time, due to the nature of the imputation. In order to avoid this, we will comment out this code-chunk after running it the first time, so that **dataImputed.csv** is only generated once.

```
# Uncomment the following code chunk only the first time you are running the script
# total_imp <- get_df_wide(QRILC_total) %>%
#   rename(Gene = name, Accession = ID) %>%
#   pivot_longer(cols = -c("Gene", "Accession", "imputed", "num_NAs"),
#                names_to = "State.Treatment_Rep", values_to = "Abundance") %>%
#   mutate(Proteome = "total") %>%
#   dplyr::rename("num_NAs_total" = num_NAs)
#
# polyUb_imp <- get_df_wide(QRILC_polyUb) %>%
#   rename(Gene = name, Accession = ID) %>%
#   pivot_longer(cols = -c("Gene", "Accession", "imputed", "num_NAs"),
#                names_to = "State.Treatment_Rep", values_to = "Abundance") %>%
#   mutate(Proteome = "polyUb") %>%
#   dplyr::rename("num_NAs_polyUb" = num_NAs)
#
# insoluble_imp <- get_df_wide(QRILC_insoluble) %>%
#   rename(Gene = name, Accession = ID) %>%
#   pivot_longer(cols = -c("Gene", "Accession", "imputed", "num_NAs"),
#                names_to = "State.Treatment_Rep", values_to = "Abundance") %>%
#   mutate(Proteome = "insoluble") %>%
#   dplyr::rename("num_NAs_insoluble" = num_NAs)
#
# dataLong <- bind_rows(total_imp, polyUb_imp, insoluble_imp) %>%
#   unite("Proteome_State.Treatment_Rep", c(Proteome, State.Treatment_Rep),
#         sep = "_", remove = FALSE) %>%
#   dplyr::mutate("Abundance" = 2^Abundance)
#
# dataLong %>%
#   dplyr::filter(Proteome == "total") %>%
#   dplyr::rename("imputed_total" = imputed) %>%
#   dplyr::select(Accession, Gene, imputed_total, num_NAs_total,
#                 Proteome_State.Treatment_Rep, Abundance) %>%
#   pivot_wider(names_from = Proteome_State.Treatment_Rep,
#               values_from = Abundance) -> data_total
#
# dataLong %>%
#   dplyr::filter(Proteome == "insoluble") %>%
#   dplyr::rename("imputed_insoluble" = imputed) %>%
#   dplyr::select(Accession, Gene, imputed_insoluble, num_NAs_insoluble,
#                 Proteome_State.Treatment_Rep, Abundance) %>%
#   pivot_wider(names_from = Proteome_State.Treatment_Rep,
#               values_from = Abundance) -> data_insoluble
#
# dataLong %>%
#   dplyr::filter(Proteome == "polyUb") %>%
#   dplyr::rename("imputed_polyUb" = imputed) %>%
#   dplyr::select(Accession, Gene, imputed_polyUb,
```

```
#                  num_NAs_polyUb, Proteome_State.Treatment_Rep, Abundance) %>%
#   pivot_wider(names_from = Proteome_State.Treatment_Rep,
#               values_from = Abundance) -> data_polyUb
#
# data_total %>%
#   left_join(data_insoluble) %>%
#   left_join(data_polyUb) -> dataImputed
#
# dataImputed %>%
#   dplyr::select(order(colnames(dataImputed))) %>%
#   write_csv("IMR90/dataImputed.csv")

dataImputed <- read_csv("IMR90/dataImputed.csv")
```

#### 2.4 Differential Analysis

DEqMS will be used for differential expression/enrichment analysis on each proteome, which is based on limma, but with some adjustments to take into account differences between transcriptomics and proteomics data (e.g., importance of the number of peptides/PSMs in variance estimates) (Zhu et al. 2020).

We need to incorporate the number of PSMs quantified per protein for each proteome back into the *dataImputed* data-frame.

```
read_csv("../../IMR90_PSQ_HS_TotUbiIns/data.csv") %>%
  dplyr::select(Accession, starts_with("# Quantitations")) %>%
  right_join(dataImputed) -> dataImputed
```

Separating *dataImputed* into the individual proteomes will allow us to use DEqMS separately on each proteome.

```
dataImputed %>%
  dplyr::select(Gene, starts_with("total"),
                starts_with("# Quantitations total")) %>%
  rowwise() %>%
  mutate(numQuants=max(c_across(
    `# Quantitations total_Prolif` : `# Quantitations total_Quiesc`))) %>%
  dplyr::select(-starts_with("#")) %>%
  dplyr::filter(numQuants != 0) -> dataTotal

dataImputed %>%
  dplyr::select(Gene, starts_with("polyUb"),
                starts_with("# Quantitations ubiquitin")) %>%
  rowwise() %>%
  mutate(numQuants=max(c_across(
    `# Quantitations ubiquitin_Prolif` : `# Quantitations ubiquitin_Quiesc`))) %>%
  dplyr::select(-starts_with("#")) %>%
  dplyr::filter(numQuants != 0) -> dataPolyUb

dataImputed %>%
  dplyr::select(Gene, starts_with("insoluble"),
                starts_with("# Quantitations insoluble")) %>%
  rowwise() %>%
```

```
  mutate(numQuants=max(c_across(
    `# Quantitations insoluble_Prolif` : `# Quantitations insoluble_Quiesc`))) %>%
  dplyr::select(-starts_with("#")) %>%
  dplyr::filter(numQuants != 0) -> dataInsoluble
```

##### 2.4.1 Differential Analysis: Total Proteome

The DEqMS workflow involves a series of data transformation and classification steps, described in detail in the DEqMS vignette.

```
# define coordinates of the quantification columns
dataTotal <- as.data.frame(dataTotal)
TMT_columns = seq(2, 19, 1)
datTotal = dataTotal[TMT_columns]
rownames(datTotal) = dataTotal$Gene

# log2-transform the data
datTotal.log = log2(datTotal)

# remove rows with NAs
datTotal.log = na.omit(datTotal.log)
```

A design table is used to describe how samples are arranged in different groups/classes.

```
# make a design table to define grouping of columns
cond = as.factor(c("Prolif.Basal", "Prolif.Basal", "Prolif.Basal",
                   "Prolif.HeatShock", "Prolif.HeatShock", "Prolif.HeatShock",
                   "Quiesc.Basal", "Quiesc.Basal", "Quiesc.Basal",
                   "Quiesc.HeatShock", "Quiesc.HeatShock", "Quiesc.HeatShock",
                   "Senesc.Basal", "Senesc.Basal", "Senesc.Basal",
                   "Senesc.HeatShock", "Senesc.HeatShock", "Senesc.HeatShock"))

# generate the design matrix
design = model.matrix(~0 + cond)
colnames(design) = gsub("cond", "", colnames(design))
```

In addition to the design, we need to define the contrast, which inform the model which group comparisons are required.

```
# define contrasts
x <- c("Prolif.HeatShock-Prolif.Basal",
            "Quiesc.HeatShock-Quiesc.Basal",
            "Senesc.HeatShock-Senesc.Basal",
            "Quiesc.Basal-Prolif.Basal",
            "Senesc.Basal-Prolif.Basal",
            "Senesc.Basal-Quiesc.Basal",
            "Quiesc.HeatShock-Prolif.HeatShock",
            "Senesc.HeatShock-Prolif.HeatShock",
            "Senesc.HeatShock-Quiesc.HeatShock")
contrasts =  makeContrasts(contrasts = x, levels = design)

# make fits
```

```r
fit1Total <- lmFit(datTotal.log, design)
fit2Total <- contrasts.fit(fit1Total, contrasts = contrasts)
fit3Total <- eBayes(fit2Total)

# assign variable `count` to fit3 object,
# telling how many PSMs are quantified for each protein
psm.count.tableTotal = data.frame(
  count = as.matrix(dataTotal[, 20]), row.names =  dataTotal$Gene)

fit3Total$count = psm.count.tableTotal[rownames(fit3Total$coefficients), "count"]
fit4Total = DEqMS::spectraCounteBayes(fit3Total)
```

We can visualise the protein variance dependence on quantified PSMs (Fig. 22 & 23).

```r
# n=30 limits the boxplot to show only proteins quantified by <= 30 PSMs.
VarianceBoxplot(fit4Total, n = 30, main = "Variance Boxplot: Total", xlab = "PSM count")
```

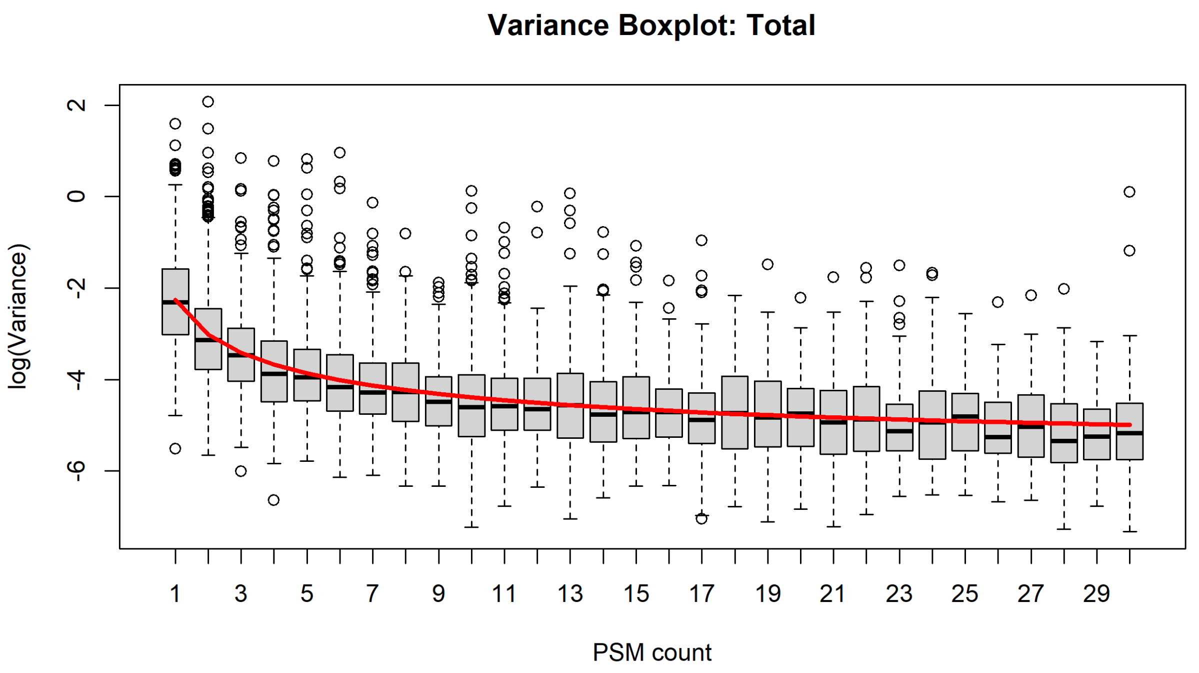

Figure 22: Boxplot showing variance of protein abundances with number of quantified PSMs in the total proteome of the IMR-90 dataset.

```r
VarianceScatterplot(fit4Total,
                    main = "Variance Scatterplot: Total", xlab = "log2(PSM count)")
```

Confirm that all the comparisons are included in the output.

```r
colnames(fit4Total$coefficients)
```

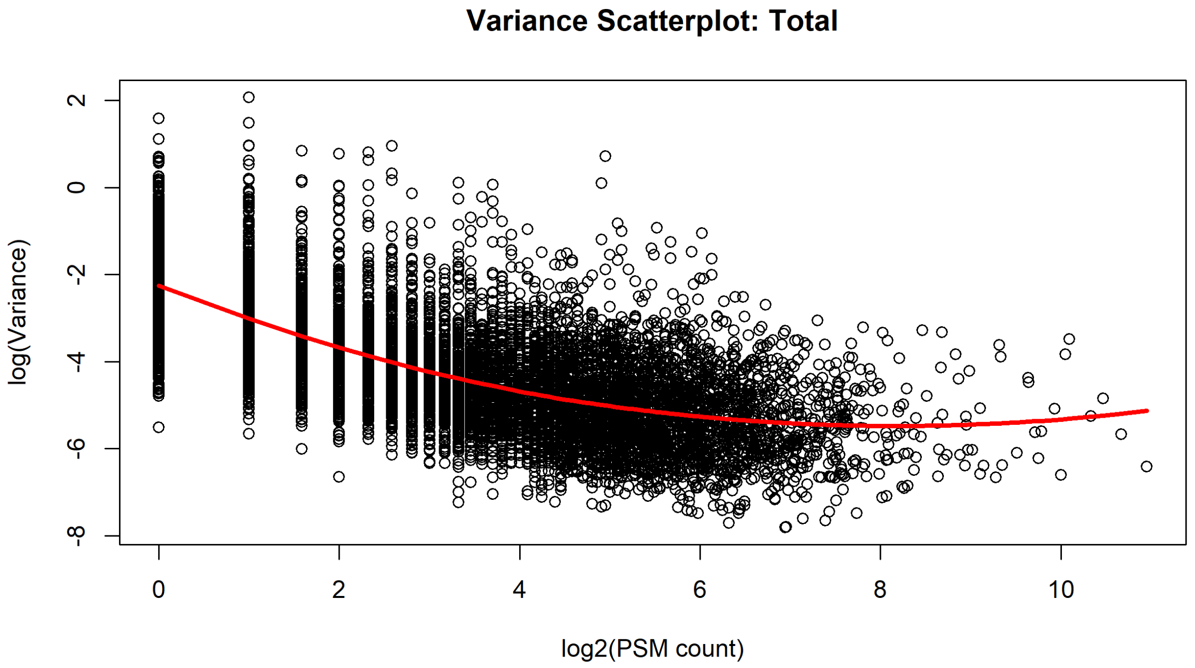

Figure 23: Scatterplot showing variance of protein abundances with number of quantified PSMs in the total proteome of the IMR-90 dataset.

```
## [1] "Prolif.HeatShock-Prolif.Basal"     "Quiesc.HeatShock-Quiesc.Basal"
## [3] "Senesc.HeatShock-Senesc.Basal"     "Quiesc.Basal-Prolif.Basal"
## [5] "Senesc.Basal-Prolif.Basal"         "Senesc.Basal-Quiesc.Basal"
## [7] "Quiesc.HeatShock-Prolif.HeatShock" "Senesc.HeatShock-Prolif.HeatShock"
## [9] "Senesc.HeatShock-Quiesc.HeatShock"
```

Using the above coordinates of the various comparisons, we can get separate data-frames for each comparison.

```
Total_Prolif.HeatShockVSProlif.Basal = outputResult(fit4Total, coef_col = 1)
Total_Quiesc.HeatShockVSQuiesc.Basal = outputResult(fit4Total, coef_col = 2)
Total_Senesc.HeatShockVSSenesc.Basal = outputResult(fit4Total, coef_col = 3)

Total_Quiesc.BasalVSProlif.Basal = outputResult(fit4Total, coef_col = 4)
Total_Senesc.BasalVSProlif.Basal = outputResult(fit4Total, coef_col = 5)
Total_Senesc.BasalVSQuiesc.Basal = outputResult(fit4Total, coef_col = 6)

Total_Quiesc.HeatShockVSProlif.HeatShock = outputResult(fit4Total, coef_col = 7)
Total_Senesc.HeatShockVSProlif.HeatShock = outputResult(fit4Total, coef_col = 8)
Total_Senesc.HeatShockVSQuiesc.HeatShock = outputResult(fit4Total, coef_col = 9)
```

Finally, we can extract only the columns we need from each data-frame, and add them to the original *dataImputed* object to make *dataAll*.

```
dataAll <- dataImputed

Total_Prolif.HeatShockVSProlif.Basal %>%
```

```r
  dplyr::select(gene, sca.P.Value, sca.adj.pval, logFC) %>%
  dplyr::rename(pval = sca.P.Value) %>%
  dplyr::rename(padj = sca.adj.pval) %>%
  rename_with(.fn = ~paste0("Total_Prolif.HeatShockVSProlif.Basal_", .),
              .cols = -gene ) %>%
  rename(Gene = gene) %>%
  right_join(dataAll) -> dataAll

Total_Quiesc.HeatShockVSQuiesc.Basal %>%
  dplyr::select(gene, sca.P.Value, sca.adj.pval, logFC) %>%
  dplyr::rename(pval = sca.P.Value) %>%
  dplyr::rename(padj = sca.adj.pval) %>%
  rename_with(.fn = ~paste0("Total_Quiesc.HeatShockVSQuiesc.Basal_", .),
              .cols = -gene ) %>%
  rename(Gene = gene) %>%
  right_join(dataAll) -> dataAll

Total_Senesc.HeatShockVSSenesc.Basal %>%
  dplyr::select(gene, sca.P.Value, sca.adj.pval, logFC) %>%
  dplyr::rename(pval = sca.P.Value) %>%
  dplyr::rename(padj = sca.adj.pval) %>%
  rename_with(.fn = ~paste0("Total_Senesc.HeatShockVSSenesc.Basal_", .),
              .cols = -gene ) %>%
  rename(Gene = gene) %>%
  right_join(dataAll) -> dataAll

Total_Quiesc.BasalVSProlif.Basal %>%
  dplyr::select(gene, sca.P.Value, sca.adj.pval, logFC) %>%
  dplyr::rename(pval = sca.P.Value) %>%
  dplyr::rename(padj = sca.adj.pval) %>%
  rename_with(.fn = ~paste0("Total_Quiesc.BasalVSProlif.Basal_", .),
              .cols = -gene ) %>%
  rename(Gene = gene) %>%
  right_join(dataAll) -> dataAll

Total_Senesc.BasalVSProlif.Basal %>%
  dplyr::select(gene, sca.P.Value, sca.adj.pval, logFC) %>%
  dplyr::rename(pval = sca.P.Value) %>%
  dplyr::rename(padj = sca.adj.pval) %>%
  rename_with(.fn = ~paste0("Total_Senesc.BasalVSProlif.Basal_", .),
              .cols = -gene ) %>%
  rename(Gene = gene) %>%
  right_join(dataAll) -> dataAll

Total_Senesc.BasalVSQuiesc.Basal %>%
  dplyr::select(gene, sca.P.Value, sca.adj.pval, logFC) %>%
  dplyr::rename(pval = sca.P.Value) %>%
  dplyr::rename(padj = sca.adj.pval) %>%
  rename_with(.fn = ~paste0("Total_Senesc.BasalVSQuiesc.Basal_", .),
              .cols = -gene ) %>%
  rename(Gene = gene) %>%
  right_join(dataAll) -> dataAll
```

```
Total_Quiesc.HeatShockVSProlif.HeatShock %>%
  dplyr::select(gene, sca.P.Value, sca.adj.pval, logFC) %>%
  dplyr::rename(pval = sca.P.Value) %>%
  dplyr::rename(padj = sca.adj.pval) %>%
  rename_with(.fn = ~paste0("Total_Quiesc.HeatShockVSProlif.HeatShock_", .),
              .cols = -gene ) %>%
  rename(Gene = gene) %>%
  right_join(dataAll) -> dataAll

Total_Senesc.HeatShockVSProlif.HeatShock %>%
  dplyr::select(gene, sca.P.Value, sca.adj.pval, logFC) %>%
  dplyr::rename(pval = sca.P.Value) %>%
  dplyr::rename(padj = sca.adj.pval) %>%
  rename_with(.fn = ~paste0("Total_Senesc.HeatShockVSProlif.HeatShock_", .),
              .cols = -gene ) %>%
  rename(Gene = gene) %>%
  right_join(dataAll) -> dataAll

Total_Senesc.HeatShockVSQuiesc.HeatShock %>%
  dplyr::select(gene, sca.P.Value, sca.adj.pval, logFC) %>%
  dplyr::rename(pval = sca.P.Value) %>%
  dplyr::rename(padj = sca.adj.pval) %>%
  rename_with(.fn = ~paste0("Total_Senesc.HeatShockVSQuiesc.HeatShock_", .),
              .cols = -gene ) %>%
  rename(Gene = gene) %>%
  right_join(dataAll) -> dataAll
```

We can repeat the same steps for the polyUb and insoluble proteomes.

###### 2.4.2 Differential Analysis: PolyUb Proteome

```
# define coordinates of the quantification columns
dataPolyUb <- as.data.frame(dataPolyUb)
datPolyUb = dataPolyUb[TMT_columns]
rownames(datPolyUb) = dataPolyUb$Gene

# log2-transform the data
datPolyUb.log = log2(datPolyUb)

# remove rows with NAs
datPolyUb.log = na.omit(datPolyUb.log)

# make fits
fit1PolyUb <- lmFit(datPolyUb.log, design)
fit2PolyUb <- contrasts.fit(fit1PolyUb, contrasts = contrasts)
fit3PolyUb <- eBayes(fit2PolyUb)

# assign variable `count` to fit3 object,
# telling how many PSMs are quantified for each protein
psm.count.tablePolyUb = data.frame(
```

```r
  count = as.matrix(dataPolyUb[, 20]), row.names =  dataPolyUb$Gene)

fit3PolyUb$count = psm.count.tablePolyUb[rownames(fit3PolyUb$coefficients), "count"]
fit4PolyUb = DEqMS::spectraCounteBayes(fit3PolyUb)
```

```r
# n=30 limits the boxplot to show only proteins quantified by <= 30 PSMs.
VarianceBoxplot(fit4PolyUb, n = 30, main = "Variance Boxplot: PolyUb", xlab = "PSM count")
```

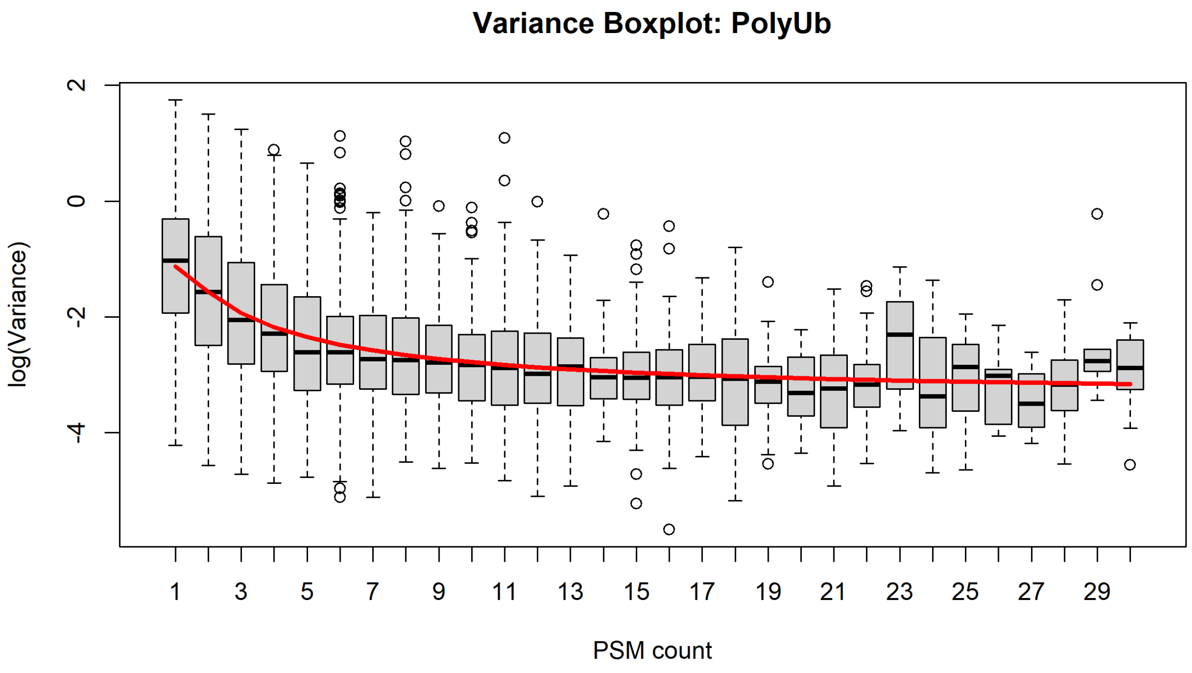

Figure 24: Boxplot showing variance of protein abundances with number of quantified PSMs in the polyUb proteome of the IMR-90 dataset.

```r
VarianceScatterplot(fit4PolyUb,
                    main = "Variance Scatterplot: PolyUb", xlab = "log2(PSM count)")
```

```r
PolyUb_Prolif.HeatShockVSProlif.Basal = outputResult(fit4PolyUb, coef_col = 1)
PolyUb_Quiesc.HeatShockVSQuiesc.Basal = outputResult(fit4PolyUb, coef_col = 2)
PolyUb_Senesc.HeatShockVSSenesc.Basal = outputResult(fit4PolyUb, coef_col = 3)

PolyUb_Quiesc.BasalVSProlif.Basal = outputResult(fit4PolyUb, coef_col = 4)
PolyUb_Senesc.BasalVSProlif.Basal = outputResult(fit4PolyUb, coef_col = 5)
PolyUb_Senesc.BasalVSQuiesc.Basal = outputResult(fit4PolyUb, coef_col = 6)

PolyUb_Quiesc.HeatShockVSProlif.HeatShock = outputResult(fit4PolyUb, coef_col = 7)
PolyUb_Senesc.HeatShockVSProlif.HeatShock = outputResult(fit4PolyUb, coef_col = 8)
PolyUb_Senesc.HeatShockVSQuiesc.HeatShock = outputResult(fit4PolyUb, coef_col = 9)
```

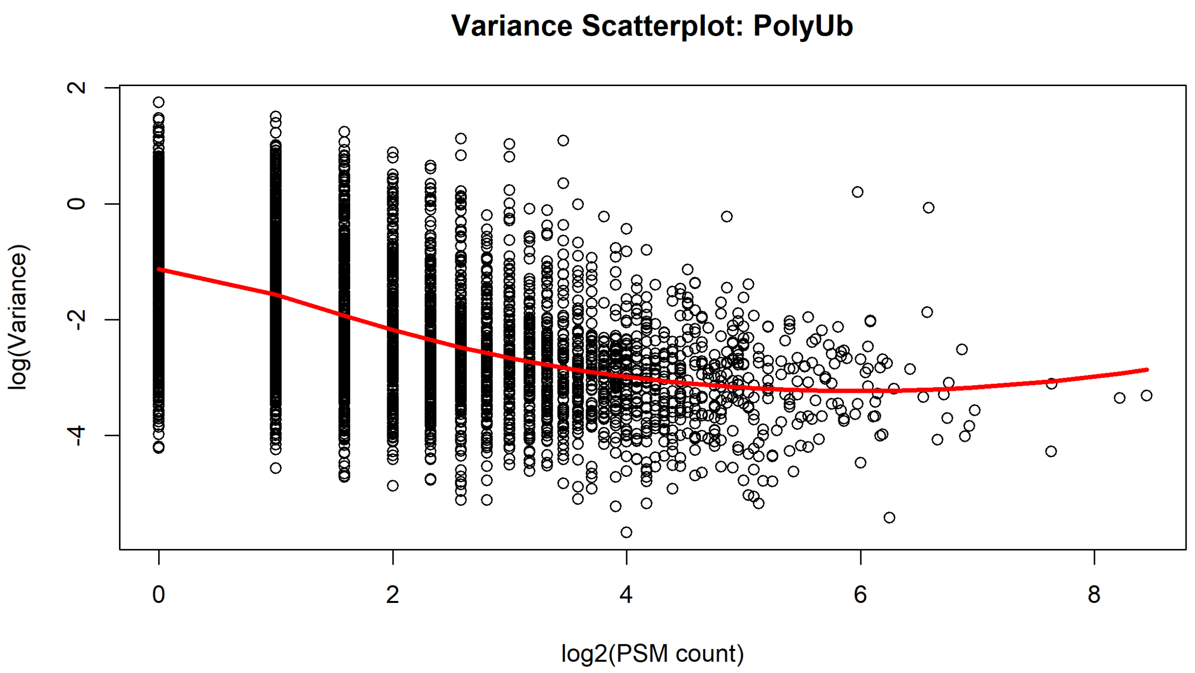

Figure 25: Scatterplot showing variance of protein abundances with number of quantified PSMs in the polyUb proteome of the IMR-90 dataset.

```
PolyUb_Prolif.HeatShockVSProlif.Basal %>%
  dplyr::select(gene, sca.P.Value, sca.adj.pval, logFC) %>%
  dplyr::rename(pval = sca.P.Value) %>%
  dplyr::rename(padj = sca.adj.pval) %>%
  rename_with(.fn = ~paste0("PolyUb_Prolif.HeatShockVSProlif.Basal_", .),
              .cols = -gene ) %>%
  rename(Gene = gene) %>%
  right_join(dataAll) -> dataAll

PolyUb_Quiesc.HeatShockVSQuiesc.Basal %>%
  dplyr::select(gene, sca.P.Value, sca.adj.pval, logFC) %>%
  dplyr::rename(pval = sca.P.Value) %>%
  dplyr::rename(padj = sca.adj.pval) %>%
  rename_with(.fn = ~paste0("PolyUb_Quiesc.HeatShockVSQuiesc.Basal_", .),
              .cols = -gene ) %>%
  rename(Gene = gene) %>%
  right_join(dataAll) -> dataAll

PolyUb_Senesc.HeatShockVSSenesc.Basal %>%
  dplyr::select(gene, sca.P.Value, sca.adj.pval, logFC) %>%
  dplyr::rename(pval = sca.P.Value) %>%
  dplyr::rename(padj = sca.adj.pval) %>%
  rename_with(.fn = ~paste0("PolyUb_Senesc.HeatShockVSSenesc.Basal_", .),
              .cols = -gene ) %>%
  rename(Gene = gene) %>%
  right_join(dataAll) -> dataAll
```

```r
PolyUb_Quiesc.BasalVSProlif.Basal %>%
  dplyr::select(gene, sca.P.Value, sca.adj.pval, logFC) %>%
  dplyr::rename(pval = sca.P.Value) %>%
  dplyr::rename(padj = sca.adj.pval) %>%
  rename_with(.fn = ~paste0("PolyUb_Quiesc.BasalVSProlif.Basal_", .),
              .cols = -gene ) %>%
  rename(Gene = gene) %>%
  right_join(dataAll) -> dataAll

PolyUb_Senesc.BasalVSProlif.Basal %>%
  dplyr::select(gene, sca.P.Value, sca.adj.pval, logFC) %>%
  dplyr::rename(pval = sca.P.Value) %>%
  dplyr::rename(padj = sca.adj.pval) %>%
  rename_with(.fn = ~paste0("PolyUb_Senesc.BasalVSProlif.Basal_", .),
              .cols = -gene ) %>%
  rename(Gene = gene) %>%
  right_join(dataAll) -> dataAll

PolyUb_Senesc.BasalVSQuiesc.Basal %>%
  dplyr::select(gene, sca.P.Value, sca.adj.pval, logFC) %>%
  dplyr::rename(pval = sca.P.Value) %>%
  dplyr::rename(padj = sca.adj.pval) %>%
  rename_with(.fn = ~paste0("PolyUb_Senesc.BasalVSQuiesc.Basal_", .),
              .cols = -gene ) %>%
  rename(Gene = gene) %>%
  right_join(dataAll) -> dataAll

PolyUb_Quiesc.HeatShockVSProlif.HeatShock %>%
  dplyr::select(gene, sca.P.Value, sca.adj.pval, logFC) %>%
  dplyr::rename(pval = sca.P.Value) %>%
  dplyr::rename(padj = sca.adj.pval) %>%
  rename_with(.fn = ~paste0("PolyUb_Quiesc.HeatShockVSProlif.HeatShock_", .),
              .cols = -gene ) %>%
  rename(Gene = gene) %>%
  right_join(dataAll) -> dataAll

PolyUb_Senesc.HeatShockVSProlif.HeatShock %>%
  dplyr::select(gene, sca.P.Value, sca.adj.pval, logFC) %>%
  dplyr::rename(pval = sca.P.Value) %>%
  dplyr::rename(padj = sca.adj.pval) %>%
  rename_with(.fn = ~paste0("PolyUb_Senesc.HeatShockVSProlif.HeatShock_", .),
              .cols = -gene ) %>%
  rename(Gene = gene) %>%
  right_join(dataAll) -> dataAll

PolyUb_Senesc.HeatShockVSQuiesc.HeatShock %>%
  dplyr::select(gene, sca.P.Value, sca.adj.pval, logFC) %>%
  dplyr::rename(pval = sca.P.Value) %>%
  dplyr::rename(padj = sca.adj.pval) %>%
  rename_with(.fn = ~paste0("PolyUb_Senesc.HeatShockVSQuiesc.HeatShock_", .),
              .cols = -gene ) %>%
```

```
  rename(Gene = gene) %>%
  right_join(dataAll) -> dataAll
```

##### 2.4.3 Differential Analysis: Insoluble Proteome

```
# define coordinates of the quantification columns
dataInsoluble <- as.data.frame(dataInsoluble)
datInsoluble = dataInsoluble[TMT_columns]
rownames(datInsoluble) = dataInsoluble$Gene

# log2-transform the data
datInsoluble.log = log2(datInsoluble)

# remove rows with NAs
datInsoluble.log = na.omit(datInsoluble.log)

# make fits
fit1Insoluble <- lmFit(datInsoluble.log, design)
fit2Insoluble <- contrasts.fit(fit1Insoluble,contrasts = contrasts)
fit3Insoluble <- eBayes(fit2Insoluble)

# assign variable `count` to fit3 object,
# telling how many PSMs are quantified for each protein
psm.count.tableInsoluble = data.frame(
  count = as.matrix(dataInsoluble[, 20]), row.names =  dataInsoluble$Gene)

fit3Insoluble$count = psm.count.tableInsoluble[rownames(
  fit3Insoluble$coefficients), "count"]
fit4Insoluble = DEqMS::spectraCounteBayes(fit3Insoluble)
```

```
# n=30 limits the boxplot to show only proteins quantified by <= 30 PSMs.
VarianceBoxplot(fit4Insoluble, n = 30,
                main = "Variance Boxplot: Insoluble", xlab = "PSM count")
```

```
VarianceScatterplot(fit4Insoluble,
                    main = "Variance Scatterplot: Insoluble", xlab = "log2(PSM count)")
```

```
Insoluble_Prolif.HeatShockVSProlif.Basal = outputResult(fit4Insoluble, coef_col = 1)
Insoluble_Quiesc.HeatShockVSQuiesc.Basal = outputResult(fit4Insoluble, coef_col = 2)
Insoluble_Senesc.HeatShockVSSenesc.Basal = outputResult(fit4Insoluble, coef_col = 3)

Insoluble_Quiesc.BasalVSProlif.Basal = outputResult(fit4Insoluble, coef_col = 4)
Insoluble_Senesc.BasalVSProlif.Basal = outputResult(fit4Insoluble, coef_col = 5)
Insoluble_Senesc.BasalVSQuiesc.Basal = outputResult(fit4Insoluble, coef_col = 6)

Insoluble_Quiesc.HeatShockVSProlif.HeatShock = outputResult(fit4Insoluble, coef_col = 7)
Insoluble_Senesc.HeatShockVSProlif.HeatShock = outputResult(fit4Insoluble, coef_col = 8)
Insoluble_Senesc.HeatShockVSQuiesc.HeatShock = outputResult(fit4Insoluble, coef_col = 9)
```

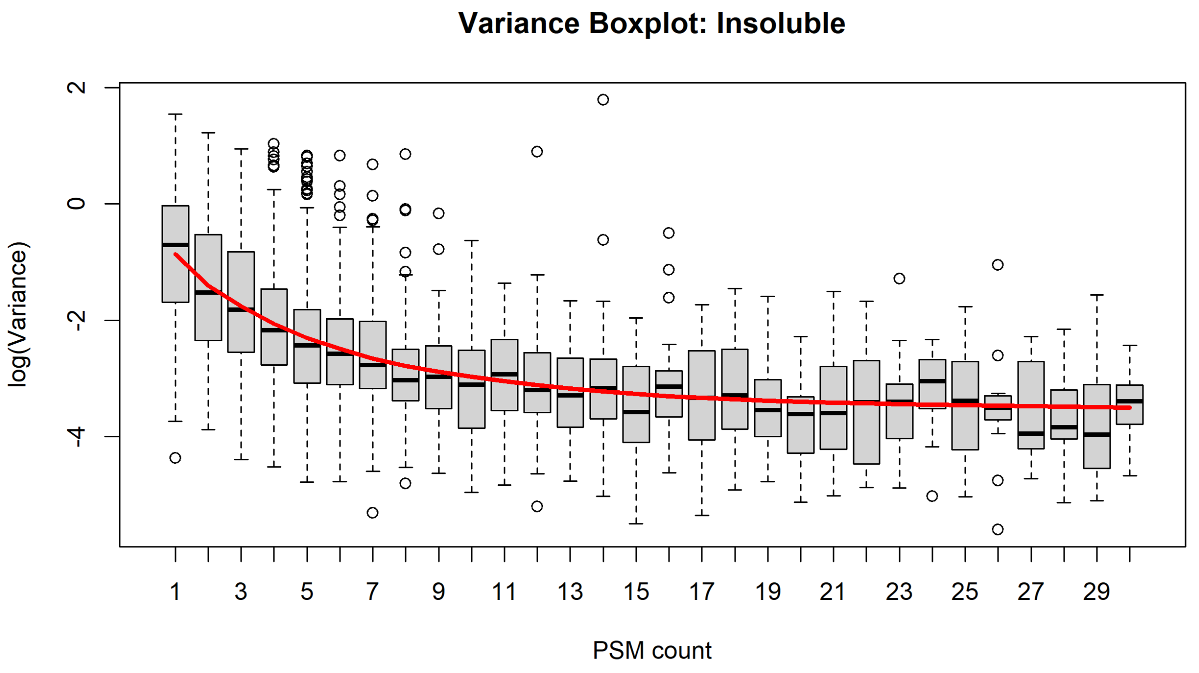

Figure 26: Boxplot showing variance of protein abundances with number of quantified PSMs in the insoluble proteome of the IMR-90 dataset.

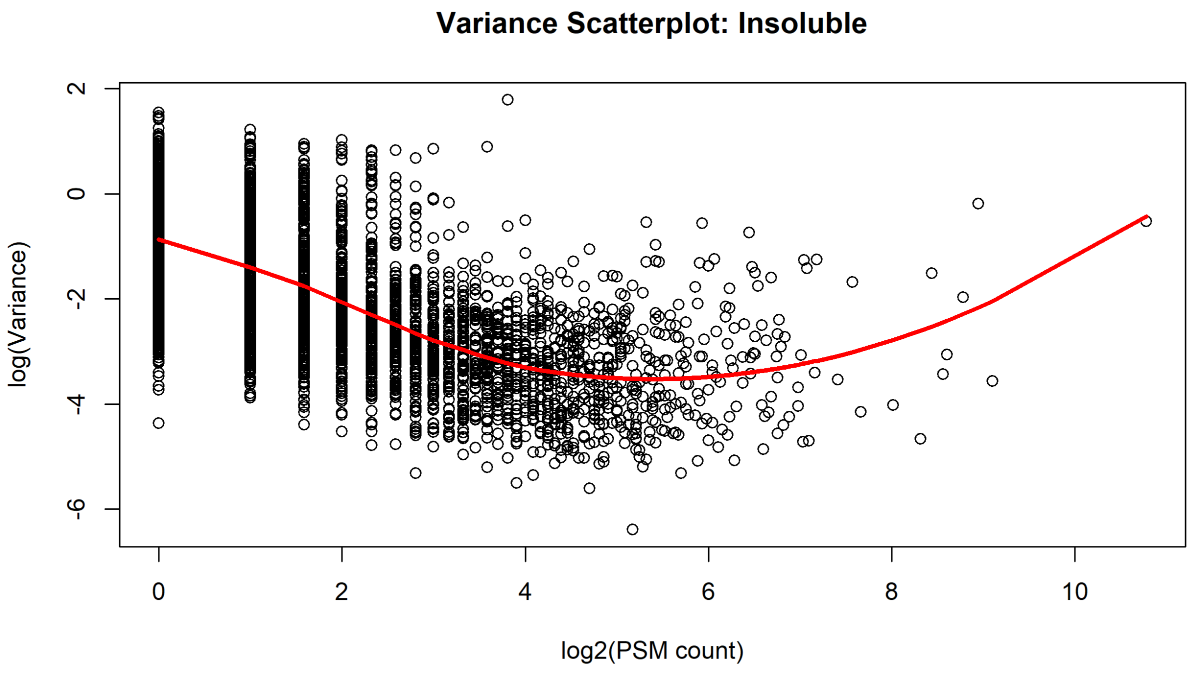

Figure 27: Scatterplot showing variance of protein abundances with number of quantified PSMs in the insoluble proteome of the IMR-90 dataset.

```r
Insoluble_Prolif.HeatShockVSProlif.Basal %>%
  dplyr::select(gene, sca.P.Value, sca.adj.pval, logFC) %>%
  dplyr::rename(pval = sca.P.Value) %>%
  dplyr::rename(padj = sca.adj.pval) %>%
  rename_with(.fn = ~paste0("Insoluble_Prolif.HeatShockVSProlif.Basal_", .),
              .cols = -gene ) %>%
  rename(Gene = gene) %>%
  right_join(dataAll) -> dataAll

Insoluble_Quiesc.HeatShockVSQuiesc.Basal %>%
  dplyr::select(gene, sca.P.Value, sca.adj.pval, logFC) %>%
  dplyr::rename(pval = sca.P.Value) %>%
  dplyr::rename(padj = sca.adj.pval) %>%
  rename_with(.fn = ~paste0("Insoluble_Quiesc.HeatShockVSQuiesc.Basal_", .),
              .cols = -gene ) %>%
  rename(Gene = gene) %>%
  right_join(dataAll) -> dataAll

Insoluble_Senesc.HeatShockVSSenesc.Basal %>%
  dplyr::select(gene, sca.P.Value, sca.adj.pval, logFC) %>%
  dplyr::rename(pval = sca.P.Value) %>%
  dplyr::rename(padj = sca.adj.pval) %>%
  rename_with(.fn = ~paste0("Insoluble_Senesc.HeatShockVSSenesc.Basal_", .),
              .cols = -gene ) %>%
  rename(Gene = gene) %>%
  right_join(dataAll) -> dataAll

Insoluble_Quiesc.BasalVSProlif.Basal %>%
  dplyr::select(gene, sca.P.Value, sca.adj.pval, logFC) %>%
  dplyr::rename(pval = sca.P.Value) %>%
  dplyr::rename(padj = sca.adj.pval) %>%
  rename_with(.fn = ~paste0("Insoluble_Quiesc.BasalVSProlif.Basal_", .),
              .cols = -gene ) %>%
  rename(Gene = gene) %>%
  right_join(dataAll) -> dataAll

Insoluble_Senesc.BasalVSProlif.Basal %>%
  dplyr::select(gene, sca.P.Value, sca.adj.pval, logFC) %>%
  dplyr::rename(pval = sca.P.Value) %>%
  dplyr::rename(padj = sca.adj.pval) %>%
  rename_with(.fn = ~paste0("Insoluble_Senesc.BasalVSProlif.Basal_", .),
              .cols = -gene ) %>%
  rename(Gene = gene) %>%
  right_join(dataAll) -> dataAll

Insoluble_Senesc.BasalVSQuiesc.Basal %>%
  dplyr::select(gene, sca.P.Value, sca.adj.pval, logFC) %>%
  dplyr::rename(pval = sca.P.Value) %>%
  dplyr::rename(padj = sca.adj.pval) %>%
  rename_with(.fn = ~paste0("Insoluble_Senesc.BasalVSQuiesc.Basal_", .),
              .cols = -gene ) %>%
  rename(Gene = gene) %>%
```

```
  right_join(dataAll) -> dataAll
```

###### 2.4.4 Export for downstream analysis

*dataAll* can now be written as **data_DEqMS.csv**, ready for downstream analysis.

```
dataAll %>%
  write_csv("IMR90/data_DEqMS.csv")
```

### 3 A549 Heat-Shock Experiment

Much of the same processing workflow performed for the IMR-90 experiment above can be used for the A549 experiments. However, these experiments differ in a few key ways.

1. There are only two cell states (proliferating and senescent); the quiescent state was not considered relevant to a cancer cell line (especially as contact inhibition cannot be achieved).
2. DNA-Damage-Induced Senescence was achieved using bleomycin (as per an established protocol with this cell line (Aoshiba, Tsuji, and Nagai 2003)), rather than doxorubicin.
3. Two different proteotoxic stresses were used: cells were either exposed to 2 h of heat-shock at 44 °C, or 24 h of the proteasome inhibitor bortezomib (100 nM). To control for the bortezomib treatment, an additional condition with the equivalent volume of vehicle control (DMSO) was also included.
4. Only the total and poly-ubiquitin (polyUb)-enriched proteomes were quantified (the insoluble proteome was not collected).
5. In order to accommodate the bortezomib and vehicle-control conditions, the samples were split into two TMT18-plex experiments. The additional channels were exploited to increase the number of replicates to four (rather than three), and the inclusion of two 'bridging' reference channels, comprising a pool of equal amounts of each of the 32 'biological' samples.

The two TMT experiments (basal vs. heat-shock, and vehicle vs. bortezomib) have been processed separately. We will start with the heat-shock experiment, as this is the most similar to the IMR-90 experiment above.

Note that code-chunks for many of the steps were identical or near-identical to the IMR-90 workflow, and therefore have mostly been omitted in the following sections.

#### 3.1 Initial Data Clean-Up and Filtering

```
rm(list = setdiff(ls(), c("get_gene_from_uniprot")))

library(readxl)
read_xlsx("../../A549_Sen44_BTZ/Sen44_Tot_TRT_Combined/TMT18_A549_44_Tot_TRT_prot.xlsx"
          ) -> data

dim(data)
```

```
## [1] 9200  178
```

The heat-shock data-frame has 9,200 entries (i.e., proteins identified in the experiment), and 178 columns.

There are 36 abundance columns for each data-frame: one for each TMT18-plex sample, across total (F1) and polyUb (F2) proteomes.

We can use these values to create our experimental_design, as before.

```
## # A tibble: 6 x 9
##   sample    sampleID TMT   conditionID label condition state treatment replicate
##   <chr>     <chr>    <chr> <chr>       <chr> <chr>     <chr> <chr>         <int>
## 1 Abundanc~ Abundan~ 127C  " 1"        Prol~ Prolif.B~ Prol~ Basal             1
## 2 Abundanc~ Abundan~ 128C  " 1"        Prol~ Prolif.B~ Prol~ Basal             2
## 3 Abundanc~ Abundan~ 129C  " 1"        Prol~ Prolif.B~ Prol~ Basal             3
## 4 Abundanc~ Abundan~ 130C  " 1"        Prol~ Prolif.B~ Prol~ Basal             4
## 5 Abundanc~ Abundan~ 131C  " 2"        Prol~ Prolif.H~ Prol~ HeatShock         1
## 6 Abundanc~ Abundan~ 132C  " 2"        Prol~ Prolif.H~ Prol~ HeatShock         2
```

###### 3.1.1 Clean-up of Gene and Accession identifiers

###### 3.1.1.1 Filling in missing Gene Symbols

```
## # A tibble: 2 x 2
## # Groups:   is.na(Gene) [2]
##   `is.na(Gene)`     n
##   <lgl>         <int>
## 1 FALSE          9135
## 2 TRUE             65
```

We can use our custom function 'get_gene_from_uniprot' to attempt to assign these 65 missing Genes automatically from UniProt.

```
## # A tibble: 2 x 2
## # Groups:   is.na(Gene) [2]
##   `is.na(Gene)`     n
##   <lgl>         <int>
## 1 FALSE          9189
## 2 TRUE             11
```

These 11 remaining missing Gene names need to be annotated manually.

```
## # A tibble: 11 x 2
##    Accession       Description
##    <chr>           <chr>
##  1 Cont_X00000     Halo-TR-TUBE protein OS=Escherichia coli OX=0000 GN=HaloTEVT~
##  2 Cont_P00761     Trypsin OS=Sus scrofa OX=9823 PE=1 SV=1
##  3 Cont_P02081     Hemoglobin fetal subunit beta OS=Bos taurus OX=9913 PE=1 SV=1
##  4 Q6ZSR9          Uncharacterized protein FLJ45252 OS=Homo sapiens OX=9606 PE=~
##  5 Cont_G5E513     Uncharacterized protein OS=Bos taurus OX=9913 PE=1 SV=2
##  6 Cont_A0A3Q1M3L6 Uncharacterized protein OS=Bos taurus OX=9913 PE=1 SV=1
##  7 Cont_P50448     Factor XIIa inhibitor OS=Bos taurus OX=9913 PE=1 SV=1
##  8 Cont_E1BCW0     HGF activator OS=Bos taurus OX=9913 GN=HGFAC PE=4 SV=3
##  9 Cont_G3N188     Ig-like domain-containing protein OS=Bos taurus OX=9913 PE=4~
## 10 Cont_Q2KIT0     Protein HP-20 homolog OS=Bos taurus OX=9913 PE=2 SV=1
## 11 Q8NFD4          Uncharacterized protein FLJ76381 OS=Homo sapiens OX=9606 PE=~
```

As before, most of the missing Gene names are contaminants. For these, we will copy the Accessions into the Gene column.

```
## # A tibble: 2 x 2
##   Accession Description
##   <chr>     <chr>
## 1 Q6ZSR9    Uncharacterized protein FLJ45252 OS=Homo sapiens OX=9606 PE=2 SV=2
## 2 Q8NFD4    Uncharacterized protein FLJ76381 OS=Homo sapiens OX=9606 PE=2 SV=1
```

For the remaining missing Genes, we will manually fill these in manually.

```
data %>%
  mutate(Gene = replace(Gene, Accession == "Q6ZSR9", "FLJ45252")) %>%
  mutate(Gene = replace(Gene, Accession == "Q8NFD4", "FLJ76381")) -> data

data %>%
  dplyr::filter(is.na(Gene)) %>%
  dplyr::count()
```

```
## # A tibble: 1 x 1
##       n
##   <int>
## 1     0
```

We will rename the ubiquitin Gene again.

```
data %>%
  mutate(Gene = replace(Gene, Accession == "P0CG47", "UBB")) -> data
```

###### 3.1.1.2 Making duplicated Genes unique

```
## # A tibble: 0 x 2
## # Groups:   Accession [0]
## # i 2 variables: Accession <chr>, n <int>
```

```
## # A tibble: 0 x 2
## # Groups:   Gene [0]
## # i 2 variables: Gene <chr>, n <int>
```

There are no duplicated Accessions or Genes.

##### 3.1.2 Adding proteome annotations

##### 3.1.3 Removing contaminants

Next, we will remove contaminants.

```
## # A tibble: 2 x 2
## # Groups:   Contaminant [2]
##   Contaminant     n
##   <lgl>       <int>
## 1 FALSE        9054
## 2 TRUE          146
```

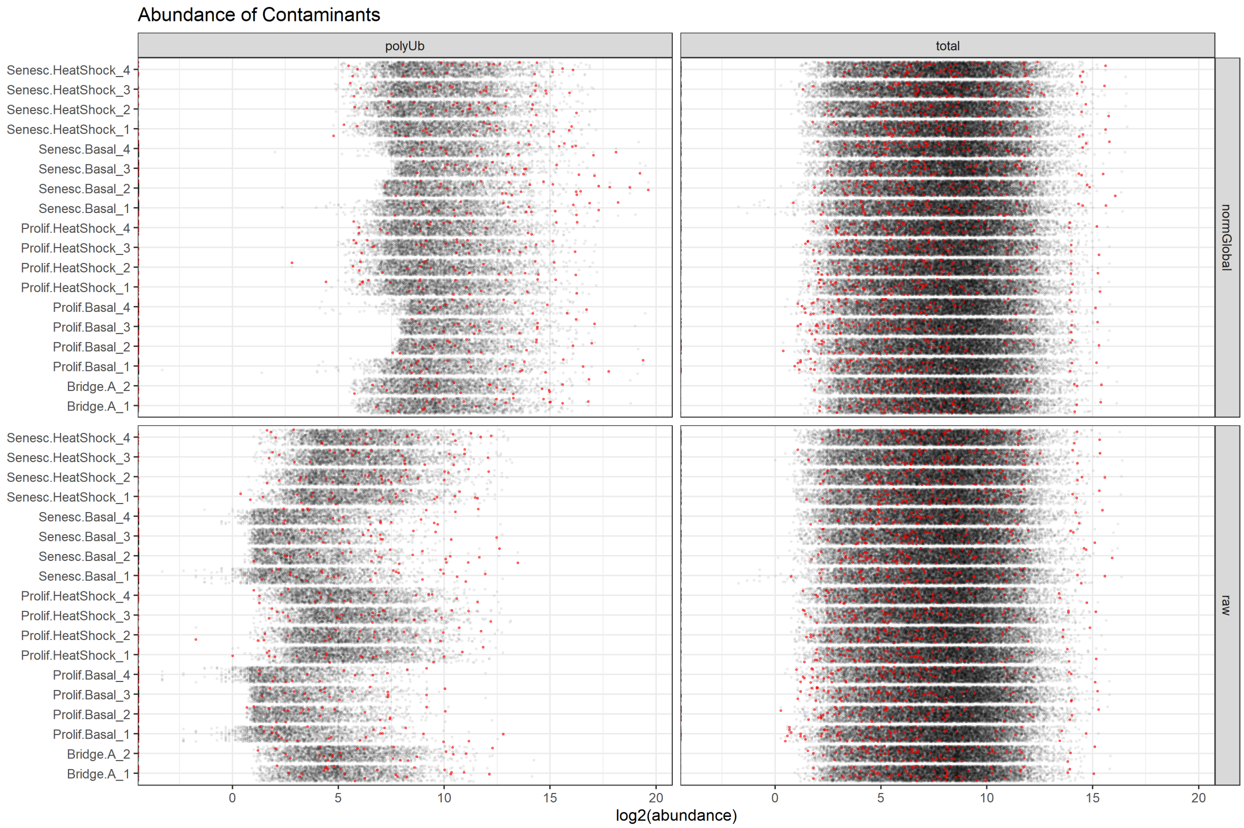

Figure 28: The raw abundance distribution of known contaminants in the A549 heat-shock dataset.

Again, based on these plots, it doesn't seem like there is any clear bias in the distribution of contaminants between the samples. We will remove these contaminants.

##### 3.1.4 Dealing with single PSM/peptide identifications

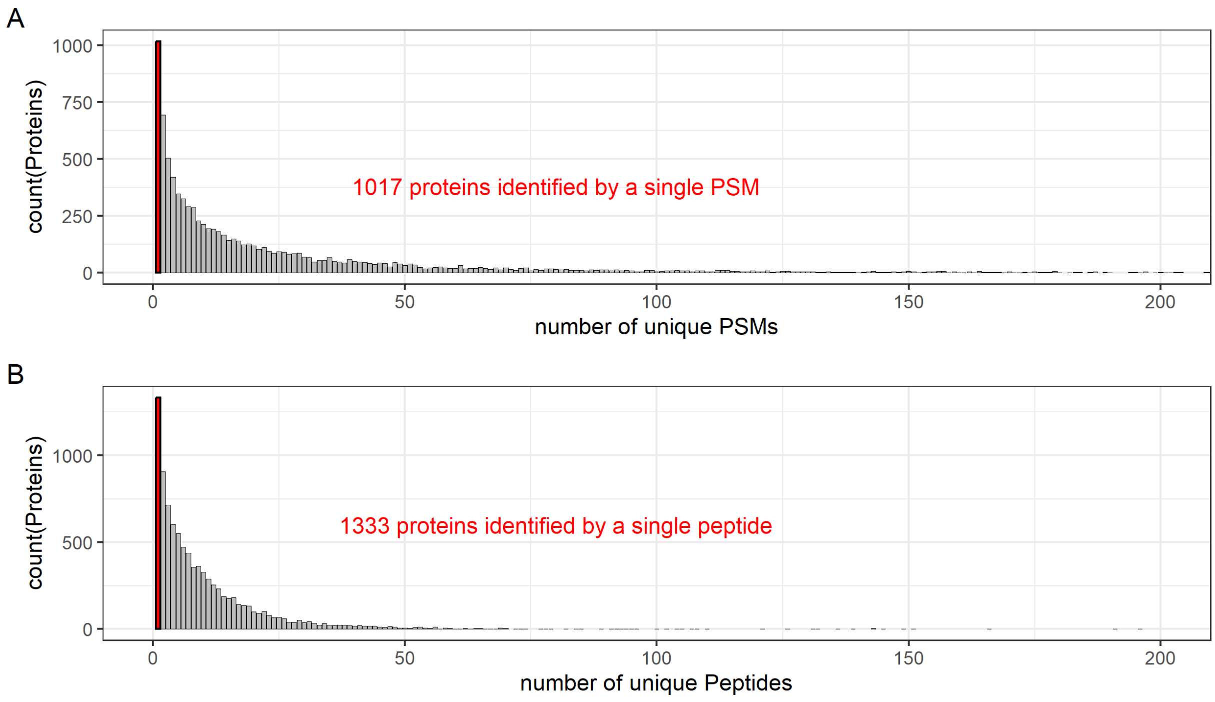

Figure 29: Histogram showing distribution of proteins identified in the A549 heat-shock dataset by the number of unique PSMs (A) or peptides (B).

```
dataNoContsNoSinglePSMs %>%
  dplyr::select(-`# PSMs`, -`# Peptides`) %>%

  write_csv("A549_heat-shock/dataFiltered.csv",
            na = "NA", append = FALSE, col_names = TRUE, escape = "double")

dim(dataNoContsNoSinglePSMs)
```

```
## [1] 8037   76
```

#### 3.2 Data Normalisation

##### 3.2.1 Assessing global normalisation

```
HS_sample_colours <- c('darkgrey', 'red4', 'red', 'turquoise4', 'cyan')

# a function 'col2hex' to find the HEX codes for these colours
# (e.g., for replicating them in Illustrator)
```

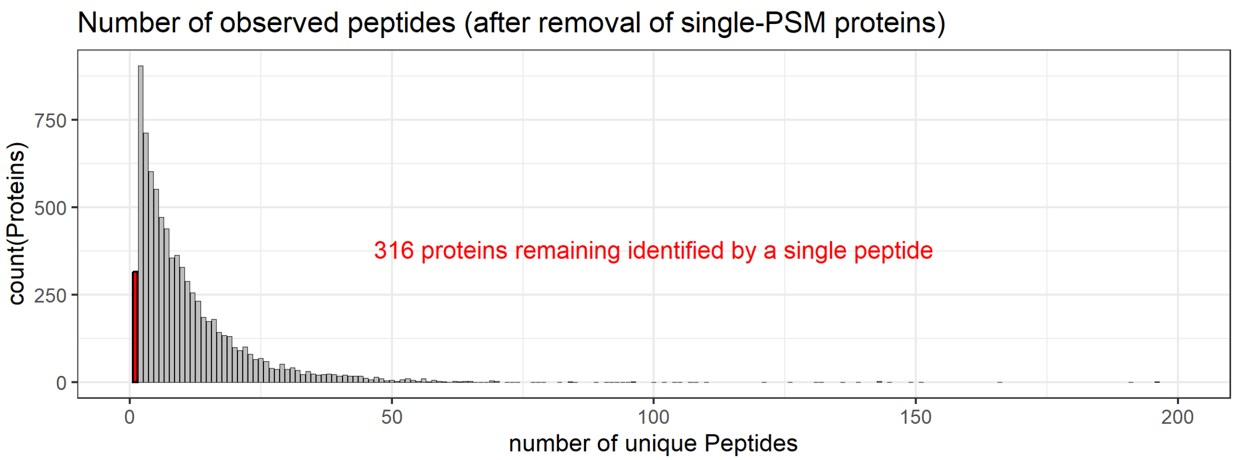

Figure 30: Histogram showing distribution of proteins identified in the A549 heat-shock dataset by the number of unique peptides, after the 1,017 proteins identified by single PSMs had been removed.

```
col2hex <- function(x, alpha = FALSE) {
  args <- as.data.frame(t(col2rgb(x, alpha = alpha)))
  args <- c(args, list(names = x, maxColorValue = 255))
  do.call(rgb, args)
}

col2hex(HS_sample_colours)
```

```
##    darkgrey        red4         red turquoise4        cyan
##   "#A9A9A9"   "#8B0000"   "#FF0000"   "#00868B"   "#00FFFF"
```

These distributions look very similar to those in the IMR-90 dataset. We can therefore use the same low-CV normalisation strategy, i.e., by defining the 10 % of the proteins in each proteome that have the lowest coefficients of variation (CVs) across all the samples, and then performing a size-factor normalisation based on these correction factors.

##### 3.2.2 Calculate low-CV genes

```
## # A tibble: 2 x 2
##   proteome nProteins
##   <chr>        <int>
## 1 polyUb         216
## 2 total          724
```

##### 3.2.3 Calculate size-factors based on low-CV genes

##### 3.2.4 Normalising data based on low-CV-calculated size-factors

For the remainder of the analysis, we will proceed with the CV-normalised abundances for the polyUb proteomes, and keep the global-normalised total proteomes.

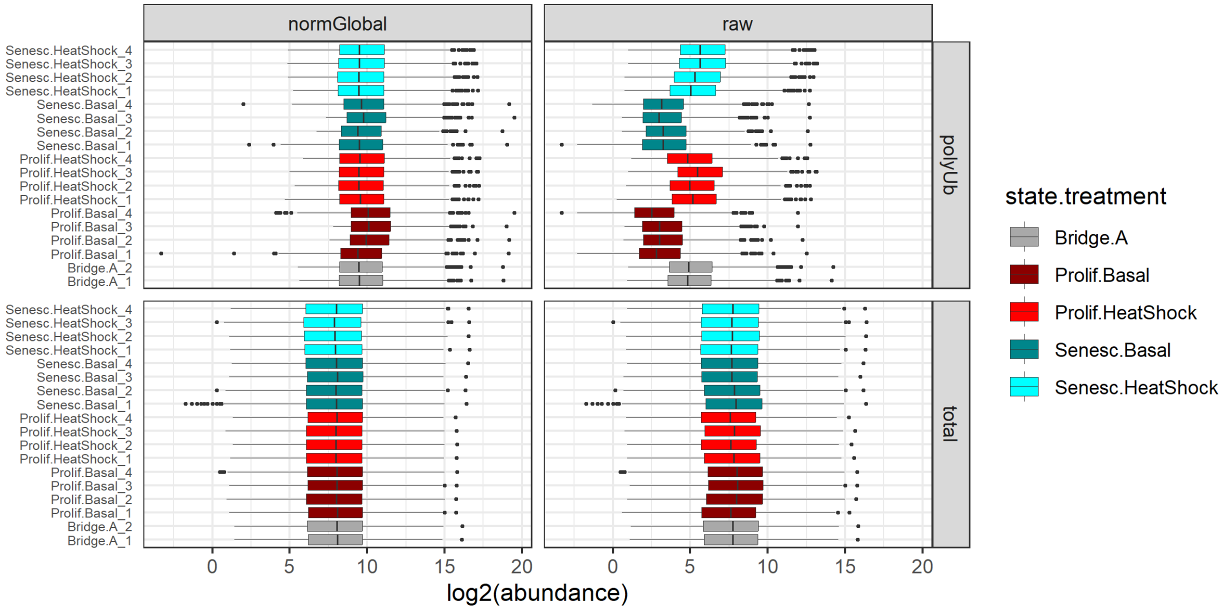

Figure 31: Raw and normalised log2-tranformed abundance distributions of all proteins identified in each sample of the A549 heat-shock experiment.

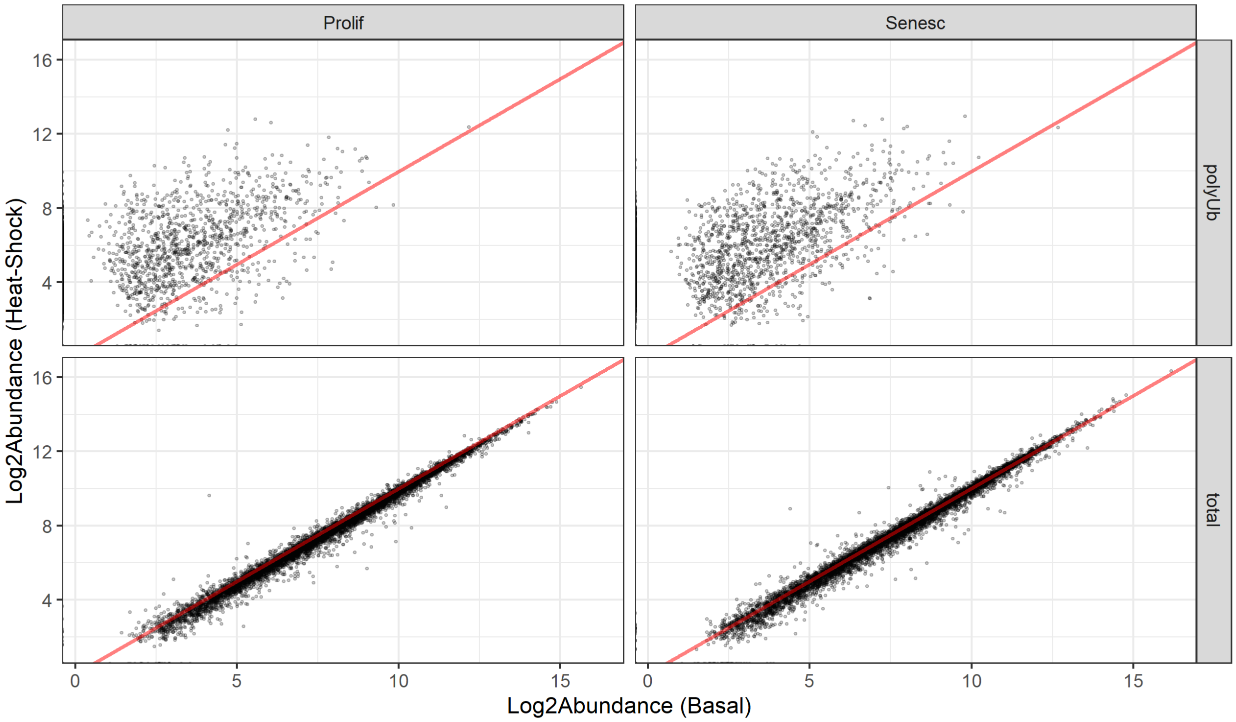

Figure 32: Scatter-plots of mean raw protein abundances for each state in the A549 heat-shock experiment. Each point represents the mean raw abundance of an individual protein in the basal vs. heat-shocked samples.

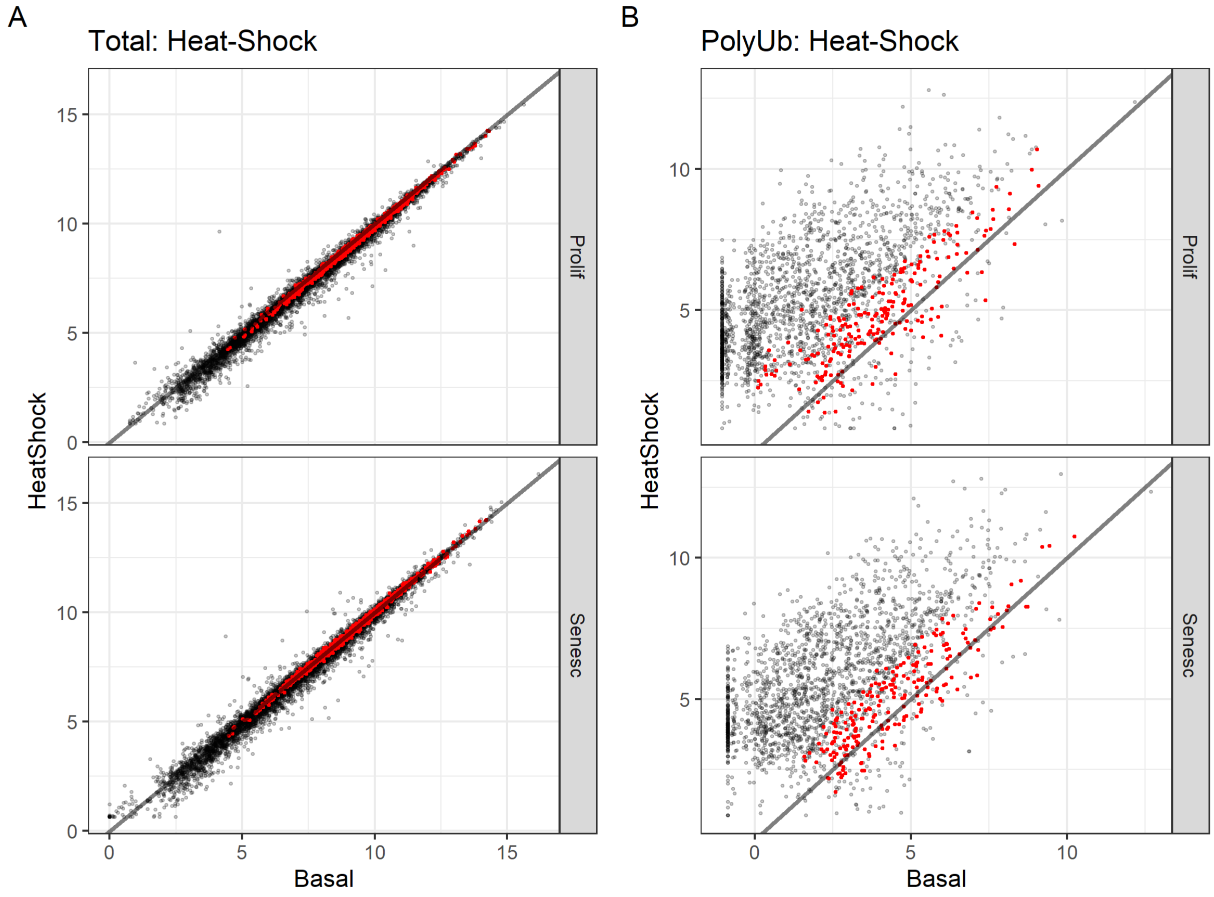

Figure 33: Lowest 10% CV proteins for each proteome from the A549 heat-shock experiment, mapped onto scatter-plots of mean raw protein abundances. Each point represents the mean raw abundance of an individual protein in the basal vs. heat-shocked samples. The proteins with the 10% lowest CVs for each proteome are plotted in red.

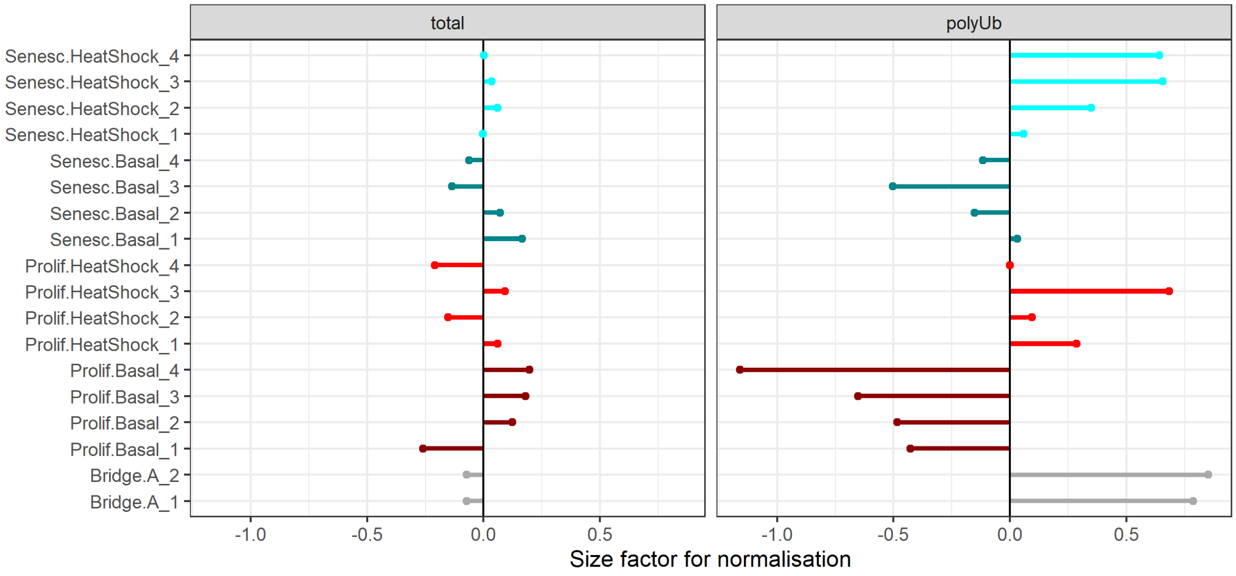

Figure 34: Size factors required for normalisation of each sample in the A549 heat-shock dataset, based on the mean 10% low-CV protein values.

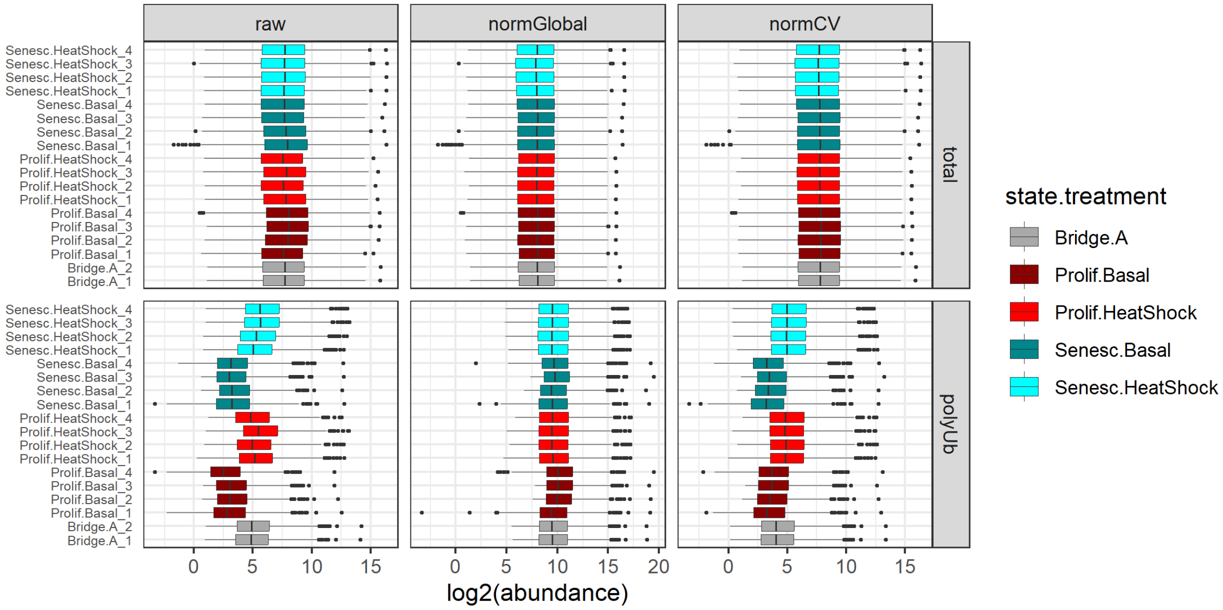

Figure 35: Raw and normalised protein abundance distributions in each sample of the A549 heat-shock dataset.

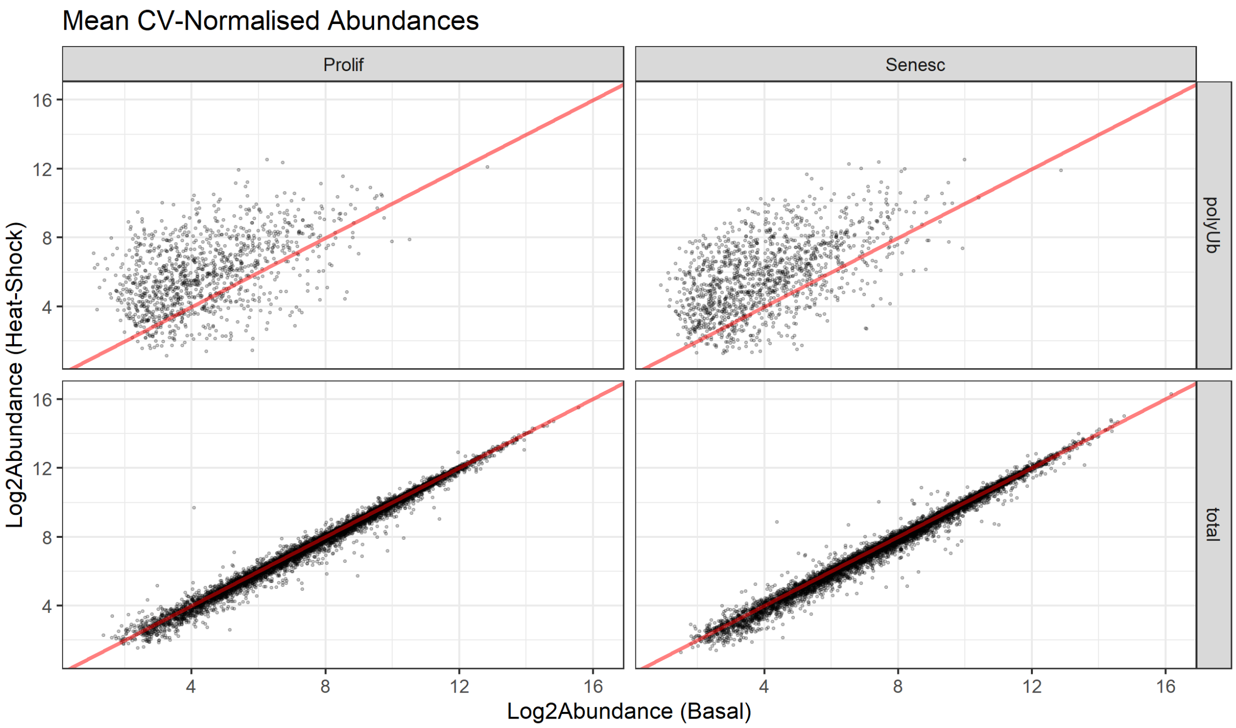

Figure 36: Scatter-plots of mean CV-normalised protein abundances of each state in the A549 heat-shock experiment. Each point represents the mean CV-normalised abundance of an individual protein in the basal vs. heat-shocked samples.

```
# save all abundance values as "dataNormalised_All.csv"
dataNormalised_All %>%
  write_csv("A549_heat-shock/dataNormalised_All.csv",
            na = "NA", append = FALSE, col_names = TRUE, escape = "double")

# save only selected normalised abundances as "dataNormalised_Selected.csv"
dataNormalised_All %>%
  dplyr::select(-starts_with("raw")) %>%
  dplyr::select(-starts_with(c("normCV_total"))) %>%
  dplyr::select(-starts_with(c("normGlobal_polyUb"))) %>%
  rename_with(~ gsub("normCV_", "", .x, fixed = TRUE)) %>%
  rename_with(~ gsub("normGlobal_", "", .x, fixed = TRUE)) -> dataNormalised_selected

dataNormalised_selected %>%
  write_csv("A549_heat-shock/dataNormalised_Selected.csv",
            na = "NA", append = FALSE, col_names = TRUE, escape = "double")
```

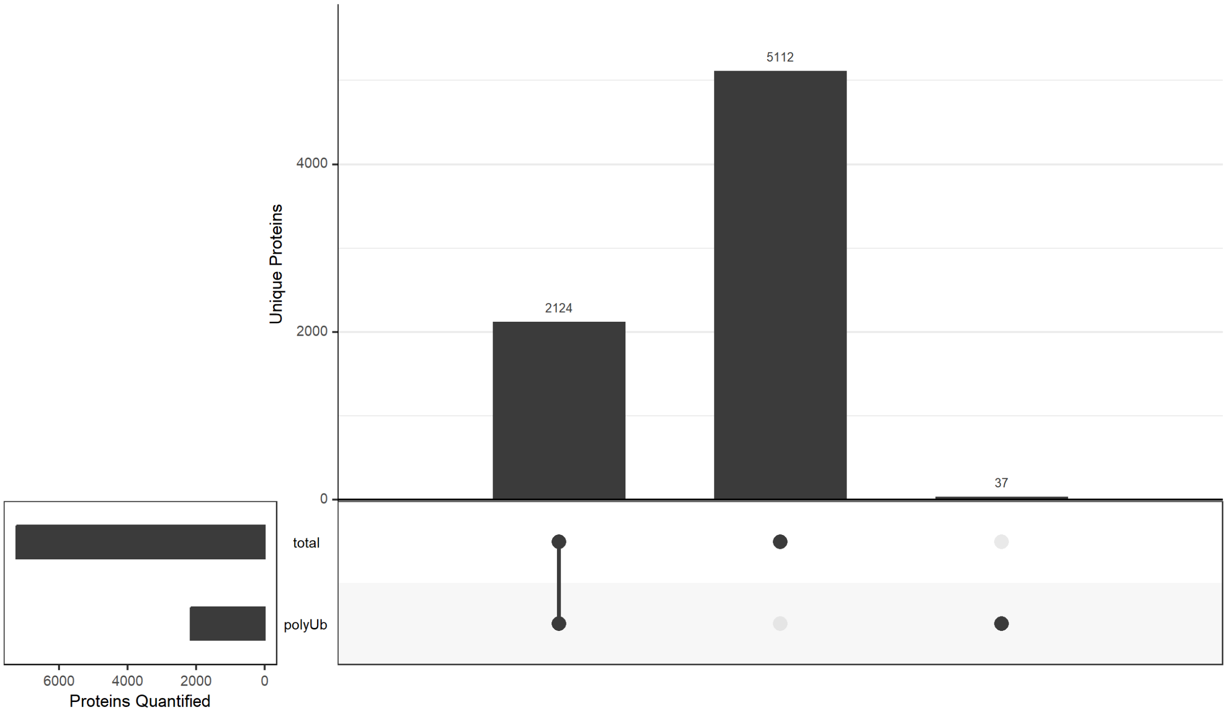

Figure 37: UpSet plot of overlap between proteins quantified in any sample between the two proteomes in the A549 heat-shock dataset.

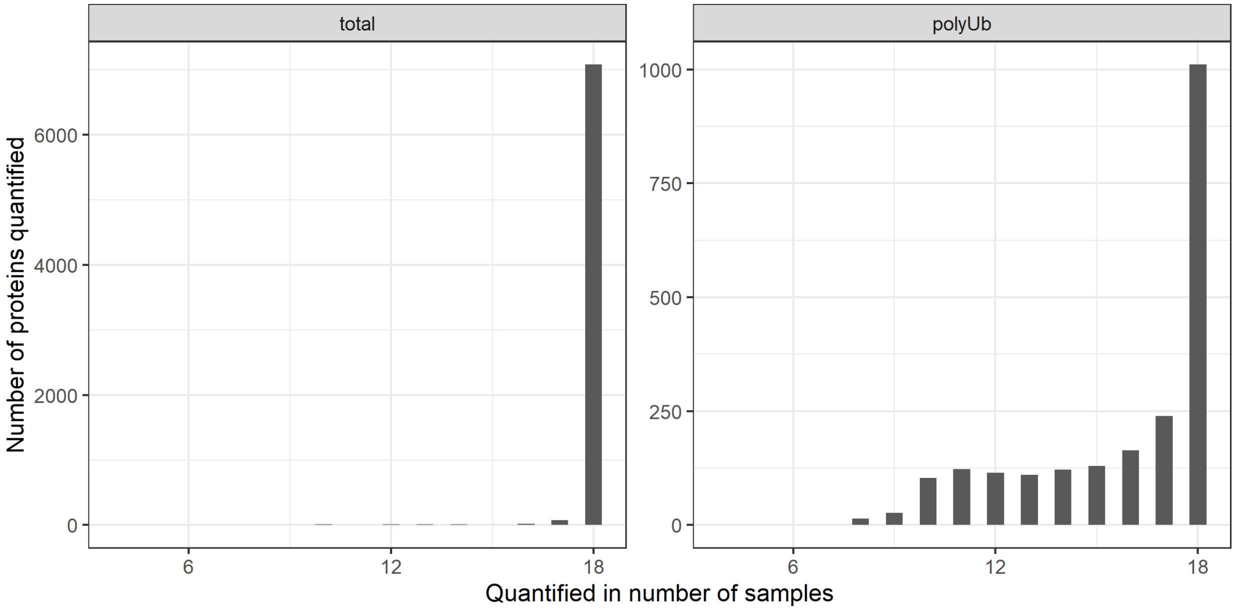

Figure 38: Channel occupancy for each proteome in the A549 heat-shock dataset.

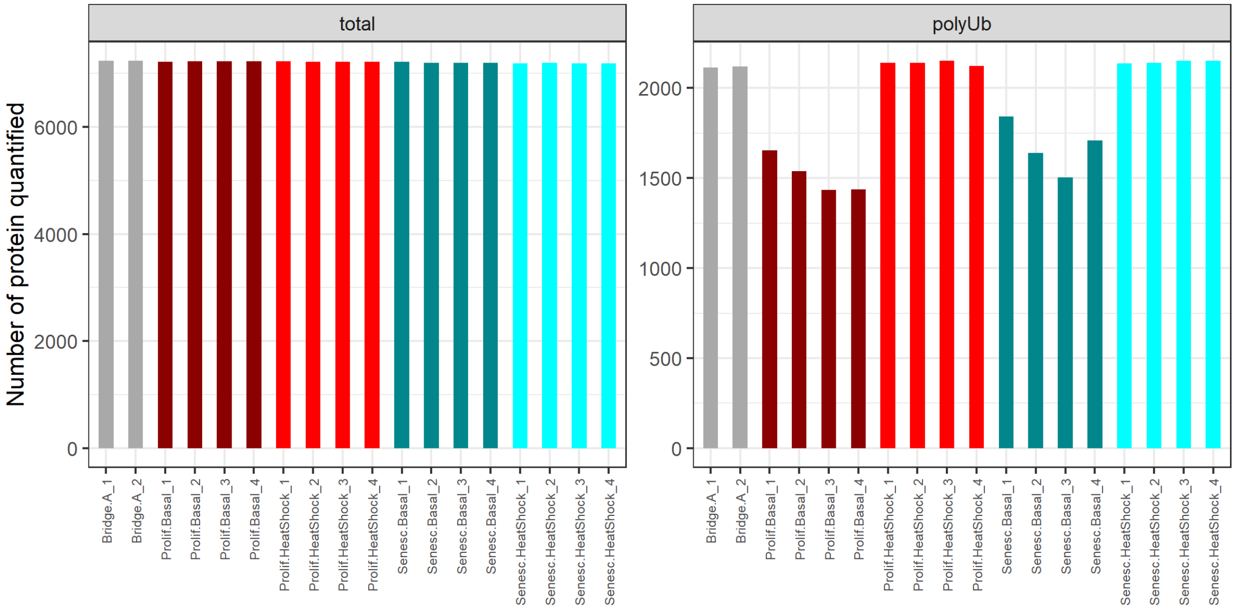

Figure 39: Number of proteins quantified in each sample of the A549 heat-shock dataset.

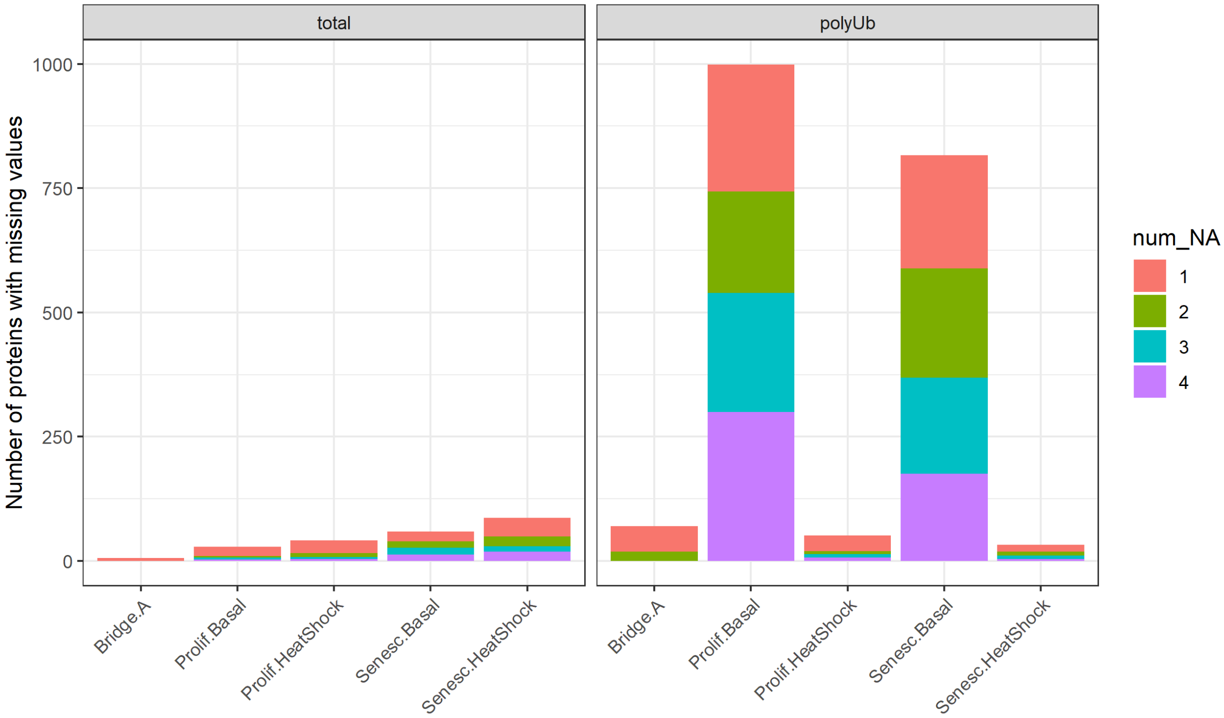

Figure 40: Number of missing values in each state.treatment per proteome for the A549 heat-shock dataset.

#### 3.3 Missing Values

##### 3.3.1 Overlapping proteins

##### 3.3.2 Channel occupancy

These plots make a few points—many being the same points as in the IMR-90 dataset. 1. The 'Bridge' samples have far fewer missing values than the rest of the samples—as expected, given that they are a pool of all the samples. 2. The total proteome has very few missing values, in general. 3. The polyUb proteome has missing values mostly in the basal/vehicle conditions and not in the heat-shock condition, as expected.

For the polyUb proteome, these are Missing Not At Random (MNAR), i.e., proteins that were not quantified in specific conditions (e.g., in basal/vehicle samples only).

##### 3.3.3 Imputing missing values

For the proteins that are missing completely in a TMT18plex (i.e., in all 16 sample channels–not necessarily the two Bridge channels), we will not impute any missing values. Rather, we will remove these proteins.

At this stage, we will also remove the Bridge channels, before proceeding with using DEP for the imputations.

```
dataLong %>%
  dplyr::filter(treatment=="Basal"|
                treatment=="HeatShock") %>%
  group_by(Gene, proteome) %>%
  dplyr::filter(!all(is.na(abundance))) %>%
  ungroup() -> dataLong

read_csv("A549_heat-shock/dataFiltered.csv") %>%
  dplyr::select(Gene, Accession) %>%
  right_join(dataLong) %>%
  dplyr::rename(name = Gene) %>%
  dplyr::rename(ID = Accession) -> dataLong

dataLong %>%
  dplyr::filter(proteome=="total") %>%
  dplyr::select(name, ID, label, abundance) %>%
  pivot_wider(names_from = label, values_from = abundance) -> dataTotal

dataLong %>%
  dplyr::filter(proteome=="polyUb") %>%
  dplyr::select(name, ID, label, abundance) %>%
  pivot_wider(names_from = label, values_from = abundance) -> dataPolyUb

# specify the abundance column names for the SummarizedExperiment
abundances_Total <- grep("Prolif|Senesc", colnames(dataTotal))
abundances_PolyUb <- grep("Prolif|Senesc", colnames(dataPolyUb))

experimental_design %>%
  dplyr::select(label, condition, replicate) %>%
  dplyr::filter(grepl("Basal$", condition)|
                  grepl("HeatShock$", condition)) %>%
  distinct() -> experimental_design
```

```r
SEforDEP_Total <- make_se(dataTotal, abundances_Total, experimental_design)
SEforDEP_PolyUb <- make_se(dataPolyUb, abundances_PolyUb, experimental_design)
```

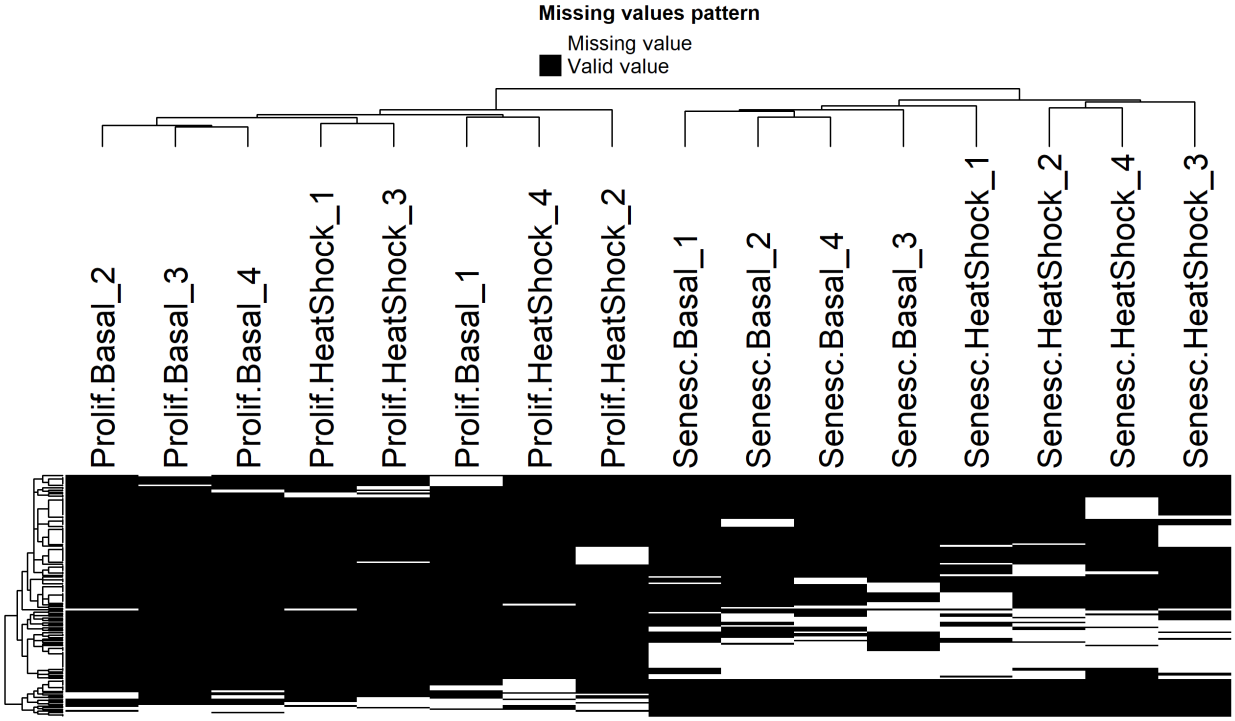

Figure 41: Heatmaps of proteins in the total proteomes, from heat-shocked and basal A549 cells, that have at least one missing value. Samples where abundance values are missing (white) or present (black) are shown.

**3.3.3.1 Deciding on imputation model** For the total proteomes, it is clear that proteins with missing values have lower intensities than the other proteins. This is far less pronounced in the polyUb proteomes.

Although for different reasons, both the total (close-to-detection-limit) and polyUb (missing-not-at-random) data can be imputed by a left-censored imputation method available within the DEP package, as for the IMR-90 heat-shock dataset.

As before, we will set some quality filters so that only proteins quantified in at least 50 % of the samples, and in all four replicates of at least one condition, are kept in the data.

```r
SEforDEP_Total <- filter_proteins(SEforDEP_Total, "fraction", min = 0.5)
SEforDEP_Total <- filter_proteins(SEforDEP_Total, "condition", thr = 0)

SEforDEP_PolyUb <- filter_proteins(SEforDEP_PolyUb, "fraction", min = 0.5)
SEforDEP_PolyUb <- filter_proteins(SEforDEP_PolyUb, "condition", thr = 0)
```

###### 3.3.3.2 Assessing imputation models

```
## [1] 0.3126329
```

```
## [1] 1.10627
```

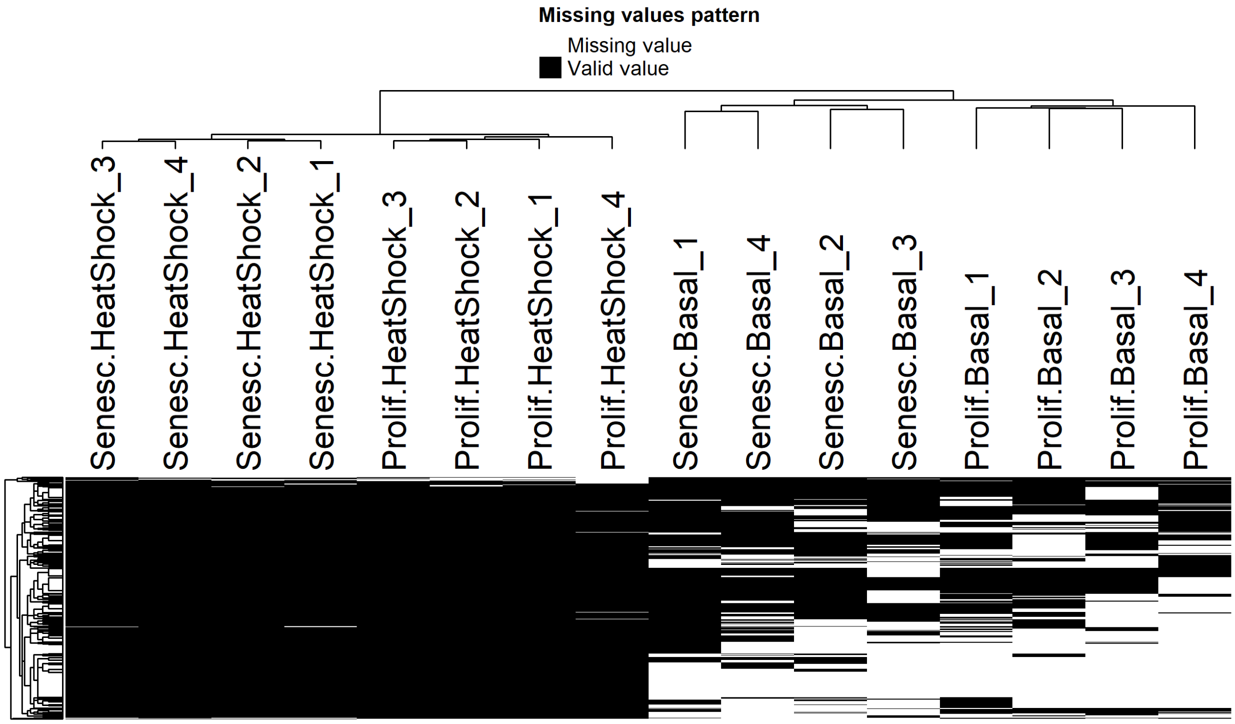

Figure 42: Heatmaps of proteins in the polyUb proteomes, from heat-shocked and basal A549 cells, that have at least one missing value. Samples where abundance values are missing (white) or present (black) are shown.

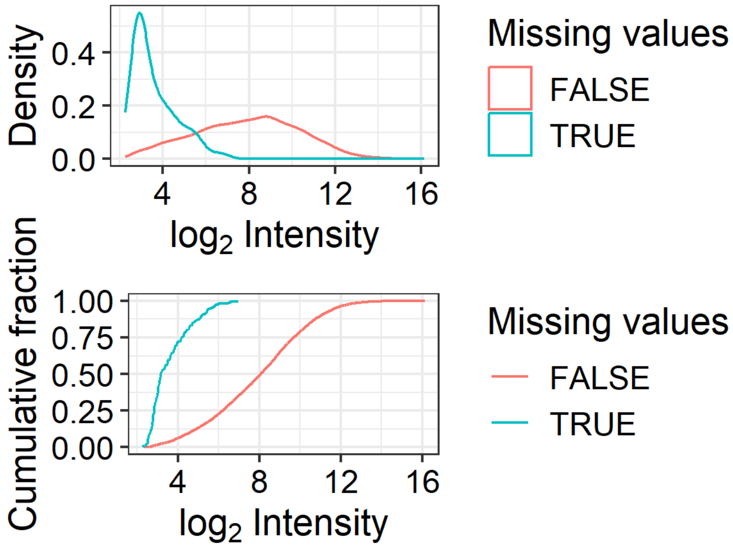

Figure 43: Density (top) and cumulative density (bottom) distributions of log2-transformed intensities, for proteins with (turquoise) or without (red) one or more missing values, in the total proteomes from heat-shocked and basal A549 cells.

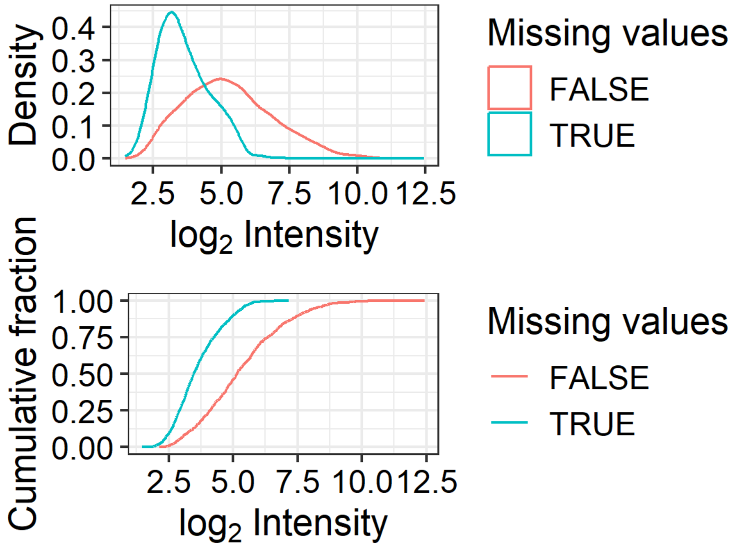

Figure 44: Density (top) and cumulative density (bottom) distributions of log2-transformed intensities, for proteins with (turquoise) or without (red) one or more missing values, in the polyUb proteomes from heat-shocked and basal A549 cells.

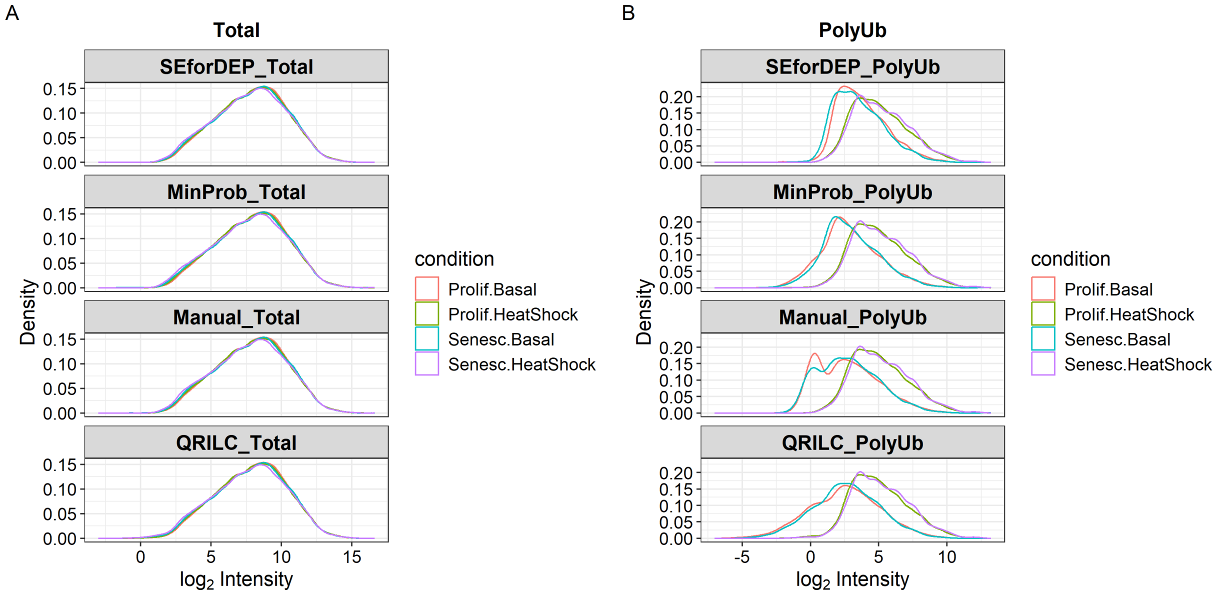

Figure 45: Density distributions of log2-transformed intensities with each imputation model across total (A) and polyUb (B) proteomes for the A549 heat-shock dataset.

The Manual model adds shoulders to the PolyUb proteome, and therefore should not be considered. The QRILC and MinProb models both seem acceptable. As we have used QRILC for the IMR-90 dataset, we will be consistent and apply the same method for this dataset.

#### 3.4 Differential Analysis

##### 3.4.1 Differential Analysis: Total Proteome

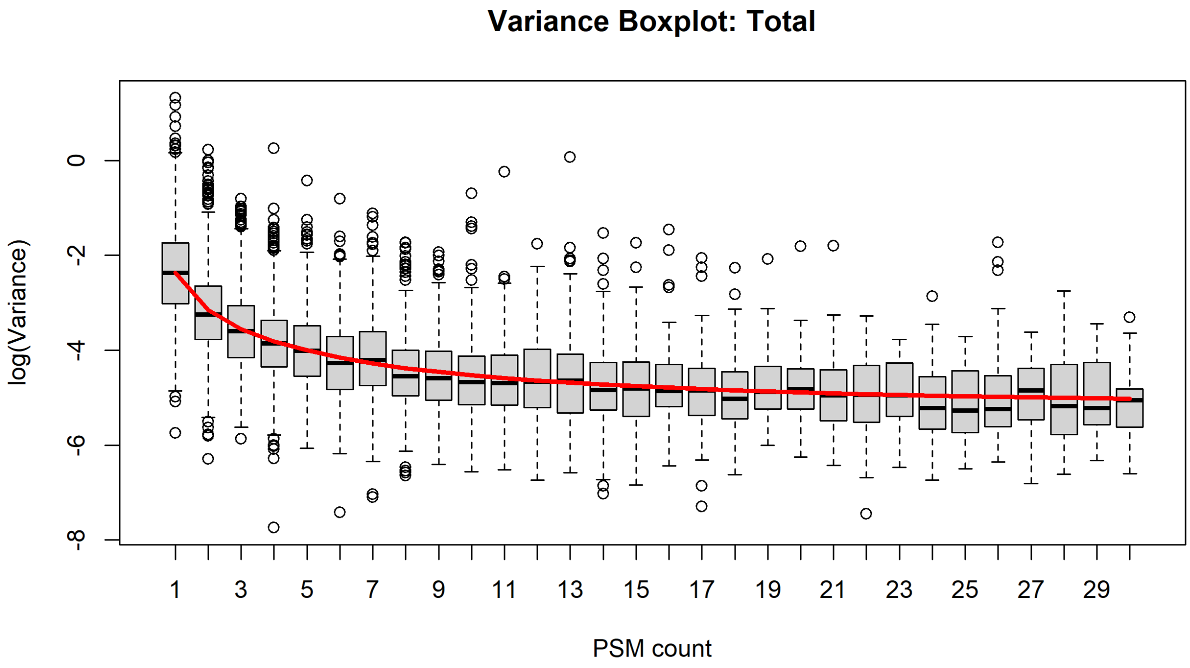

Figure 46: Boxplot showing variance of protein abundances with number of quantified PSMs in the total proteome from basal and heat-shocked samples in the A549 heat-shock experiment.

```
## [1] "Prolif.HeatShock-Prolif.Basal"     "Senesc.HeatShock-Senesc.Basal"
## [3] "Senesc.Basal-Prolif.Basal"         "Senesc.HeatShock-Prolif.HeatShock"
```

##### 3.4.2 Differential Analysis: PolyUb Proteome

##### 3.4.3 Export for downstream analysis

```
data %>%
write_csv("A549_heat-shock/data_DEqMS.csv")
```

### 4 A549 Bortezomib Experiment

Finally, we can perform the same workflow on the bortezomib-treated A549 samples.

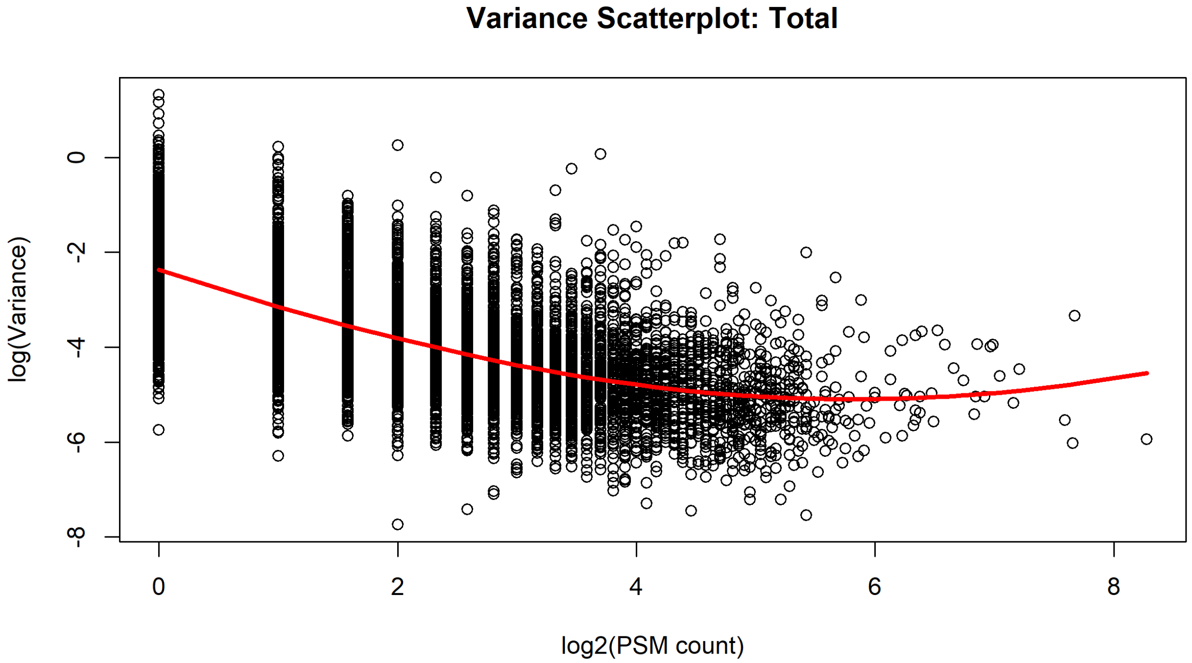

Figure 47: Scatterplot showing variance of protein abundances with number of quantified PSMs in the total proteome from basal and heat-shocked samples in the A549 heat-shock experiment.

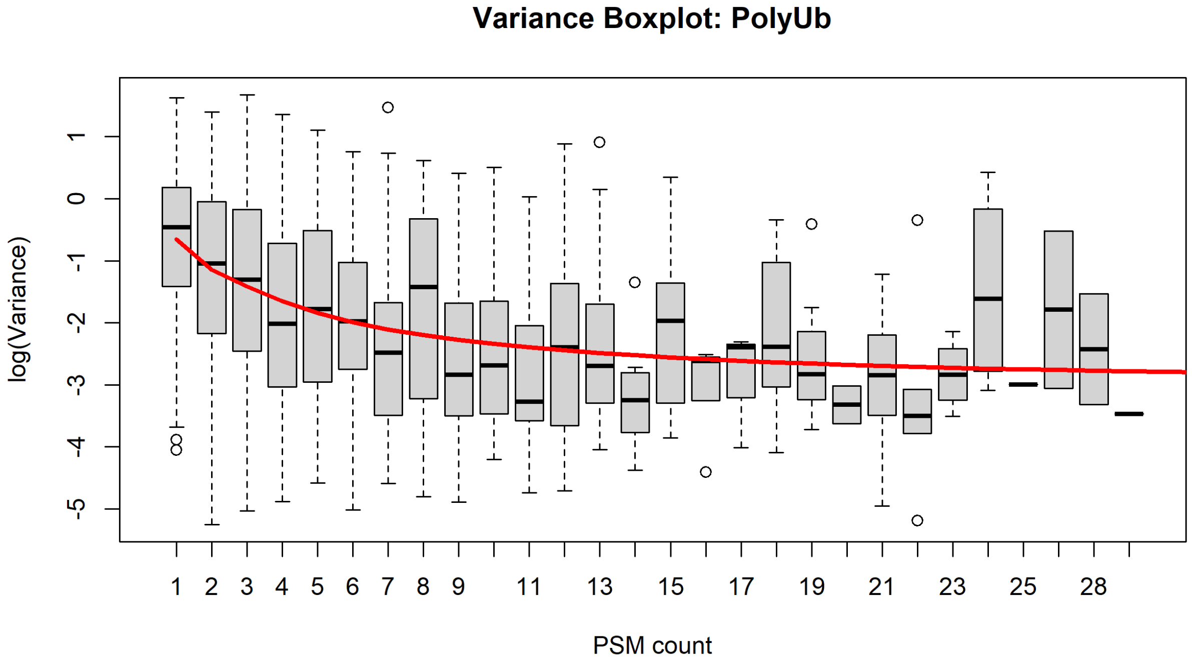

Figure 48: Boxplot showing variance of protein abundances with number of quantified PSMs in the total proteome for basal and heat-shocked samples in the A549 heat-shock experiment.

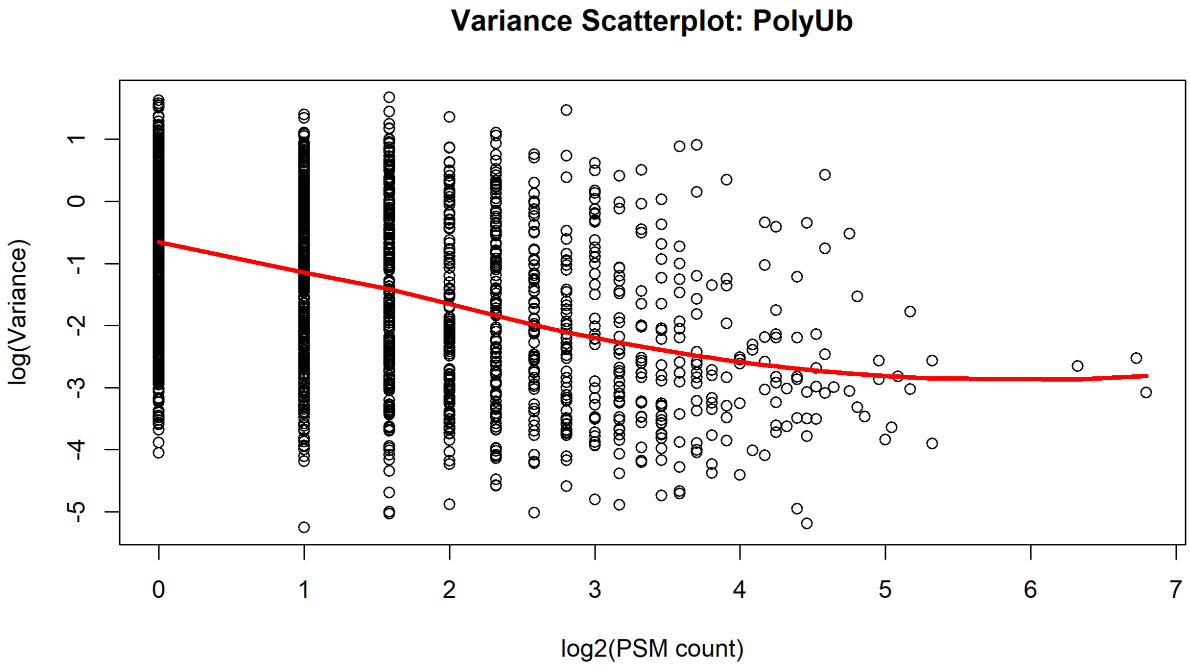

Figure 49: Scatterplot showing variance of protein abundances with number of quantified PSMs in the total proteome for basal and heat-shocked samples in the A549 heat-shock experiment.

#### 4.1 Initial Data Clean-Up and Filtering

```
rm(list = setdiff(ls(), c("get_gene_from_uniprot")))

library(readxl)
read_xlsx("../../A549_Sen44_BTZ/SenBTZ_Tot_TRT_Combined/TMT18_A549_BTZ_Tot_TRT_prot.xlsx"
          ) -> data

dim(data)
```

```
## [1] 9541  178
```

```
## # A tibble: 6 x 9
##   sample    sampleID TMT   conditionID label condition state treatment replicate
##   <chr>     <chr>    <chr> <chr>       <chr> <chr>     <chr> <chr>         <int>
## 1 Abundanc~ Abundan~ 127C  " 5"        Prol~ Prolif.V~ Prol~ Vehicle           1
## 2 Abundanc~ Abundan~ 128C  " 5"        Prol~ Prolif.V~ Prol~ Vehicle           2
## 3 Abundanc~ Abundan~ 129C  " 5"        Prol~ Prolif.V~ Prol~ Vehicle           3
## 4 Abundanc~ Abundan~ 130C  " 5"        Prol~ Prolif.V~ Prol~ Vehicle           4
## 5 Abundanc~ Abundan~ 131C  " 6"        Prol~ Prolif.B~ Prol~ Bortezom~         1
## 6 Abundanc~ Abundan~ 132C  " 6"        Prol~ Prolif.B~ Prol~ Bortezom~         2
```

##### 4.1.1 Clean-up of Gene and Accession identifiers

###### 4.1.1.1 Filling in missing Gene Symbols

```
## # A tibble: 2 x 2
## # Groups:   is.na(Gene) [2]
##   `is.na(Gene)`     n
##   <lgl>         <int>
## 1 FALSE          9473
## 2 TRUE             68
```

After using our function 'get_gene_from_uniprot'…

```
## # A tibble: 2 x 2
## # Groups:   is.na(Gene) [2]
##   `is.na(Gene)`     n
##   <lgl>         <int>
## 1 FALSE          9526
## 2 TRUE             15
```

…there are 15 remaining missing Gene names to be annotated manually. Performed as before (identical code; not shown here).

```
## # A tibble: 15 x 2
##    Accession       Description
##    <chr>           <chr>
##  1 Cont_X00000     Halo-TR-TUBE protein OS=Escherichia coli OX=0000 GN=HaloTEVT~
##  2 Cont_P00761     Trypsin OS=Sus scrofa OX=9823 PE=1 SV=1
##  3 Cont_P02081     Hemoglobin fetal subunit beta OS=Bos taurus OX=9913 PE=1 SV=1
##  4 Q6ZSR9          Uncharacterized protein FLJ45252 OS=Homo sapiens OX=9606 PE=~
##  5 Cont_A0A3Q1M3L6 Uncharacterized protein OS=Bos taurus OX=9913 PE=1 SV=1
##  6 Cont_P50448     Factor XIIa inhibitor OS=Bos taurus OX=9913 PE=1 SV=1
##  7 Cont_G5E513     Uncharacterized protein OS=Bos taurus OX=9913 PE=1 SV=2
##  8 Cont_E1BCW0     HGF activator OS=Bos taurus OX=9913 GN=HGFAC PE=4 SV=3
##  9 Q9UF83          Uncharacterized protein DKFZp434B061 OS=Homo sapiens OX=9606~
## 10 Q08AF8          Putative golgin subfamily A member 8F/8G OS=Homo sapiens OX=~
## 11 Cont_Q29437     Primary amine oxidase, liver isozyme OS=Bos taurus OX=9913 P~
## 12 Q9HAA7          Putative uncharacterized protein FLJ11871 OS=Homo sapiens OX~
## 13 Cont_Q2KIT0     Protein HP-20 homolog OS=Bos taurus OX=9913 PE=2 SV=1
## 14 Cont_O76011     Keratin, type I cuticular Ha4 OS=Homo sapiens OX=9606 GN=KRT~
## 15 A8MVM7          Putative uncharacterized protein ENSP00000382790 OS=Homo sap~
```

As before, most of the missing Gene names are contaminants, which we will handle by copying over the Accession field.

```
## # A tibble: 5 x 2
##   Accession Description
##   <chr>     <chr>
## 1 Q6ZSR9    Uncharacterized protein FLJ45252 OS=Homo sapiens OX=9606 PE=2 SV=2
## 2 Q9UF83    Uncharacterized protein DKFZp434B061 OS=Homo sapiens OX=9606 PE=2 S~
## 3 Q08AF8    Putative golgin subfamily A member 8F/8G OS=Homo sapiens OX=9606 GN~
## 4 Q9HAA7    Putative uncharacterized protein FLJ11871 OS=Homo sapiens OX=9606 P~
## 5 A8MVM7    Putative uncharacterized protein ENSP00000382790 OS=Homo sapiens OX~
```

For the remaining 5 missing Genes, we will manually fill these in manually.

```
data %>%
  mutate(Gene = replace(Gene, Accession == "Q9HAA7", "FLJ11871")) %>%
  mutate(Gene = replace(Gene, Accession == "Q9UF83", "DKFZp434B061")) %>%
  mutate(Gene = replace(Gene, Accession == "Q08AF8", "GOLGA8F")) %>%
  mutate(Gene = replace(Gene, Accession == "Q6ZSR9", "FLJ45252")) %>%
  mutate(Gene = replace(Gene, Accession == "A8MVM7", "ENSP00000382790")) -> data

data %>%
  dplyr::filter(is.na(Gene)) %>%
  dplyr::count()
```

```
## # A tibble: 1 x 1
##       n
##   <int>
## 1     0
```

We will also rename the ubiquitin Gene again.

```
# Check for duplicated Accessions or Genes
data %>%
  group_by(Accession) %>%
  filter(n()>1) %>%
  dplyr::count()
```

###### 4.1.1.2 Making duplicated Genes unique

```
## # A tibble: 0 x 2
## # Groups:   Accession [0]
## # i 2 variables: Accession <chr>, n <int>
```

```
data %>%
  group_by(Gene) %>%
  filter(n()>1) %>%
  dplyr::count()
```

```
## # A tibble: 0 x 2
## # Groups:   Gene [0]
## # i 2 variables: Gene <chr>, n <int>
```

Once again, there are no duplicated Accessions or Genes.

##### 4.1.2 Adding proteome annotations

##### 4.1.3 Removing contaminants

```
## # A tibble: 2 x 2
## # Groups:   Contaminant [2]
##   Contaminant     n
```

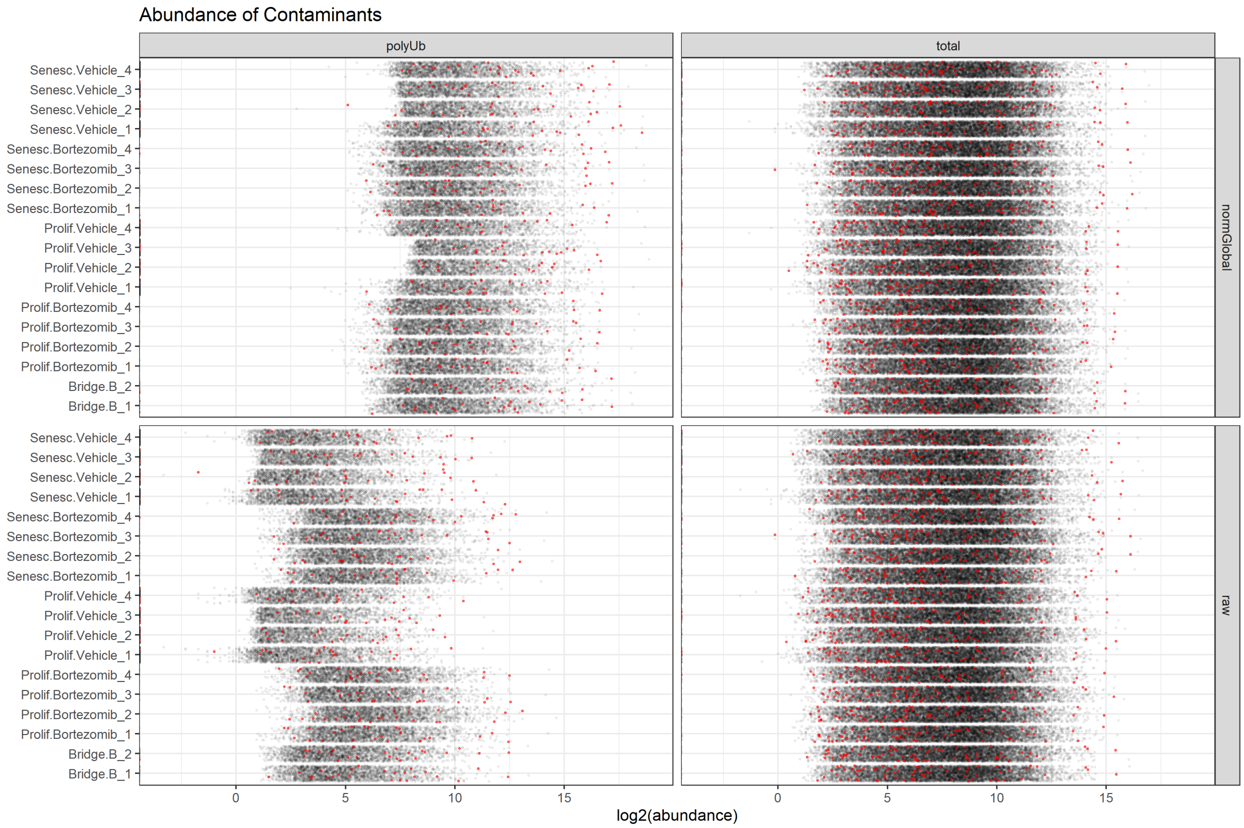

Figure 50: The raw abundance distribution of known contaminants in the A549 bortezomib dataset.

```
##   <lgl>       <int>
## 1 FALSE        9399
## 2 TRUE          142
```

We removed these contaminants.

##### 4.1.4 Dealing with single PSM/peptide identifications

```
data %>%
  dplyr::select(Gene, Accession, `# Peptides`, `# PSMs`) %>%
  right_join(dataNoConts) -> dataNoConts

# Plot number of proteins identified by a single PSM
dataNoConts %>%
  dplyr::rename(numPSMs = `# PSMs`) %>%
  group_by(numPSMs > 1) %>%
  dplyr::count() %>%
  ungroup() %>%
  dplyr::filter(`numPSMs > 1` == FALSE) %>%
  dplyr::select(`n`) -> singletonNumber

dataNoConts %>%
    ggplot(aes(x = `# PSMs`)) +
    geom_bar(fill = "grey", colour = "black", size = 0.2) +
    geom_bar(data = ~filter(.x, `# PSMs` == 1), fill = "red", colour = "black") +
    coord_cartesian(xlim = c(0, 200)) +
  annotate("text", label = paste0(
    singletonNumber, " proteins identified by a single PSM"),
    x = 80, y = 380, color = "red", size = 4) +
    xlab("number of unique PSMs") +
  ylab("count(Proteins)") -> plot_numPSMs

# Count number of proteins identified by a single peptide
dataNoConts %>%
  dplyr::rename(numPeptides = `# Peptides`) %>%
  group_by(numPeptides > 1) %>%
  dplyr::count() %>%
  ungroup() %>%
  dplyr::filter(`numPeptides > 1` == FALSE) %>%
  dplyr::select(`n`) -> singletonNumber

dataNoConts %>%
    ggplot(aes(x = `# Peptides`)) +
    geom_bar(fill = "grey", colour = "black", size = 0.2) +
    geom_bar(data = ~filter(.x, `# Peptides` == 1), fill = "red", colour = "black") +
    coord_cartesian(xlim = c(0, 200)) +
  annotate("text", label = paste0(
    singletonNumber, " proteins identified by a single peptide"),
    x = 80, y = 600, color = "red", size = 4) +
    xlab("number of unique Peptides") +
  ylab("count(Proteins)") -> plot_numPeptides
```

```
plot_numPSMs + plot_numPeptides +
  plot_layout(nrow = 2) + plot_annotation(tag_levels = 'A')
```

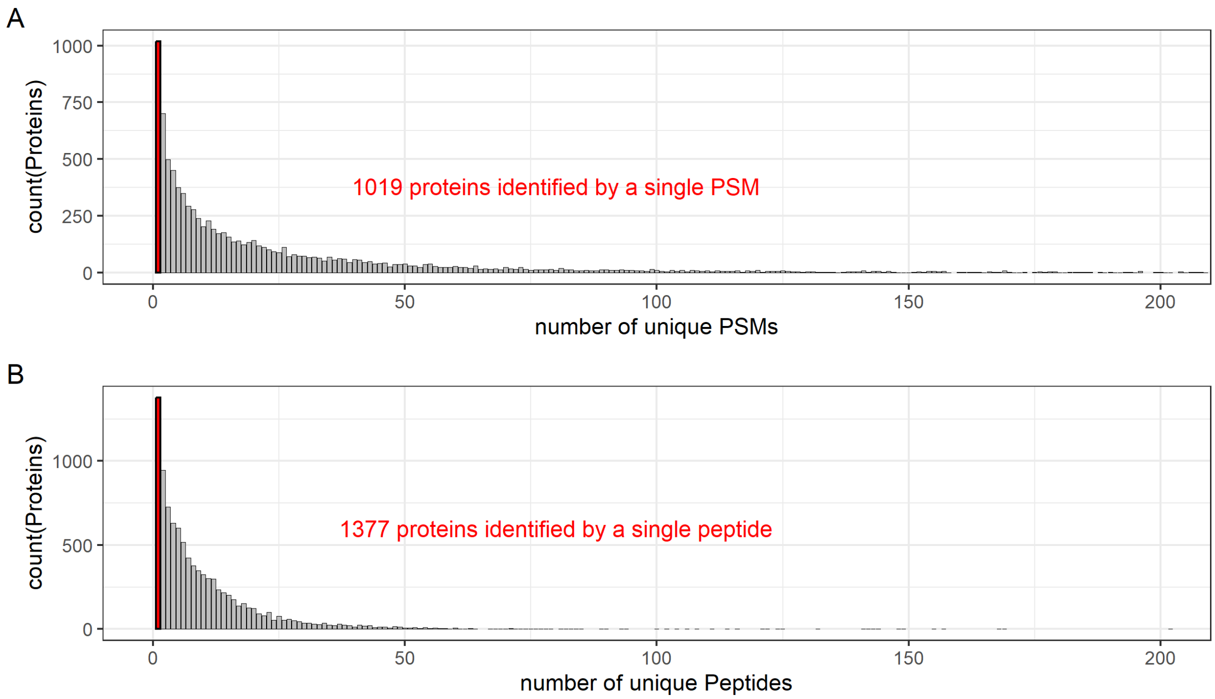

Figure 51: Histogram showing distribution of proteins identified by the number of unique PSMs (A) or peptides (B) in the A549 bortezomib experiment.

```
dataNoContsNoSinglePSMs %>%
  dplyr::select(-`# PSMs`, -`# Peptides`) %>%

  write_csv("A549_bortezomib/dataFiltered.csv",
            na = "NA", append = FALSE, col_names = TRUE, escape = "double")

dim(dataNoContsNoSinglePSMs)
```

```
## [1] 8380   76
```

#### 4.2 Data Normalisation

##### 4.2.1 Assessing global normalisation

Here, we will have a slightly different colour palette, so that we can clearly distinguish between the different stress used here and in the heat-shock experiments.

```
Btz_sample_colours <- c('lightgrey', 'orangered3','orange', 'aquamarine3', 'aquamarine')

# a function 'col2hex' to find the HEX codes for these colours
# (e.g., for replicating them in Illustrator)
```

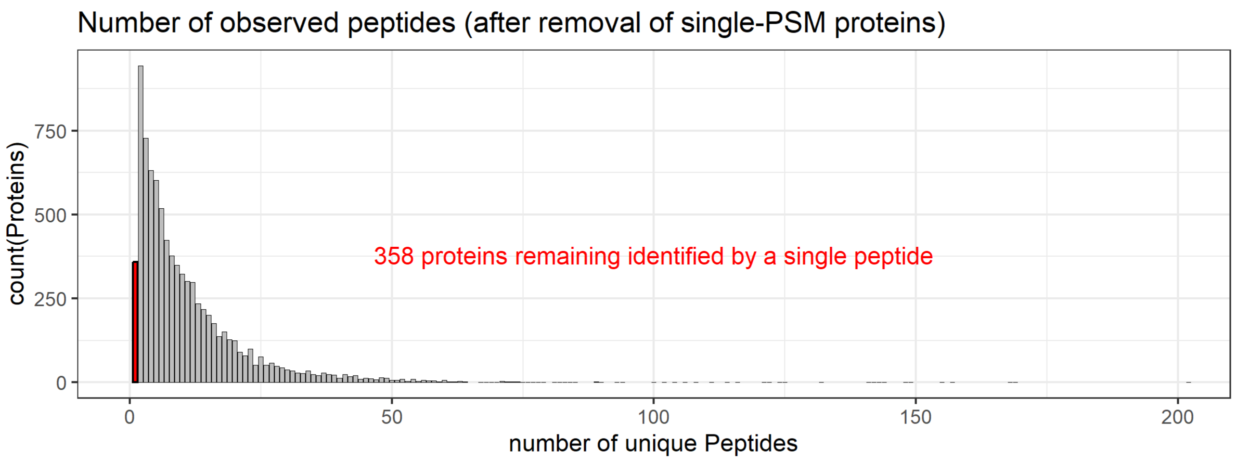

Figure 52: Histogram showing distribution of proteins identified in the A549 bortezomib experiment by the number of unique peptides, after the 1,019 proteins identified by single PSMs had been removed.

```
col2hex <- function(x, alpha = FALSE) {
  args <- as.data.frame(t(col2rgb(x, alpha = alpha)))
  args <- c(args, list(names = x, maxColorValue = 255))
  do.call(rgb, args)
}

col2hex(Btz_sample_colours)
```

```
##    lightgrey  orangered3      orange aquamarine3  aquamarine
##    "#D3D3D3"   "#CD3700"   "#FFA500"   "#66CDAA"   "#7FFFD4"
```

Plot the distributions of the data.

Use the same low-CV normalisation strategy.

###### 4.2.2 Calculate low-CV genes

```
## # A tibble: 2 x 2
##   proteome nPromteins
```

###### 4.2.3 Calculate size-factors based on low-CV genes

###### 4.2.4 Normalising data based on low-CV-calculated size-factors

```
# save all abundance values as "dataNormalised_All.csv"
dataNormalised_All %>%
  write_csv("A549_bortezomib/dataNormalised_All.csv",
            na = "NA", append = FALSE, col_names = TRUE, escape = "double")
```

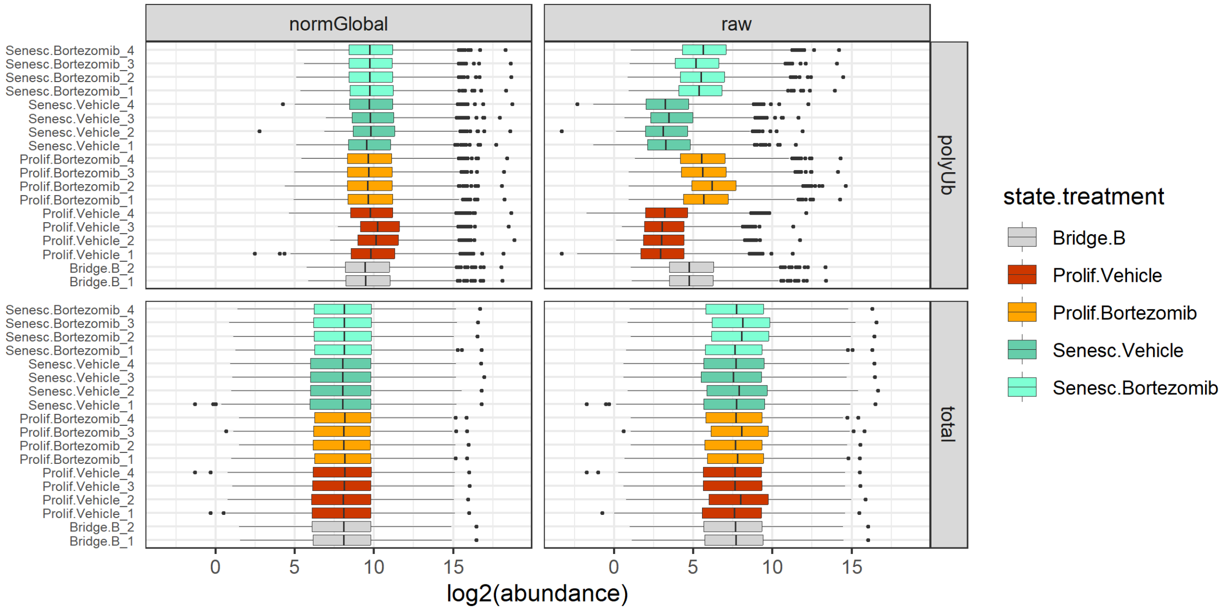

Figure 53: Raw and normalised log2-tranformed abundance distributions of all proteins identified in each sample of the A549 bortezomib experiment.

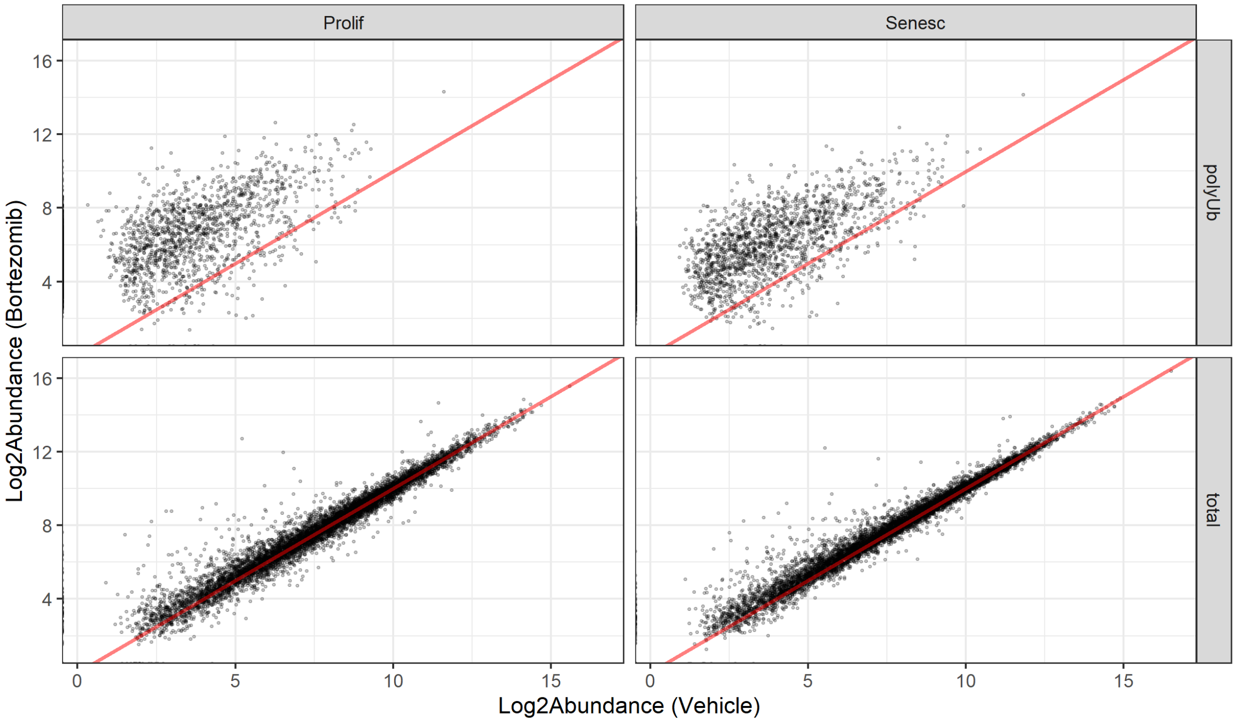

Figure 54: Scatter-plots of mean raw protein abundances for each state in the A549 bortezomib experiment. Each point represents the mean raw abundance of an individual protein in the vehicle- vs. bortezomib-treated samples.

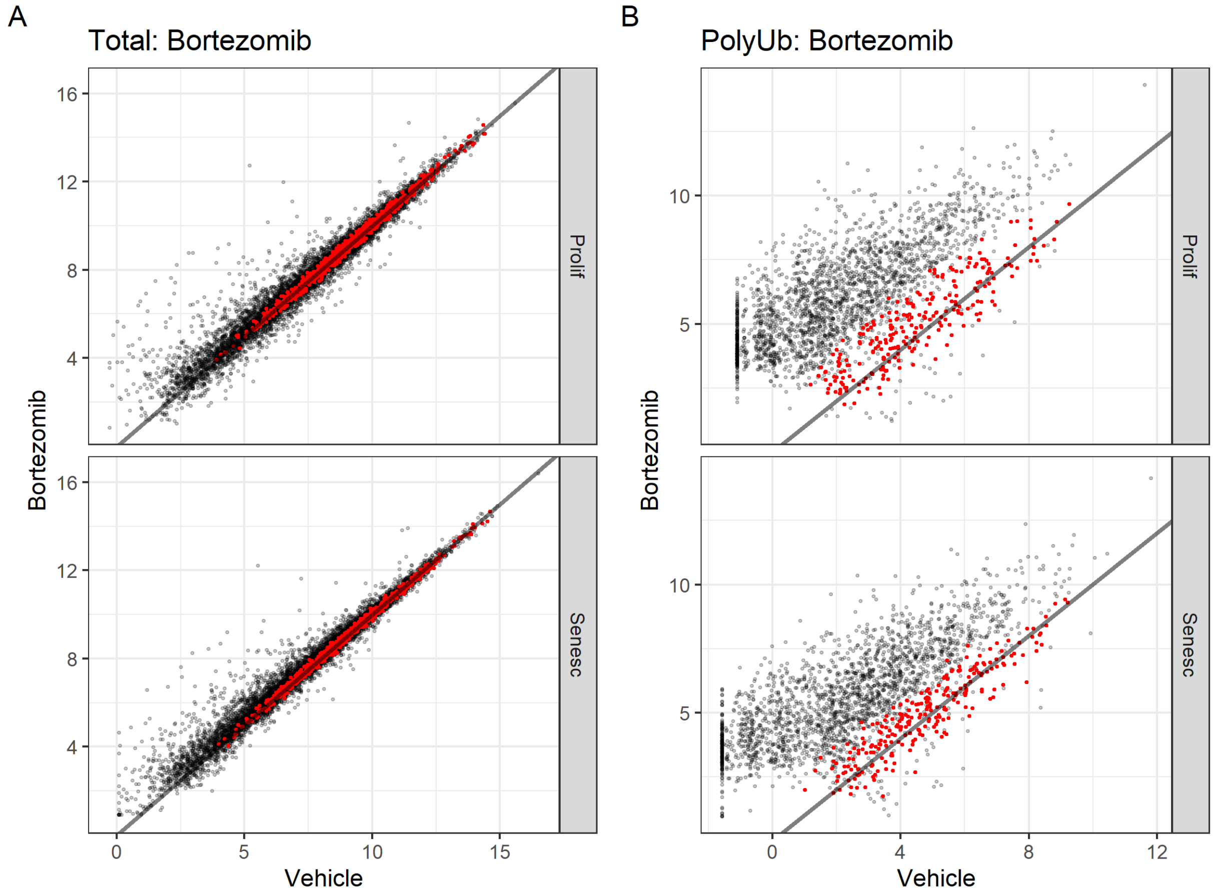

Figure 55: Lowest 10% CV proteins mapped onto scatter-plots of mean raw protein abundances for the A549 bortezomib dataset. Each point represents the mean raw abundance of an individual protein in the vehicle vs. bortezomib-treated samples. The proteins with the 10% lowest CVs for each proteome are plotted in red

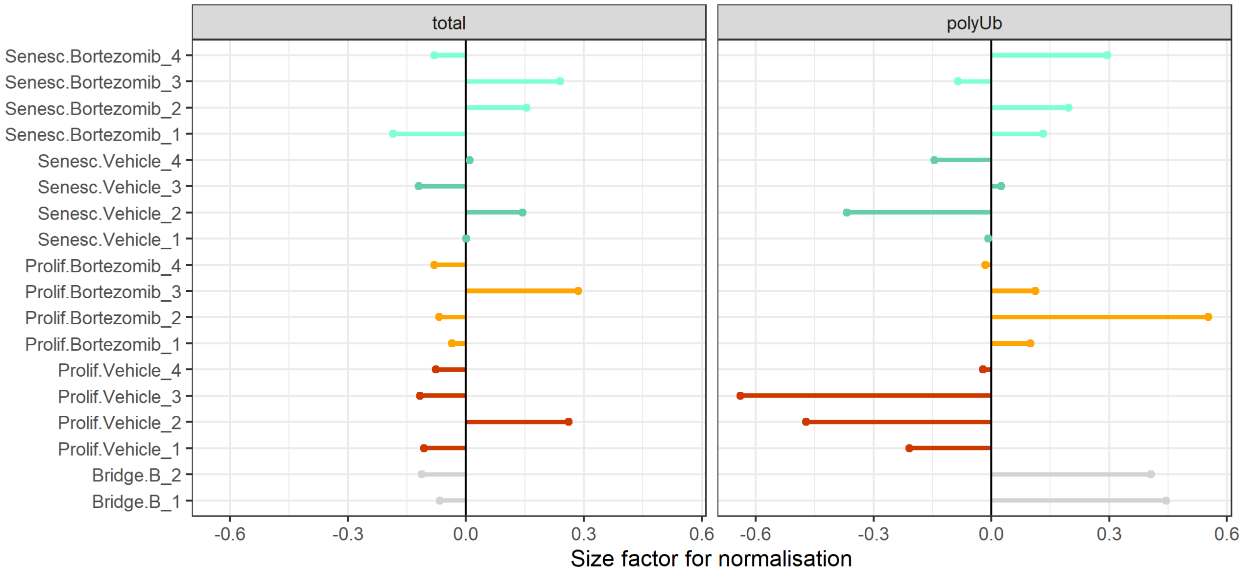

Figure 56: Size factors required for normalisation of each sample in the A549 bortezomib dataset, based on the mean 10% low-CV protein values.

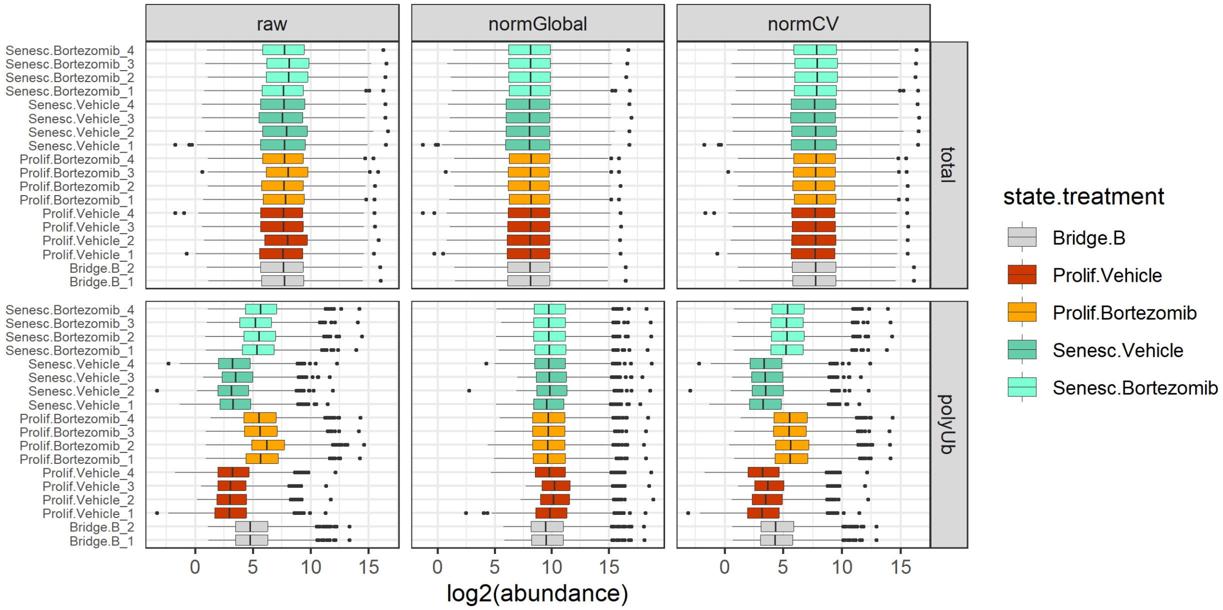

Figure 57: Raw and normalised protein abundance distributions in each sample from the A549 bortezomib experiment.

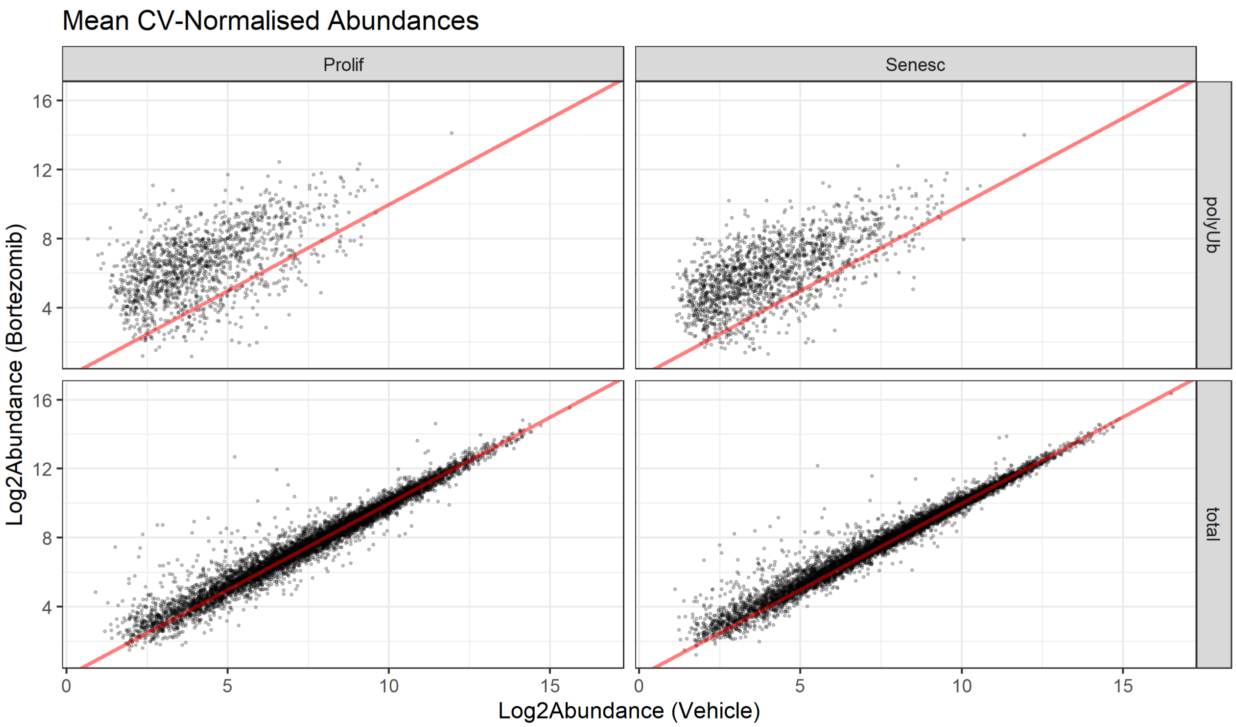

Figure 58: Scatter-plots of mean CV-normalised protein abundances for each state in the A549 bortezomib experiment. Each point represents the mean CV-normalised abundance of an individual protein in the vehicle vs. bortezomib-treated samples.

```
# save only selected normalised abundances as "dataNormalised_Selected.csv"
dataNormalised_All %>%
  dplyr::select(-starts_with("raw")) %>%
  dplyr::select(-starts_with(c("normCV_total"))) %>%
  dplyr::select(-starts_with(c("normGlobal_polyUb"))) %>%
  rename_with(~ gsub("normCV_", "", .x, fixed = TRUE)) %>%
  rename_with(~ gsub("normGlobal_", "", .x, fixed = TRUE)) -> dataNormalised_selected

dataNormalised_selected %>%
  write_csv("A549_bortezomib/dataNormalised_Selected.csv",
            na = "NA", append = FALSE, col_names = TRUE, escape = "double")
```

#### 4.3 Missing Values

##### 4.3.1 Overlapping proteins

##### 4.3.2 Channel occupancy

```
dataLong %>%
  group_by(Gene, proteome) %>%
  dplyr::filter(!all(is.na(abundance))) %>%
```

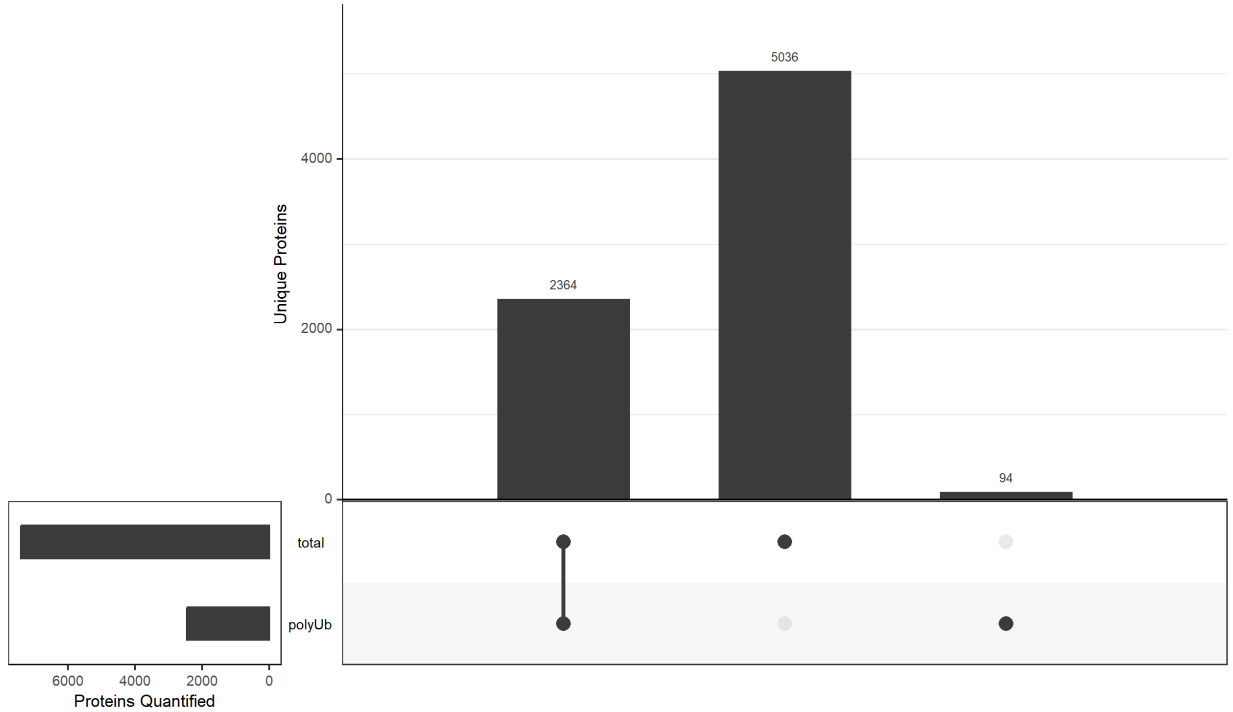

Figure 59: UpSet plot of overlap between proteins quantified in any sample between the two proteomes for the A549 bortezomib dataset.

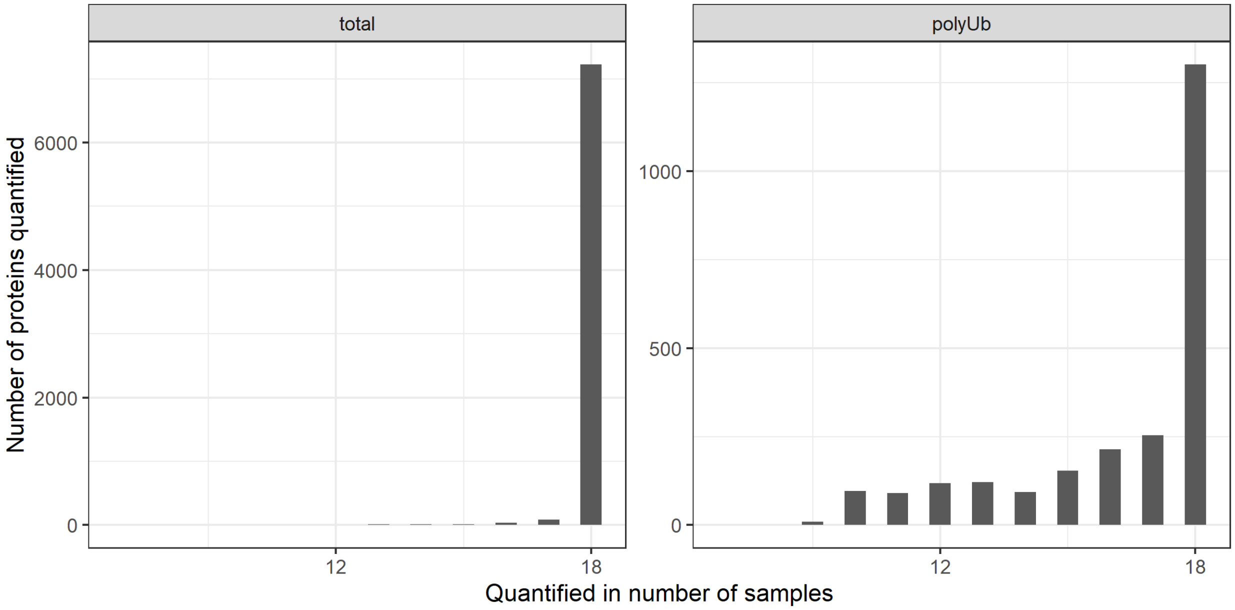

Figure 60: Channel occupancy for each proteome in the A549 bortezomib experiment.

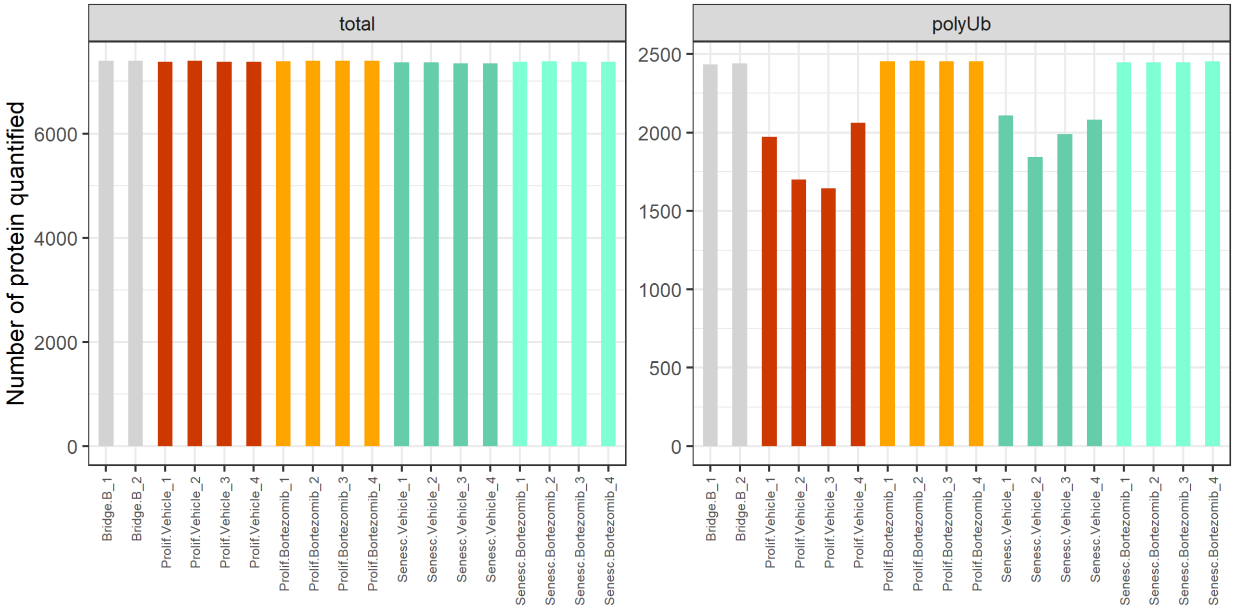

Figure 61: Number of proteins quantified in each sample of the A549 bortezomib dataset.

```
  dplyr::group_by(Gene, proteome, state.treatment) %>%
  dplyr::summarise(num_NA = sum(is.na(abundance))) %>%
  ungroup() %>%
  dplyr::filter(num_NA > 0) %>%
  dplyr::mutate(num_NA = as.factor(num_NA)) %>%

  mutate(state.treatment = factor(
    state.treatment, levels = c("Bridge.B",
                                "Prolif.Vehicle", "Prolif.Bortezomib",
                                "Senesc.Vehicle", "Senesc.Bortezomib"))) %>%

  ggplot(aes(x = state.treatment, fill = num_NA)) +
  geom_bar() +
  ylab("Number of proteins with missing values") +
  theme(axis.text.x = element_text(angle = 45, hjust = 1),
        axis.title.x = element_blank()) +
  facet_grid(~factor(proteome, levels = c("total", "polyUb")))
```

As these plots are very similar to the heat-shock data (in terms of missing-value distributions), we can proceed with the same strategy.

##### 4.3.3 Imputing missing values

**4.3.3.1 Deciding on imputation model** Again, very similar distributions as for the heat-shock data. Let's proceed as before.

```
SEforDEP_Total <- filter_proteins(SEforDEP_Total, "fraction", min = 0.5)
SEforDEP_Total <- filter_proteins(SEforDEP_Total, "condition", thr = 0)
```

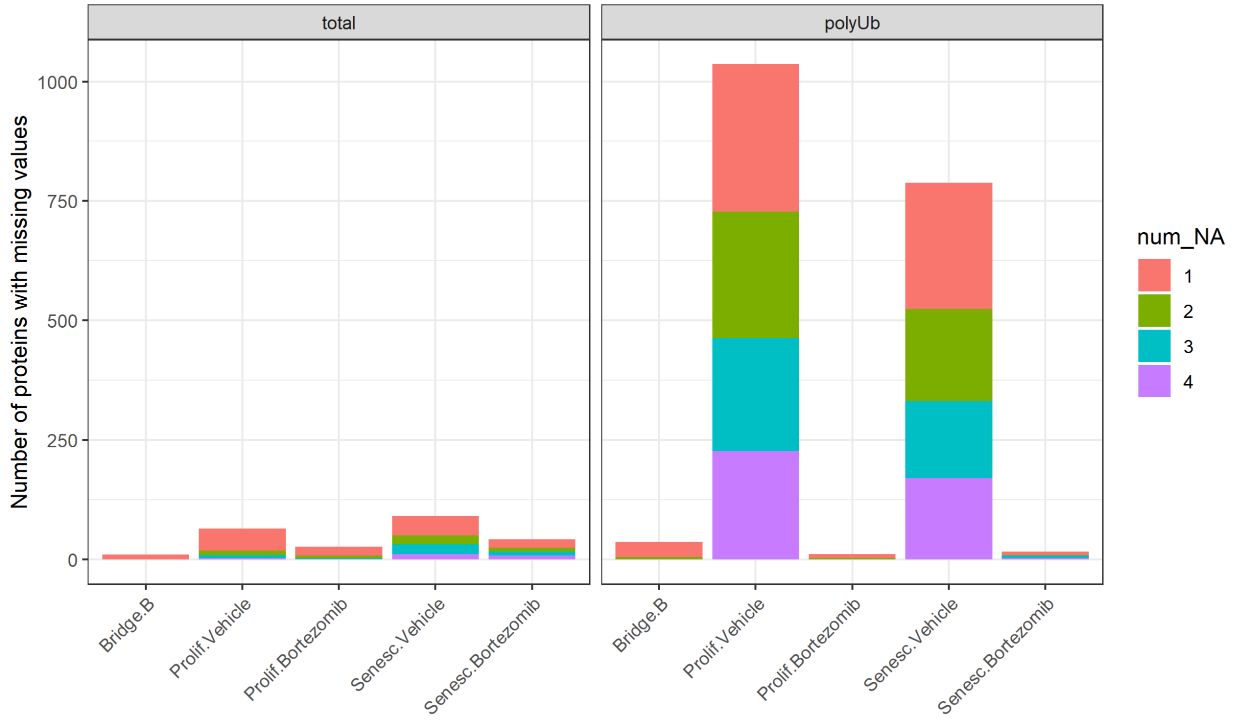

Figure 62: Number of missing values in each state.treatment per proteome of the A549 bortezomib dataset.

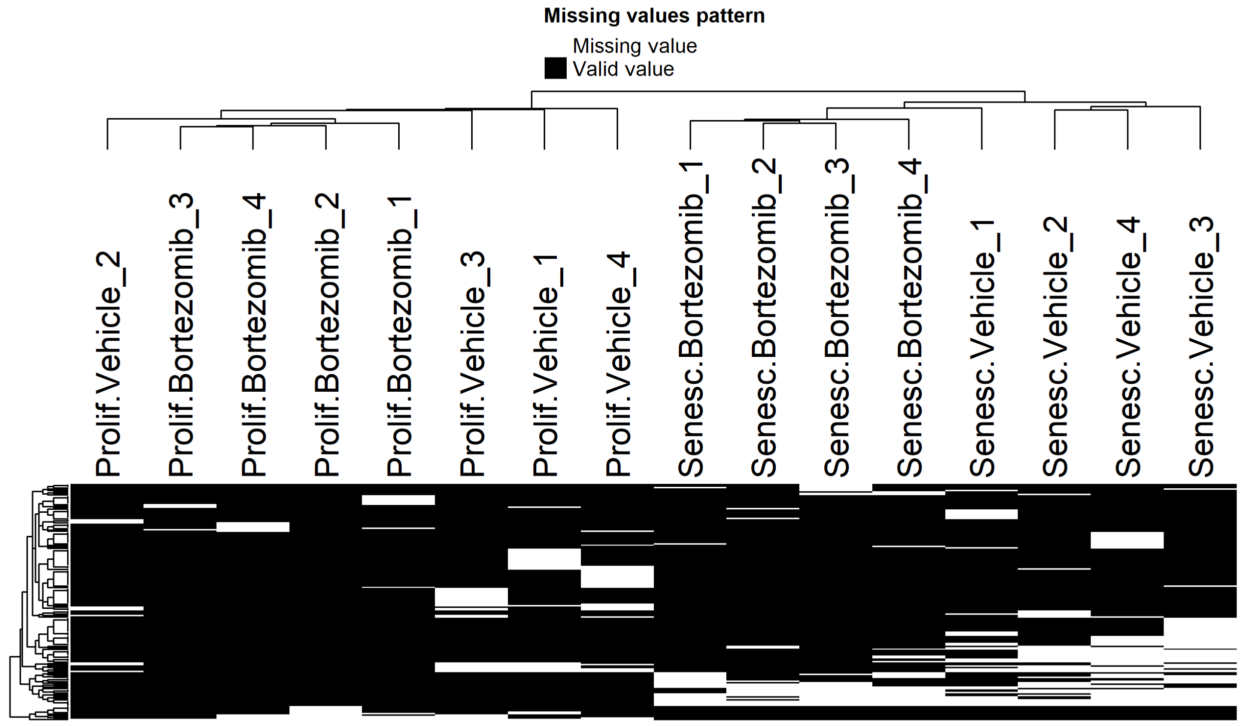

Figure 63: Heatmaps of proteins in the total proteomes, from vehicle- and bortezomib-treated A549 cells, that have at least one missing value. Samples where abundance values are missing (white) or present (black) are shown.

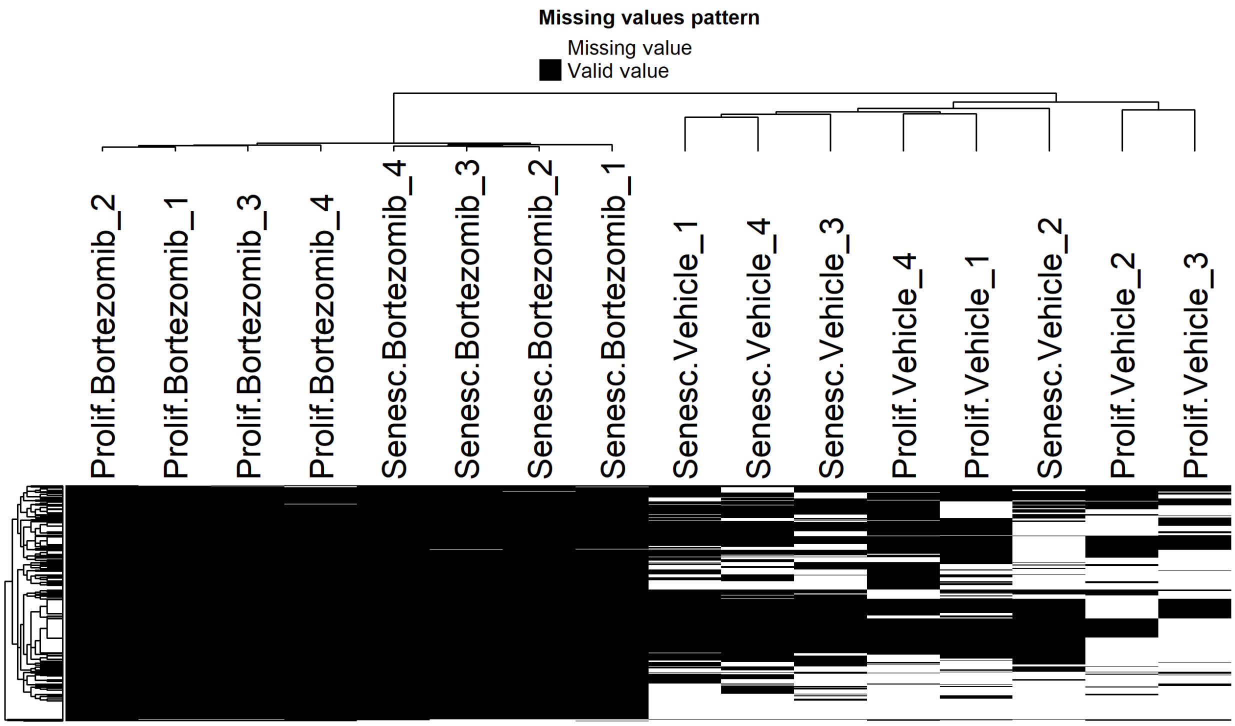

Figure 64: Heatmaps of proteins in the polyUb proteomes, from vehicle- and bortezomib-treated A549 cells, that have at least one missing value. Samples where abundance values are missing (white) or present (black) are shown.

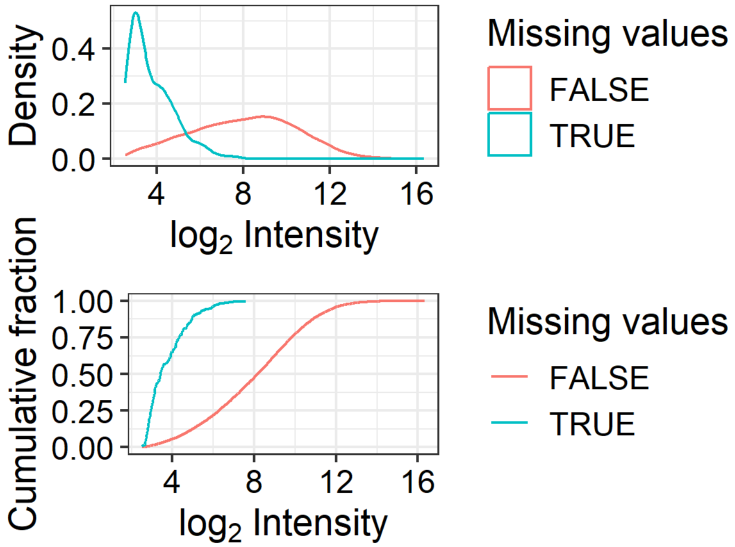

Figure 65: Density (top) and cumulative density (bottom) distributions of log2-transformed intensities, for proteins with (turquoise) or without (red) one or more missing values, in the total proteomes from vehicle- and bortezomib-treated A549 cells.

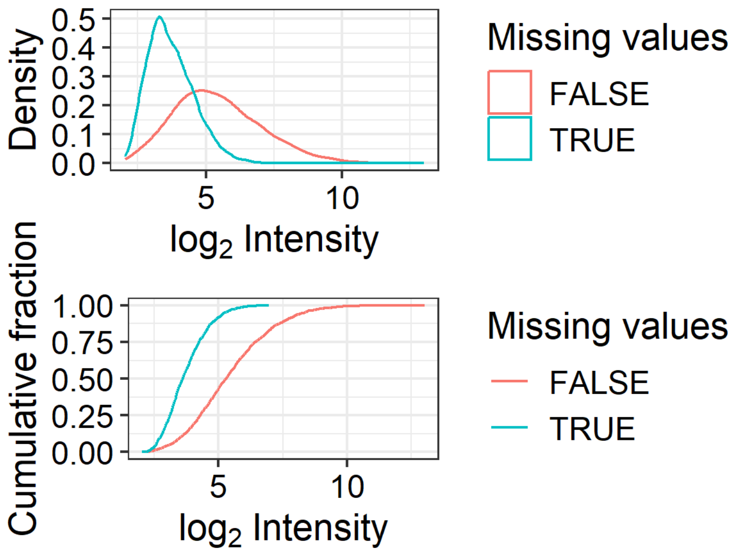

Figure 66: Density (top) and cumulative density (bottom) distributions of log2-transformed intensities, for proteins with (turquoise) or without (red) one or more missing values, in the polyUb proteomes from vehicle- and bortezomib-treated A549 cells.

```
SEforDEP_PolyUb <- filter_proteins(SEforDEP_PolyUb, "fraction", min = 0.5)
SEforDEP_PolyUb <- filter_proteins(SEforDEP_PolyUb, "condition", thr = 0)
```

###### 4.3.3.2 Assessing imputation models

```
## [1] 0.3286718
```

```
## [1] 1.300777
```

As before, we will keep the QRILC-imputed data.

#### 4.4 Differential Analysis

##### 4.4.1 Differential Analysis: Total Proteome

```
## [1] "Prolif.Bortezomib-Prolif.Vehicle"    "Senesc.Bortezomib-Senesc.Vehicle"
## [3] "Senesc.Vehicle-Prolif.Vehicle"       "Senesc.Bortezomib-Prolif.Bortezomib"
```

##### 4.4.2 Differential Analysis: PolyUb Proteome

##### 4.4.3 Export for downstream analysis

```
data %>%
write_csv("A549_bortezomib/data_DEqMS.csv")
```

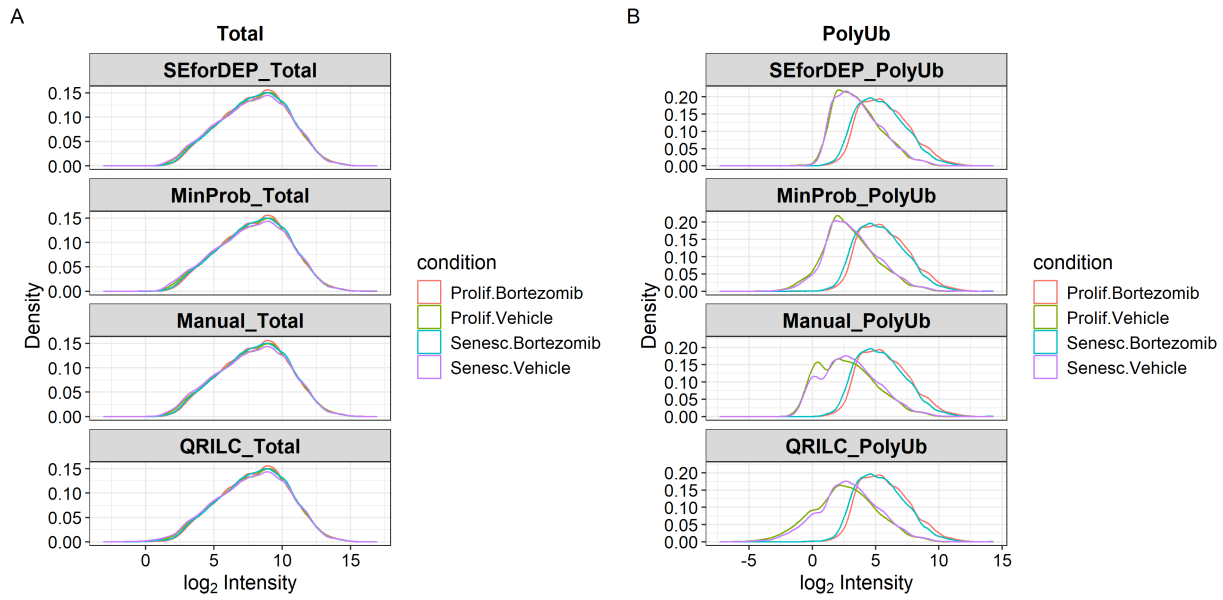

Figure 67: Density distributions of log2-transformed intensities with each imputation model across total (A) and polyUb (B) proteomes from the A549 bortezomib dataset.

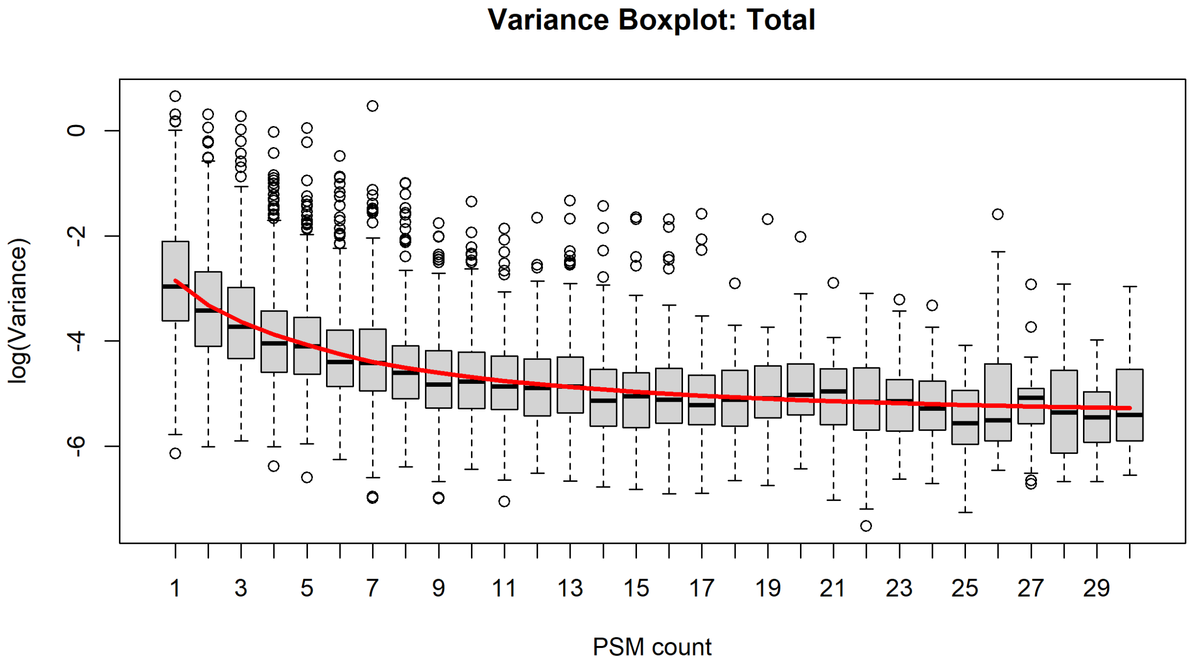

Figure 68: Boxplot showing variance of protein abundances with number of quantified PSMs in the total proteome from vehicle- and bortezomib-treated samples for the A549 bortezomib experiment.

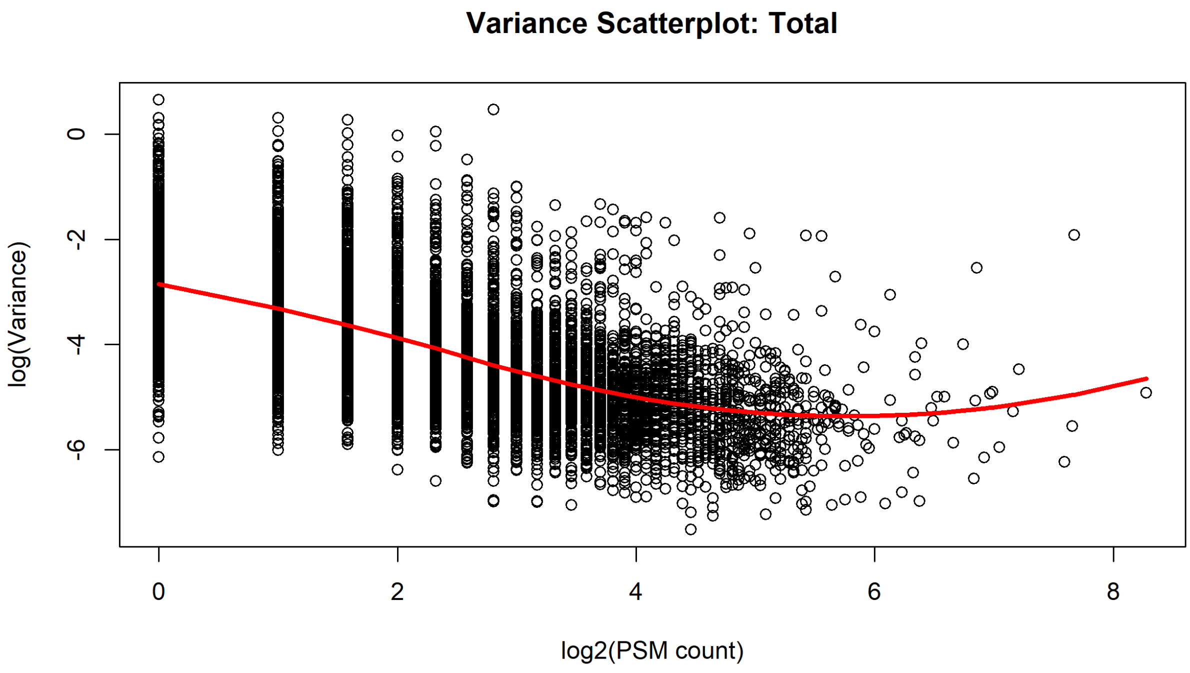

Figure 69: Scatterplot showing variance of protein abundances with number of quantified PSMs in the total proteome from vehicle- and bortezomib-treated samples for the A549 bortezomib experiment.

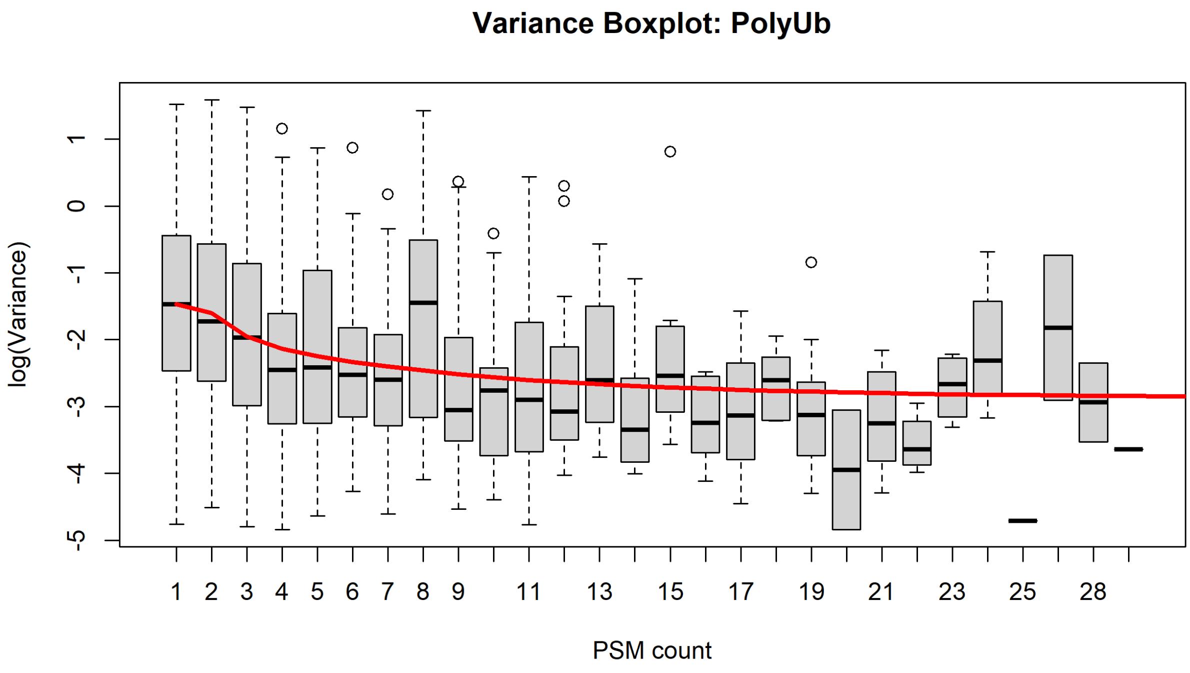

Figure 70: Boxplot showing variance of protein abundances with number of quantified PSMs in the total proteome for vehicle- and bortezomib-treated samples for the A549 bortezomib experiment.

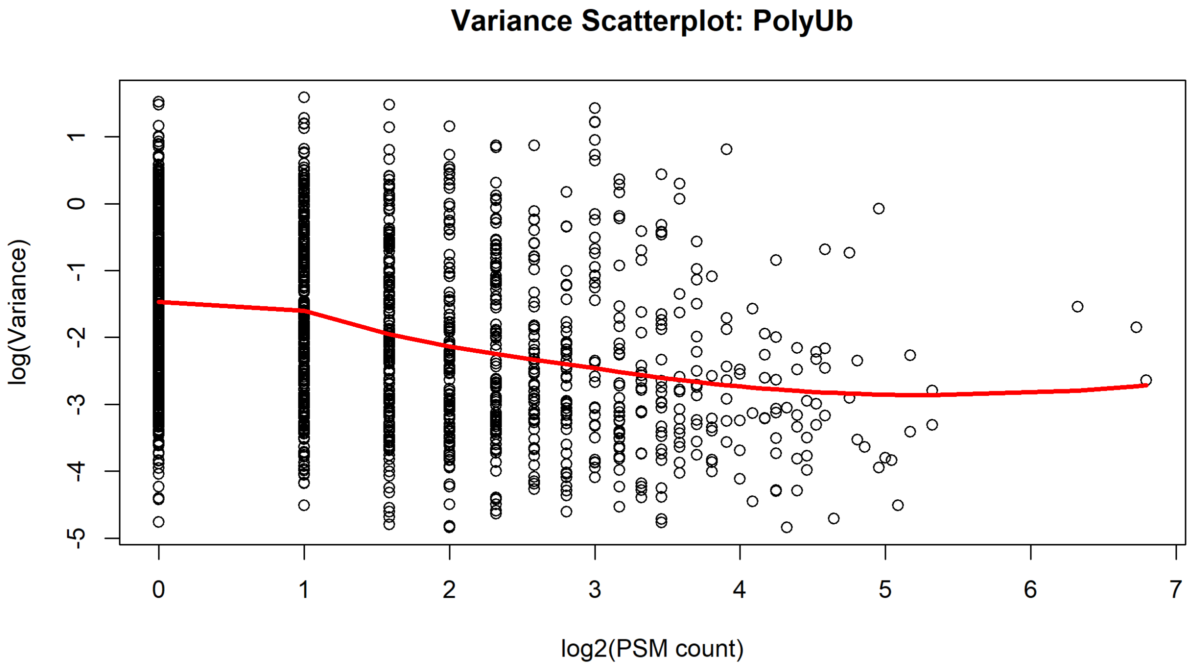

Figure 71: Scatterplot showing variance of protein abundances with number of quantified PSMs in the total proteome for vehicle- and bortezomib-treated samples for the A549 bortezomib experiment.
