## Supplementary material for "Declining intracellular proteostasis capacity drives misfolded protein secretion in senescent human cells": Key Resources Table

| **REAGENT or RESOURCE** | **SOURCE** | **IDENTIFIER** |
| --- | --- | --- |
| **Antibodies** | | |
| 53-BP1, rabbit polyclonal | Bethyl | Cat# A300-273A; RRID:AB_185521 |
| Vimentin (clone V9), mouse monoclonal | Thermo Fisher Scientific | Cat# MA5-11883; RRID:AB_10985392 |
| K48-Ubiquitin, sdAb, unconjugated C-terminal Cysteine, alpaca monoclonal | NanoTag Biotechnologies | Cat# N1805-250ug; RRID:AB_3076010 |
| Ubiquitin, Lys63-Specific (clone Apu3), rabbit monoclonal | Millipore | Cat# 05-1308; RRID:AB_1587580 |
| p62/SQSTM1 Lck ligand, mouse monoclonal | BD Biosciences | Cat# 610833; RRID:AB_398152 |
| GFP, rabbit polyclonal | ChromoTek | Cat# PABG1-20;  RRID:AB_2749857 |
| G3BP1 (clone 23), mouse monoclonal | BD Biosciences | Cat# 611127;  RRID:AB_398438 |
| MATR3, rabbit polyclonal | Proteintech | Cat# 12202-2-AP;  RRID:AB_2281752 |
| FUS/TLS (clone 4H11), mouse monoclonal | Santa Cruz Biotechnology | Cat# sc-47711;  RRID:AB_2105208 |
| TARDBP (TDP-43), rabbit monoclonal | Sigma | Cat# T1580;  RRID:AB_2532125 |
| TARDBP (TDP-43), mouse monoclonal | R&D Systems | Cat# MAB77781;  RRID:AB_3658921 |
| RAB5 (EPR21801), rabbit monoclonal | Abcam | Cat# ab218624;  RRID:AB_2892717 |
| TDP-43, rabbit polyclonal | Proteintech | Cat# 10782-2-AP; RRID:AB_615042 |
| DNAJC5, rabbit polyclonal | GeneTex | Cat# GTX66031 |
| beta-Tubulin (clone TUB 2.1), mouse monoclonal | Sigma | Cat# T4026; RRID:AB_477577 |
| Anti-Mouse IgG (H+L) Cross-Adsorbed, Alexa Fluor 488, goat polyclonal | Thermo Fisher Scientific | Cat# A-11001  RRID:AB_2534069 |
| Anti-Rabbit IgG (H+L) Cross-Adsorbed, Alexa Fluor 633, goat polyclonal | Thermo Fisher Scientific | Cat# A-21070  RRID:AB_2535731 |
| Anti-Rabbit IgG (Heavy-chain) Superclonal, Alexa Fluor 488, goat recombinant | Thermo Fisher Scientific | Cat# A27034; RRID:AB_2536097 |
| Anti-Mouse IgG (H+L) Superclonal, Alexa Fluor 488, goat recombinant | Thermo Fisher Scientific | Cat# A28175; RRID:AB_2536161 |
| Anti-Rabbit IgG (Heavy-chain) Superclonal, Alexa Fluor 647, goat recombinant | Thermo Fisher Scientific | Cat# A27040; RRID:AB_2536101 |
| Anti-Mouse IgG (H+L) Superclonal, Alexa Fluor 647, goat recombinant | Thermo Fisher Scientific | Cat# A28181; RRID:AB_2536165 |
| Anti-Rabbit IgG (H+L), DyLight 680, goat | Cell Signaling Technology | Cat# 5366; RRID:AB_10693812 |
| Anti-Mouse IgG (H+L), DyLight 680, goat | Cell Signaling Technology | Cat# 5470; RRID:AB_10696895 |
| Anti-Rabbit IgG (H+L), DyLight 800, goat | Cell Signaling Technology | Cat #5151; RRID:AB_10697505 |
| Anti-Mouse IgG (H+L), DyLight 800, goat | Cell Signaling Technology | Cat #5257; RRID:AB_10693543 |
| **Dyes** | | |
| Sulfo-Cy5 azide | Jena Bioscience | CLK-AZ118-1 |
| Janelia Fluor 549 maleimide | Tocris | 6500 |
| DAPI | Thermo Fisher Scientific | D1306 |
| Hoechst 33342, hydrochloride | Thermo Fisher Scientific | H3570 |
| SYTOX Orange | Thermo Fisher Scientific | S11368 |
| **Bacterial and virus strains** |  |  |
| *E. coli:* DH5-alpha (sub-cloning efficiency) | Thermo Fisher Scientific | 18265017 |
| *E. coli:* DH5-alpha (library efficiency) | Thermo Fisher Scientific | 18263012 |
| *E. coli: ccd*B Survival 2 T1^R^ | Thermo Fisher Scientific | A10460 |
| *E. coli:* NEB Stable | New England Biolabs | C3040I |
| *E. coli:* dam^–^/dcm^–^ | New England Biolabs | C2925I |
| **Chemicals, peptides, and recombinant proteins** | | |
| DMEM, high glucose | Gibco | 41965039 |
| L-Glutamine (200 mM) | Gibco | 25030081 |
| Penicillin-Streptomycin (10,000 U/mL) | Gibco | 15140122 |
| MEM Non-Essential Amino Acids Solution (100X) | Gibco | 11140050 |
| HyClone Characterized Fetal Bovine Serum, U.S. Origin, Heat-Inactivated | Cytiva | SH30071.02HI |
| Trypsin-EDTA (0.05%), phenol red | Gibco | 25300054 |
| Opti-MEM I Reduced Serum Medium | Gibco | 31985070 |
| Doxorubicin hydrochloride | Sigma-Aldrich | D1515 |
| Bleomycin (sulfate) | Cayman Chemicals | CAY13877 |
| Doxycycline monohydrate, 97% | Thermo Scientific Chemicals | 458890050 |
| Bortezomib | BOC Sciences | B0084-293315 |
| Bafilomycin A1 | Selleck Chemicals | S1413 |
| BGP-15 | Sigma-Aldrich | B4813 |
| SYBR Safe DNA Gel Stain | Invitrogen | S33102 |
| Polybrene | Sigma | TR-1003-G |
| DMSO, anhydrous | Molecular Probes | D12345 |
| Trypan blue solution | Sigma | T8154 |
| 16% Formaldehyde (w/v), Methanol-Free | Thermo Fisher Scientific | 28908 |
| Copper(II) sulfate pentahydrate | Sigma | 939315 |
| Ascorbic acid | Sigma | 1043003 |
| X-gal | Sigma | 11680293001 |
| Sodium phosphate (Na_2_HPO4) | Sigma | 71643 |
| Potassium ferricyanide(III) | Sigma | 702587 |
| Potassium ferrocyanide | Sigma | P3289 |
| Sodium chloride | Sigma | S9888 |
| Magnesium chloride solution | Sigma | M1028 |
| Phosphate buffered saline | Sigma | P2272 |
| EdU (5-ethynyl-2'-deoxyuridine) | Thermo Fisher Scientific | A10044 |
| Pierce Universal Nuclease for Cell Lysis | Thermo Fisher Scientific | 88701 |
| Sequencing-Grade Modified Trypsin | Promega | V5111 |
| TMTpro 18-plex Isobaric Label Reagent | Thermo Fisher Scientific | A52045 |
| Triethylammonium bicarbonate | Thermo Fisher Scientific | 90114 |
| Hydroxylamine | Thermo Fisher Scientific | 90115 |
| **Critical commercial assays** | | |
| Human TDP-43 ELISA kit | Proteintech | KE00005 |
| ChromoTek GFP-Trap Multiwell Plate | Proteintech | gtp-96 |
| Monarch Spin DNA Gel Purification Kit | New England Biolabs | T1120S |
| QIAprep Spin Miniprep Kit | Qiagen | 27104 |
| Gateway LR Clonase Enzyme mix | Invitrogen | 11791019 |
| GenElute HP Endotoxin-Free Plasmid Maxiprep Kit | Sigma | NA0410 |
| **Deposited data** | | |
| IMR-90 proliferating/quiescent/senescent, +/- HS, transcriptomes | This study | GEO: GSE307082 |
| IMR-90 proliferating/quiescent/senescent, +/- HS, total/polyUb/insoluble proteomes | This study | ProteomeXchange:PXD067225 |
| A549 proliferating/senescent, +/- HS, total/polyUb proteomes | This study | ProteomeXchange:PXD067153 |
| A549 proliferating/senescent, +/- bortezomib, total/polyUb proteomes | This study | ProteomeXchange:PXD067220 |
| **Experimental models: Cell lines** | | |
| IMR-90 | Coriell Institute, NJ USA | Cat# I90-10; RRID:CVCL_0347 |
| WI-38 | Coriell Institute, NJ USA | Cat# AG06814-M, RRID:CVCL_0579 |
| hTERT RPE-1 | ATCC (via René Medema, NCI, Netherlands) | Cat# CRL-4000, RRID:CVCL_4388 |
| A549 | ATCC | CCL-185 |
| A549 TetOn::GFP—Htt-exon1[polyQ97] | This study |  |
| A549 TetOn::GFP—Htt-exon1[polyQ25] | This study |  |
| **Oligonucleotides (all sequences 5’ to 3’)** | | |
| Cloning PCR primer Htt_For-BglII: tatcagatctATGGCGACCCTGGAAAAGC | This study |  |
| Cloing PCR primer Htt_Rev-NheI-STOP: attgctagctaAGGTCGGTGCAGAGGCT | This study |  |
| RT-qPCR primer HSPA1A-For: ACCTTCGACGTGTCCATCCTGA | This study |  |
| RT-qPCR primer HSPA1A-Rev:  TCCTCCACGAAGTGGTTCACCA | This study |  |
| RT-qPCR primer PUM1-For: CGGTCGTCCTGAGGATAAAA | This study |  |
| RT-qPCR primer PUM1_Rev: CGTACGTGAGGCGTGAGTAA | This study |  |
| **Recombinant DNA** | | |
| pMNLucPAUM 6HSE | Thomas Czerny,  FH Campus Wien, Austria | Ortner et al. (2015) <https://doi.org/10.1007/s12192-014-0540-5> |
| pME tdTomato−NLS | David Tobin (Addgene plasmid # 135305; <http://n2t.net/addgene:135205> | RRID: Addgene_135205 |
| pLenti CMV Puro DEST (w118-1) | Eric Campeau & Paul Kaufman (Addgene plasmid #17452; <http://n2t.net/addgene:17452>) | RRID: Addgene_17452 |
| pLenti CMV Puro tdTomato−NLS | This study |  |
| pLVX−ATF4 mScarlet NLS | David Andrews (Addgene plasmid # 115969; <http://n2t.net/addgene:115969>) | RRID: Addgene_115969 |
| pLVX 6xHSEp::mScarlet−NLS | This study | Internal # pRSS131 |
| pET28a−HaloTag−[tev]−[MCS] | MRC-PPU Reagents & Services, UK | Cat# DU23222 |
| pET28a T7pCONS TIR-2 sfGFP | Daniel Daley, (Addgene plasmid #154464; <http://n2t.net/addgene:154464>) | RRID: Addgene_154464 |
| pRSET−6xTR-TUBE | Yasushi Saeki (Addgene plasmid #110313; <http://n2t.net/addgene:110313>) | RRID: Addgene_110313 |
| pET28a HaloTag−[tev]−trTUBE_6_ | This study | Internal # pRSS114 |
| EYFP−CRAF | Laurence Pearl, U. Sussex, UK | Li et al. (2017) <https://doi.org/10.1016/j.celrep.2017.05.078> |
| pCW57.1 | David Root (Addgene plasmid #41393; <http://n2t.net/addgene:41393>) | RRID:  Addgene_41393 |
| pENTREGFP2 | Nathan Lawson (Addgene plasmid #22450; <http://n2t.net/addgene:22450>) | RRID:  Addgene_22450 |
| Halo−Sec61−C-18 | Kevin McGowan (Addgene plasmid #123285; <http://n2t.net/addgene:123285>) | RRID:  Addgene_123285 |
| pENTR HaloTag−[MCS] | This study | Internal # pRSS131 |
| pcDNA3 Htt-exon1[polyQ25]−mTurboID−HA | Judith Frydman, Stanford U., CA USA | Internal # pRSS063 |
| pcDNA3 Htt-exon1[polyQ97]−mTurboID−HA | Judith Frydman, Stanford U., CA USA | Internal # pRSS064 |
| pENTR HaloTag−Htt-exon1[polyQ25] | This study | Internal # pRSS141 |
| pENTR EGFP−Htt-exon1[polyQ25] | This study | Internal # pRSS145 |
| pENTR HaloTag−Htt-exon1[polyQ97] | This study | Internal # pRSS147 |
| pENTR EGFP−Htt-exon1[polyQ97] | This study | Internal # pRSS148 |
| pCW57.1 HaloTag−Htt-exon1[polyQ25] | This study | Internal # pRSS230 |
| pCW57.1 HaloTag−Htt-exon1[polyQ97] | This study | Internal # pRSS231 |
| pCW57.1 EGFP−Htt-exon1[polyQ25] | This study | Internal # pRSS238 |
| pCW57.1 EGFP−Htt-exon1[polyQ97] | This study | Internal # pRSS239 |
| pMD2.G | Didier Trono (Addgene plasmid #12259; <http://n2t.net/addgene:12259>) | RRID: Addgene_12259 |
| psPAX2 | Didier Trono (Addgene plasmid #12260; <http://n2t.net/addgene:12260>) | RRID: Addgene_12260 |
| pLenti eGFP Puro (w159-1) | Eric Campeau & Paul Kaufman (Addgene plasmid #17481; <http://n2t.net/addgene:17481>) | RRID: Addgene_17481 |
| pmScarlet-HSP70 | Vincent Timmerman (Addgene plasmid # 163790; <http://n2t.net/addgene:163790>) | RRIDL Addgene_163790 |
| pcDNA3.1 Flag-HA-HSP90AB1 | This study | Internal # pRSS027 |
| **Software and algorithms** | | |
| Fiji ImageJ (v1.52p) | <https://imagej.net/Fiji> | Schindelin et al. (2012) <https://doi.org/10.1038/nmeth.2019> |
| CellProfiler (v3.1.9) | <https://cellprofiler.org/> | McQuin et al. (2018) <https://doi.org/10.1371/journal.pbio.2005970> |
| StarDist plugin for ImageJ – Object Detection with Star-convex Shapes (v0.3.0) | <https://github.com/stardist/stardist-imagej/> | Schmidt et al. (2018) <https://arxiv.org/abs/1806.03535> |
| PerkinElmer Harmony 4.9 High-Content Imaging and Analysis Software | Revvity, UK | <https://www.revvity.com/gb-en/category/cellular-imaging-software> |
| R (v4.3.1) | <https://cran.r-project.org/> |  |
| tidyverse: Easily Install and Load the ‘Tidyverse’ (v2.0.0) | <https://doi.org/10.32614/CRAN.package.tidyverse> | <https://www.tidyverse.org> |
| drc: Analysis of Dose–Response Curves (v3.0-1) | <https://doi.org/10.32614/CRAN.package.drc> | Ritz et al. (2015) <https://doi.org/10.1371/journal.pone.0146021> |
| DEP: Differential Enrichment analysis of Proteomics data | <https://doi.org/doi:10.18129/B9.bioc.DEP> | Zhang et al. (2018)  <https://doi.org/10.1038/nprot.2017.147> |
| DEqMS: A tool to perform statistical analysis of differential protein expression for quantitative proteomics data (v1.26.0) | <https://doi.org/doi:10.18129/B9.bioc.DEqMS> | Zhu et al. (2020) <https://doi.org/10.1074/mcp.tir119.001646> |
| ggcorrplot: Visualization of a correlation matrix using ggplot2 (v0.1.4.1) | <https://doi.org/10.32614/CRAN.package.ggcorrplot> | Fissuh (2023) <https://rpubs.com/Alema/1000474> |
| org.Hs.eg.db: Genome wide annotation for Human (v3.20.0) | <https://doi.org/doi:10.18129/B9.bioc.org.Hs.eg.db> |  |
| VennDiagram: Generate High-Resolution Venn and Euler Plots (v1.7.3) | <https://doi.org/10.32614/CRAN.package.VennDiagram> | Chen et al. (2011)  <https://doi.org/10.1186/1471-2105-12-35> |
| eulerr: Area-Proportional Euler and Venn Diagrams with Ellipses (v7.0.2) | <https://doi.org/10.32614/CRAN.package.eulerr> | Larsson (2018) <http://lup.lub.lu.se/student-papers/record/8934042> |
| rstatix: Pipe-Friendly Framework for Basic Statistical Tests (v0.7.2) | <https://doi.org/10.32614/CRAN.package.rstatix> |  |
| STRING protein-protein interaction database (v12) | <https://string-db.org/> | Szklarczyk et al. (2021) <https://doi.org/10.1093/nar/gkaa1074> |
| Image Studio (v6.0) | <https://www.licorbio.com/image-studio> |  |
| **Other** | | |
| IncuCyte SX5 Live-Cell Analysis System | Sartorius | <https://www.sartorius.com/en/products/live-cell-imaging-analysis/live-cell-analysis-instruments/sx5-live-cell-analysis-instrument> |
| ImageXpress Confocal HT.ai High-Content Imaging System | Molecular Devices, CA USA | <https://www.moleculardevices.com/products/cellular-imaging-systems/high-content-imaging/imagexpress-confocal-ht-ai> |
| EVOS FL AMF4300 Cell Imaging System | Thermo Scientific | <https://assets.thermofisher.com/TFS-Assets/BID/manuals/MAN0007988_EVOS_FL_and_EVOS_FL_Color_UG.pdf> |
| Orbitrap Eclipse Tribrid Mass Spectrometer | Thermo Scientific | <https://www.thermofisher.com/order/catalog/product/FSN04-10000> |
| PerkinElmer Opera Phenix High-Content Screening System | Revvity | <https://www.revvity.com/gb-en/product/opera-phenix-plus-system-hh14001000> |
| LiCor Odyssey CLx Imager | LI-COR Biotech | <https://www.licorbio.com/support/answer-portal/imaging-systems/odyssey-clx.html> |
| Savant SpeedVac SPD210 Vacuum Concentrator | Thermo Scientific | SPD210P2-230 |
| CFX Opus 96 Real-Time PCR System | Bio-Rad | 12011319 |
| Whatman Puradisc-25 0.45μm PES syringe filter | Sigma-Aldrich | WHA67802504 |
| PhenoPlate 96-well, black, optically clear flat-bottom, tissue-culture treated | Revvity, UK | 6055300 |
| Nunc Microwell 96-well, Nunclon Delta-treated, flat-bottom microplate | Thermo Scientific | 167008 |
| Zeba Spin Desalting Columns 7 kDa MWCO | Thermo Scientific | 89882 |
| 0.5 mL Protein LoBind Tube | Eppendorf | 0030108094 |
| 2 mL Protein LoBind Tube | Eppendorf | 0030108132 |
| Phasemaker TRIzol Reagent and Phasemaker Tubes Complete System | Thermo Scientific | A33251 |
| GlycoBlue Coprecipitant | Thermo Scientific | AM9516 |
| Invitrogen DNA-free DNA Removal Kit | Thermo Scientific | AM1906 |
| High-Capacity cDNA Reverse Transcription Kit | Thermo Scientific | 4368814 |
| iTaq Universal SYBR Green Supermix | Bio-Rad | 1725121 |
| Magne HaloTag Beads | Promega | G7281 |
| Countess Cell Counting Chamber Slides | Thermo Scientific | C10228 |
| Pierce Protein Concentrator, PES, 3 kDa MWCO, | Thermo Scientific | 88526 |
| Pierce Protein Concentrator, PES, 3 kDa MWCO | Thermo Scientific | 88515 |
