## Supplementary Information for "Declining intracellular proteostasis capacity drives misfolded protein secretion in senescent human cells"

#### Supplementary Figures S1–S14 (and Supplementary Figure Legends)

**Supplementary Figure S1:** Validation of senescence models used in this study.

**Supplementary Figure S2:** Quiescent and senescent states exhibit increased proteotoxic stress resilience across cell types and models.

**Supplementary Figure S3:** Global transcriptomics and proteomics reveals delayed heat-shock-response induction in senescent and quiescent states.

**Supplementary Figure S4:** Differential analysis of total proteome reveals limited differences between heat-shock-modulated proteins across cell states.

**Supplementary Figure S5:** Differential analysis of polyUb-enriched proteome reveals limited differences between heat-shock-modulated proteins across cell states.

**Supplementary Figure S6:** Total and polyUb-enriched proteomics of A549 human lung adenocarcinoma cells reveals similarly conserved responses between proliferating and senescent states to heat-shock and proteasome inhibition.

**Supplementary Figure S7:** Differential analysis of insoluble proteome reveals dampened RNA-related protein granule formation in the senescent state.

**Supplementary Figure S8.** Nuclear RBPs TDP-43 and MATR3, but not FUS, form stress-induced nuclear foci that diverge in magnitude across non-proliferative states.

**Supplementary Figure S9.** Misfolded but not well-folded proteins are secreted upon stress in senescent states, across cell types and senescence-induction models.

**Supplementary Figure S10.** Peri-centriolar satellite proteins shown to be important for aggresome formation are consistently declined in the senescent state.

**Supplementary Figure S11.** Aggresome formation is impaired in the senescent state.

**Supplementary Figure S12.** A549 cells form aggresomes containing misfolded proteins in proliferating but not senescent states.

**Supplementary Figure S13.** Limited molecular chaperone alterations in senescence that are not shared with quiescence.

**Supplementary Figure S14.** DNACJ5 is not important for senescence induction trajectories, but does impact misfolded protein secretion in a manner that can be rescued by chaperone induction.

#### Supplementary Methods

### Supplementary Files (separate files)

**Key Resources Table.docx**
**Supplementary-File-S1.pdf** Processing of multi-dimensional proteomics data.

### Supplementary Figure S1

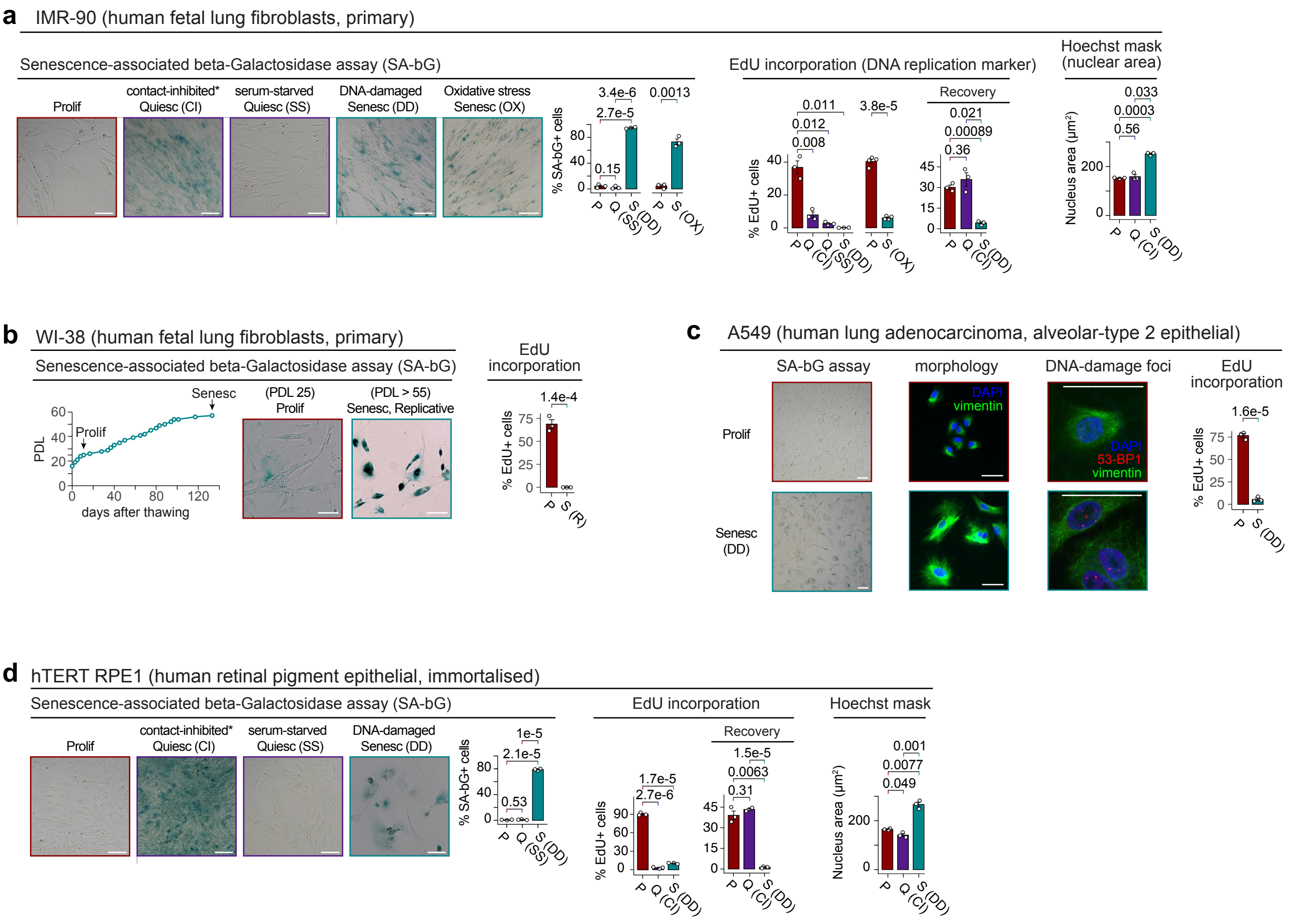

### Supplementary Figure S2

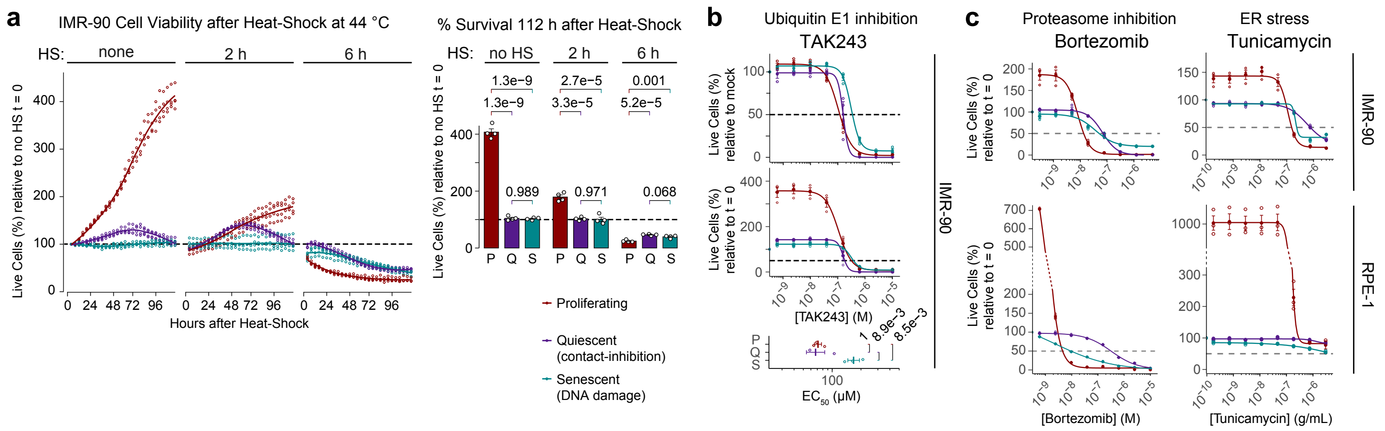

### Supplementary Figure S3

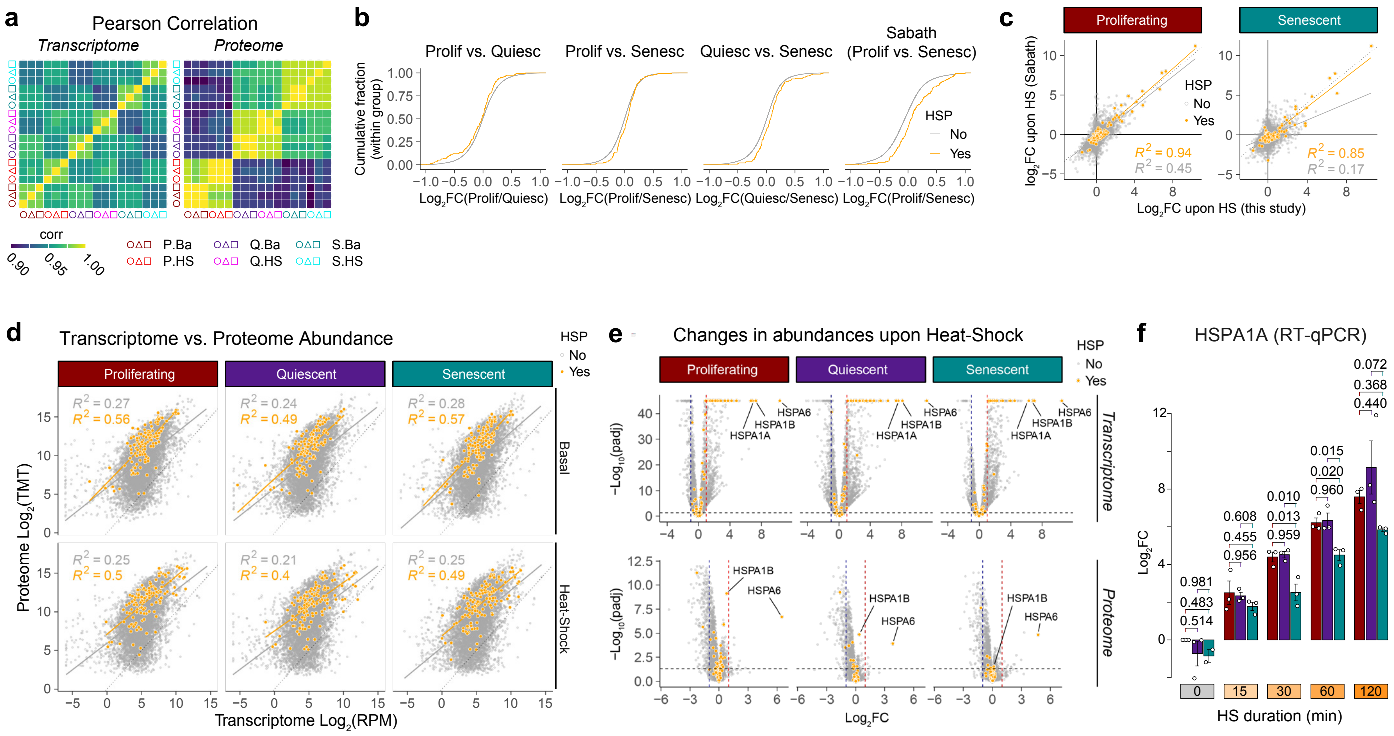

### Supplementary Figure S4

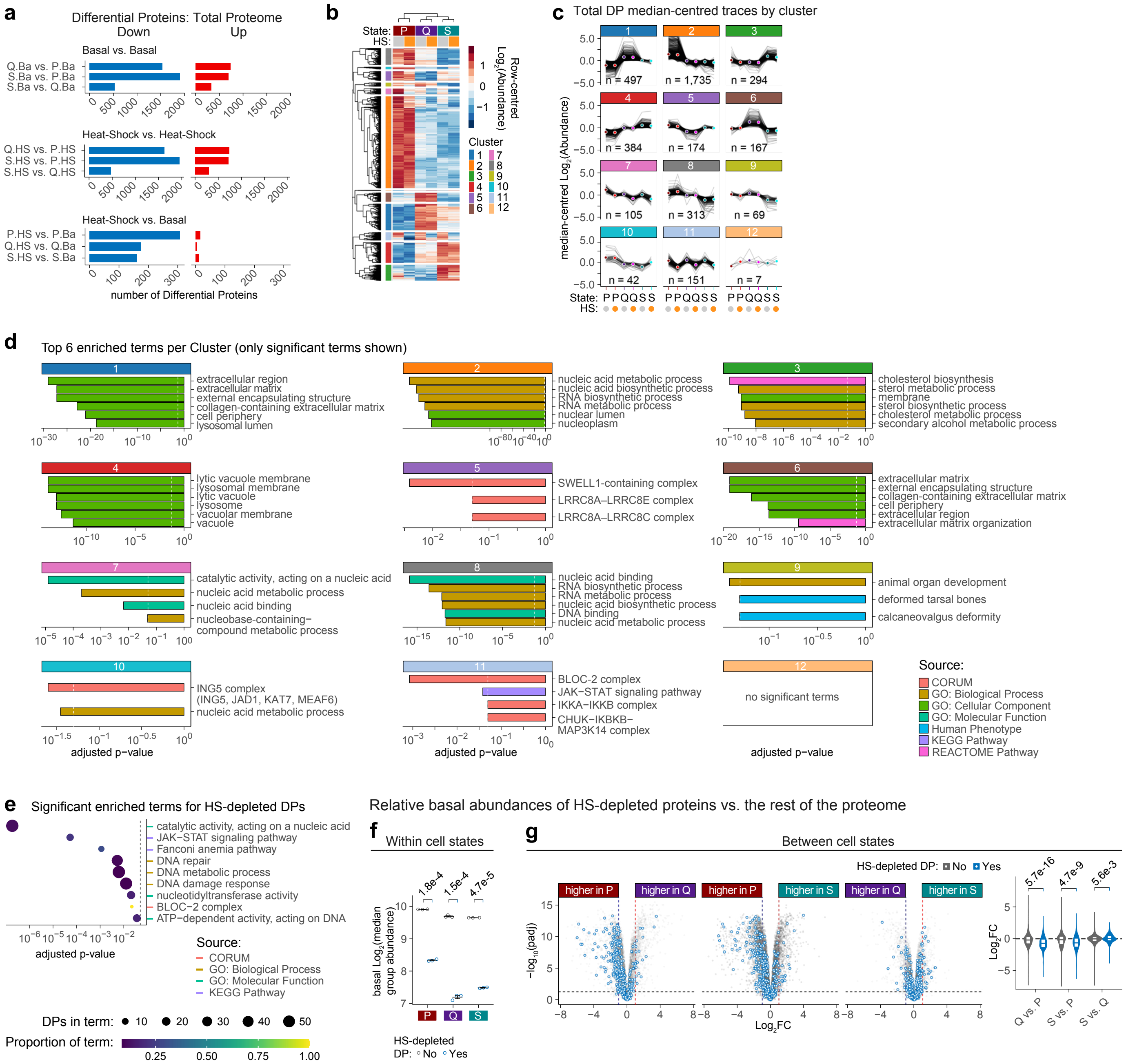

#### Supplementary Figure S5

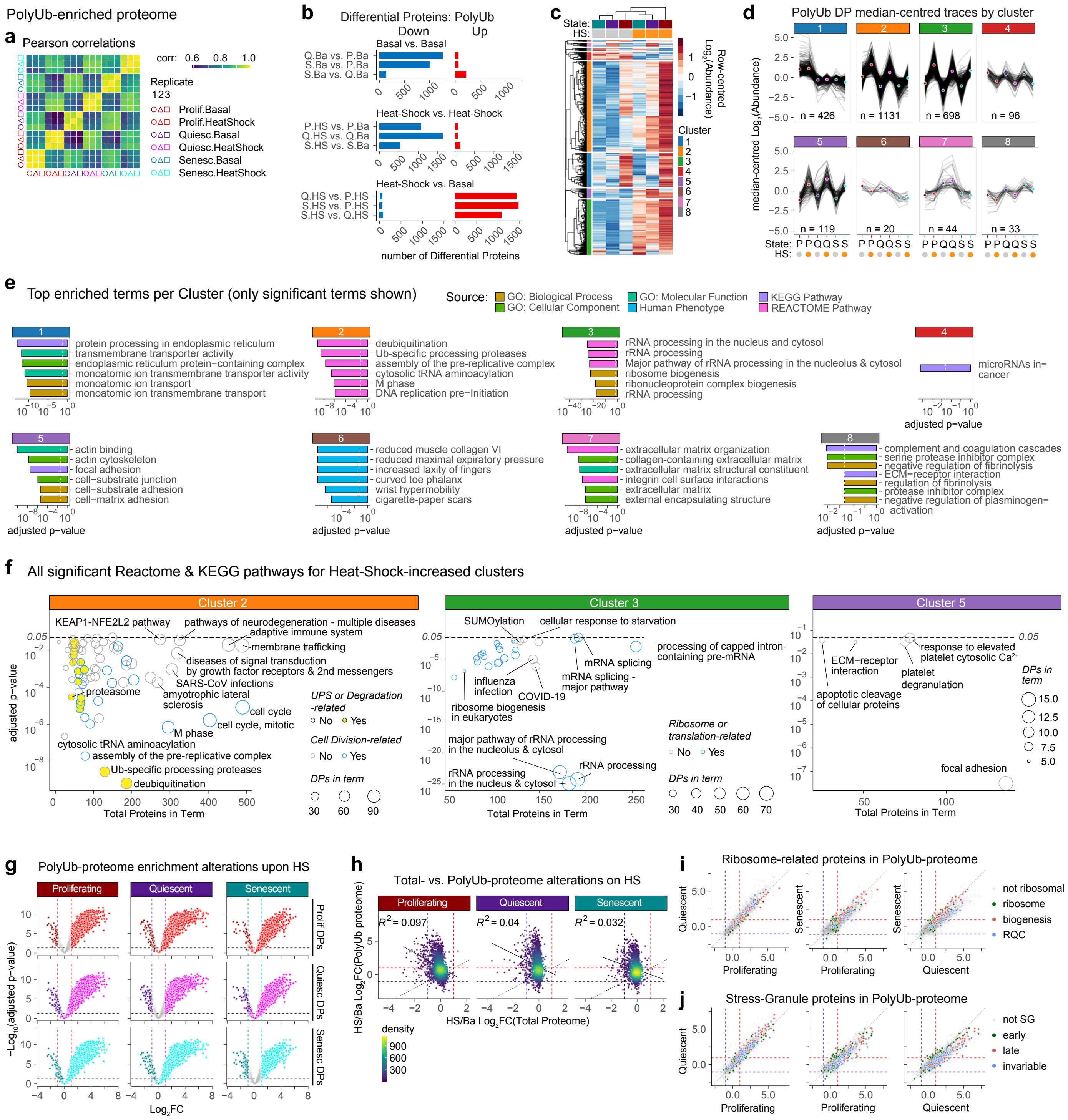

### Supplementary Figure S6

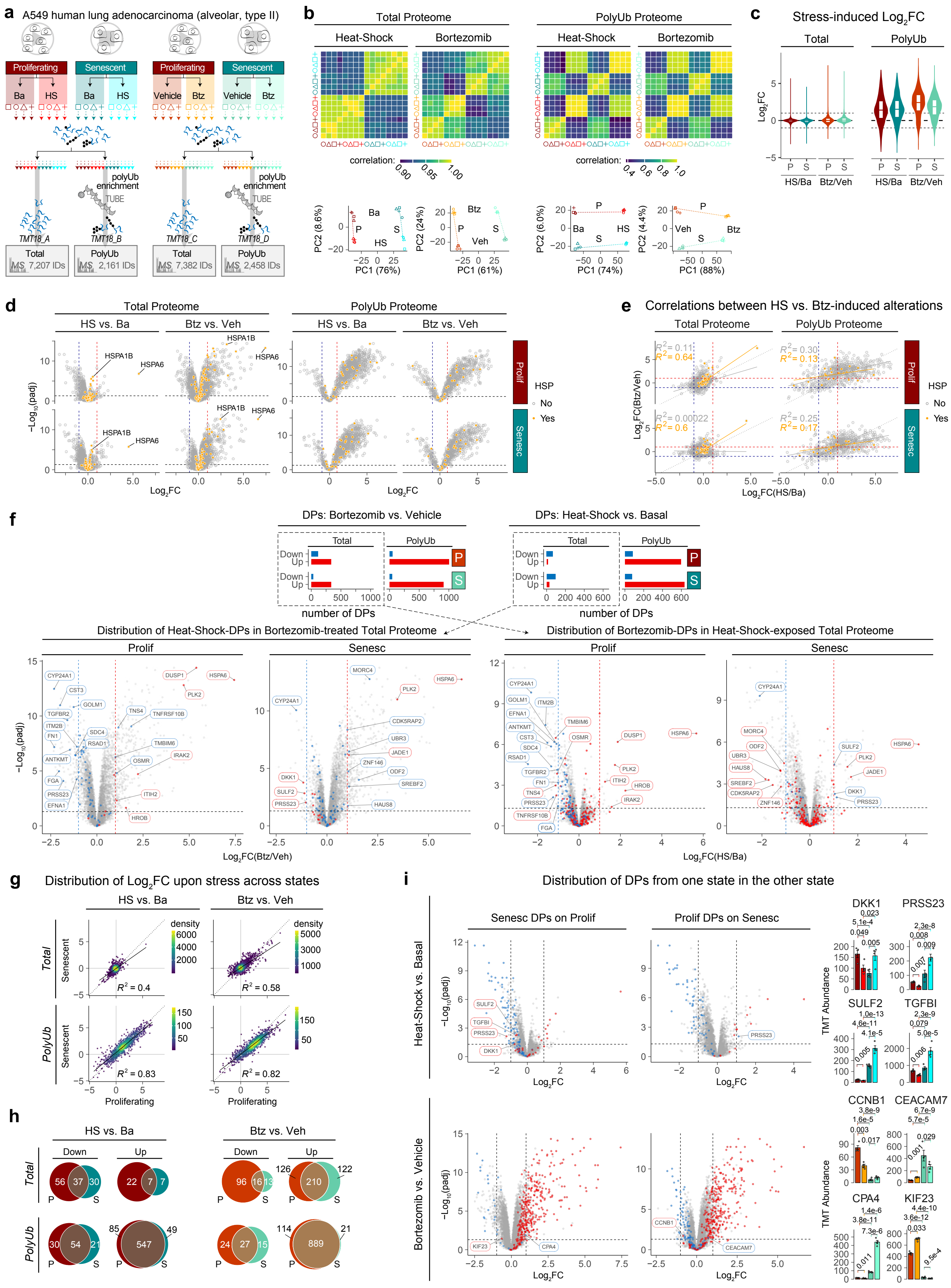

### Supplementary Figure S7

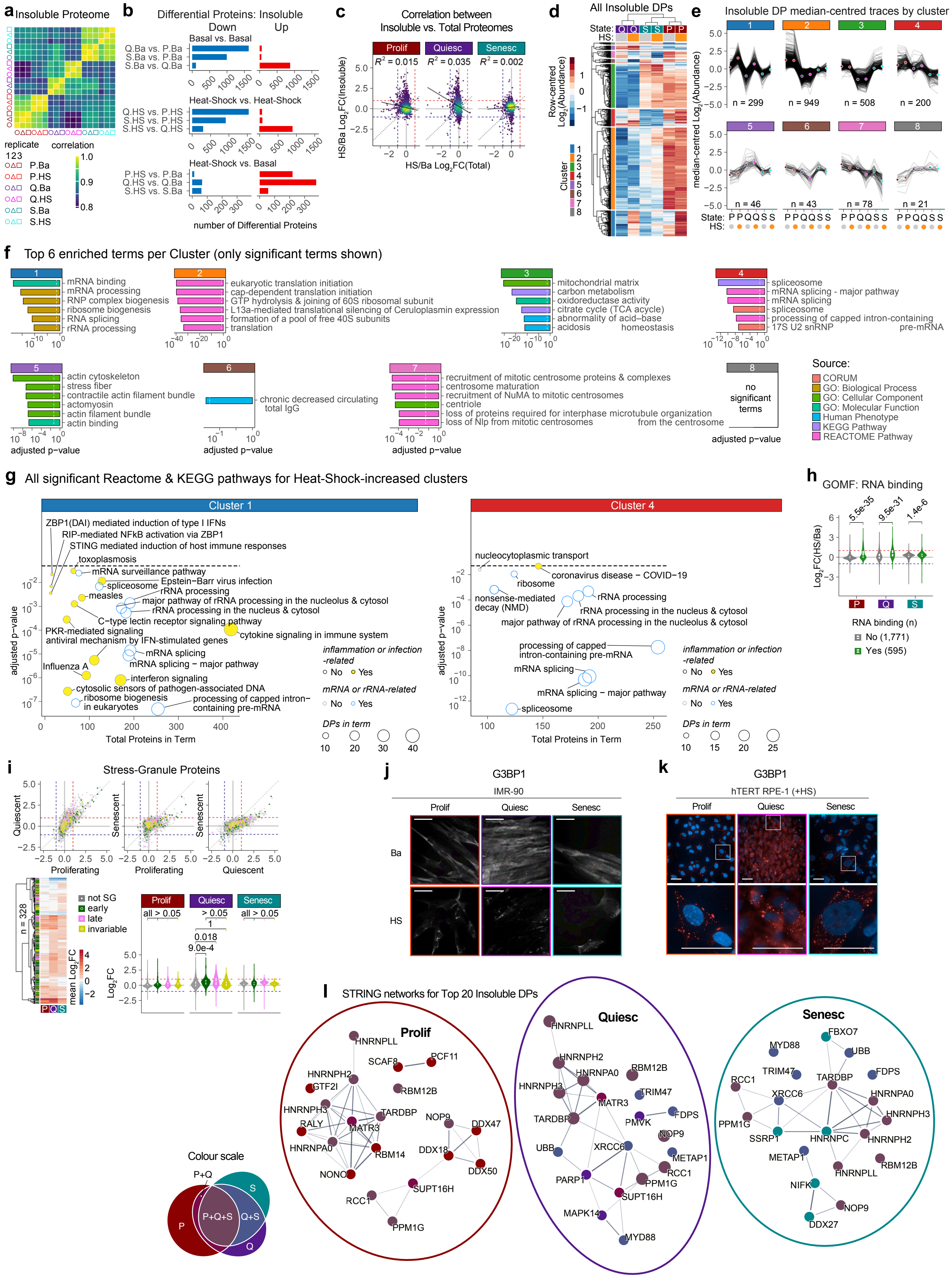

#### Supplementary Figure S8

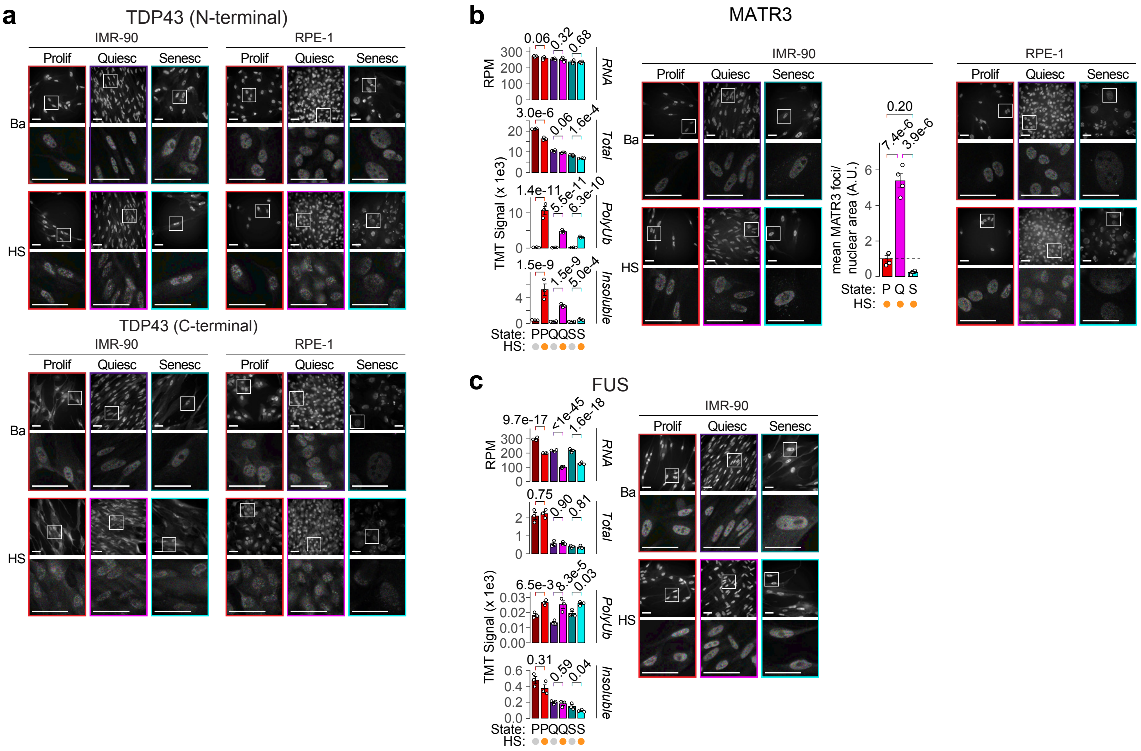

### Supplementary Figure S9

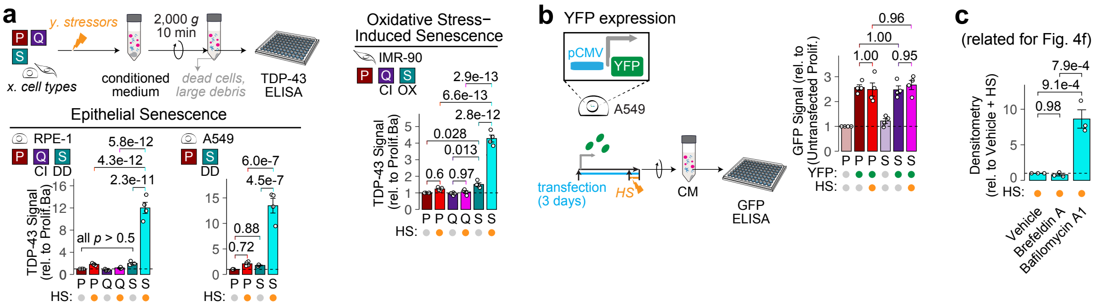

### Supplementary Figure S10

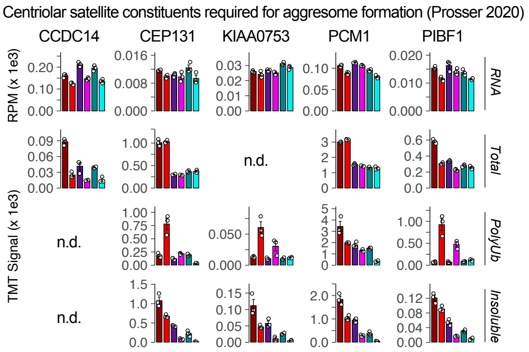

#### Supplementary Figure S11

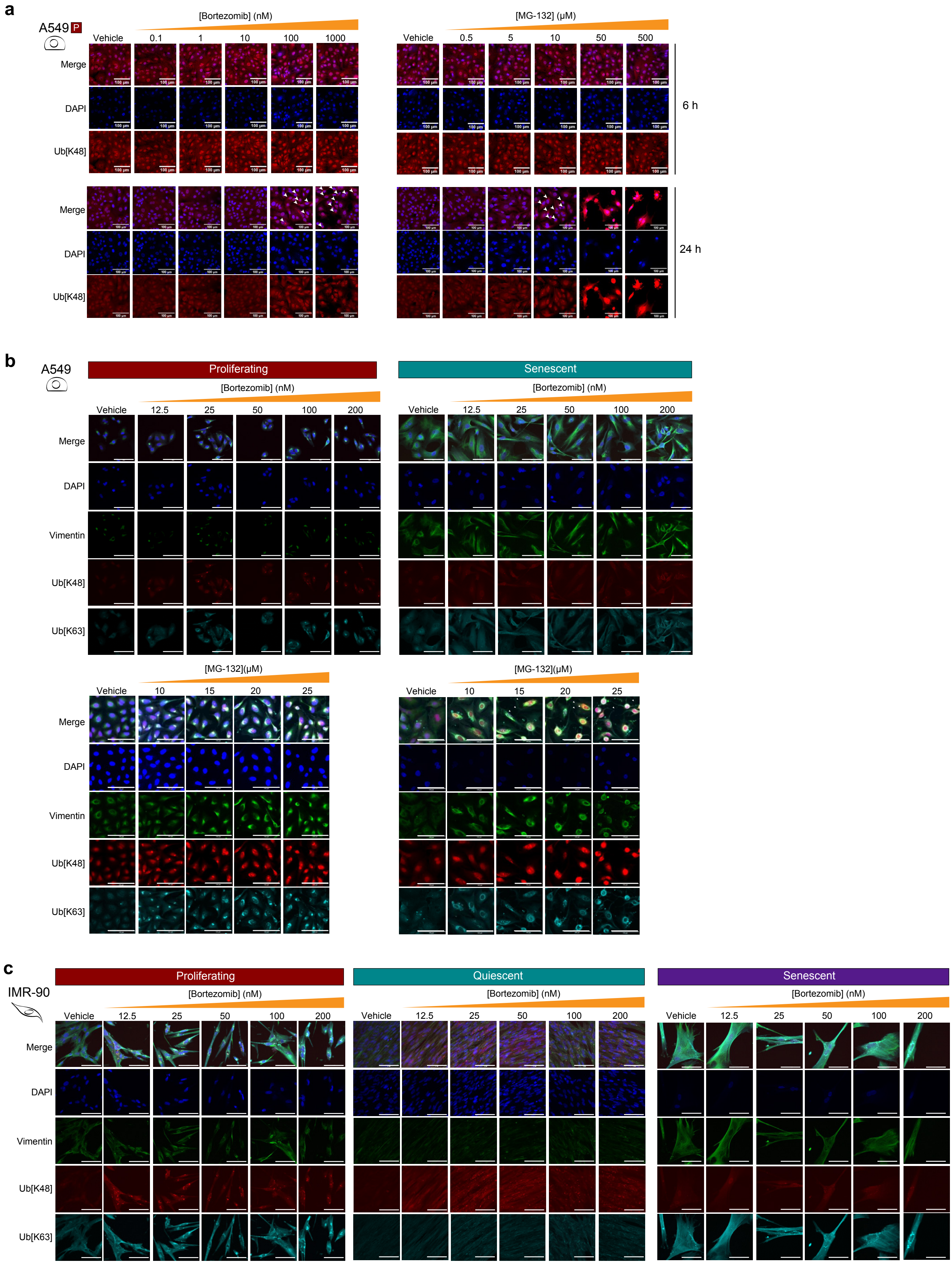

#### Supplementary Figure S12

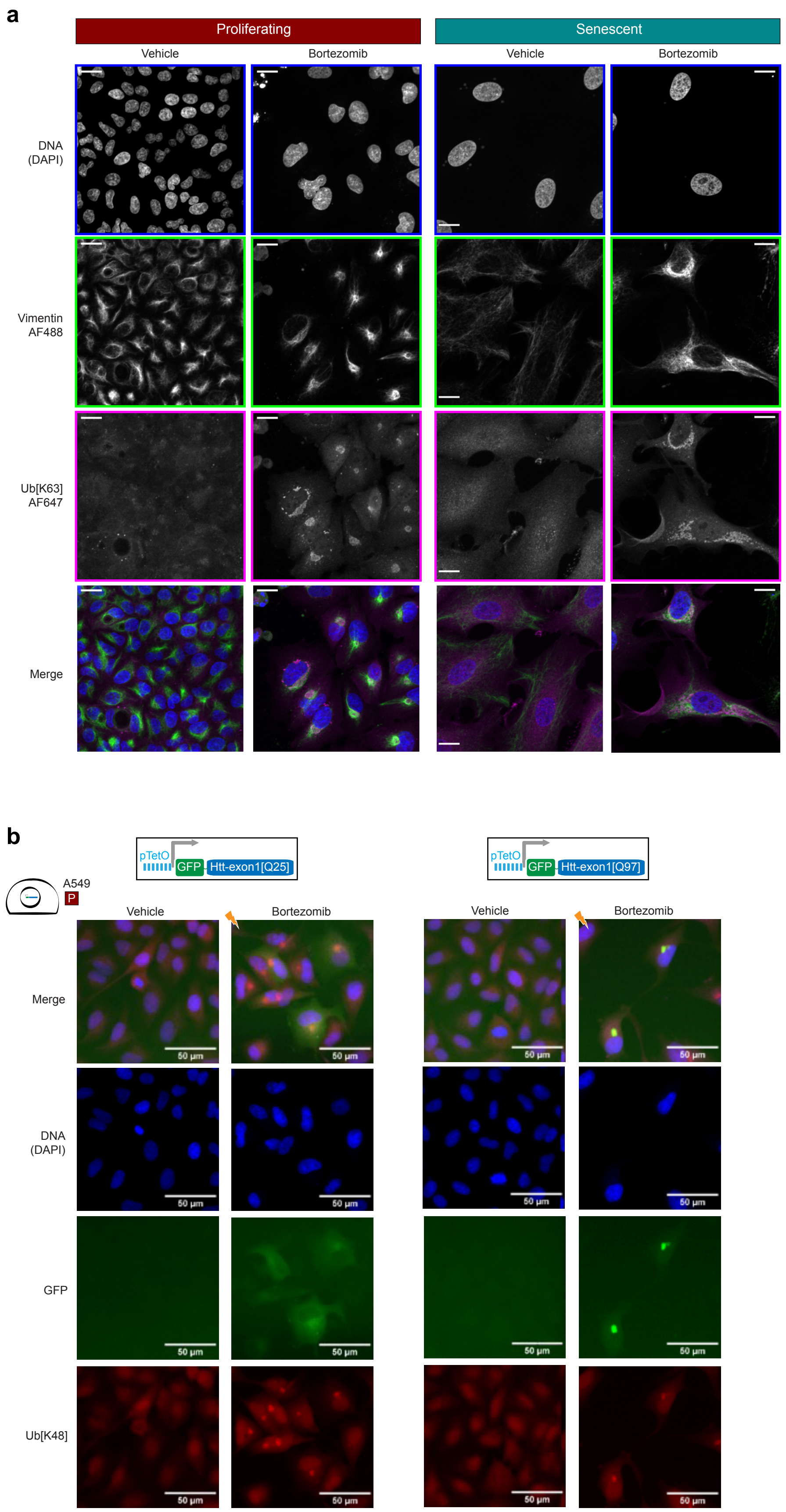

### Supplementary Figure S13

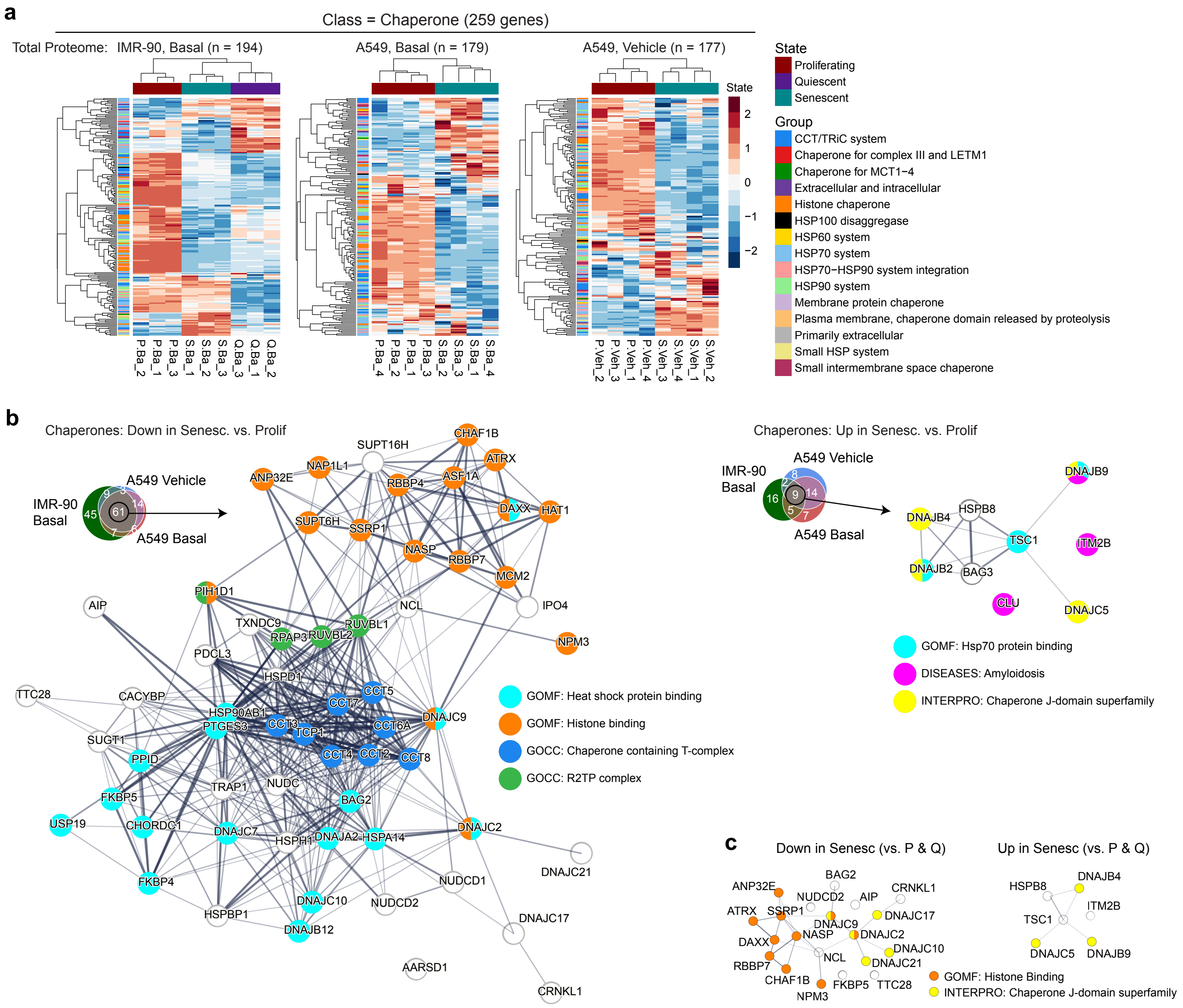

### Supplementary Figure S14

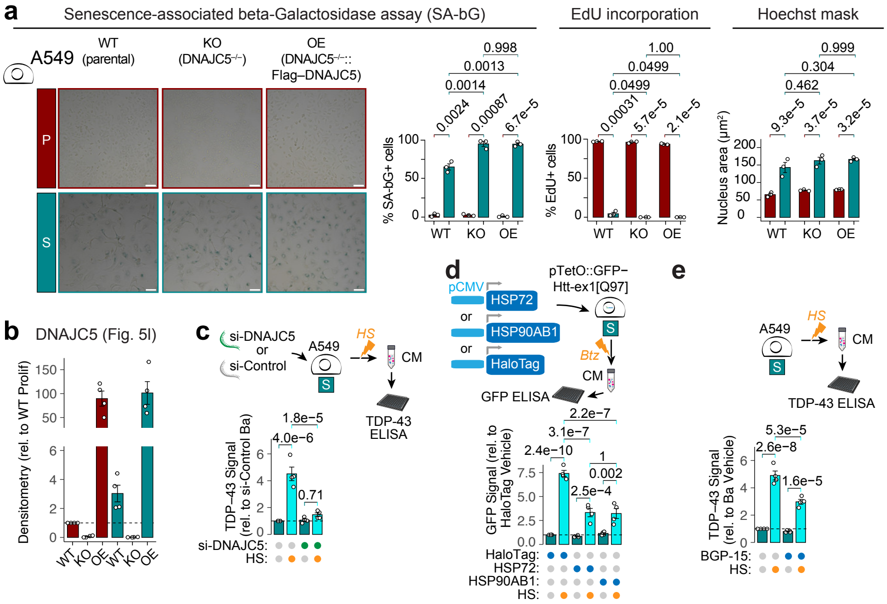

#### Supplementary Figure Legends

**Supplementary Figure S1. Validation of senescence models used in this study.**

Validation of senescence models used in this study, by SA-beta-Galactosidase assay (SA-bG), 24 h EdU incorporation, and area of nuclear mask by Hoechst 33342 or DAPI staining. *Note that contact-inhibited quiescent states also stained positive in the SA-bG assay, as per previous reports.[1] To distinguish these contact-inhibited states from senescent states, cells were trypsinised and returned to sub-confluent culture conditions for two days prior to 24 h EdU incubation ('Recovery'). All scale bars represent 100 µm, except for DNA-damage foci images in C (50 µm).

**Supplementary Figure S2. Quiescent and senescent states exhibit increased proteotoxic stress resilience across cell types and models.**

**a.** IMR-90 fibroblasts expressing tdTomato–NLS nuclear marker in proliferating, contact-inhibited quiescent, or DNA-damage-induced senescent states were heat-shocked for 2 h or 6 h at 44 °C, or kept at 37 °C ('none'), before returning to regular culture conditions (t = 0) with SYTOX Green dead-cell stain, and monitoring every 4 h on an Incucyte SX5 Live-Cell Analysis System. Left, number of live cells were calculated at each time-point by subtracting SYTOX+ from tdTomato+ nuclei counts, both quantified using the StarDist plugin in ImageJ. Right, Differences in percentage of surviving cells between each state per heat-shock condition at the end of the experiment (112 h time-point). See also Fig. 1b.

**b–c.** IMR-90 fibroblasts or RPE1 epithelia in proliferating, contact-inhibited quiescent, or DNA-damage-induced senescent states were treated with a range of concentrations TAK243 or bortezomib for 72 h, or tunicamycin for 120 h, before staining with Hoechst 33342 and SYTOX Orange dead-cell stain, and imaging immediately on an ImageXpress Confocal HT.ai High-Content Imaging System. Number of live cells was calculated as in (a), and normalised as a percentage of either the DMSO-treated vehicle control for that cell state at the end of the experiment (t = 72 or 120 h), or the live-cell numbers at the start of the experiment (t = 0). Dose-response curves and $EC_{50}$ values were calculated using the drc R package. See also Fig. 1c–d.

Open-circle data-points represent individual biological replicates (n = 3/4), and filled-circles or bars +/– error-bars represent mean +/– standard-error. Adjusted p-values from Tukey's HSD post-hoc test following significant ($p < 0.05$) one-way ANOVA are shown.

**Supplementary Figure S3. Global transcriptomics and proteomics reveals delayed heat-shock-response induction in senescent and quiescent states.**

**a.** Pairwise Pearson correlation coefficients of the $Log_2$-transformed normalised RPM (transcriptome) or TMT abundances (total proteome) between all 18 samples.

**b.** HS-induction of HSPs is mildly impaired in quiescent and senescent vs. proliferating states, but not to the same extent as replicatively-senescent vs. low PDL WI-38 fibroblasts from Sabath et al. (2020).[2] Cumulative distribution function (CDF) plots showing the difference in HS-induced $Log_2FC$ values for HSPs (orange) vs. the rest of the transcriptome (grey) between states in our transcriptomics data, or from Sabath et al.

**c.** HS-induced $Log_2FC$ values for our transcriptomics data (x-axis) vs. Sabath et al. (y-axis), indicates strong correlation for HSPs (orange) between studies across proliferating and senescent states, but poorer correlation for the rest of the transcriptome (grey), especially for the senescent state.

**d.** Global-normalised and $Log_2$-transformed mean RPM (transcriptome, x-axes) or TMT abundances (proteome, y-axes) for each cell state and condition.

For c–d: For each pairwise comparison, linear regression lines for $y \sim x$, and squared Pearson's correlation coefficients, for HSPs (orange) vs. the rest of the transcriptome (grey), are shown. Dotted lines represent $y = x$.

**e.** Volcano plots showing heat-shock-induced transcriptome (top) or proteome (bottom) alterations in proliferating, quiescent, and senescent states. HSPs (orange) are highlighted vs. the rest of the transcriptome/proteome, and the HSP70s HSPA1A, HSPA1B, and HSPA6 are labelled (note that HSPA1A and HSPA1B have identical amino-acid sequences, and peptides corresponding to these are arbitrarily assigned 'HSPA1B' in the proteomics data). $Log_2FC$ = 1, –1, and adjusted p = 0.05, are represented by dashed red, blue, and black lines, respectively.

**f.** HS time-course suggests a mild dampening of HSPA1A mRNA induction in the senescent state. IMR-90 cells in proliferating, contact-inhibited quiescent, or DNA-damage-induced senescent states were exposed to HS at 44 °C for the indicated duration, before immediate extraction nucleic acids in Trizol, and quantification of mRNA for HSPA1A, and the housekeeping genes PUM1 for normalisation. Open-circle data-points represent individual biological replicates (n = 3, except for Senescent 0 h, where one replicate did not yield sufficient RNA), and filled-circles or bars +/– error-bars represent mean +/– standard-error. Adjusted p-values from Tukey's HSD post-hoc test following significant ($p < 0.05$) one-way ANOVA are shown.

**Supplementary Figure S4. Differential analysis of total proteome reveals limited differences between heat-shock-modulated proteins across cell states.**

**a.** Total numbers of differential proteins (DPs) based on DEqMS-calculated cut-offs of adjusted $p < 0.05$ and $Log_2FC >= 1$ for increased (up, red) or <= –1 for decreased (down, blue), between the pairwise comparisons indicated for proliferating (P), quiescent (Q), or senescent (S) states under basal (Ba) or heat-shock (HS) conditions.

**b.** Heatmap of row-centred mean $Log_2$-transformed normalised TMT abundances for all DPs in the total proteome, based on the cut-offs described in A. Hierarchical clustering was performed using the complete agglomeration method with Euclidean distance measure for both columns (conditions) and rows (proteins). The row dendrograms were separated into 12 clusters.

**c.** Traces of the median-scaled $Log_2FCs$ for each DP across the six conditions, separated by cluster. The mean of the median-scale $Log_2FCs$ for all proteins in the cluster per condition is overlayed as circles.

**d–e.** Top significant (adjusted $p < 0.05$, dashed line) annotation terms by over-representation analysis (ORA) for the proteins in each cluster (d), or all 368 heat-shock-depleted proteins (e). For (e), Size of the circle depicts the number of DPs in each term, the fill colour of the circle depicts what proportion of the total number of proteins in the term is represented by the DPs here, and y-axis tick colour depicts the source of the enriched term. ORA was performed using the gprofiler2 R package, with all 8,449 quantified in the IMR-90 total proteome as the custom background. Sources for annotation terms are indicated by the bar colours: CORUM, CORUM mammalian protein complex database; GO BP, CC, MF, Gene Ontology Biological Process, Cellular Compartment, Molecular Function; HP, Human Phenotype ontology; KEGG, Kyoto Encyclopaedia of Genes and Genomes pathway; REAC, Reactome Pathway Database.

**f.** Heat-shock-depleted DPs have on average lower basal abundance values than the rest of the quantified total proteome within each cell state. Each point represents the median $Log_2$-transformed abundance in the basal (i.e., non-heat-shocked) condition of all 368 heat-shock-depleted DPs (blue) vs. the rest of the quantified proteome (grey) for a replicate. Mean and standard-error summary statistics per group are overlayed, and adjusted p-values from two-tailed tests with Bonferroni multiple-testing correction between groups per cell state are shown.

**g.** Heat-shock-depleted DPs have on average lower abundance values in non-proliferating states, when compared with the rest of the quantified proteome. Left, volcano plots showing pair-wise comparisons between the three states in their basal (i.e., non-heat-shocked) protein total abundances, with the heat-shock-depleted DPs (blue) highlighted against the rest of the proteome (grey). Right, violin plots showing the distribution of Log2-transformed fold-change values for the total protein abundances of all 368 heat-shock-depleted DPs (blue) or the rest of the proteome (grey), at basal conditions. Adjusted p-values from two-tailed t-tests with Bonferroni multiple-testing correction between the two groups are shown for each pairwise comparison.

**Supplementary Figure S5. Differential analysis of polyUb-enriched proteome reveals limited differences between HS-modulated proteins across cell states.**

**a.** Pairwise Pearson correlation coefficients of the $Log_2$-transformed normalised TMT abundances of the PolyUb–proteomes between all 18 samples indicate the highest degree of similarity between samples in the same condition (Ba or HS), and then by cell state.

**b.** Total numbers of differential proteins (DPs) increased (up, red) or decreased (down, blue) between the pairwise comparisons indicated for proliferating (P), quiescent (Q), or senescent (S) states under basal (Ba) or heat-shock (HS) conditions in the PolyUb–proteome.

**c.** Heatmap of row-centred mean $Log_2$-transformed normalised TMT abundances for all DPs in the PolyUb–proteome, based on the cut-offs described in (b). Hierarchical clustering was performed using the complete agglomeration method with Euclidean distance measure for both columns (conditions) and rows (proteins). The row dendrograms were separated into 8 clusters.

**d.** Traces of the median-scaled $Log_2FCs$ for each DP across the six conditions in the PolyUb–proteome, separated by cluster. The mean of the median-scale $Log_2FCs$ for all proteins in the cluster per condition is overlayed as circles.

**e.** Top significant (adjusted $p < 0.05$, dashed line) annotation terms by over-representation analysis (ORA) for the proteins in each cluster in the PolyUb–proteome. ORA was performed using the gprofiler2 R package, with all 8,449 quantified in the IMR-90 total proteome as the custom background. Sources for annotation terms are indicated by the bar colours: GO BP, CC, MF, Gene Ontology Biological Process, Cellular Compartment, Molecular Function; HP, Human Phenotype ontology; KEGG, Kyoto Encyclopaedia of Genes and Genomes pathway; REAC, Reactome Pathway Database.

**f.** All significant (adjusted $p < 0.05$, dashed line) Reactome and KEGG pathway annotation terms by ORA (from (e)) for proteins in the HS-increased clusters from the PolyUb–proteome. Size of the circle depicts the number of DPs in each term. Terms related to specific biological features (e.g., UPS, ubiquitin–proteasome system) were manually annotated and coloured as shown in the legend for each cluster.

**g.** Volcano plots showing heat-shock-induced polyUb–proteome alterations in proliferating, quiescent, and senescent states. The values for DPs from proliferating (reds), quiescent (purples), or senescent (turquoises) states are plotted onto the other two states, to determine whether direction of changes are conserved across states. Absolute $Log_2FC = 1$, and adjusted $p = 0.05$, are represented by dashed vertical and horizontal lines, respectively.

**h.** HS-induced $Log_2FC$ values across each cell state for the total (x-axis) vs. PolyUb-enriched (y-axis) proteomes. For each pairwise comparison, linear regression lines for $y \sim x$, and squared Pearson's correlation coefficients, are shown. Density of overlapping points are indicated with the colour scale.

**i–j.** Heat-shock-induced $Log_2FC$ values in the PolyUb–proteome for each state, with ribosome–[3] (i) or stress-granule–[4] (j) annotated proteins highlighted.

For h–j: Dotted lines represent $y = x$, and dashed lines represent $Log_2FC = –1$ (blue) or $= 1$ (red).

**Supplementary Figure S6. Total and polyUb-enriched proteomics of A549 human lung adenocarcinoma cells reveals similarly conserved responses between proliferating and senescent states to HS and proteasome inhibition.**

**a.** Experimental design for total proteome and tandem-ubiquitin-binding-entity (TUBE)–based polyUb–proteome quantification of A549 human lung adenocarcinoma cells. Cells in proliferating or senescent (by DNA-damage via bleomycin) states were exposed to two different proteotoxic stressors across two experiments. For the HS experiment, cells were moved to a 44 °C (Heat-Shock, HS), or kept at 37 °C (Basal, Ba) for 2 h. For the proteasome inhibition experiment, cells were treated with 100 nM bortezomib (Bortezomib, Btz), or equal volume of the DMSO vehicle (Vehicle, Veh), for 24 h. Samples were harvested and lysed immediately at the end of each stress exposure, and processed for total and polyUb-enriched proteome quantification, as described for the IMR-90 fibroblasts.

**b.** Top, pairwise Pearson correlation coefficients of the $Log_2$-transformed normalised TMT abundances between all 16 A549 samples per experiment. Bottom, raw component scores from Principal Component Analysis (PCA) of the top two PCs for each experiment. Percentage of variance explained by each PC is indicated in the axis titles.

**c.** Global distribution of $Log_2FC$ values (from DEqMS differential analysis) upon heat-shock or proteasome inhibition in each proteome and state are plotted as violin and overlaid boxplots (median + inter-quartile range); dashed black and grey horizontal lines represent absolute $Log_2FC$ = 0 and 1, respectively.

**d.** Volcano plots showing $Log_2$-transformed fold-change ($Log_2FC$) and negative-$Log_{10}$-transformed adjusted p-values ($–Log_{10}$(padj)), both calculated by DEqMS, for each A549 proteomics experiment. HSPs are highlighted in orange, and HSPA6 and HSPA1B are labelled for the total proteome. Dashed lines represent the padj = 0.05 (black), $Log_2FC$ = 1 (red) or $Log_2FC$ = –1 (blue) thresholds used to classify DPs.

**e.** Direct comparison of $Log_2FCs$ upon heat-shock (x-axis) vs. proteasome inhibition (y-axis) for the A549 total (left) or polyUb–enriched (right) proteomes. HSPs are highlighted in orange vs. the rest of the proteome. For each pairwise comparison, linear regression lines for y ~ x, and squared Pearson's correlation coefficients, for HSPs vs. the rest of the proteome are shown. Dotted lines represent y = x, and dashed lines represent $Log_2FC$ = 1 (red) or $Log_2FC$ = –1 (blue).

**f.** Top, Total numbers of differential proteins (DPs) increased (up, red) or decreased (down, blue) between the pairwise comparisons indicated for each A549 total proteome experiment. Bottom, Volcano plots (plotted as in (d)) for each proteomics experiment, with DPs that increased (red) or decreased (blue) with the other proteotoxic stressor (i.e., in the other experiment) are indicated, and labelled only if they also passed the DP thresholds in the current experiment (regardless of direction). DPs from each state (proliferating or senescent) and stress (HS or Btz) were kept together, i.e., proliferating HS-modulated total proteome DPs were only plotted for the proliferating Btz total proteome data. Dashed lines represent adjusted p = 0.05 (black), $Log_2FC$ = 1 (red), or $Log_2FC$ = –1 (blue).

**g.** $Log_2FCs$ upon HS (left) or proteasome inhibition (right) across proliferating vs. senescent states for the total (top) or polyUb-enriched (bottom) proteomes. For each pairwise comparison, linear regression lines for y ~ x, and squared Pearson's correlation coefficients, are shown. Dotted lines represent y = x, and density of overlapping points are indicated with the colour scale.

**h.** Venn diagram showing overlap between proteins decreased (Down) or increased (Up) upon heat-shock (left) or proteasome inhibition (right) in the total (top) or polyUb-enriched (bottom) proteomes across the cellular states.

**I.** Left, Volcano plots showing $Log_2FC$ and $–Log_{10}$(padj)), both calculated by DEqMS, for each A549 total proteome experiment. Differential proteins (DPs) that increased (red) or decreased (blue) with the same stress in the other cell state (i.e., within the same TMT experiment) are indicated, and were labelled only if they also passed the p-value threshold in the current state (padj < 0.05), but in the opposite direction (regardless of whether they passed the additional threshold of absolute $Log_2FC$ >= 1). DPs from each state (proliferating or senescent) and stress (heat-shock or bortezomib) were kept together, i.e., proliferating heat-shock-modulated total proteome DPs were only plotted for the proliferating bortezomib total proteome data. Dashed lines represent padj = 0.05 (black), $Log_2FC$ = 1 (red) or $Log_2FC$ = -1 (blue).

**Supplementary Figure S7. Differential analysis of insoluble proteome reveals dampened RNA-related protein granule formation in the senescent state.**

**a.** Pairwise Pearson correlation coefficients of the $Log_2$-transformed normalised TMT abundances of the insoluble proteomes between all 18 samples.

**b.** Total numbers of differential proteins (DPs) increased (up, red) or decreased (down, blue) between the pairwise comparisons indicated for proliferating (P), quiescent (Q), or senescent (S) states under basal (Ba) or heat-shock (HS) condition in the insoluble proteome.

**c.** HS-induced $Log_2FC$ values across each cell state for the total (x-axis) vs. insoluble (y-axis) proteomes. For each pairwise comparison, linear regression lines for y ~ x, and squared Pearson's correlation coefficients, are shown. Density of overlapping points are indicated with the colour scale.

**d.** Heatmap of row-centred mean $Log_2$-transformed normalised TMT abundances for all DPs in the polyUb-proteome, based on the cut-offs described in B. Hierarchical clustering was performed using the complete agglomeration method with Euclidean distance measure for both columns (conditions) and rows (proteins). The row dendrograms were separated into 8 clusters.

**e.** Traces of the median-scaled $Log_2FCs$ for each DP across the six conditions in the insoluble proteome, separated by cluster. The mean of the median-scale $Log_2FCs$ for all proteins in the cluster per condition is overlayed as circles.

**f.** Top significant (adjusted $p < 0.05$, dashed line) annotation terms by over-representation analysis (ORA) for the proteins in each cluster in the insoluble proteome. ORA was performed using the gprofiler2 R package, with all 8,449 quantified in the IMR-90 total proteome as the custom background. Sources for annotation terms are indicated by the bar colours: CORUM, CORUM mammalian protein complex database; GO BP, CC, MF, Gene Ontology Biological Process, Cellular Compartment, Molecular Function; HP, Human Phenotype ontology; KEGG, gyoto Encyclopaedia of Genes and Genomes pathway; REAC, Reactome Pathway Database.

**g.** All significant (adjusted $p < 0.05$, dashed line) Reactome and KEGG pathway annotation terms by ORA (from (e)) for proteins in the HS-increased clusters from the insoluble proteome. Size of the circle depicts the number of DPs in each term. Terms related to specific biological features were manually annotated and coloured as shown in the legend for each cluster.

**h.** Comparison of proteins belonging to the 'RNA binding GOMF term (green) in the insoluble proteome suggests senescence-specific dampening at the insoluble proteome dimension vs. non-senescent states. Distribution of HS-induced $Log_2FCs$ in the insoluble proteomes, shown as violin + overlaid boxplots (i) or scatter-plots (j). Dashed lines represent $Log_2FC = 1$ (red) or –1 (blue), and grey horizontal line represents $Log_2FC = 0$.

**i.** HS-induced insoluble proteome accumulation of SG proteins is increased in the quiescent state. HS-induced alterations in the insoluble proteome for each state, with SG-annoted proteins highlighted based on classification by Hu et al. (2023).[4] $Log_2FCs$ for each group are shown as scatter-plots or violin and overlaid boxplots. Heatmap shows row-centred mean $Log_2$-transformed normalised TMT abundances in the insoluble proteome for all quantified SG proteins. Hierarchical clustering was performed using the complete agglomeration method with Euclidean distance measure for both columns (cell states) and rows (proteins).

For violin and overlaid boxplots in (h) and (i), adjusted p-values from non-parametric Kruskal-Wallis test (two-tailed, unpaired) are shown; horizontal grey, red, and blue horizontal lines represent $Log_2FC = 0, 1,$ and –1, respectively.

**j–k.** SG formation upon HS is conserved across cell states in IMR-90 fibroblasts and RPE-1 epithelia. IMR-90 (j) or RPE-1 (k) cells in proliferating, contact-inhibited quiescent, or DNA-damage-induced senescent states were heat-shocked (44 °C for 2 h, HS), or kept at 37 °C, followed by immediate fixation. Cells were permeabilised and immuno-stained with anti-G3BP1 (canonical SG marker) and counter-stained with DAPI, before imaging and automated quantification of cytoplasmic SGs (G3BP1 foci) and number of cells (DAPI objects). See also Fig. 3h.

**l.** STRING-database protein–protein interaction networks for the top 20 HS-increased insoluble proteome DPs in each cell state. The full STRING network (i.e., both functional and physical interactions) was included at medium confidence (minimum interaction score = 0.4). Fill colour of circles represents the degree of overlap for the protein in the top-20 lists. Line thickness represents the interaction confidence (as evaluated by the STRING database). See also 3l.

**Supplementary Figure S8. Nuclear RBPs TDP-43 and MATR3, but not FUS, form stress-induced nuclear foci that diverge in magnitude across non-proliferative states.**

IMR-90 fibroblasts or RPE-1 epithelia in proliferating, contact-inhibited quiescent, or DNA-damage-induced senescent states were heat-shocked (44 °C for 2 h, HS), or kept at 37 °C, followed by immediate fixation. Cells were permeabilised and immuno-stained with anti-TDP-43 (raised against either the N- or C- terminal region)(a), MATR3 (b), or FUS (c), and counter-stained with DAPI, before imaging and automated quantification of nuclear foci and number of cells (DAPI objects). See also Supplementary Fig. S8.

For (b) and (c), bar-charts on the left show normalised RPM (transcriptome) or TMT abundances (total, polyUb, and insoluble proteomes) for each protein, with adjusted p-values from DESeq2 (transcriptome) or DEqMS (proteomes).

For MATR3, additional bar-chart shows mean nuclear foci per cell after adjusting for differences in nuclear area (as calculated by the mean nuclear mask area per state and condition).

**Supplementary Figure S9. Misfolded but not well-folded proteins are secreted upon stress in senescent states, across cell types and senescence-induction models.**

**a.** IMR-90 fibroblasts, or A549 or RPE-1 epithelia, in proliferating, contact-inhibited quiescent, or DNA-damage/oxidative-stress-induced senescent states, were heat-shocked (44 °C for 2 h, HS), or kept at 37 °C, in fresh complete growth medium, followed by collection of the CM, centrifugation and passing through a 0.2 μm filter to remove intact cells and debris, and analysis by TDP-43 sandwich ELISA as a readout, according to manufacturer's instructions. See also Fig. 4c.

**b.** Well-folded YFP does not accumulate in the CM of senescent A549 epithelia upon HS. A549 cells transiently-transfected with a plasmid for expression of YFP were established in proliferating or DNA-damage-induced senescent states, and heat-shocked for 2 h. CM collection and quantification was performed as in (a), except with the GFP ELISA as a readout.

**c.** Densitometry-based quantification of the TDP-43 immunoblot shown in Fig. 4f. Bands were quantified using a LiCor Odyssey and Image Studio.

**Supplementary Figure S10. Peri-centriolar satellite proteins shown to be important for aggresome formation are consistently declined in the senescent state.**

Multi-omics profile for the five proteins whose knockdown was shown to impair aggresome formation.[5] Bar-charts show normalised RPM (transcriptome) or TMT abundances (total, polyUb, and insoluble proteomes) for each protein, with adjusted p-values from DESeq2 (transcriptome) or DEqMS (proteomes).

**Supplementary Figure S11. Aggresome formation is impaired in the senescent state.**

**a.** Imaging of aggresomes formed following proteasome inhibition with bortezomib (Btz) or MG-132, or DMSO-vehicle control (Vehicle), for 6 h or 24 h, in A549 human lung adenocarcinoma cells in various cell states. At the end of 24 h treatment, cells were fixed, permeabilised, and immuno-stained with anti–Ub[K48], and counter-stained with DAPI, before imaging immediately on an ImageXpress Confocal HT.ai High-Content Imaging System. White arrowheads depict putative aggresomes.

**b–c.** Dose-response of aggresome formation in A549 (b) or IMR-90 (c) cells in different states, upon proteasome inhibition with bortezomib or MG-132, following treatment and processing of cells as described in (a).

**Supplementary Figure S12. A549 cells form aggresomes containing misfolded proteins in proliferating but not senescent states.**

**a.** Imaging of aggresomes formed following proteasome inhibition with bortezomib (Btz) or DMSO-vehicle control (Vehicle), for 24 h, in A549 human lung adenocarcinoma cells in proliferating or DNA-damage-induced senescent cell states. At the end of 24 h treatment, cells were fixed, permeabilised, and immuno-stained with anti–Ub[K48] and anti-vimentin, and counter-stained with DAPI, before imaging immediately on an ImageXpress Confocal HT.ai High-Content Imaging System.

**b.** GFP–Htt-exon1[Q97], but not GFP–Htt-exon1[Q25], accumulates at the aggresome in proliferating A549 cells. Cells were treated and processed as described in (a).

**Supplementary Figure S13. Limited molecular chaperone alterations in senescence that are not shared with quiescence.**

**a.** Heatmap of row-centred mean $Log_2$-transformed normalised TMT abundances for all chaperones in the total proteome, with classifications of chaperones according to the Proteostasis Consortium annotation.[3] Hierarchical clustering was performed using the complete agglomeration method with Euclidean distance measure for both columns (conditions) and rows (proteins).

**b.** STRING-database protein–protein interaction networks for the chaperones statistically lower (left) or higher (right) in abundance in senescent vs. proliferating states.

**c.** STRING-database protein–protein interaction networks for the chaperones statistically lower (left) or higher (right) in abundance in senescent vs. proliferating and quiescent states. The full STRING network (i.e., both functional and physical interactions) was included at medium confidence (minimum interaction score = 0.4). Fill colour of circles represents specific annotation terms. Line thickness represents the interaction confidence (as evaluated by the STRING database).

**Supplementary Figure S14. DNACJ5 is not important for senescence induction trajectories, but does impact misfolded protein secretion in a manner that can be rescued by chaperone induction.**

**a.** Changes to SA-bG, EdU, and nuclear size are conserved regardless of DNAJC5 levels in A549 cells. Assay were performed as described in Supplementary Fig. S1.

**b.** Densitometry-based quantification of the DNAJC5 immunoblot shown in Fig. 5l. Bands were quantified using a LiCor Odyssey and Image Studio.

**c.** Transient knockdown of DNAJC5 significantly reduces extracellular TDP-43 levels in senescent A549 cells upon heat-shock. A549 cells were made senescent by DNA-damage for 7 days, followed by lipid-based transfected with siRNA targeting DNAJC5, or non-targetting control siRNA, for three days. Subsequent HS, CM collection, and TDP-43 quantification was performed as in Fig. 5o.

**d.** Heterologous over-expression of HSPA1 (HSP72) or HSP90AB1 (HSP90-beta) significantly reduces extracellular GFP–Htt-exon1[Q97] levels in senescent A549 cells upon heat-shock. A549 cells were made senescent by DNA-damage for 7 days, followed by lipid-based transfected with pCMV::mScarlet–HSP72, pCMV::Flag–HA–HSP90AB1, or pCMV::HaloTag for three days. Subsequent HS, CM collection, and GFP quantification were performed as in Fig. 5o.

**e.** Pharmacologic potentiation of the heat-shock-response by BGP-15 significantly reduces extracellular TDP-43 release upon HS in senescent A549 epithelia. A549 cells were made senescent by DNA-damage for 10 days, followed by acute HS as described in Fig. 5m, +/- 10 µM BGP-15 (added 30 min before HS). CM collection and TDP-43 quantification was performed as in Fig. 5m.

#### Supplementary Methods

##### Senescence verification by EdU and senescence-associated beta-galactosidase (SA-bG) staining.

Cells were seeded into 6 or 12-well TC-treated plates and allowed to achieve their desired cell states *in situ*. Cells were then pulsed with 10 μM EdU for 24 h, and fixed in 4 % formaldehyde (Thermo, Cat# 28908) for 15 min. Cells were incubated in SA-β-galactosidase staining solution (Table 1),[6] adjusted to pH 6.0 (except for the positive control, which was at pH 4.0). Samples were wrapped tightly in plastic to avoid evaporation and incubated in a non-$CO_2$-controlled incubator overnight at 37 °C. Note that the assay relies on residual SA-bG activity present in the short window post fixation, and thus staining must be carried out immediately following fixation. The next day, cells were washed 3x with ice-cold PBS and stored at 4 °C until ready for EdU fluorophore conjugation. Cells were permeabilised with 0.1 % Triton X-100 in ice-cold PBS for 5 min, washed 3x in ice-cold PBS, then incubated in fresh EdU fluorophore conjugation solution (Table 2) for 30 min at room temperature in the dark. Cells were further stained with 0.5 μg/mL DAPI for 10 min at room temperature in the dark. Samples were imaged on a wide-field fluorescence microscope (Nikon Ti2) and analysed for senescence indicators: cell-cycle arrest (percentage of EdU+ nuclei as a percentage of the total DAPI count), increase in nuclear size (DAPI mask area), and SA-β-gal-positive staining compared to control cells.

| Reagent | Stock concentration | Final concentration |
|---|---|---|
| X-gal solution | 20 mg/mL | 1 mg/mL, always made fresh |
| Citric acid/sodium phosphate buffer* | 0.2 M | 40 mM (pH 6.0) |
| Potassium ferricyanide | 100 mM | 5 mM |
| Potassium ferrocyanide | 100 mM | 5 mM |
| NaCl | 1 M | 150 mM |
| $MgCl_2$ | 100 mM | 2 mM |
| PBS pH 7.4 | - | - |

*Citric acid/sodium phosphate buffer (0.2 M, pH 6.0): mix 36.85 mL of 0.1 M citric acid solution with 63.15 mL of 0.2 M sodium phosphate ($Na_2HPO_4$) solution. Verify that the pH is 6.0.

**Table 1: SA-bG activity staining solution**

| Reagent | Stock concentration | Final concentration |
|---|---|---|
| PBS pH 7.4 | - | - |
| Sulfo-Cy5-Azide | 4 mM | 8 µM |
| $CuSO_4.5H_2O$ | 200 mM | 2 mM |
| Ascorbic Acid | 1.13 M | 113 mM |

**Table 2: EdU–fluorophore conjugation solution**

**Cloning of plasmids.**

To construct pLenti CMV Puro tdTomato–NLS, the pENTR plasmid pME tdTomato–NLS (gift from David Tobin, Addgene plasmid #135205) was inserted into the lentiviral Gateway destination vector pLenti CMV Puro DEST (w118-1) (gift from Eric Campeau & Paul Kaufman, Addgene plasmid #17452) using the LR reaction (Thermo Fisher Scientific, Cat# 11791100), according to manufacturer's protocols.

To construct pLVX 6xHSE::mScarlet–NLS, the 6xHSE artificial promoter fragment from pMNLucPAUM 6HSE (gift from Thomas Czerny, FH Campus Wien, Austria)[7] was double-digested with NheI-HF + StyI, and ligated into NheI-HF + BspHI double-digested pLVX–ATF4 mScarlet–NLS (gift from David Andrews, Addgene #115969).

To construct pET28a HaloTag–[tev]–trTUBE$_6$, the HaloTag-[tev]-[MCS] fragment from pET28a-HaloTag-[tev]-[MCS] (DU23222, MRC-PPU Reagents & Services) was inserted into the optimised pET28a T7pCONS TIR-2 sfGFP (gift from Daniel Daley, Addgene #154464)[8] backbone through PCR amplification of the HaloTag-[tev]-[MCS] fragment with custom primers 108-BlpI-F1 and pET28a-BlpI-R1, followed by BlpI digests and ligation to yield pET28aOPT-HaloTag-[tev]-[MCS]. The 6xTR-TUBE insert from pRSET-6xTR-TUBE (gift from Yasushi Saeki, Addgene #110313)[9] was double-digested with EcoRV + DraIII and ligated into Eco53kI + DraIII double-digested pET28aOPT-HaloTag–[tev]–[MCS] to generate the final pET28a HaloTag–[tev]–trTUBE$_6$ expression plasmid.

To construct the doxycycline-inducible reporters pCW57.1 EGFP–Htt-exon1[polyQ97] and pCW57.1 EGFP–Htt-exon1[polyQ25], Gateway cloning was used. PCR fragments encoding Htt-exon1[Q97] and Htt-exon1[Q25] were amplified from pcDNA3 Htt-exon1[Q25]–mTurboID–HA, or pcDNA3 Htt-exon1[Q97]–mTurboID–HA (gifts from Judith Frydman, Stanford University, CA, USA) using custom primers containing restriction sites for

BglII and NheI (Htt_For-BglII and Htt_Rev-STOP-NheI), and ligated after BglII + NheI-HF double-digestion into similarly digested pENTREGFP2 (gift from Nathan Lawson, Addgene plasmid #22450) purified from *dam-/dcm-* competent *E. coli*. The resultant pENTR plasmids were inserted into the lentiviral Gateway destination vector pCW57.1 (gift from David Root, Addgene plasmid #41393) using the LR reaction according to manufacturer's protocols.
All plasmids were verified by whole-plasmid sequencing (Plasmidsaurus, NJ USA).

**Lentivirus preparation.**

For each lentivirus to be generated, 600,000 early-passage HEK293T cells were seeded into a well of a 6-well plate, and allowed to attach overnight under regular culture conditions. The following day, 0.2 pmol of endotoxin-free DNA of the desired lentiviral transfer vector was mixed with 0.27 pmol packaging (psPAX2, gift from Didier Trono, Addgene plasmid #12260) and 0.07 pmol envelope (pMD2.G, gift from Didier Trono, Addgene plasmid #12259) plasmids, in 150 µL Opti-MEM I Reduced Serum Medium (Gibco, Cat# 31985070), and mixed with another solution containing 7 µL Lipofectamine-2000 in 150 µL Opti-MEM before incubating at room temperature for 10 min, to allow DNA-enclosed liposomes to form. The liposome mixture was added drop-wise onto the HEK293T cells, and incubated for 24 h under regular culture conditions, before replacing the transfection media with 3 mL complete DMEM. After another 72 h under regular culture conditions, the lentivirus-containing media was collected, centrifuged at 1,000 *g* for 5 min to pellet any detached HEK293T cells and larger debris, and passed through a sterile Whatman Puradisc-25 0.45 *µ*m PES syringe filter (Sigma-Aldrich, Cat# WHA67802504) to remove smaller debris. This lentiviral supernatant was either used for cell transduction immediately, or stored at 4 °C for a maximum of 7 days.

**Lentiviral transduction and selection for stable transgenic cell lines.**

To generate stable transgenic cell lines, cells to be transduced were seeded into 24-well plates in their complete growth media and allowed to attach overnight under regular culture conditions. The following day, media in each well was replaced with a 1:1 mixture of complete growth media and filtered lentiviral supernatant with 2.5 µg/mL polybrene, and incubated for 48 h, before replacing the lentivirus-containing media with complete growth media. After another 48 h (i.e., 96 h post-transduction), fluorescence of the GFP-positive control (pLenti eGFP Puro (w159-1), gift from Eric Campeau and Paul Kaufman, Addgene plasmid #17481) was assessed using an EVOS FL AMF4300 Cell Imaging System.

For the tdTomato–NLS and heat-shock-response (HSR) reporter IMR-90 cell lines, the transduced cell population was selected in bulk by flow sorting, based on baseline red fluorescence, on a FACSAria Fusion (BD Biosciences, UK). For inducible Htt-exon1[Q25] and Htt-exon1[Q97] A549 cell lines, transduced cells were cultured in complete media supplemented with puromycin (2 µg/mL) for 21 days, to select for successfully-transduced cell populations.

**Generation of HaloTag-[tev]-trTUBE$_6$ magnetic beads.**

Rosetta 2(DE3) Competent *E. coli* transformed with the pET28a-HaloTag—[tev]—trTUBE$_6$ plasmid were cultured at 30 °C for 18 h to $OD_{600}$ ~0.7, induced with 1 mM IPTG, and incubated for 24 h at 16 °C. Cells were pelleted at 7,000 *g* for 10 min at 4 °C and lysed with three pellet volumes (w/v) bacterial lysis buffer (50 mM Tris-HCl, pH 7.5, 150 mM NaCl, 1% Triton X-100, 20 mM TCEP, 1x cOmplete EDTA-free Protease Inhibitor Cocktail, 0.2 mM PMSF), disrupted by sonication on ice for 2 min total at 20 W (10 s intervals) using non-microtip probe (Covaris), and cleared by centrifugation (30 min, 16,000 *g*, 4 °C). 200 µL of the resultant cleared lysate was used to saturate 1 mL of 20 % Magne HaloTag Beads (Promega Cat# G7281). Conjugated beads were washed 3x with 1 mL IP buffer (50 mM Tris-HCl, pH 7.5, 150 mM NaCl, 1% IGEPAL CA-630) and incubated for 2 h at room temperature with end-over-end rotation at 20 rpm. Beads were subsequently washed 3x with IP wash buffer, and stored at 4 °C in a 1:10 bead:IP buffer ratio for up to 6 months.

**Fluorophore-conjugation of Ub[K48] nanobody.**

To label the Ub[K48] single-domain antibody (sdAb), lyophilised sdAb anti-Ubiquitin K48 (NanoTag Biotechnologies, Germany; Cat# N1805) was reconstituted in 100 mM Tris-HCl pH 8.0 (made up in molecular biology-grade water) and immediately incubated with a 5-fold molar excess of Janelia Fluor 549 maleimide (Tocris Biosciences; Cat# 6500) or Janelia Fluor 646 maleimide (Tocris Biosciences; Cat# 6590) that had been freshly reconstituted in anhydrous DMSO (Molecular Probes, Cat# D12345). The reaction tube was flushed with argon to minimise cysteine oxidation, and incubated on ice for 2 h in the dark, after which free unlabelled fluorophore was removed by passing the reaction mixture through a Zeba Spin Desalting Column (Thermo Fisher Scientific; Cat# 89882), according to manufacturer's instructions. The purified conjugated nanobody was aliquoted and stored at –70 °C. Dilution factor was newly ascertained for each batch, but 1:500 was commonly found to provide the optimal signal:noise ratio.

**Image processing, quantification, and display.**

Images obtained in proprietary file formats (e.g., Nikon .nd2; Olympus .oif) were converted to .tiff files in ImageJ via the Bio-Formats importer. For display (e.g., in figure panels), tiff files were batch processed to equalise look-up tables (LUTs) between images to be compared, cropped to the desired dimensions, fluorescence channels separated, converted from 16-bit to 8-bit, inlaid with scale-bars, and saved as png files.

For aggresome quantification, all files were acquired using the same settings on the Molecular Devices HT.ai high-content imaging system, and processed using a custom pipeline on CellProfiler (v3.1.9) (Table 3). This pipeline was developed to be relatively flexible across cell types, e.g., the same modules were used to identify aggresomes in fibroblasts and epithelial cells in our study.

| **Module** | **Settings** | **Function** |
|---|---|---|
| Identify Primary Objects | Global thresholding using minimum cross-entropy of DAPI stain; smoothening scale 1.3488; based on object size 50–200; discard objects touching border | Segments nuclei. |
| Identify Secondary Objects | Propagation using nuclei; Global thresholding using minimum cross-entropy; | Segments cells using nuclei as seed, via cytoplasmic stain (e.g., vimentin). |
| Measure Object Size Shape | Nuclei; no advanced features | Measures size and shape of nuclei. |
| Filter Objects | Nuclei; filter on limits -> area shape -> form factor; minimum value 0.3 | Discards any nuclei which are not round enough, e.g., apoptotic blebbing. |
| Relate Objects | Set parents (Nuclei) and children (Cells) – Save as new set | Only keeps cells which have a matching live nucleus. |

| | | |
|---|---|---|
| Measure Object Size Shape | Cells; no advanced features | Measures size and shape of cells. |
| Filter Objects | Cells; filter on limits -> area shape -> form factor; maximum value 0.4 | Discards any cells which are ‘too round’, e.g., cells that are dead or detached. |
| Measure Image Intensity | Select post-filtered cells | Measures fluorescence intensity of each channel within selected objects. |
| Identify Primary Objects | Object size 20–80; global thresholding using mean image intensity of Ub[K48] stain (note that Ub[K48] signal must be strong, otherwise set manually); correction factor 1.5; lower threshold bound 0.007; distinguish clumped objects by shape | Finds Ub[K48]+ puncta large enough to be considered an aggresome. |
| Identify Secondary Objects | Set input image as DAPI stain; filtered nucleus as object, use distance – N to identify secondary object, 50 pixels by which to expand the primary object | Draws a juxtanuclear ring (JR). |
| Relate Object | Set parents (JR) and children (K48 aggresomes) – Save as new set | Only keeps Ub[K48]+ aggresomes inside the JR. |
| Split or Merge Objects | Set K48 aggresomes as the input, use merge operation per-parent with the JR as the parent object | Merges any aggresomes found within JR. Pipeline occasionally labels one aggresome as two smaller side-by-side aggresomes; this step merges them back together. |

| | | |
|---|---|---|
| Identify Primary Objects | Object size 20–100; global thresholding using mean image intensity of the vimentin stain (note signal must be strong, otherwise set manually); correction factor 1.3; lower threshold bound 0.15; distinguish clumped objects by shape; never fill holes in identify objects | Identifies vimentin cages |
| Relate Objects | Set parents (Cell) and children (vimentin cage) – Save as new set | Only keeps vimentin cages found within a cell mask. |
| Identify Secondary Objects | Set input image as vimentin stain; input object K48 aggresomes; use distance – N to identify secondary object, 50 pixels by which to expand the primary object | Draws a ring around Ub[K48]+ aggresomes. |
| Relate Objects | Set parents (Ub[K48] aggresome ring) and children (vimentin cage) – Save as new set | Keeps only vimentin cages if found near an aggresome (module needed to de-noise vimentin signal). |
| Measure Object Intensity | Select objects to measure | Measures fluorescence intensity of each channel within selected objects. |
| Enhance or Supress Features | Enhance Polyubiquitin-K63 signal; feature type speckles at size 10 | Finds Ub[K63]+ aggregates by enhancing the Ub[K63] image. |
| Identify Primary Objects | Object size 2–20; Global thresholding using minimum cross-entropy; smoothening scale 1.3488; distinguish clumped objects by intensity | Identifies Ub[K63]+ aggregates. |

| | | |
|---|---|---|
| Relate Objects | Set parents (Cell) and children (Ub[K63] aggregate) – Save as new set | Filters out Ub[K63]+ aggregates not within a cell mask. |
| Gray to Colour | Select images to be pseudo-coloured in RGB, relative weight for each 1.0 | Pseudo-colours each channel.<br><br>Sanity-check for pipeline; removed when running analysis. |
| Overlay Outlines | Overlay onto RGB image, select desired objects and their preferred outline colour | Draws an outline around each nucleus, cell, aggresome, and vimentin cage. Allows visual confirmation that the pipeline is accurately segmenting images.<br><br>Sanity-check for pipeline; removed when running analysis |
| Export to Spreadsheet | Select features to export; add image file and folder names to data file; calculate per-image mean values for object measurements | Exports all data into excel format. |

**Table 3: 23-step CellProfiler pipeline to segment nuclei, cells, aggresomes, aggregates, and vimentin cages**.
